# A small molecule inhibitor of NVL suppresses tumor growth by blocking ribosome biogenesis

**DOI:** 10.1101/2025.07.31.667081

**Authors:** Holly H. Guo, Ye Tao, Victor E. Cruz, Min Fang, Vishal Khivansara, Shanhai Xie, Ashley Leach, Divya Reddy, Johann Peterson, Jiwoong Kim, Noelle S. Williams, Arin Aurora, Jan P. Erzberger, Jef K. De Brabander, Deepak Nijhawan

## Abstract

A longstanding hypothesis, stemming from the enlarged nucleoli typical of cancer cells, posits ribosome production as a selective cancer liability. Certain genotoxic chemotherapies work partly by disrupting ribosome biogenesis, highlighting the need for selective inhibitors of this pathway. Using forward genetics, we identified mutations in the essential 60S ribosomal subunit assembly factor NVL that confer resistance to MM17, a dibenzothiazepinone with anticancer activity. Cryo-EM reconstructions of the NVL hexameric assembly reveal two MM17 docking sites adjacent to resistance mutations. NVL inhibition by MM17 arrests 60S biogenesis in the nucleolus and induces cell cycle arrest or apoptosis through both MDM2/p53-dependent and p53-independent pathways, without causing DNA damage. A bioavailable analog, MM927, suppresses tumor growth in mouse models of leukemia and colorectal cancer without observable toxicity. These findings establish NVL inhibitors as a promising new class of targeted therapeutics and validate ribosome biogenesis as a cancer-specific vulnerability.

## INTRODUCTION

Our ability to evaluate many cancer-relevant proteins as potential drug targets is limited by a general lack of specific inhibitors, particularly for protein families with little precedent for engagement by drug-like molecules. To circumvent this, cell-based phenotypic screens for anticancer compounds offer an unbiased approach to identify protein targets for which there is no prior example of small molecule modulation (*1*). We and others have leveraged this strategy by combining phenotypic approaches with unbiased forward genetics, biochemical reconstitution, and structural biology to uncover druggable cancer targets in cellular pathways that are poorly characterized or underexplored (*2*).

For example, cancer cells ramp up ribosome synthesis to meet the elevated protein production demands of uncontrolled proliferation, making the ribosome biogenesis pathway a compelling drug target (*3, 4*). Interestingly, the cancer drugs 5-fluorouracil (5-FU) and oxaliplatin have recently been found to interfere with ribosome biogenesis, despite being classified as DNA-damaging agents (*5, 6*). Specifically, 5-FU incorporation into rRNA disrupts 47S rRNA nucleolytic processing (*7, 8*), while oxaliplatin has been proposed to inhibit ribosome production by disrupting nucleolar condensate formation (*6, 7, 9, 10*).

These findings underscore the need for specific inhibitors to definitively determine whether ribosome biogenesis is a valid cancer target. With over 200 known assembly factors involved in this process, candidate targets include polymerases, rRNA processing and modifying enzymes, structural proteins that template assembly steps, and various ATPases that remodel rRNA or remove biogenesis factors at critical junctions to guide ribosome maturation (*11–13*). In ongoing efforts to inhibit ribosome biogenesis, several compounds have been nominated as RNA polymerase I inhibitors (*14–18*) including CX-5461 (*19*), which has advanced to Phase II clinical trials in cancer patients (Table S1) (*20, 21*). In all of these cases, however, RNA polymerase I has not been validated as the functional target, and recent studies suggest that the anticancer activity of CX-5461 is actually the result of DNA damage caused by topoisomerase TOP2B inhibition (*22–25*). Although a few other compounds have been proposed to disrupt mammalian ribosome assembly (*26–28*), none have a defined molecular target or a well-characterized mode of action (Table S1) (*29*).

AAA+ ATPase enzymes catalyze critical steps in ribosome biogenesis by coupling ATP hydrolysis to the mechanical removal of specific biogenesis factors and are therefore attractive pharmacological targets. Assembly of the large ribosomal subunit (60S) is mediated by several AAA+ ATPases including NVL, MDN1 and SPATA5/5L1 (*12, 30–35*). Small molecules that disrupt AAA+ ATPase function during 60S biogenesis have been developed for Drg1, the *S.cerevisiae* homolog of SPATA5 (*36, 37*) and Mdn1, the *S.pombe* homolog of MDN1 (*38*). However, no small-molecule inhibitors have been reported for any of the mammalian AAA+ orthologs (Table S1).

In this study, we identified the human AAA+ ATPase NVL as the target of a previously undescribed compound, MM17, that exhibits anticancer activity. A cryo-EM structure of NVL bound to MM17 reveals two analogous binding sites within the hexameric assembly. Resistance-conferring mutations in NVL cluster around these sites and reduce compound binding. The NVL-MM17 interaction specifically arrests 60S subunit maturation in the nucleolus. NVL inhibition stabilizes p53 through MDM2 and leads to cell cycle arrest or apoptosis, depending on the type of cell, without causing DNA damage. In the absence of p53, MM17 induces a more delayed cell-cycle arrest via an alternate pathway. MM927 is a potent and bioavailable analog of MM17 that suppresses tumor growth in two different cancer models with no evidence of overt toxicity. Together, our findings demonstrate that ribosome biogenesis can be selectively targeted in cancer and identifies MM17 as a lead molecule for the development of a new class of anticancer therapeutics.

## RESULTS

### NVL is the target of MM17

We have an ongoing research effort to use unbiased forward genetics to identify the targets of novel small molecules emerging from high-throughput screens that reduce the viability of cancer cells. We previously reported 53 compounds from a high-throughput screen of 99,599 small molecules that inhibited proliferation of the colorectal cancer cell line HCT116 (*2*). Among these, the dibenzothiazepinone MM17 inhibited HCT116 growth in a dose-dependent manner (IC50 = 0.46 µM) (Fig. 1A, B). We have previously derived a cell line, iHCT116, in which addition of indole acetic acid (IAA) triggers ubiquitin-mediated degradation of the DNA mismatch repair protein MLH1, transiently increasing mutation rates (*2*). Barcoded iHCT116 cells, initially cultured either with (Mutagenesis-on) or without IAA (Mutagenesis-off), were exposed to escalating doses of MM17. The ratio of surviving clones between the Mutagenesis-on to -off conditions increased in a dose-dependent manner, suggesting a genetic basis for resistance (Fig. 1C).

**Figure 1.**
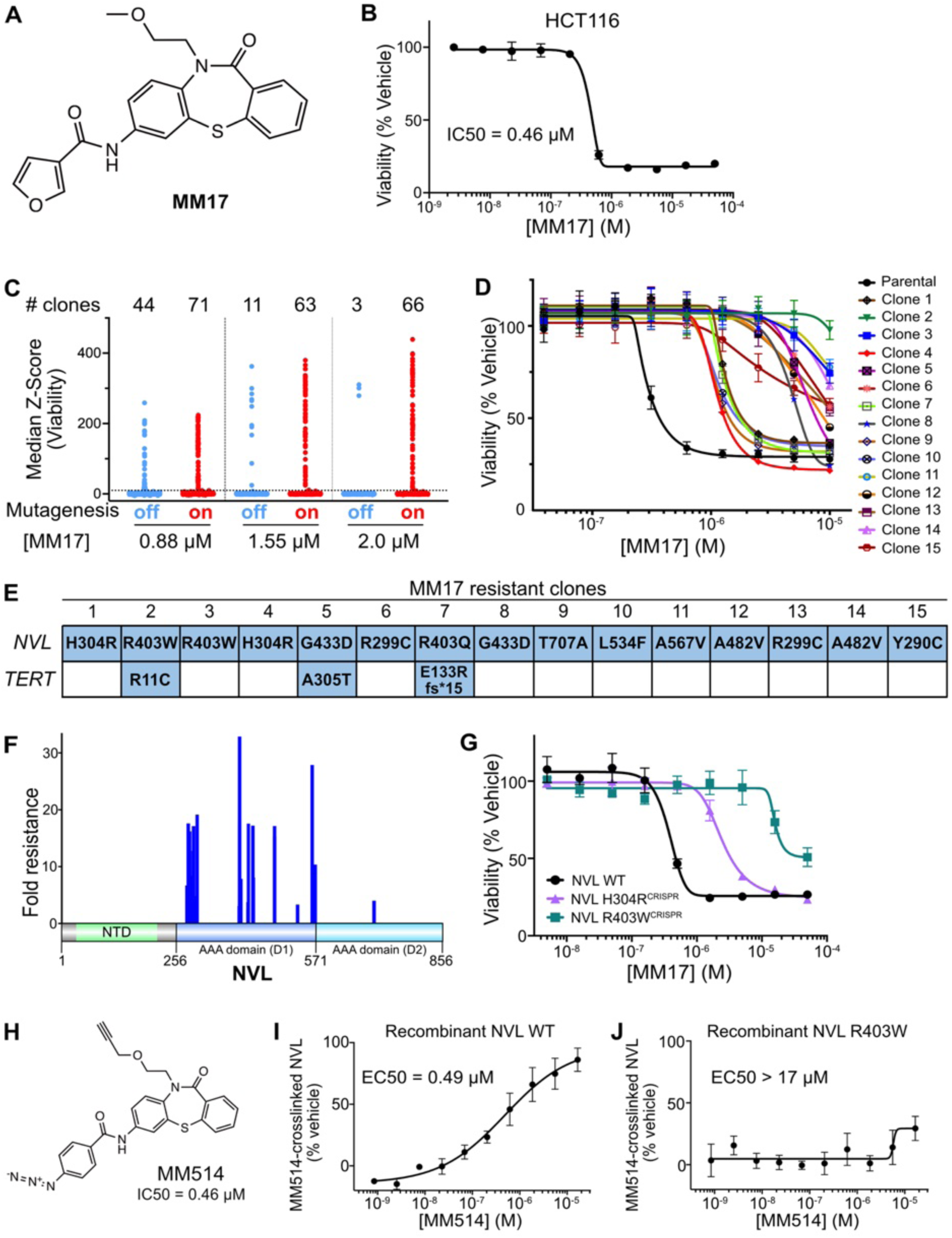
NVL is the target of MM17. **A.** Chemical structure of MM17. **B.** Viability of iHCT116 cells treated with MM17 for 72 hours (n = 2 biological replicates, mean ± sem). **C.** Cell viability after selection with MM17 in a forward genetics screen of iHCT116 cells without (Mutagenesis-off) or with (Mutagenesis-on) IAA treatment performed according to recently published procedures (*2*). **D.** Viability assay in parental iHCT116 cells and 15 independent resistant clones treated with MM17 for 72 hours (n = 2 biological replicates, mean ± sem). **E.** Genes mutated in at least 3 out of 15 resistant clones analyzed by whole exome sequencing. **F.** NVL mutations found in 57 out of 59 unique clones mapped onto a schematic of the NVL domain structure. **G.** Viability assay of HCT116 NVL WT, NVL H304R^CRISPR^, and NVL R403W^CRISPR^ cells treated with MM17 for 72 hours (n = 3 biological replicates, mean ± sem). **H.** Chemical structure of MM514 and its viability inhibition IC50 in HCT116. **I, J.** Quantitation of MM514-crosslinked recombinant NVL WT (**I**) or NVL R403W (**J**). (n = 3 biological replicates, mean ± sem).

We identified 59 clones with unique barcode sequences (Table S2), consistent with a distinct founder event for each clone. The MM17 IC50 values for these clones ranged from 2.45 to 32.89-fold higher than that of the parental cell line, supporting acquired resistance and indicating that the clones likely harbored a diverse set of mutations. We therefore performed whole-exome sequencing on 15 representative clones spanning the full resistance spectrum (Fig. 1D). Only two genes contained variants in 3 or more of the selected clones: *TERT* was mutated in 3 clones, while *NVL* was mutated in all 15 (Fig. 1E). These results implicate *NVL* as the gene most likely associated with MM17 resistance. We sequenced *NVL* exons in the remaining clones using PCR-based methods, focusing on regions mutated in the exome sequencing data. In total, 57 of 59 clones harbored mutations in *NVL.* NVL is a hexameric type II AAA+ protein implicated in ribosome biogenesis, with each subunit containing two ATPase domains (D1 and D2). The majority of mutations clustered within the D1 domain (Fig. 1F), with several amino acid positions mutated to different residues across independent clones (Fig. S1A and Table S2).

To directly test whether the spontaneous *NVL* mutations confer MM17 resistance, we used CRISPR/Cas9 (clustered regularly interspaced short palindromic repeats) to introduce focal edits in two frequently mutated regions of NVL (residues 298-304 and 402-406; n=8 for each region) (Figure. S1A). CRISPR-induced double-strand breaks can result in random mutations via non-homologous end-joining (NHEJ) or, in the presence of a template, specific edits via homology-directed repair (HDR) (*39*). Expression of Cas9 with sgRNA targeting two of these regions resulted in a higher frequency of MM17-resistant colonies than a mock sgRNA control (Fig. S1B-D). Co-delivery of repair templates encoding the most frequent substitutions in each region, H304R and R403W, further increased resistance, demonstrating that these specific mutations are sufficient to confer MM17 resistance. The most frequent missense variants in each population were the designed alleles H304R and R403W, consistent with efficient HDR, although we also observed frequent frameshift and in-frame mutations consistent with NHEJ (Fig. S1E). The prevalence of frameshift mutations suggests that resistant cells have silenced one allele and are likely heterozygous for H304R or R403W suggesting that these amino acid substitutions preserve the essential AAA+ translocase function of NVL. These populations (hereafter referred to as NVL H304R^CRISPR^ and NVL R403W^CRISPR^) grew similarly to parental cells (Fig. S1F), were more resistant to MM17 compared to NVL WT (IC50: NVL WT = 0.38 µM; NVL H304R^CRISPR^ = 2.4 µM; NVL R403W^CRISPR^ > 50 µM) (Fig. 1G), and remained equally sensitive to the unrelated cytotoxin paclitaxel (Fig. S1G). These results demonstrate that focal editing in *NVL* resistance hotspots, particularly H304R and R403W, is sufficient to confer MM17 resistance, and furthermore, support the conclusion that the spontaneous *NVL* mutations identified in iHCT116 confer MM17 resistance.

NVL mutations may confer MM17 resistance by affecting a downstream pathway or alternatively by preventing compound binding. To test whether MM17 binds directly to NVL in cells, we synthesized MM514, a photoactivatable analog that forms covalent adducts with bound proteins upon UV exposure and contains an alkyne for azide-dye conjugation via click chemistry (Fig. 1H). Importantly, MM514 inhibited HCT116 viability (IC50 = 0.46 µM) with potency comparable to MM17, and NVL R403W^CRISPR^ cells were resistant to MM514, suggesting a similar mechanism of action (Fig. S2A). Recombinant NVL-SNAP purified from cells treated with MM514 and UV light was covalently bound to MM514, providing evidence of direct target binding (Fig. S2B-E). To determine whether this interaction is functionally relevant, we performed probe competition assays using two structurally related MM17 derivatives with differing potencies (Fig. S2F, G). A single methoxy group shift from the para (MM524) to the ortho (MM691) position increased potency by 28-fold (IC50: 0.43 µM vs 12 µM).

Correspondingly, MM524 more effectively displaced MM514 crosslinking (EC50 = 2.6 µM) than MM691 (EC50 = 26 µM) (Fig. S2H, I). The correlation between structure, NVL binding, and cellular activity indicates that MM514 crosslinking to NVL is both specific and related to its anticancer activity. We next purified wild type and R403W-mutated NVL-SNAP from UV-treated cells exposed to increasing concentrations of MM514. MM514 crosslinked to wild-type NVL-SNAP (EC50 = 0.49 µM) more efficiently than the R403W variant (EC50 > 17 µM), suggesting that the resistance mutation impairs compound binding (Fig. 1I, J, S2J, K). Taken together with the probe competition data, these findings show that NVL is the direct target of MM17 and related derivatives.

### Cryo-EM reconstruction of the NVL/MM17 complex

To investigate the molecular basis of the MM17-NVL interaction, we determined the cryo-EM structure of NVL in complex with MM17. Expression and purification of the D1/D2 core of wild-type NVL yielded low amounts of aggregated protein, and cryo-EM analysis revealed that it assembles into long, filament-like spirals rather than the expected hexamers (Fig. S3A). Double mutations in the Walker B motif of the *C. thermophilum* NVL homolog Rix7 have previously been shown to stabilize the hexameric conformation (*40*). Applying the same strategy to NVL, recombinant expression and purification of NVL^E366Q/E683Q^ (NVL^dEQ^) yielded stable, monodisperse hexamers (Fig. S3B, C), with hexamer stoichiometry unaffected by MM17 addition, as measured by mass photometry (Fig. S3D).

A single particle cryo-EM reconstruction of NVL^dEQ^ in complex with MM17 produced a map with an overall resolution of 3.05 Å (Fig. S4 and Table S2). As observed in reconstructions of *C. thermophilum* Rix7, *H. sapiens* P97 (Fig. S5A) and *S. cerevisiae* Drg1, the NVL hexamer adopts a staircase configuration, with pore loops from five D1 and D2 AAA+ modules engaging an extended peptide within the central channel (Figs. 2A, S3E and S5B,C) (*32*) (*41–43*).

**Figure 2.**
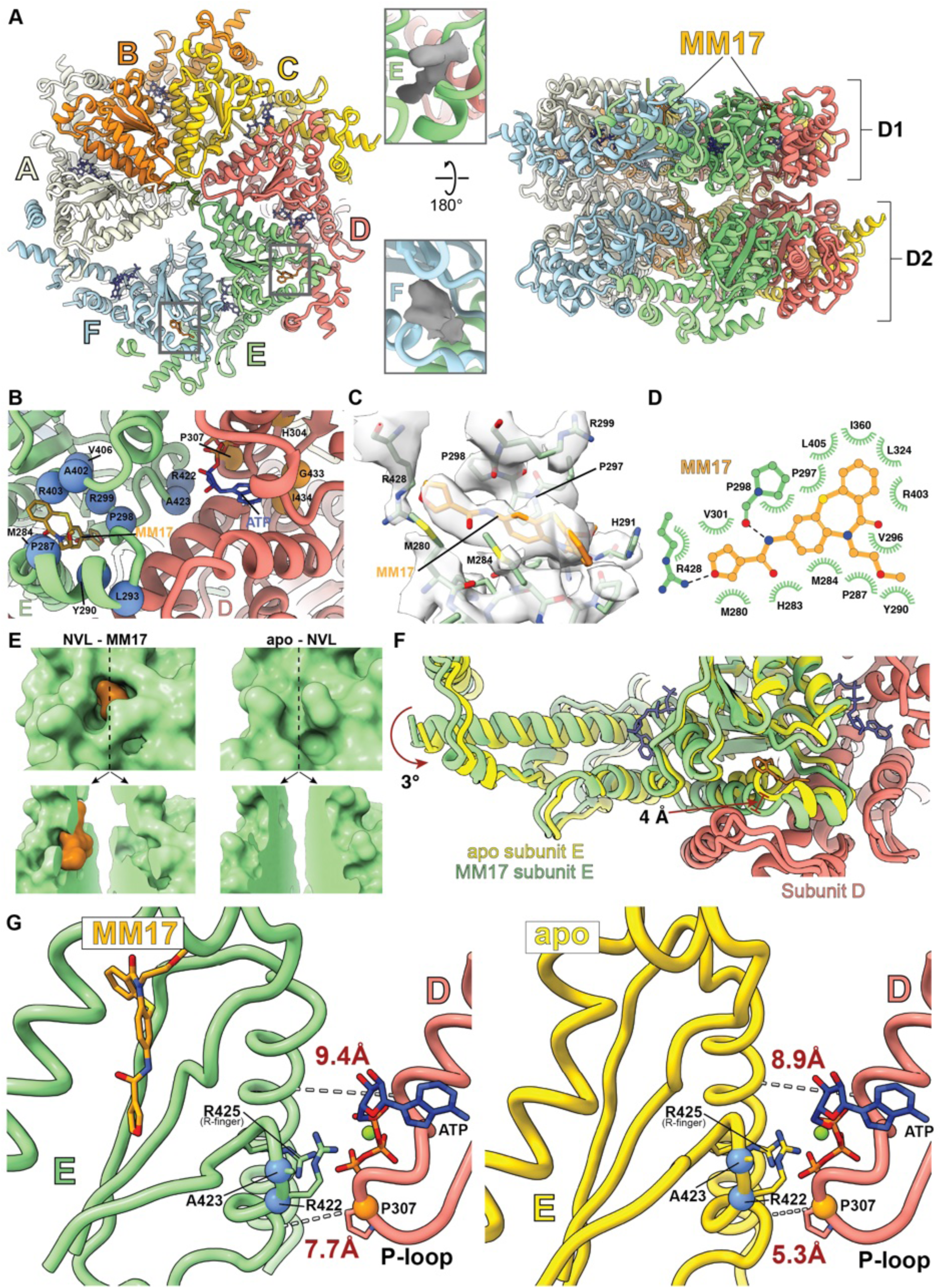
Cryo-EM structure of NVL^dEQ^ in complex with MM17. **A.** Overview of the NVL^dEQ^ hexamer model viewed along (left) and perpendicular to (right) the central pore axis. D1 and D2 AAA+ ATPase modules are indicated. Monomers are colored individually and labeled A-F. ATP molecules at the subunit interfaces are shown as dark blue sticks. MM17 (orange sticks) binds to subunits E and F. Insets show cryo-EM density for MM17 in each subunit. **B.** Close-up view of the MM17 binding pocket in subunit E. NVL subunits are shown as a cartoon models; MM17 is shown as sticks. Residues mutated in MM17-resistant clones and located near the MM17 binding pocket are shown as blue spheres; those near the adjacent ATP binding pocket are shown in orange. **C.** Detailed view of MM17 in the subunit E binding pocket. Protein residues and MM17 are shown as sticks; the cryo-EM map density for MM17 is rendered as a transparent surface. **D.** Interaction diagram of MM17 (orange) and surrounding NVL residues (green). Hydrophobic contacts are depicted as fans; hydrogen bonds are indicated by dashed lines. **E.** (Top) surface representations of the MM17 binding pocket in the MM17-bound (left) and apo (right) NVL structures. The protein surface is shown in green and MM17 in orange. (Bottom) Splayed open surfaces highlighting the absence of a pre-formed binding pocket in the apo structure. **F.** Structural overlay of the D and E subunits from NVL^dEQ^-MM17 (colored as in A) and apo-NVL^dEQ^ (subunit E in yellow). Arrows indicate MM17-induced swiveling (in degrees) and backbone displacement (in Å). **G.** Comparison of the ATP binding interface between D1 domains of subunits D and E in the MM17-bound (left) and apo (right) structures. MM17, ATP, arginine finger, and P307 are shown as sticks. Dashed lines indicate distances between selected Cα atoms (T312-D393 and P307-A419) to highlight allosteric rearrangements at the D/E interface.

Following convention, we designate the topmost subunit in the staircase as A and the lowest as E. As in other AAA+ ATPases, the sixth subunit (F) is dynamic, transitioning between positions E and A, and is therefore poorly resolved in the cryo-EM map (Figs. S3E and S4F). Based on bulky side-chain density in the channel and by analogy to the prior Rix7 analyses (*41*), we hypothesize that the substrate mimic engaged by NVL corresponds to one of the N-terminal 14x-HIS tag present on our construct. We therefore modeled this density as a polyhistidine chain, although the tag also contains non-histidine linker residues.

Initial cryo-EM map analysis revealed additional density consistent with MM17 in a single site within subunit E (Fig. 2A-C). MM17 is inserted into a predominantly hydrophobic pocket in the core domain of the D1 AAA+ module. This pocket is enclosed on one side by the central D1 β-sheet (strands 5, 1 and 4) and on the other side by the N-terminal α0 helix and the adjacent loop element preceding strand β1 (Fig. 2B). The α0 helix also engages the AAA+ lid-domain of subunit D, which forms the peripheral edge of the MM17 binding pocket (Fig. 2A). No equivalent pocket is present in subunits A-D, where the α0 helix is tightly packed against the central β-sheet of the AAA+ ATPase domain, precluding ligand engagement. To determine if MM17 also binds the mobile and poorly resolved F subunit, we performed a skip-align classification using a mask around the subunit F D1 module (Fig. S4C). The resulting reconstruction showed MM17 density at an analogous position in subunit F (Fig. 2A). Our analysis focused on the better resolved subunit E binding pocket near the functionally critical D/E subunit interface. Resistance-conferring mutations cluster either around the ligand-binding pocket, consistent with structural disruption of the binding pocket, or near the nucleotide-binding interface between subunits D and E, suggesting an allosteric mechanism that links ligand binding to nucleotide occupancy (Fig. 2B). MM17 is an elongated molecule with a ∼115° kink introduced by the dibenzothiazepinone scaffold (Fig. 2C, D). Its furan ring is exposed to the central solvent channel separating the D1 and D2 hexameric tiers, where it forms hydrophobic interactions with M280 and V301 and a hydrogen bond with R428 (Fig. 2C, D). Adjacent to the furan moiety, the planar amide linker hydrogen bonds with the backbone carbonyl of P298 (Fig. 2C, D). The dibenzothiazepinone ring is deeply embedded within a hydrophobic pocket formed by M280, M284, P297, L324, R403 and L405, four of which (M284, P297, R403 and L405) are either mutated or adjacent to resistance-conferring residues (Fig. 2C, D, Table S2). The carbonyl group of the central ring is positioned within hydrogen bonding distance of the ε-amino group of R403. Finally, the methoxyethyl tail projects out near the loop connecting α0 and β1 (Fig. 2B, D).

The presence of resistance mutations near the nucleotide-binding pocket adjacent to the MM17 binding pocket (Fig. 2B) prompted us to carry out single-particle cryo-EM reconstruction of apo-NVL^dEQ^ at 2.83 Å resolution, to examine how ligand binding influences inter-subunit dynamics that regulate nucleotide binding at NVL subunit interfaces (Fig. S6). In the apo structure, the NVL hexamer retains the staircase conformation, but the ligand binding pocket in subunit E is closed, adopting the same configuration as in subunits A-D (Fig. 2E). In the absence of ligand, the D1 domain of subunit E shifts closer to the neighboring AAA+ module in subunit D, closing the bipartite nucleotide-binding site at the D/E interface and bringing the arginine finger of subunit E into closer proximity to the bound ATP compared to the MM17-bound state (Fig. 2F, G, Movie S1). These structural changes suggest that MM17 disrupts key nucleotide interactions at the D/E interface within the D1 ring. Four resistance mutations (H304R, P307T, R422C and D423P) map to either the P-loop of subunit D or the helix containing the arginine-finger in subunit E (Fig. 2B), consistent with a model in which compensatory substitutions that enhance nucleotide binding and/or weaken the allosteric coupling between ligand and nucleotide sites alleviate the MM17-mediated inhibition of NVL function (Fig. 2G, Movie S1). The apo structure also offers clues to the selective binding of MM17 to subunits E and F. In the staircase hexamer arrangement, these subunits are less constrained by neighboring subunits and therefore exhibit greater conformational variability, as evidenced by lower local resolution and more diffuse map densities compared to tightly engaged subunit junctions (Fig. S5D). This increased structural flexibility underlies the selective access of MM17 to the occluded binding pockets in subunits E and F.

### MM17 inhibits NVL and blocks 60S biogenesis

We hypothesized that MM17 exerts its effects by inhibiting NVL activity. To test this, we modeled NVL loss of function by engineering a FLAG-tagged auxin-inducible degron (AID) into the endogenous *NVL* locus in cells expressing an F74A mutant of the plant OsTIR1 E3 ligase (Fig. S7A) (*44*). NVL^AID/AID^ and NVL^AID/+^ cells grew similarly to the cells expressing untagged NVL (Fig. S7B). Treatment with 5-Phenyl-indole-3-acetic acid (Ph-IAA) led to dose-dependent depletion of NVL-AID but not untagged NVL (Fig. S7C) and reduced viability of NVL^AID/AID^ cells (Fig. 3A). The viability of NVL^+/+^ or NVL^AID/+^ cells was unaffected by Ph-IAA (Fig. 3A), consistent with prior results demonstrating that a single allele of NVL is sufficient to support growth (Fig. S7B).

**Figure 3.**
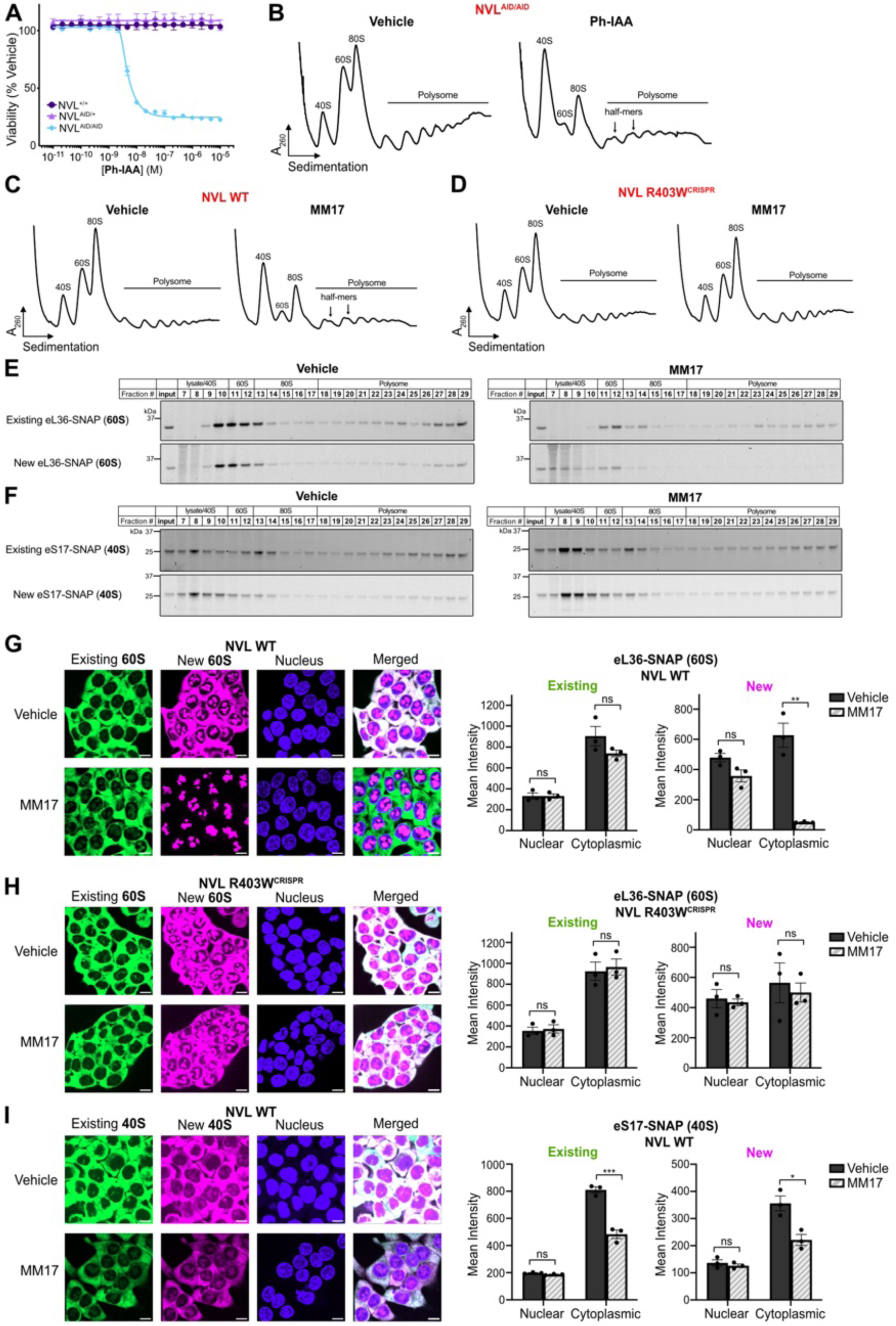
MM17 inhibits NVL and suppresses 60S biogenesis. **A.** Viability assay of HCT116 NVL^+/+^, NVL^AID/+^, and NVL^AID/AID^ cells treated with Ph-IAA for 72 hours. (n = 3 biological replicates, mean ± sem). **B.** Polysome profiling of HCT116 NVL^AID/AID^ treated with 100 nM Ph-IAA for 24 hours. **C, D.** Polysome profiling of HCT116 NVL WT (**C**) and NVL R403W^CRISPR^ (D) cells treated with 3 µM MM17 for 24 hours. **E, F.** Fluorescence scans of fractions separated by sucrose gradient centrifugation, showing the distribution of existing or newly synthesized 60S protein eL36-SNAP (**E**) or 40S protein eS17-SNAP (**F**), following 3 µM MM17 treatment for 6 hours. Existing and newly synthesized SNAP tagged proteins are fluorescently labeled with Oregon green and 647-SiR benzylguanine, respectively. **G, H.** Representative microscopy images and quantitation of existing (green) and newly synthesized (magenta) eL36-SNAP in NVL WT (**G**) and NVL R403W^CRISPR^ (**H**) HCT116 cells treated with 3 µM MM17 for 24 hours. (Scale bar = 10 µm; n = 3 biological replicates, mean ± sem; two-tailed unpaired t-test, **p<0.01, ns = non-significant). **I.** Representative microscopy images and quantitation of existing (green) and newly synthesized (magenta) eS17-SNAP in NVL WT HCT116 cells treated 3 µM MM17 for 24 hours. (Scale bar = 10 µm; n = 3 biological replicates, mean ± sem; two-tailed unpaired t-test, ***p<0.001, *p<0.05, ns = non-significant).

We next used polysome profiling to assess how NVL depletion alters the distribution of ribosomal species. Ph-IAA-induced NVL degradation in NVL^AID/AID^ cells, but not NVL^+/+^ cells, caused a time-dependent reduction in free 60S and 80S peaks (based on absorbance at 260 nm) and a corresponding increase in free 40S levels (Fig. 3B, Fig. S7D). Ph-IAA addition also induced half-mer polysomes, representing unjoined 40S subunits stalled at translation initiation sites, a hallmark of defective 60S biogenesis (*45*). MM17 treatment in NVL WT cells phenocopied the ribosomal distribution from Ph-IAA-treated NVL^AID/AID^ cells but had no effect in NVL R403W^CRISPR^ cells (Fig. 3C, D). The altered ribosomal subunit distribution following NVL disruption in HCT116 cells is similar to profiles reported in yeast with *RIX7* mutations (*46*) or in human cells overexpressing catalytic mutants of *NVL* (*47*).

The observed reduction in free 60S could reflect either decreased 60S synthesis or subunit turnover, but distinguishing between these mechanisms requires differentiating newly synthesized from pre-existing 60S subunits. We therefore engineered HCT116 cells to express a SNAP-tag on eL36, a ribosomal protein incorporated into the 60S subunit prior to NVL binding (*48*) or, as a control, on the small ribosomal subunit protein eS17 to monitor 40S biogenesis (Fig. S7E) (*48*). We first labeled pre-existing SNAP tagged eL36 or eS17 in cells with Oregon green benzylguanine (green), then treated cells with vehicle or MM17 for 6 hours, followed by a second labeling step with 647-SiR benzylguanine (magenta) to label newly synthesized ribosomal proteins. We prepared polysome profiles from lysates of these cells by sucrose gradient centrifugation and analyzed each gradient fraction for pre-existing (green) or new (magenta) fluorescence to determine the distribution of labeled ribosomal proteins. Existing eL36-SNAP was present in 60S, 80S, and polysome fractions, confirming its incorporation into mature translating ribosomes (Fig. 3E, S7F, G). MM17 did not affect the levels or distribution pattern of pre-existing eL36-SNAP. In contrast, newly synthesized eL36-SNAP was shifted to the non-ribosomal upper fractions and depleted from the 60S, 80S, and polysome fractions following MM17 treatment, consistent with a block in 60S biogenesis. In vehicle- or MM17-cells, both existing and newly synthesized eS17-SNAP remained distributed across the 40S, 80S, and polysome fractions, indicating that MM17 has limited impact on 40S production (Fig. 3F, S7H, I).

Next, we examined the effect of MM17 on the subcellular localization of eL36-SNAP (60S) or eS17-SNAP (40S). Ribosome maturation begins in the nucleolus and proceeds through concerted steps that accompany release into the nucleoplasm and subsequent export to the cytoplasm, where final maturation occurs before subunits can assemble for translation. MM17 had no impact on the localization of pre-existing eL36-SNAP compared to vehicle-treated cells (Fig. 3G). By contrast, MM17 caused a 12-fold reduction in cytoplasmic signal from newly synthesized eL36-SNAP, which instead localized exclusively to nucleolar foci as defined by colocalization with Fibrillarin (Fig. S7J, K). This is consistent with the known subcellular localization of NVL (*49*) and indicative of nucleolar retention of 60S subunits.

To test whether the biogenesis block is caused by MM17 binding to NVL, we engineered cells expressing eL36-SNAP and NVL R403W^CRISPR^ (Fig. S7L). In these cells, MM17 treatment did not alter the localization of newly synthesized eL36-SNAP, which remained both nuclear and cytoplasmic, indistinguishable from the vehicle-treated cells (Fig. 3H). These findings indicate that MM17-induced nuclear accumulation of immature 60S subunits is due to inhibition of NVL. Finally, we asked whether MM17 affects 40S production by performing analogous experiments in eS17-SNAP-expressing cells. Compared to vehicle, MM17 led to a modest 1.7- and 1.6-fold reduction in the cytoplasmic localization of both pre-existing and newly synthesized eS17-SNAP, respectively, suggesting a slight feedback effect between 60S and 40S production (Fig. 3I). Together, these results demonstrate that MM17 blocks nucleolar 60S assembly by directly inhibiting NVL.

The preservation of existing ribosomes and the persistence of polysomes following MM17 treatment (Fig. 3C) suggested that global translation may be preserved. To directly assess the impact of MM17 on global protein synthesis, we measured O-propargyl puromycin incorporation. While markedly reduced by the protein synthesis inhibitor cycloheximide, MM17 does not affect global translation over 24 hours (Fig. S7M), although we cannot exclude selective effects on specific mRNAs. These findings indicate that pre-existing ribosomes are sufficient to sustain bulk translation during acute inhibition of 60S biogenesis. Thus, MM17 likely acts through a mechanism distinct from canonical translation inhibitors such as cycloheximide.

### NVL inhibition leads to p53 dependent and independent cell cycle arrest

To better understand how NVL inhibition affects cell growth, we performed an unbiased genome-wide pooled CRISPR/Cas9 knockout screen in HCT116 cells using a library of guide RNAs targeting 19,114 genes. Cells were intermittently exposed to 24-hour pulses of either vehicle or MM17 over 21 days, during which MM17-treated cells underwent 9.79 fewer population doublings (Fig. S8A). Massively parallel sequencing of PCR products amplified from genomic DNA was used to determine relative levels of sgRNA sequences to infer the relative impact of each gene on MM17 sensitivity (Table S4). *RPF1*, which encodes the human ortholog of one of the three biogenesis factors removed by Rix7 in yeast (*33*), is the 8^th^ most depleted gene (Fig. 4A), consistent with our conclusion that MM17 targets NVL. The tumor suppressor *TP53,* which encodes the p53 protein, and one of its effector genes, *CDKN1A,* which encodes p21, were the most enriched genes in this screen, indicating that loss of p53 and, to a lesser degree, p21, confers a degree of resistance to MM17. To validate these findings, we performed growth competition experiments between cells expressing ZsGreen (green) with either p53 or p21 knockout (KO) and mock cells expressing mCherry (red), using 24-hour pulses of MM17 (Fig. S8B, C). In the absence of treatment, the ratio of green to red cells remained constant over time, whereas MM17 induced a dose-dependent enrichment of both p53 and p21 knockout cells. (Fig. S8D). This effect was specific to MM17, as paclitaxel, a microtubule-stabilizing toxin, did not confer a similar advantage (Fig. S8E).

**Figure 4.**
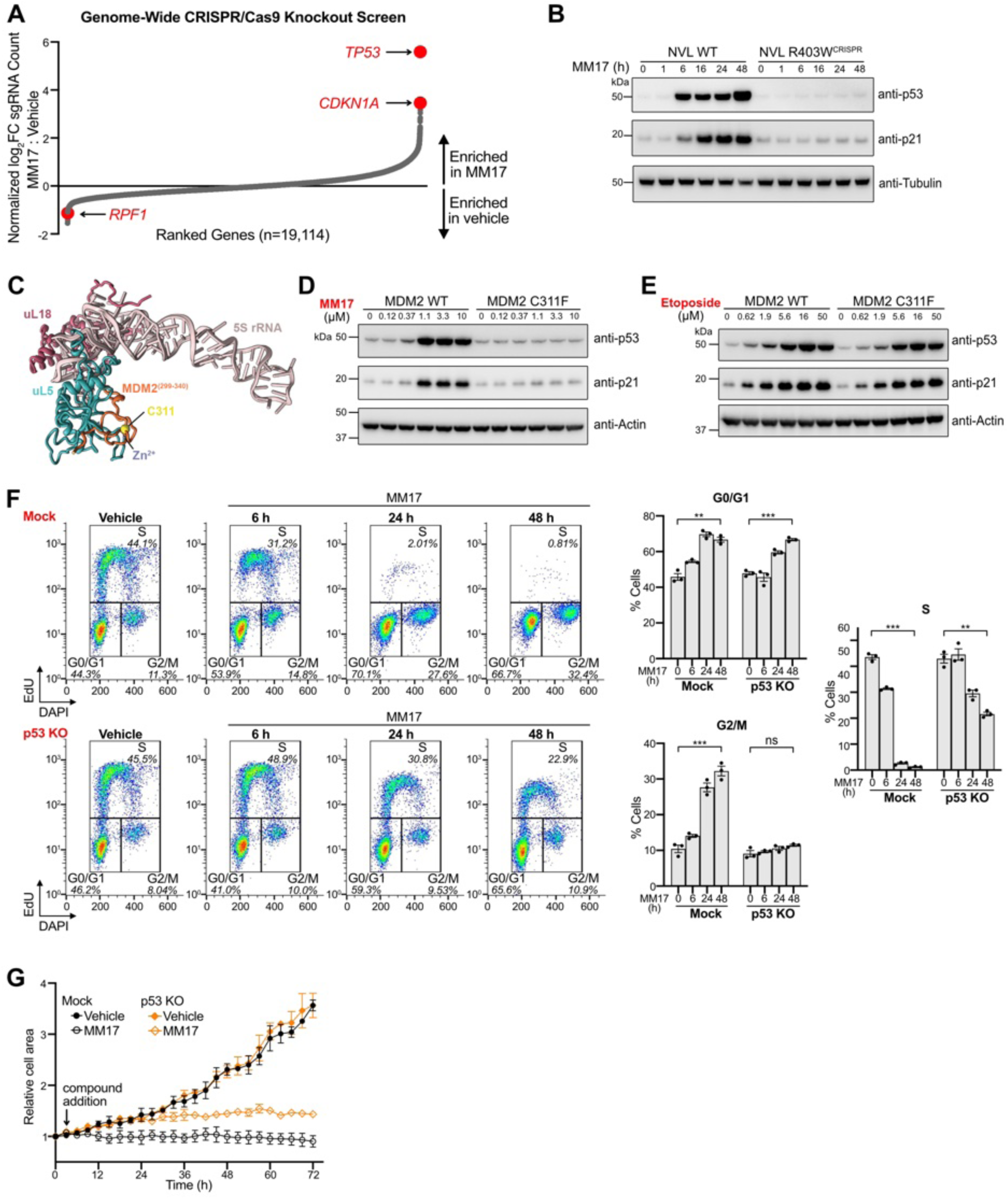
NVL inhibition leads to p53 dependent and independent cell cycle arrest. **A.** Genome wide CRISPR/Cas9 knockout screen in HCT116 cells. The log2 fold change for each gene comparing samples treated with vehicle or 1 µM MM17 was determined by averaging the log2 fold changes of its associated sgRNAs and subtracting the average log2 fold change of non-targeting sgRNAs. The top two most enriched genes, and one of the biogenesis factors removed by Rix7 are indicated in red and labeled. **B.** Immunoblots of p53, p21, and tubulin (loading control) in HCT116 NVL WT and NVL R403W^CRISPR^ cells treated with 3 µM MM17. **C**. Cryo-EM structure of the MDM2^299-340^ –5S RNP complex (PDB ID - 8BGU; MDM2 - orange, uL5 - turquoise, uL18 - red, 5S rRNA - pink) (*50*). MDM2 C311 residue (yellow) and Zn^2+^ (purple) at the MDM2-uL5 binding interface are labeled. **D, E.** Immunoblots of p53, p21 and actin (loading control) in HCT116 MDM2 WT and MDM2 C311F cells treated with MM17 (**D**) or etoposide (E) for 24 hours. **F.** Cell cycle analysis of HCT116 mock and p53 KO cells treated with 3 µM MM17. (n = 3 biological replicates, mean ± sem; two-tailed unpaired t-test, ***p<0.001, **p<0.01, ns = non-significant). **G.** Live cell imaging analysis of relative cell area of HCT116 mock and p53 KO cells treated with 3 µM MM17 for 72 hours. (n = 3 biological replicates, mean ± sem).

We then assessed whether there was a time-dependent induction of p53 or p21 in HCT116 cells following exposure to MM17. p53 levels increased starting at 6 hours, followed by the upregulation of p21, consistent with its role as a p53 target gene (Figs. 4B, S8F). Importantly, NVL R403W^CRISPR^ cells did not respond, indicating that p53 and p21 activation results from direct NVL inhibition. The p53 protein can be stabilized by the impaired ribosome biogenesis checkpoint (IRBC), triggered by accumulation of free 5S RNP particles (components of the 60S ribosome) that bind to the zinc finger domain of MDM2, blocking p53 ubiquitination and degradation (*50–53*). Mutations in the Zn-coordinating cysteines in this MDM2 domain, including C311F, prevent 5S RNP binding, and render cells resistant to the IRBC without impacting other p53 activating pathways, including DNA damage response (Fig 4C) (*50, 54, 55*). To test whether MM17 triggered the IRBC, we compared p53 activation in MDM2 knockout cells that ectopically expressed wild type or MDM2 C311F (Fig. S8G-I). MM17 treatment robustly upregulated p53 in MDM WT controls but failed to induce p53 in cells expressing MDM2 C311F (Fig. 4D, S8J). In contrast, treatment of MDM2 WT and MDM2 C311F cells with the DNA damaging agent etoposide resulted in comparable increases in p53, confirming that MDM2 C311F specifically impairs the IRBC (Fig. 4E, S8K, L). Together, these findings provide evidence that MM17 triggers p53 induction through the IRBC by targeting NVL.

p53 stabilization can lead to cell cycle arrest or apoptosis (*56*). To assess whether MM17 induces apoptosis, we analyzed procaspase-3 cleavage into its active form. As a positive control, we used MLN4924, a distinct toxin that induces apoptosis by inhibiting the neddylation activation enzyme complex, which caused a marked decrease in procaspase-3 levels and increase in cleaved caspase-3, consistent with apoptotic activation (Fig. S9A). In contrast, prolonged MM17 treatment for up to 3 days had no effect on either procaspase-3 or activated caspase-3 levels. These results suggest that the mechanism by which MM17 reduces cell viability is not primarily through apoptotic cell death in HCT116 cells. We therefore examined cell cycle progression following MM17 exposure. MM17 caused an accumulation of cells in both G1 and G2 phases of the cell cycle, accompanied by a corresponding decrease in S-phase cells (Fig. S9B), indicating that MM17 blocks progression through both G1/S and G2/M transitions.

To evaluate the role of p53 on MM17-induced cell cycle arrest, we used CRISPR/Cas9 to silence p53 expression in HCT116 cells (Fig. S9C). The distribution of cells across various cell cycle phases was comparable between vehicle treated mock and p53 knockout (KO) cell lines (Fig. 4F). In mock cells, MM17 caused a time-dependent increase in both G0/G1 and G2/M cell percentages with a concomitant decrease in S-phase cell percentages, consistent with both G1/S and G2/M arrest. In p53 KO cells treated with MM17, the G1/S arrest was delayed relative to mock cells, suggesting that while p53 may initially accelerate G1/S arrest, it ultimately is p53-independent. Strikingly, G2/M accumulation was absent in p53 KO cells, indicating that this checkpoint arrest is p53-dependent. Together these results indicate that NVL inhibition leads to a p53-independent G1/S arrest and a p53-dependent G2/M arrest. Consistent with these findings, live cell imaging revealed a biphasic effect of MM17 inhibition on cell growth based on p53. In the first 24 hours, MM17 slowed growth of mock-edited cells but not p53 KO cells (Fig. 4G).

These findings align with the results of our CRISPR screen in which cell populations were intermittently pulsed with MM17 for 24 hours over 21 days. After 24 hours, MM17 arrested growth in both the mock and p53 KO populations. We concluded that MM17 acutely inhibits proliferation in a p53- and p21-dependent manner, but that prolonged NVL inhibition ultimately suppresses cell growth independent of p53.

The TP53 gene is a tumor suppressor and one of the most frequently mutated or deleted genes in human cancer (*57*). To broadly assess whether p53 status influences MM17 activity in other cell types, we used a PRISM (profiling relative inhibition simultaneously in mixtures) screen consisting of 855 cancer cell lines representing 26 distinct lineages to compare the viability between cell lines with or without functional (“hotspot”) *TP53* mutations or *TP53* deletions (*58*) following MM17 treatment. Cell lines with *TP53* mutations or deletions correlated with reduced MM17 activity (Fig. S9D, Table S5), however, the average difference in sensitivity was minimal, consistent with our conclusion that MM17 reduces viability in the absence of p53.

### A bioavailable analog reduces tumor growth in mouse xenografts without overt toxicity

MM17 represents a potential tool to determine whether selective inhibition of ribosome biogenesis can exhibit an anti-tumor therapeutic index. However, its short half-life (t ½=2.68 min) in the presence of mouse liver microsomes predicted poor bioavailability in mice (Fig. S10A). To overcome this limitation, we synthesized MM927, a structurally related compound with improved metabolic stability (t ½ = 25.34 min) and nearly 10-fold greater potency (IC50 = 0.053 µM) (Fig. 5A, S10B, C). To assess bioavailability, we quantified MM927 plasma levels at different time points following a 10 mg/kg intraperitoneal (IP) injection. MM927 reached a peak concentration (Cmax) of 2510 ng/mL (5.15 µM) with an elimination half-life of 101.33 minutes (Fig. 5B, S10D). Plasma exposure increased in a dose-dependent manner, reaching a maximum concentration of 17.86 µM following injection with 35 mg/kg (Fig. S10E).

**Figure 5.**
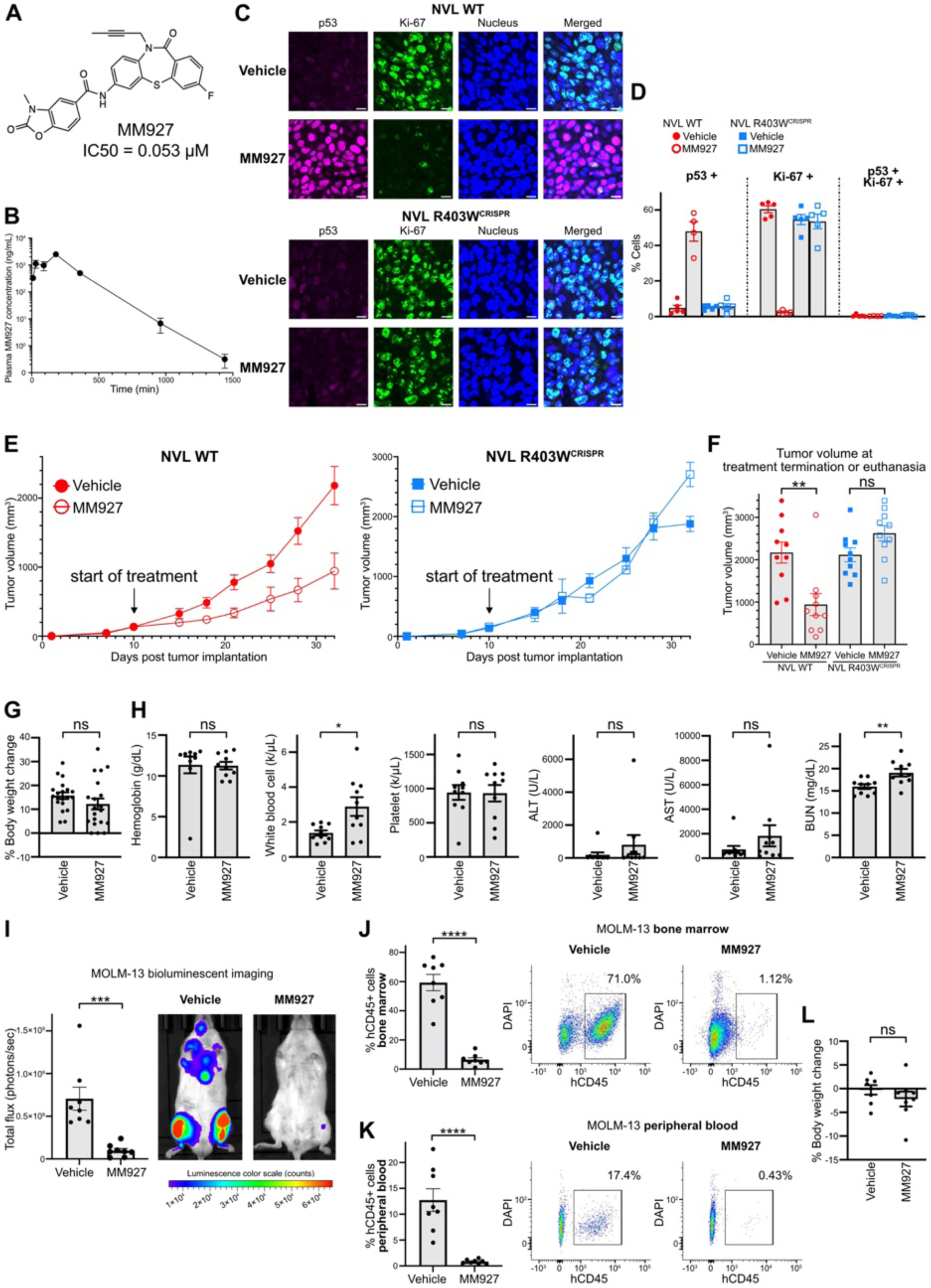
A bioavailable analog MM927 inhibits colorectal and leukemia tumor xenograft growth *in vivo* without overt toxicity. A. Chemical structure of MM927 and the potency (IC50) of its impact on the viability of HCT116. **B.** Pharmacokinetic analysis of MM927 in mouse plasma after a 10 mg/kg intraperitoneal (IP) injection (n = 3 mice/group, mean ± sem). **C, D.** Representative microscopy images (**C**) and quantitation (**D**) showing immunofluorescence of p53 (magenta), Ki-67 (green), and nucleus (DAPI, blue) in NVL WT and NVL R403W^CRISPR^ tumor xenografts implanted in either flank of the same mouse. Mice were treated with 35 mg/kg MM927 IP twice daily for 3 days, and tumors were harvested 6 hours after the last dose. (Scale bar = 10 µm; n = 5 mice/group, mean ± sem). **E.** Effect of MM927 (35 mg/kg IP twice daily for 21 days) on HCT116 NVL WT or NVL R403W^CRISPR^ tumor xenograft growth (n = 10 mice/group, mean ± sem). **F**. Tumor volume measurements in (E) at the end of 21 days or euthanasia due to tumor diameter exceeding 2 cm (n = 10 mice per group, mean ± sem; two-tailed unpaired t-test, ** p < 0.01, ns = non-significant). **G.** Body weight % changes in HCT116-tumor bearing mice at the end of MM927 treatment (n = 20 mice/group, mean ± sem; two-tailed unpaired t-test, ns = non-significant). **H.** Levels of hemoglobin, white blood cells, platelets, liver enzymes AST and ALT, and renal clearance of blood urea nitrogen (BUN) 3 hours after the last dose (n = 10 mice per group, mean ± sem; two-tailed unpaired t-test, **p<0.01, *p<0.01, ns = non-significant). **I.** Representative images and quantitation of leukemic disease burden by bioluminescent imaging of luciferase expressing MOLM-13 cells. (n = 8 mice/group, mean ± sem; two-tailed unpaired t-test, *** p < 0.001). **J, K**. Flow cytometry-based quantitation of MOLM-13 cells (human CD45 positive) in the mouse bone marrow (**J**) and peripheral blood (**K**) (n = 8 mice/group, mean ± sem; two-tailed unpaired t-test, ****p<0.0001). **L**. Body weight changes at the end of MM927 treatment in mice xenografted with MOLM-13 (n = 8 mice/group, mean ± sem; two-tailed unpaired t-test, ns = non-significant).

Prior to evaluating MM927 in mouse models of human cancer, we asked if MM927 has the same binding mode and mechanism of action as MM17. To test this, we determined a 2.86 Å cryo-EM structure of NVL^dEQ^ in complex with MM927, focusing on the ligand-binding pocket in subunit E (Fig. S11, S12A). The dibenzothiazepinone rings of MM17 and MM927 overlap nearly perfectly when the two structures were superposed (Fig. S12B). The benzoxazolone moiety of MM927 occupies the same region as the furan ring of MM17 but makes additional contacts with surrounding residues (Fig. S12C, D). In addition, the rigid and more hydrophobic butynyl tail of MM927 appears to stabilize the α0/β1 loop, which is comparatively more flexible in the presence of the methoxyethyl tail of MM17 (Fig. S12D). Apart from these minor differences, the overall binding conformation and resulting disruption of the D/E subunit interface closely resemble those observed with MM17, indicating that MM927 shares a common binding site and mechanism of action. Cell-based assays confirmed that MM927 reduces viability through the same mechanism as MM17, by blocking 60S biogenesis via NVL inhibition. Like MM17, MM927 induced half-mer polysomes in polysome profiles and arrests 60S ribosome biogenesis in the nucleolus (Fig. S13A-D). MM927 also triggered cell cycle arrest at both the G1/S and G2/M transitions, mirroring the effects of MM17, with G1/S arrest partially and G2/M arrest fully dependent on p53 (Fig. S13E). Consistent with target engagement, the IC50 of MM927 was nearly 100-fold higher in NVL R403W^CRISPR^ cells compared to parental controls (Fig. S13F).

To interpret the consequence of blocking NVL in normal tissues, we next determined whether MM927 inhibits mouse NVL at concentrations comparable to the human enzyme. We ectopically expressed either human or mouse NVL in NVL^AID/AID^ cells (Fig. S14A) and treated with Ph-IAA so that the cells must rely solely on ectopically expressed NVL. As expected, Ph-IAA reduced cell viability in mock-infected cells, while expression of either mouse or human NVL fully rescued growth (Fig. S14B). Under these conditions, MM927 inhibited cells expressing ectopic human NVL with an IC50 of 0.051 µM (Fig. S14C), comparable to its potency with endogenous NVL, suggesting that NVL overexpression does not alter sensitivity. MM927 also inhibited mouse NVL with an IC50 of 0.207 µM, providing a benchmark plasma concentration that would be capable of inhibiting NVL in normal mouse tissue. Importantly, maximum plasma levels of MM927 following a 35 mg/kg injection exceed 15 μM (Fig. S10C, E), well above the viability inhibition IC50 of MM927 in cells expressing either human or mouse NVL (Fig. S14C), confirming that therapeutically relevant exposures are achieved *in vivo*.

To directly evaluate target engagement *in vivo*, we measured the kinetics of p53 and p21 induction in HCT116 xenograft tumors in mice treated with MM927. We implanted HCT116 NVL WT or NVL R403W^CRISPR^ cells subcutaneously on opposite flanks of an individual mouse and evaluated the levels of p53 and p21 in each tumor at various time points after IP administration of 35 mg/kg MM927. Compared to vehicle treated mice, p53 protein levels increased beginning at 3 hours post-injection, followed by p21 induction soon thereafter (Fig. S14D, E). Both p53 and p21 remained elevated at 12 hours and returned to basal levels by 24 hours, suggesting that twice daily dosing is required for persistent target engagement.

Importantly, the absolute levels of p53 and p21 induction in tumor xenografts were comparable to what we observed in cultured cells treated with 24 hours of continuous MM17. Critically, NVL R403W^CRISPR^ tumors implanted on the opposite flank of the same mice showed no change in p53 or p21, suggesting that the observed changes in NVL WT tumors were the result of NVL inhibition and establishing p53 and p21 as viable pharmacodynamic (PD) markers.

Having established the kinetics of NVL inhibition by MM927 *in vivo*, we next assessed markers of tumor cell growth following treatment with 35 mg/kg over three days. Mice bearing matched NVL WT or NVL R403W^CRISPR^ xenografts on opposite flanks were treated with MM927 twice daily for 3 days. Tumors were stained for p53 and Ki-67, a marker of proliferating cells. MM927 treatment substantially increased nuclear p53 staining and substantially reduced Ki-67-positive cell staining (Fig. 5C, D). Consistent with a cell cycle arrest phenotype, few to no cells were stained with both markers. In contrast, NVL R403W^CRISPR^ derived tumors showed no change in either marker. Together, these results indicate that MM927 is a bioavailable inhibitor of both mouse and human NVL and a promising candidate for evaluating efficacy and toxicity in mouse models of cancer.

We treated mice bearing either HCT116-derived NVL WT or NVL R403W^CRISPR^ tumors with 35 mg/kg MM927 by IP twice daily for 21 days. MM927 treatment reduced the growth of NVL WT tumors but had no impact on NVL R403W^CRISPR^ tumors (Fig. 5E, S14F). Tumor measurements collected either at the end of treatment or the time of euthanasia demonstrated that MM927 reduced NVL WT tumor burden, but had no effect on NVL R403W^CRISPR^ tumors (Fig. 5F). MM927 was detected at micromolar levels in the plasma and tumors of all treated mice 3 hours after the last treatment dose, demonstrating adequate drug exposure was achieved in both cohorts (Fig. S14G). Notably, mice treated with MM927 exhibited no signs of overt toxicity.

There were no differences in body weight changes, hemoglobin, platelets, or the liver enzymes ALT and AST (Fig. 5G, H), and only small increases in white blood cells and blood urea nitrogen (Fig. 5H). Taken together, these findings suggest that the potent and bioavailable analog MM927 reduces tumor growth in HCT116 mouse xenografts by inhibiting NVL, without overt or dose limiting toxicities.

To test whether NVL inhibition could be effective across cancer types, we tested the potency of MM17 in MOLM-13 cells, a p53 wild-type cell line derived from a patient with acute myelogenous leukemia (AML) (*59*). We used CRISPR to generate isogenic MOLM-13 p53 KO cell lines to assess the importance of p53 to MM17 sensitivity (Fig.S15A). MM17 reduced viability in both mock and p53 KO MOLM-13 cells with comparable IC50s (0.35 µM and 0.39 µM, respectively), demonstrating that p53 is not required for target engagement (Fig. S15B).

Unlike in HCT116 cells, MM17 treatment triggered apoptosis in MOLM-13 cells, evident by decreased procaspase-3 and corresponding increased cleaved caspase-3 that was comparable to the MLN4924 control (Fig. S15C). Consistent with apoptosis, the majority of MOLM-13 cells stained positive for Annexin V and were non-viable following treatment with MM17 for 2- or 3-days (Fig. S15D). In both assays for apoptosis and viability, p53 silencing slightly delayed but did not prevent apoptosis, indicating that NVL inhibition can induce either cell cycle arrest or apoptosis in a cell type-dependent and p53-independent manner (Fig. S15E).

To test the efficacy of MM927 in MOLM-13 mouse xenografts, we first tested how MOLM-13 cells ectopically expressing NVL R403W respond to MM927 (Fig. S15F).

Expression of either the NVL WT or NVL R403W transgene had no impact on cell viability (Fig. S15G), and NVL WT expression did not affect the potency of MM927 (Fig. S15H). The expression of NVL R403W, by contrast, increased the IC50 of MM927 from 0.092 μM to over 5 μM. These findings demonstrate that the anti-cancer activity of MM927 is through NVL binding. Next, we established xenografts by intravenous injection of MOLM-13 cells. Fifteen days post-injection, we treated mice with either vehicle or 35 mg/kg MM927 twice daily for 14 days.

MM927 treatment led to a 7-fold reduction of tumor burden by bioluminescence imaging, and 9- and 15-fold decreases of MOLM-13 cell percentages in the bone marrow and peripheral blood, respectively (Fig. 5I-K). Again, we observed no overt toxicity and no change in body weight between vehicle- and drug-treated mice (Fig. 5L).

### NVL inhibitors exhibit a response profile similar to 5-FU and oxaliplatin in cancer cell lines without inducing DNA damage

The cancer drugs 5-fluorouracil (5-FU), oxaliplatin, and the investigational RNA polymerase I inhibitor CX-5461 are small molecules that impair ribosome biogenesis but also induce DNA damage, complicating interpretation of their primary mechanism of action. To test whether NVL inhibitors share this liability, we quantified γH2AX foci by immunofluorescence as a marker of DNA damage. At IC100 concentrations, both 5-FU and oxaliplatin induced a marked increase in γH2AX foci, consistent with genotoxic stress (Fig. 6A, S16A-C). In contrast, treatment with MM17 or MM927—even at 50 µM, concentrations far exceeding their respective IC50s did not increase γH2AX signal above vehicle, indicating an absence of DNA damage (Fig. 6A, S16A-C). These findings demonstrate that MM17 and MM927 inhibit ribosome biogenesis without causing DNA damage.

**Figure 6.**
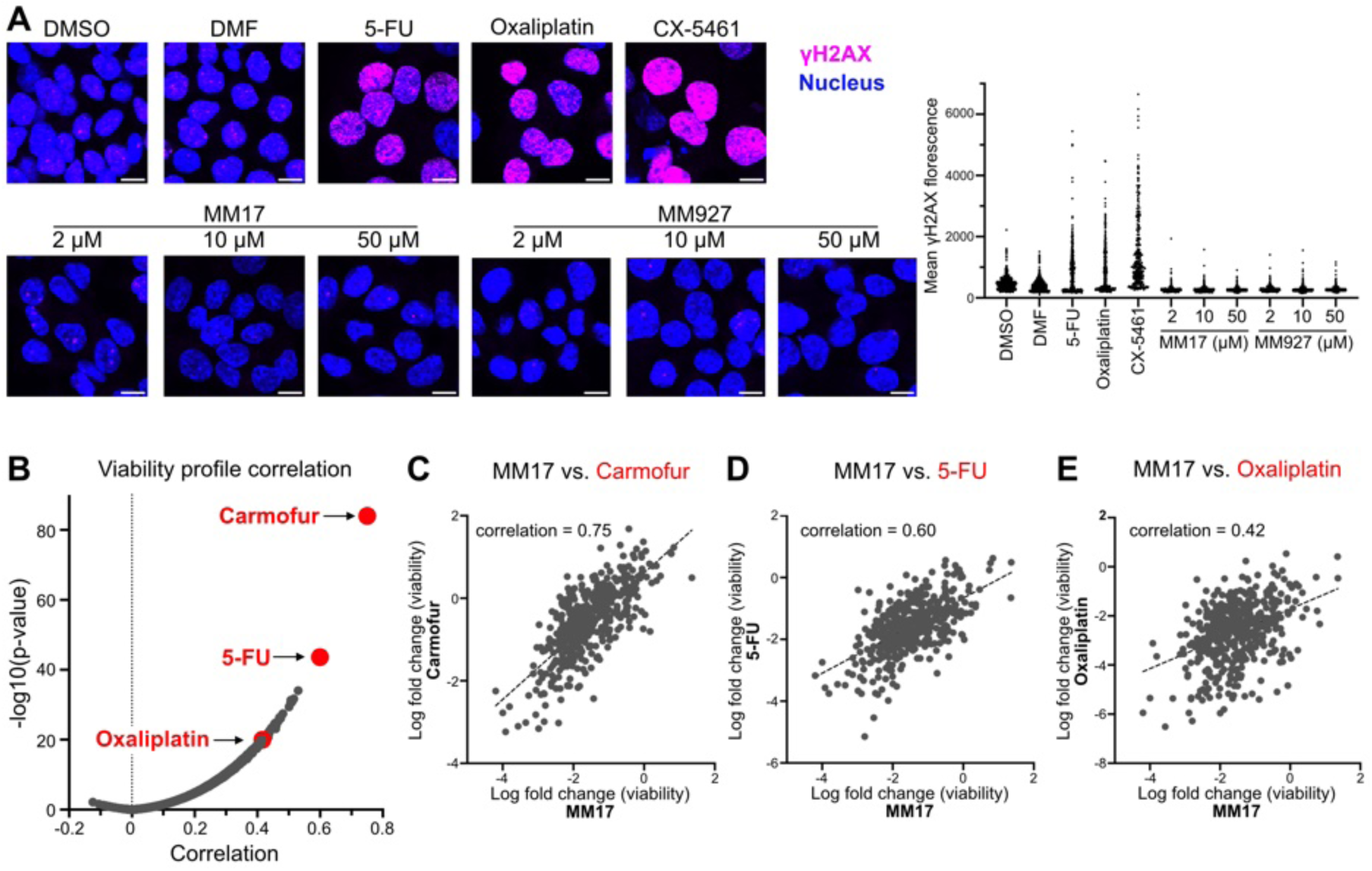
NVL inhibitors target cancer cells sensitive to 5-FU and oxaliplatin without causing DNA damage. **A.** Representative microscopy images of γH2AX immunofluorescence (magenta) and nucleus (DAPI, blue) in HCT116 cells treated with MM17, MM927, 150 µM 5-FU, 50 µM oxaliplatin, or 5 µM CX-5461 for 24 hours (Scale bar = 10 µm). Mean γH2AX fluorescence of individual nuclei is plotted. **B.** Correlation of MM17’s viability profile in 858 cancer cell lines with that of 1,182 other compounds previously reported in the PRISM screen. Each point represents a compound. **C-E.** Correlation of the viability of cancer cell lines between 10 µM MM17 and **(C)** 10 µM carmofur (n=462), **(D)** 10 µM 5-FU (n=440), **(E)** 10.3 µM oxaliplatin (n=460) where each point represents a cell line.

Despite their broad polypharmacology and dose-limiting toxicities, 5-FU and oxaliplatin remain cornerstone therapies for several cancers, including colorectal malignancies (*60*). Given that their clinical efficacy may derive, at least in part, from ribosome-targeting activity rather than their historical classification as DNA-damaging agents, we hypothesized that cancer cells sensitive to 5-FU or oxaliplatin might also respond to selective NVL inhibition. To explore this possibility, we compared the PRISM viability profile of MM17 to those of 1,184 compounds with PRISM datasets. Strikingly, 5-FU and its prodrug carmofur (*61*) ranked among the most highly correlated profiles, with oxaliplatin showing a similarly strong relationship (Fig. 6B-E, Table S6). Although the precise mechanisms of 5-FU and oxaliplatin remain incompletely defined, these findings reinforce the idea that inhibition of ribosome biogenesis is a shared effector mechanism and suggest that NVL inhibitors may benefit patient populations currently treated with 5-FU-based regimens.

## DISCUSSION

We report potent, specific, and bioavailable inhibitors of the ribosome biogenesis AAA+ ATPase NVL that block 60S subunit maturation and suppress tumor growth in both colorectal cancer and acute myeloid leukemia models, without overt toxicity. NVL inhibition did not impact existing ribosomes, and acutely, we did not observe a decrease in global translation. Rather, the immediate antiproliferative effect of NVL inhibition was mediated by p53 stabilization through the impaired ribosome biogenesis checkpoint (IRBC). Whereas direct translation inhibitors that impair mature ribosomes cause widespread cytotoxicity (*62*), 21 days of continuous treatment with NVL inhibitors did not lead to toxicity in mouse models. One explanation for this difference is that the on-target toxicity of NVL inhibition is mediated through p53 activation rather than translation inhibition. NVL inhibitors also triggered a delayed G1/S arrest in the absence of p53, suggesting a parallel mechanism by which cells sense ribosome output to regulate cell cycle progression. It remains to be determined if this mechanism is related to reduced protein translation stemming from fewer ribosomes. Even though we did not observe toxicity, we expect that more potent derivatives with higher target occupancy are likely to lead to on-target toxicity. Insights from ribosomopathies, genetic syndromes caused by defects in ribosome production, suggest that certain lineages may be especially susceptible, including hematopoietic stem cells and neural crest-derived cells (*63, 64*).

Conventional chemotherapies are remarkably effective, yet their therapeutic windows remain poorly understood, and treatment-related toxicities, often resulting from off-target polypharmacology, continue to constrain clinical benefit (*65*). Recent strategies to improve chemotherapy have centered on identifying predictive biomarkers of response or developing agents with greater mechanistic precision to reduce systemic toxicity (*66*). Selective targeting of ribosome biogenesis offers a complementary approach that may redefine the boundaries of cytotoxic therapy. We found that 5-fluorouracil (5-FU) and oxaliplatin exhibit strikingly similar viability profiles to MM17, aligning with previous reports implicating these agents in ribosome biogenesis inhibition. If the clinical efficacy of 5-FU and oxaliplatin, cornerstones of first-line colorectal cancer treatment, derives from disrupting ribosome production, then selective NVL inhibition may retain therapeutic benefit while avoiding genotoxicity, offering a targeted and potentially safer alternative to existing chemotherapy regimens.

Beyond therapeutic implications, NVL inhibitors offer a tractable system for probing the mechanism of type II AAA+ ATPases. Cryo-EM structures of catalytically inactive, substrate-bound NVL revealed that MM17 binds near the D/E subunit interface, a key site for ATP hydrolysis and translocase activity (*32, 67*), disrupting nucleotide coordination and subunit progression along the hexameric staircase. Indeed, a subset of resistance mutations cluster near the nucleotide binding pocket, consistent with allosteric disruption of the ATP-binding pocket at the D/E subunit interface by MM17. However, because these studies relied on a catalytically inactive hexamer and a substrate mimic, a complete understanding of NVL inhibition will require the *in vitro* reconstitution of the full catalytic cycle to fully understand the impact of MM17 on hexamer assembly, substrate engagement, ATP turnover, and translocation dynamics. Future efforts should also focus on identifying the endogenous substrates of NVL and elucidating the mechanism by which 60S assembly disruption leads to the accumulation of free 5S RNP particles.

Dibenzothiazepinone-based NVL inhibitors are first-in-class, selective inhibitors of ribosome biogenesis. By targeting an essential AAA+ translocase without eliciting a DNA damage response, they provide a molecular foothold for therapeutic development and a tool to dissect the biology of ribosome maturation. Further medicinal chemistry optimization and biomarker discovery will be critical for evaluating NVL inhibition as a viable strategy in cancer therapy.

## ACKNOWLEDGMENTS

We thank the institutionally supported UTSW Preclinical Pharmacology Core for evaluation of the ADME properties of MM17 and bioavailable analogs, Marcel Mettlen at the UTSW Quantitative Light Microscopy Core for help with image analysis, Alejandro Daniel-Olivas at the UTSW Histopathology Core for tissue immunofluorescence staining, the UTSW Medicinal Chemistry Core for synthesizing Oregon green benzylguanine and Alexa fluor 532 azide, and Carlos Arteaga’s laboratory for access to Incucyte. We thank Neha Barrows for technical support in processing resistant clones, and Giomar Rivera Cancel for helpful discussion regarding expression of recombinant NVL. We acknowledge Andrew Boghossian, Matthew Rees, Melissa Ronan, and Jennifer Roth at the Broad Institute of MIT and Harvard for conducting the PRISM screen. We thank Daniel Stoddard and the staff of the UTSW Cryo-Electron Microscopy Facility, funded in part by the CPRIT RP170644. A portion of this research was supported by NIH grant R24GM154185 and performed at the Pacific Northwest Center for Cryo-EM (PNCC) with assistance from Marcelo de Farias. Mass photometry data presented in this report were acquired with a mass photometer supported by NIH award S10OD030312-01. JPE is a Eugene McDermott Scholar in Biomedical Research and a CPRIT Scholar in Cancer Research. This work is supported by the Program in Molecular Medicine, an initiative funded by an Anonymous donor. JKDB holds the Julie and Louis Beecherl, Jr., Chair in Medical Science. DN is a UT Presidential Scholar and holds the Joseph F. Sambrook, Ph.D. Distinguished Chair in Biomedical Science.

## AUTHOR CONTRIBUTIONS

HHG and SX performed ribosome biogenesis inhibition and mode of action studies. HHG, AL, and AA conducted i*n vivo* studies. YT designed and synthesized compounds. VEC performed cryo-EM reconstructions. MF performed compound resistance screens. VK performed CRISPR screen, resistant clone sequencing, and cell line engineering. DR conducted chemical probe crosslinking studies. JK and JP performed bioinformatic analysis. AL, NW, and AA performed pharmacokinetic studies. HHG, VEC, JPE, JKDB, DN wrote the original draft, which is reviewed and edited by all authors.

## DATA AND MATERIALS AVAILABILITY

The cryo-EM maps and protein models have been deposited in the EMDB and the PDB, respectively with the following accession codes: (apo-NVL^dEQ^) EMD-44633 /PDB-9BJI, (NVL^dEQ^ – MM17) EMD-44634 /PDB-9BJJ, (NVL^dEQ^ – MM927) EMD-49941 /PDB-9NYP.

## SUPPLEMENTARY MATERIALS

### MATERIALS AND METHODS

#### Cell culture

The MOLM-13 (a gift from Omar Abdel-Wahab) and HCT116 (ATCC) cell lines were cultured in RPMI-1640 medium (Millipore Sigma #R8758) supplemented with 10% fetal bovine serum (FBS) (Sigma-Aldrich #F2442) and 2 mM L-Glutamine (Sigma Aldrich #G7513), or Dulbecco’s Modified Eagle’s Medium (DMEM) – high glucose (Sigma Aldrich #D5796) supplemented with 10% FBS and 2 mM L-Glutamine, respectively. Cells were cultured at 37°C in 5% CO_2_. The identity of all cell lines was confirmed by short tandem repeat analysis and confirmed to be mycoplasma-free using a PCR based assay (Genatlantis).

#### Lentiviral packaging and stable cell line generation

Lenti-X HEK293T cells (Clonetech #632180) were plated at 500,000 cells per well in a 6-well plate. The next day, cells were transfected with 1 µg of the DNA construct, 900 ng psPAX2 (Addgene #12260), and 100 ng pMD2.G (Addgene #12259) with TransIT-Lenti (Mirus, #MIR6603). At 24 hours, FBS was added to a final concentration of 30%. At 72 hours, lentiviral supernatants were collected and filtered with 0.45 µm PES filter. If not used immediately, lentivirus was aliquoted and stored at −80 °C. Cells were incubated with lentivirus and in the case of HCT116 8 µg/mL polybrene (NEB # H9268) for 18 hr. Following removal of the virus, cells were selected with the relevant antibiotic blasticidin (10 μg/mL), puromycin (2 μg/mL), or hygromycin (10 μg/mL) for at least 7 days.

#### SDS-PAGE and immunoblots

Cells were washed with PBS once before lysis using Buffer A (50 mM HEPES pH 8.0, 10 mM KCl, 2 mM MgCl_2_) with 1% SDS and 20 U/mL benzoase (Sigma-Aldrich # E1014). Protein concentrations were measured and normalized by absorbance at 280 nm using Nanodrop 1000 spectrophotometer (Fisher). Lysates were prepared with 4x Laemmli sample buffer (Bio-Rad #1610747) containing 50 mM 2-mercaptoethanol. Samples were separated on a Bis-Tris gel and transferred to 0.2-micron nitrocellulose membrane using Trans-Blot Turbo transfer system (Bio-Rad). Membranes were stained with Ponceau S (0.1% w/v in 5% acetic acid) followed by wash with 5% acetic acid. Membranes were blocked with 5% milk and incubated with primary antibodies then secondary antibodies: anti-rabbit IgG HRP-linked (Cell Signaling Technologies #7074) or goat anti-mouse IgG HRP-conjugate (Bio-Rad #1706516). Peroxidase signal was visualized with chemiluminescent substrate (Thermo Scientific #34580) and imaged using ChemiDoc (Bio-Rad).

#### Primary Antibodies

The following primary antibodies were used: anti-NVL (Bethyl Laboratories #A304-863A); anti-eL36 (Proteintech #15145-1-AP); anti-eS17 (Abcam #ab128671); anti-p53 (Cell Signaling #9282); anti-p21 (Cell Signaling #2947); anti-Fibrillarin (Cell Signaling #2639); anti caspase-3 (D3R6Y) (Cell Signaling #14220); anti-FLAG m2 peroxidase HRP conjugated (Cell Signaling #a8592); anti-MDM2 (Cell Signaling #86934); anti-SNAP (NEB #P9310S); anti-alpha tubulin HRP conjugated (Cell Signaling #9099); anti-beta actin HRP conjugated (Santa Cruz Biotechnology #47778). For quantitation, gel images were color inverted, and target protein band intensity is measured and normalized to that of loading control using Fiji v2.16 (ImageJ2).

#### Isolation of MM17 resistant clones

10,000 iHCT116 cells/well were plated in five 96 well plates and MM17 was added at the indicated concentrations using a using a D300e Digital Dispenser (Tecan) with a final concentration of DMSO at 0.5%. 17 days after compound addition, viability was determined by incubating with 50 µM glycyl-phenylalanyl-aminofluorocoumarin (provided by the UTSW Medicinal Chemistry Core) for four hours at which point viability was determined by quantifying fluorescence (excitation 380 nm, emission 505 nm) using a Synergy 2 microplate reader (BioTek). Wells with viable cells were manually inspected to confirm clonal growth and expanded and approximately 3 million cells were used to extract genomic DNA, which was used for barcode sequencing and whole exome sequencing according to recently published procedures (*2*).

### Generation of NVL H304R^CRISPR^ and NVL R403W^CRISPR^ cell lines

#### Guide RNA selection using surveyor assay

Guide RNAs were cloned into pX330 vector (Addgene #42230). Lenti-X HEK293T cells (Clontech #632180) were transfected with either pX330 only or with pX330 + guides using TransIT-Lenti (Mirus #MIR 6603). 48 hours after transfection, the cells were collected and genomic DNA was extracted using Zymo genomic DNA extraction kit. The region near H304 was amplified with CloneAmp Hifi PCR mix (Takara Bio #639298) using 5’ TTGTAAGCAAGCTTGTTGTAATATCG and 5’ CTCCTCATGTGGCACTGAAC primers. The region near R403 was amplified with 5’ TAACCATGAAATGATTCTGTAACCA and 5’CCCAGGCATAAATGATTTTCTTAAT. 200ng of PCR product were denatured at 95 °C for 10 min and re-annealed at −1 °C per second ramp to 25 °C. The heterocomplexed PCR product was incubated with 10 units of T7 endonuclease I (NEB, #M0302S) at 37 °C for 1hr. The PCR products were run on 1% agarose gels and analyzed for presence of cleaved products at the expected size.

#### CRISPR/Cas9 focal NVL mutagenesis and knock-in

HCT116 cells were transfected with TransIT-LT (Mirus #MIR2304) according to manufacturer’s protocol with a guide RNA (either mock or NVL targeting) in the px330 vector (Addgene, #42230), and 10 nM single stranded deoxyoligonucleotide encoding a specific NVL mutation (see below for specific sequences). 72 hours after transfection, cells were selected by MM17 treatment as described below. The resulting cells were evaluated by cresyl violet staining and PCR-based amplicon sequencing of the target region. Paired-end sequencing reads were aligned to the human reference genome (hg38) using Burrows-Wheeler Aligner (BWA-MEM, v0.7.17). The aligned reads were sorted and indexed using SAMtools (v1.15.1). Paired-end reads were merged into consensus sequences using a custom Perl script (https://github.com/jiwoongbio/NGS_scripts), which detects overlapping read pairs and reconstructs high-confidence sequences. The merged reads were then realigned to the reference genome with BWA-MEM, followed by sorting and indexing with SAMtools. Variants were identified using SAMtools mpileup, and the detected variants were annotated using the Annomen (https://github.com/jiwoongbio/Annomen), incorporating gene annotations and functional effect predictions based on RefSeq transcript and protein databases. For each mutation category, the number of reads supporting each variant type was counted. Mutation categories included missense, nonsense, frameshift, in-frame, silent, and non-variant reads.

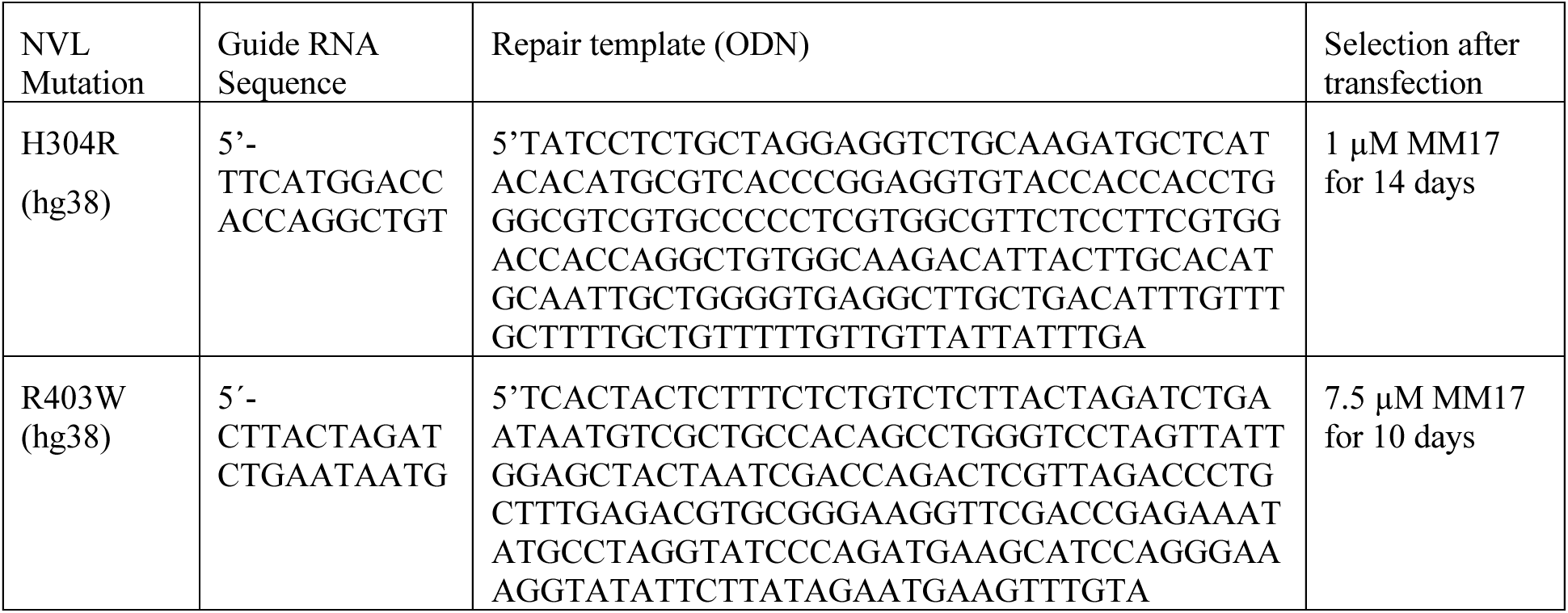

#### Generation of knock-in tags in cell lines

Guide RNA targeting genomic region surrounding the stop codon of the targeted gene was cloned into pX330. The repair template was constructed in a pGEM-T Easy Vector (Promega) to include 1 kilobase homology arms matching upstream and downstream sequences of the genomic locus. The repair template sequence contains the knock-in construct, as well as an IRES Neo cassette flanked by two LoxP sites. Cells were transfected with the guide RNA and the repair template using TransIT LT. Cells were selected with G418 (1 mg/mL) for 2 weeks. Single clones were verified by Western blotting.

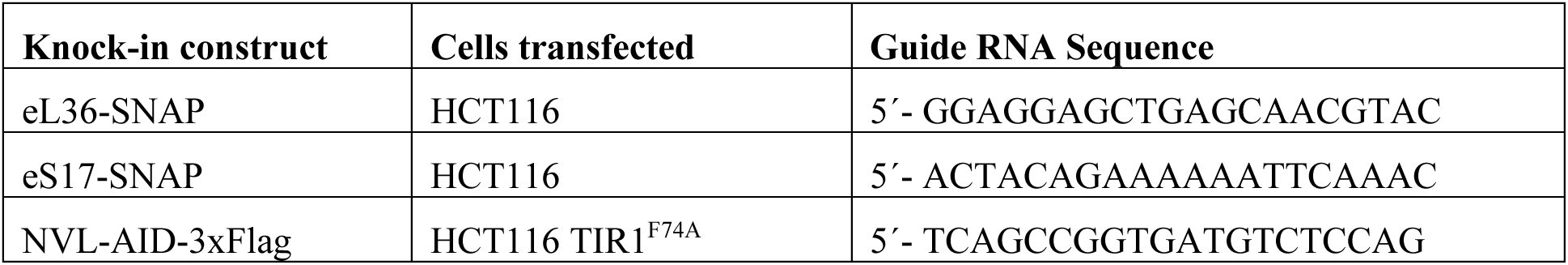

#### CRISPR/Cas9 Lentiviral plasmids cloned into pLenticrisprV2 (Addgene #52961) sequence verified

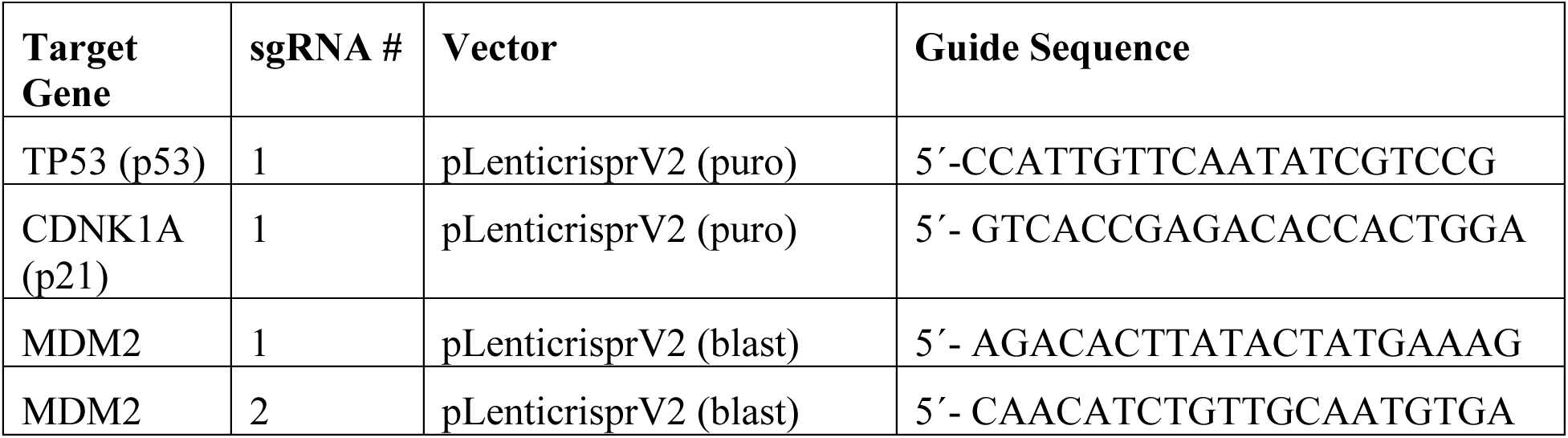
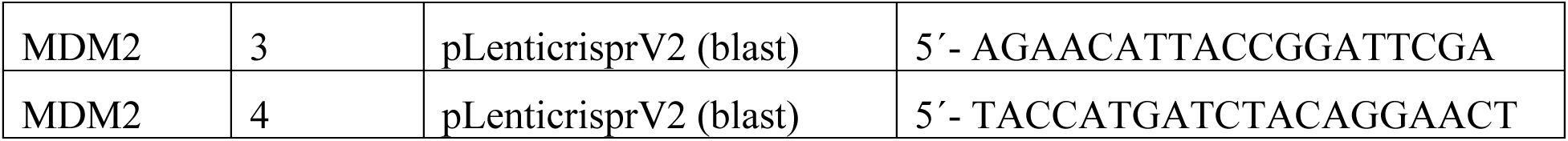

#### Mammalian ORF expression plasmids cloned by Gibson assembly and sequence verified

pLVX IRES Puromycin, Fibrillarin (NM_001436.3) fused to a C-terminal HALO tag

pLVX IRES Blasticidin, Human NVL (NM_002533.4)

pLVX IRES Blasticidin, Human NVL (NM_002533.4) with an R403W mutation

pLVX puromycin, Mouse NVL (NM_026171.2)

pLVX puromycin, Firefly Luciferase (Luc2P)

pLVX IRES Puromycin, MDM2 (NM_002392.6) both wild type and C311F mutation

pLVX IRES Hygromycin ZsGreen

pLVX IRES Hygromycin mCherry

#### Amplicon sequencing of resistant clones

Genomic DNA was extracted using QIAamp DNA Mini Kit (Qiagen # 51304) from resistant clones raised against MM17 and were used as template for amplification of DNA sequence flanking the following regions of NVL.

*Exon 9*: (5’-GATATTGCTGAAGAGCTTGGGTACCAGAGCCAGGATTCA and 5’-AGAGGTTGATTGTTCCAGAAAATGTCAGCAAGCCTCACC)

*Exon 12*: (5’-GATATTGCTGAAGAGCTTGCCATTATTGCCGTCAGTGTC and 5’-AGAGGTTGATTGTTCCAGAAGAGAAAGAAAAGGCTACTTGGT)

*Exon 14*: (5’-GATATTGCTGAAGAGCTTGAACTTTTCAATAGCTGTTGTGGTG and 5’-AGAGGTTGATTGTTCCAGACATGCATGGAGCTTTATCTCTG)

*Exon 18*: (5’-GATATTGCTGAAGAGCTTGTTGAGAGCTGATGGAACCAG and 5’-AGAGGTTGATTGTTCCAGAAAATGGATTGCCCAAATTTAATC)

Indexed PCR products were pooled together and submitted for Amplicon-EZ next generation sequencing (Azenta).

#### Cell viability assay

For dose response assays, cells were plated in 96-well assay plates (Corning # 2903), and compounds suspended in DMSO or DMF were dispensed using a D300e Digital Dispenser (Tecan). All wells were normalized to 0.5% DMSO or DMF. 72 hours after incubation at 37°C in 5% CO_2_, cell viability was assayed using an ATP-based luminescent viability assay, CellTiter-Glo (Promega # G8462) per manufacturer’s instructions. Luminescence was recorded using a Synergy 2 microplate reader (BioTek) or Cytation 5 cell imaging multimode reader (BioTek). In Prism 10 software (GraphPad), ten-point dose response curves were plotted with baseline-correction to the vehicle control, and IC50 values were calculated by curve fitting using variable slope (four parameters) fit.

#### Image-based cell growth assay

Cells plated in 96-well assay plates (Corning # 2903) were allowed to attach overnight and were then incubated in Incucyte (Sartorius). After 3 hours, compounds suspended in DMSO were dispensed using a D300e Digital Dispenser (Tecan), and all wells were normalized to 0.5% DMSO. Cells were imaged every 3 hours during 72 hours of treatment. Average cell area per image (µm^2^) relative that of time 0 was quantified.

#### Cell growth assays

For HCT116, 1.5 million cells were plated in 10-cm plate, and cells were trypsinized and counted every three days after plating. For MOLM-13, 2 million cells / mL were plated in 10-cm plates, and cells were collected and counted every two days after plating. The number of viable cells was counted in technical duplicates using Vi-Cell XR (Beckman Coulter), an automated cell analyzer based on exclusion of trypan blue dye, per manufacturer’s instructions. Cumulative doubling was calculated by 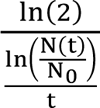, where N(t) = viable cell number at time t, and N_0_ = viable cell number at time 0.

#### Cresyl Violet Staining

The media is removed from the cells are washed with PBS once. Cresyl violet solution (0.5% cresyl violet and 20% methanol) is added to the dish. Following shaking with 5 minutes, the cresyl violet solution is removed, the cells are washed with water twice, and the dish is allowed to dry overnight prior to imaging.

#### Genome-wide CRISPR/Cas9 knockout screen

HCT116 cells stably expressing Cas9 were generated by infecting cells with lentiCas9-Blast (Addgene #52962) followed by selection with 10 µg/mL blasticidin. A Cas9-expressing clone was isolated and expanded to confirm Cas9 protein expression by Western blotting. Lentivirus carrying the human Brunello CRISPR knockout pooled sgRNA library was prepared and provided by Anderson Frank in the McFadden lab. 300 million cells were infected in 8 µg/mL polybrene (Sigma) at a multiplicity of infection of approximately 0.3 and selected with puromycin (2 µg/mL) for 7 days. 150 million cells were treated for 24 hours with either DMSO or 1 µM MM17 and allowed to recover in fresh media without compound until confluency.

Viable cells were counted at each passage and collected after 21 days. Genomic DNA was extracted using Blood & Cell Culture DNA Maxi kit (QIAGEN #13362), and concentrations were determined using Quant-iT PicoGreen dsDNA assay kits (Invitrogen #P7589). To ensure the diversity and coverage of sgRNAs, multiple 100 μL PCR reactions were set up using 5 µg of genomic DNA per condition. sgRNAs were PCR-amplified using 28 cycles from approximately 650 µg genomic DNA (5 µg DNA/100 µL reaction) using Ex Taq DNA Polymerase (TaKaRa #RR001B), 8 staggered P5 primers (12.5 μM), 1 unique P7 index primer (10 µM), and 2.5 mM each of dNTPs per sample. Pooled PCR products were purified using SPRIselect bead-based reagent (Beckman #B23318) per manufacturer’s instructions before sequencing on NextSeq 2000 P2 flowcell (100 cycles, 400M reads) at UTSW McDermott Center Next Generation Sequencing Core. BWA was used to map the sequencing reads to the sgRNA sequences concatenated with 5’ and 3’ flanking regions. The mapped reads were counted for each sgRNA using SAMtools (v1.9), with the options ‘-F 2308 -q 1’. Counts per million (CPM) were calculated as CPM = (Read Count / Total Reads in the Sample) × 10^6^, and log2-transformed CPM (log2CPM) values were obtained using log2(CPM + 1). Log2 fold changes for each sgRNA were calculated as the difference between the log2CPM of control and treatment samples. The log2 fold change for each gene was determined by averaging the log2 fold changes of its associated sgRNAs and subtracting the average log2 fold change of non-targeting sgRNAs.

#### Polysome profiling

Following vehicle (DMSO) or compound treatment, 80% confluent HCT116 cells were incubated with 100 µg/mL cycloheximide (CHX) for 15 minutes and lysed in 5 mM Tris-HCl pH 7.5, 2.5 mM MgCl_2_, 1.5 mM KCl, 0.5% Triton X-100, 0.5% sodium deoxycholate, 100 µg/mL CHX, 1 mM DTT, 100 U RNAase inhibitor (Invitrogen #AM2696), and protease inhibitor (Thermo Scientific #YD3832015). After centrifugation at 16,000*g* at 4 °C for 7 minutes, lysates were collected, and RNA concentrations were normalized by absorbance at 260 nm using Nanodrop 1000 spectrophotometer (Fisher). 350 µL of lysate was loaded on top of a 10-50% (w/v) sucrose gradient containing 20 mM Tris-HCl pH 7.5, 140 mM KCl, 5 mM MgCl_2_•6H_2_O, and 100 µg/mL CHX. Sucrose gradient was made in open-top centrifuge tubes (Seton Scientific #7030) using Gradient Master 108v5.37 (BioComp). Following centrifugation in a SW 41 Ti rotor (Beckman Coulter #333790) at 41,000 rpm at 4°C for 2 hours, absorbance at 260 nm was recorded using Piston Gradient Fractionator 1.2 (BioComp).

#### Polysome fractionation of existing and newly synthesized ribosomes

To label existing 60S or 40S ribosomes, respectively, HCT116 eL36-SNAP or eS17-SNAP cells were incubated with SNAP substrate (500 nM Oregon green or 647-SiR benzylguanine) for 15 minutes. Oregon green benzylguanine (green) was synthesized by UTSW Medicinal Chemistry Core, and 647-SiR benzylguanine (magenta) was from NEB # S9102S. To block any unlabeled ribosomes, 500 nM SNAP-Cell Block (NEB #S9106S) was then added for 15 minutes and removed. Cells were treated with vehicle (DMSO) or 3 µM MM17 for 6 hours. After 6 hours, to label the newly synthesized ribosomes, cells were incubated with a different SNAP substrate (500 nM Oregon green or 647-SiR benzylguanine) and 100 µg/mL cycloheximide (CHX) for 15 minutes. For 60S labeling, two biological replicates used Oregon green benzylguanine to label pre-existing eL36-SNAP and 647-SiR benzylguanine to label newly synthesized eL36-SNAP, while one replicate used the reverse labeling scheme. For 40S labeling, one biological replicate used Oregon green benzylguanine to label pre-existing eL36-SNAP and 647-SiR benzylguanine to label newly synthesized eL36-SNAP, while one replicate used the reverse labeling scheme.

Following polysome profiling (as described above in “Polysome profiling”), 32 fractions were collected, and fractions associated with ribosomes were spun at 200*g* for 5 minutes, and proteins were precipitated with 20 % trichloroacetic acid. Samples were suspended in 25 µL Buffer A (50 mM HEPES pH 8.0, 10 mM KCl, 2 mM MgCl_2_) with 1% SDS and prepared with 4x Laemmli sample buffer (Bio-Rad #1610747) containing 50 mM 2-mercaptoethanol. After boiling at 98°C for 10 minutes, 15 µL of fractioned samples and 2 µL of input lysates were subjected to SDS-PAGE, and the gels were scanned using Typhoon FLA 9000 (GE Healthcare). Green and magenta fluorescence were visualized with 495 nm and 653 nm excitation lasers, respectively.

### Microscopy localization of existing and newly synthesized ribosomes

#### Labeling of existing versus newly synthesized eL36-SNAP (60S) and eS17-SNAP (40S)

To label existing 60S or 40S ribosomes, respectively, HCT116 eL36-SNAP or eS17-SNAP cells plated on coated coverslips (Neuvitro #GG-18-PLL) were incubated with 3 µM Oregon green benzylguanine for 15 minutes. Cells were treated with vehicle (DMSO) or 3 µM MM17 for 24 hours. To label newly synthesized ribosomes, cells were incubated with 1 µM 647-SiR benzylguanine. Cells were then fixed with 4% paraformaldehyde (Fischer #J19943.K2) for 20 minutes, permeabilized with 0.3% Triton X-100 for 15 minutes, and the nucleus were stained with 2 µg/mL DAPI (Fischer # 62247) for 15 minutes. Coverslips were mounted on microscope slides (Sigma #CLS294875X2) using antifade mountant (Fisher #P36984) and sealant (Fisher #P56128). Slides were imaged using 60x oil objective on Nikon CSU-W1 spinning disk confocal microscope at UTSW Quantitative Light Microscopy Core. DAPI, 647-SiR benzylguanine- and Oregon green benzylguanine-conjugated SNAP protein was visualized using DAPI, Cy5, and GFP settings, respectively. The same image acquisition settings, such as laser power and exposure time, were used for all samples. Representative images were magnified 4x, pseudo-colored according to the acquisition color channel, and the same linear brightness and contrast adjustment was used for all conditions using Fiji v2.16 (ImageJ2). Mean nuclear and cytoplasmic fluorescence of Oregon green or 647-SiR benzylguanine conjugated SNAP protein were quantified using a custom Fiji macro “ribosomeLocalization.ijm” (Data S6). In brief, nuclei were first outlined using DAPI fluorescence, and the mean fluorescence of SNAP proteins localized within and outside of the nucleus was quantified as mean nuclear and cytoplasmic fluorescence, respectively. For each sample, at least nine images (304 to 886 cells in total) were quantified.

#### Co-localization of newly synthesized eL36-SNAP (60S) and fibrillarin-HALO (nucleolus)

HCT116 eL36-SNAP fibrillarin-HALO were plated in a 35-mm dish with glass coverslip (MatTek #P35GC-1.5-14-C). Cells were incubated with 250 nM SNAP-Cell Block (NEB #S9106S) for 20 minutes to block existing eL36-SNAP. Following treatment with vehicle or 3 µM MM17 for 24 hours, cells were incubated for 15 minutes with 1 µM 647-SiR benzylguanine (magenta) to label newly synthesized eL36-SNAP and 1 µM Oregon green HALO ligand (green) (Promega #G2801) to label fibrillarin-HALO. Cells were imaged using 100x oil objective on Zeiss LSM 980 laser scanning confocal microscope. Oregon green labeled fibrillarin-HALO and 647-SiR labeled eL36-SNAP was visualized using the GFP and Cy5 settings, respectively. The same image acquisition settings, such as laser power and exposure time, were used for all conditions in the experiment. Representative images were pseudo-colored according to the acquisition color channel, and the same linear brightness and contrast adjustment was used for all conditions using Fiji v2.16 (ImageJ2).

#### γH2AX immunofluorescence staining

Cells were plated on coated coverslips (Neuvitro #GG-18-PLL). Following 24-hour vehicle (DMSO or DMF) or compound treatment, cells were fixed with 4% paraformaldehyde (Fischer #J19943.K2) for 20 minutes at room temperature. Coverslips were incubated with 1:500 anti-phospho-histone H2A.X (Ser 139) antibody (Cell Signaling #2577) diluted in 1% BSA/0.3% Triton X-100/1x PBS overnight at 4 °C, followed by incubation with 1:1000 Alexa fluor 647 conjugated goat anti-rabbit IgG (Cell Signaling #4414) for 1 hour at room temperature, and 2 µg/mL DAPI (Fischer # 62247) for 15 minutes. Coverslips were mounted on microscope slides (Sigma #CLS294875X2) using antifade mountant (Fisher #P36984) and sealant (Fisher #P56128). Slides were imaged using 60x oil objective on Nikon CSU-W1 spinning disk confocal microscope at UTSW Quantitative Light Microscopy Core. Alexa fluor 647 conjugated γH2AX fluorescence was visualized using the DAPI and Cy5 settings, respectively. The same image acquisition settings, such as laser power and exposure time, were used for all conditions in the experiment. Representative images were magnified 3x for, pseudo-colored according to the acquisition color channel, and the same linear brightness and contrast adjustment was used for all conditions using Fiji v2.16 (ImageJ2). Mean γH2AX fluorescence for each nucleus was quantified using a custom Fiji macro “gammaH2AX.ijm” (Data S6). Each individual nucleus was outlined by DAPI fluorescence, and the mean γH2AX fluorescence of each nucleus was measured and plotted. Ten images (254 to 1095 total nuclei) were quantified for each condition.

### Flow cytometry-based cell cycle and apoptosis analysis

#### Propidium iodine (PI) staining

Following vehicle (DMSO) or compound treatment of HCT116 cells, both attached and detached cells were collected and fixed with 70% ethanol at 4°C overnight. Cells were stained with 50 µg/mL PI (Fisher #P1304MP), 100 µg/mL RNAase A (Sigma #R6148), 0.1% Triton X-100 at 37°C for 30 minutes.

#### DAPI and 5-Ethynyl-2’-deoxyuridine (EdU) staining

Following compound treatment, 10 µM EdU suspended in DMSO (Med Chem Express #HY-118411) was added to cells for 2 hours. All attached and detached cells were collected, fixed with 2% paraformaldehyde, and lysed with 0.1% saponin/3% FBS/1x PBS. Lysates were subjected to click reaction with Alexa fluor 532 azide (synthesized by UTSW Medicinal Chemistry Core) using Click-iT cell reaction kit (Fischer # C10269) per manufacturer’s instructions. Cells were stained with 1 µg/mL DAPI (Fischer # 62247) at 4°C overnight.

#### PI and Annexin V staining

Following treatment, MOLM-13 cells were collected and suspended in 100 µL Annexin V binding buffer (BioLegend #422201), 5 µL FITC-Annexin V (BioLegend #640906), and 1 µg/mL PI for 15 minutes at room temperature. 400 µL Annexin V binding buffer was added and samples were kept on ice until analysis.

#### FACS analysis

Stained cells were filtered using a cell strainer (Fisher Scientific #08-771-23). PI, DAPI, EdU-Alexa fluor 532, and FITC Annexin V fluorescence was recorded using PI, DAPI, PE, and FITC settings, respectively, on FACSAria Fusion (BD Biosciences) cytometer at UTSW Children’s Research Institute Moody Flow Cytometry Facility, and data was analyzed using FlowJo v10.10 (BD).

### Recombinant NVL crosslinking assay

#### Cloning of NVL expression constructs

A two-fragment insertional cloning strategy was employed to clone NVL as a fusion with a SNAP tag in a modified pcDNA vector containing an inbuilt 3xFlag. The first fragment - SNAP tag was amplified by PCR using the primers 5’ TTTCACCTGCTTATGCGGCCGCCACCATGGACAAAGACTGCGAAATG 3’ and 5’

ATACACCTGCGCCGCTTCCCCCAGCCCAGGCTTG 3’, while the second fragment – wildtype or R403W mutant NVL was amplified using 5’ GATCACCTGCTTCGGAAGCGGCAGTAAGCCCAGACCTGCAGGG 3’ and 5’

AGTCACCTGCACGCCGGACCGGCTGAGGGACTC 3’ from NVL expression plasmids in pLVX-IRES-Blast vector as described above. Plasmids were sequence verified, and the protein is expressed as SNAP-NVL-3xFlag.

#### UV-crosslinking of MM514 to recombinant NVL

293F suspension cells were cultured in Freestyle 293 media (Gibco # 12338018) at 37°C with 8% CO_2_ and 220 rpm. 30 million cells were transfected with 30 µg of expression plasmids for either wildtype of R403W mutated SNAP-NVL-3xFlag with a PEI reagent (Polysciences #23966). After 48 hours of transfection 5 million cells were seeded per well in a 12 well plate. Compounds suspended in DMSO were dispensed to the wells using a Multi drop Pico 8 Digital Dispenser (Thermo Scientific). All wells were normalized to 0.5% DMSO. Post 1 hour of compound addition the cells were subjected to UV crosslinking at 306 nm for 15 minutes on ice. The cells were then harvested and resuspended in a buffer containing 20 mM HEPES pH 7.5, 150 mM NaCl, 2% CHAPS, 10% Glycerol and 10 mM MgCl_2_ incubated on ice for 10 min. To the supernatant obtained post centrifugation at 15000*g* at 4C, 1 µM of 647-SiR benzylguanine (magenta, NEB # S9102S) was added to label total NVL-SNAP-3xFlag and incubated for 30 min at 37 ° C in the dark after which Flag antibody conjugated magnetic beads were added and the lysate was incubated at 4°C for 15 minutes on a rocker. The beads were washed three times in the binding buffer and eluted with 300 ng/µl of 3xFlag peptide (Sigma) at 4 °C for 30 min. SDS at a final concentration of 1% was added and subjected to click reactions with 100 μM Tris[(1-benzyl-1H-1,2,3-triazol-4-yl)methyl]amine (TBTA) dissolved in 4:1 DMSO:t-butanol (Tokyo Chemical Industry, # T2993), 1 mM freshly prepared sodium ascorbate (Sigma # A7631), 2 mM CuSO_4_ (EM Science # CX21851) and 25 μM Alexa fluor 532 azide (red, synthesized by UTSW Medicinal Chemistry Core) for 1 hour at room temperature in the dark. For competition assays, competitors were dispensed before MM514. Protein samples were subjected to SDS-PAGE, and the gels were scanned using Typhoon FLA 9000 (GE Healthcare). Red and magenta fluorescence were visualized with 532 nm and 653 nm excitation lasers, respectively.

#### O-propargyl-puromycin protein synthesis assay

HCT116 cells were treated with vehicle (DMSO), 10 µg/mL cycloheximide, or 3 µM MM17 at the indicated time points. At the end of the treatment, cells were incubated with or without 10 µM o-propargyl-puromycin (Click Chemistry Tools # 1407) for 2 hours. Lysates were prepared and subjected to click reactions as described in “Recombinant NVL crosslinking assay”. Alexa fluor 532 azide conjugated o-propargyl-puromycin was visualized on a Typhoon FLA 9000 (GE Healthcare) using 532 nm excitation laser. Gels were stained with Coomassie Blue for total protein.

#### Growth competition assay

Equal numbers of HCT116 mCherry (red) mock-engineered cells and ZsGreen (green) p53 KO-1, ZsGreen p53 KO-2, or ZsGreen p21 KO cells were plated in 12-well culture plates. The next day, cells were treated with indicated concentrations of MM17, paclitaxel (Selleckchem #S1150) for 24 hours. At the end of treatment, compounds were removed, and cells were allowed to recover to approximately 80% confluency, at which point they were spilt 1:10. The next day, cell images were taken using 10x objective on Cytation 5 cell imaging multimode reader (BioTek).

The number of red and green cells were quantified from four images per sample using Gen5 3.05 (Biotek). Green and red fluorescent cells were imaged with 469 nm and 586 nm excitation lasers, respectively.

#### Pharmacokinetic studies

Animal work described in this manuscript has been approved and conducted under the oversight of the UTSW Institutional Animal Care and Use Committee. UTSW uses the “Guide for the Care and Use of Laboratory Animals” when establishing animal research standards. Pharmacokinetic studies were performed by dosing 6–7 week old CD-1 female mice with analog MM927 formulated as 10% DMSO/10% Kolliphor EL/10% Solutal/70% D5W via the intraperitoneal route. Animals were dosed in groups of three, blood was obtained by cheek bleeds at each time point using the anticoagulant ACD (acidified citrate dextrose) and plasma isolated by centrifugation. For the dose escalation study, mice were dosed with 20 mg/kg, 35 mg/kg, or 60 mg/kg and blood was collected at various time points post dose. For the efficacy study, blood and tumor tissue was collected from tumor bearing NOD.CB17-*Prkdc^scid^ Il2rg^tm1Wjl^*/SzJ (NSG) mice (Jackson Laboratories,) 3 hours after the final dose of MM927 or vehicle (from 10 mice per group). Aliquots of plasma or tumor tissue homogenate were mixed with a two-fold volume of methanol containing 0.15% formic acid and 37.5 ng/mL N-benzylbenzamide internal standard. The samples were vortexed 15 s, incubated at room temp for 10’ and spun twice at 16,100 x g 4°C in a refrigerated microcentrifuge. The resulting supernatants were evaluated by LC-MS/MSusing a Sciex (Framingham, MA) 4500 Triple Quad coupled to a Shimadzu (Columbia) Prominence LC. Briefly, MM927 was detected in multiple reaction monitoring mode (MRM) using the (m+1) transition 488.059/176.1. N-benzylbenzamide (transition 212.12/91.2) was used as an internal standard. Samples were chromatographed prior to introduction into the mass spectrometer using a C18 column (Agilent XDB column, 5 micron packing, 50 x 4.6 mm) with the following conditions. Buffer A (water + 0.1% formic acid) and Buffer B (MeOH + 0.1% formic acid) were used to create a gradient over the following times: 0-1.5 min 3% B, 1.5-2.0 min gradient to 100% B, 2.0-3.5 min 100% B, 3.5-3.6 min gradient to 3% B, 3.6-4.5 min 3%B. Standard curves were generated using naïve tumor tissue or blank plasma (Bioreclamation) spiked with known concentrations of analog MM927 and processed as described above. The concentrations of drug in each time-point sample were quantified using Analyst software (Sciex). A value of 3-fold above the signal obtained from blank plasma was designated the limit of detection (LOD). The limit of quantitation (LOQ) was defined as the lowest concentration at which back calculation yielded a concentration within 20% of theoretical and which was above the LOD. Cmax and AUC values were calculated using the noncompartmental analysis tool of Phoenix WinNonLin (Certara/Pharsight).

#### Mouse liver microsome stability assay

Levels of MM17 were monitored by LC-MS/MS using an AB Sciex (Framingham, MA) TripleQuad™ 4500 mass spectrometer coupled to a Shimadzu (Columbia, MD) Prominence LC. The compound was detected with the mass spectrometer in positive MRM (multiple reaction monitoring) mode by following the precursor to fragment ion transition 395.05/363.0. Buffer A (Water + 0.1% formic acid/2mM NH_4_ Acetate) and Buffer B (Methanol + 0.1% formic acid/2mM NH_4_ Acetate) were used to create a gradient over the following times: 0 - 1.0 min 5% B, 1.0 - 2.0 min gradient to 95% B, 2.0 - 3.0 min 95% B, 3.0 - 3.2 min gradient to 5% B, 3.2 - 4.5 5% B with a flow rate of 1.5 mL/min using an Agilent C18 XDB column (5 micron, 50 × 4.6 mm). N-benzylbenzamide (transition 212.1 to 91.1 from Sigma (St. Louis, MO) or tolbutamide (transition 271.1 to 91.1 also from Sigma) was used as an internal standard (IS). Levels of MM927 was monitored by LC-MS/MS as described in “Pharmacokinetic studies”. Male ICR/CD-1 mouse microsome fractions (lot OYJ) were purchased from BioIVT (Westbury, NY). Microsome protein (0.5 mg/mL) was placed in a glass screw cap tube; a 2 mM DMSO compound stock was spiked into a 50 mM Tris, pH 7.5 solution and this was added to the microsome solution on ice. The final concentration of compound after addition of all reagents was 2 µM. An NADPH-regenerating system (1.7 mg/mL NADP, 7.8 mg/mL glucose-6-phosphate, 6 U/mL glucose-6-phosphate dehydrogenase in 2% w/v NaHCO_3_/10 mm MgCl_2_) was added for analysis of Phase I metabolism after heating both the regenerating solution and the sample tubes to 37°C for 5 min a shaking water bath. The incubation was continued and at varying time points after addition of phase I cofactors, the reaction was stopped by the addition of 0.5 mL methanol containing IS, formic acid, and NH_4_ acetate with final concentrations of 100 ng/mL, 0.1%, and 2 mM respectively. Time 0 samples were stopped with the methanol solution while still on ice prior to addition of the NADPH regenerating system and compound, which were subsequently added. The samples were incubated 10’ at room temperature and then spun at 500*g* in a tabletop centrifuge. The supernatant was spun a second time at 16,100*g* for 5 min in a microcentrifuge at 4°C and analyzed by LC-MS/MS. A “% remaining” value was used to assess metabolic stability of a compound over time. The LC-MS/MS peak area of the incubated sample at each time point was divided by the LC-MS/MS peak area of the time 0 (T0) sample and multiplied by 100. The natural Log (LN) of the % remaining of compound was then plotted versus time (in min) and a linear regression curve plotted going through y-intercept at LN(100). The half-life (T ½) was calculated as T½ = 0.693/slope.

#### Pharmacodynamic studies

Five million HCT116 NVL WT or HCT116 NVL R403W^CRISPR^ in a volume of 0.1 mL were implanted using a subcutaneous injection in the left and right side of four-to-eight-week-old NOD.CB17-*Prkdc^scid^ Il2rg^tm1Wjl^*/SzJ (NSG) mice, respectively. Tumor volumes were measured every 2 to 4 days with calipers. When the tumor diameter reached 0.5 cm, mice were dosed with 35 mg/kg MM927 or vehicle formulated as 10% DMSO/10% Kolliphor EL/10% Solutal/70% D5W and administered via the intraperitoneal route.

#### Immunoblot analysis of p53 and p21

Tumor-bearing mice were given one dose of vehicle or MM927, and tumors were harvested and flash frozen at indicated time points. Tumors were grinded using a pellet pestle (Fisher Scientific # K749521-1590) in Buffer A (50 mM HEPES pH 8.0, 10 mM KCl, 2 mM MgCl_2_) with 1% SDS and 20 U/mL benzoase (Sigma-Aldrich # E1014), and immunoblotting was performed as described in “SDS-PAGE and immunoblots”.

#### Immunofluorescence staining of p53 and Ki-67

Tumor-bearing mice were treated with vehicle or MM927 twice daily for 3 days, and tumors were harvested 6 hours after the last dose and fixed in 4% paraformaldehyde for 48 hours. Fixed tumors were paraffin processed, embedded, and sectioned at UTSW Histopathology Core. In brief, samples were dehydrated through 7 increasing grades of ethanol, cleared through 3 exchanges of xylene and infiltrated with 3 exchanges of paraffin wax (Paraplast Plus, Leica Microsystems) with vacuum-assist on a Thermo-Shandon Excelsior Tissue Processor (Kalamazoo). Embeds to obtain tumor anatomy inclusive of capsule contiguous to core were prepared on a Sakura Finetek Tec 6 Embedding Center (Torrance). Paraffin sections were concomitantly prepared and checked by dark-field microscopy to ensure quality without rarefaction. One MM927 treated wildtype tumor was excluded due to improper fixation. A battery of 10 slides was prepared serially from each block with a second step-level applied to each slide at a 200µm deeper interval from the first section. Multiplex immunofluorescence was done for Ki-67 (MM1 clone, Leica) and p53 (7A5 clone, Cell Signaling). In brief, multiplex immunofluorescence sections were deparaffinized, and antigens retrieved by heating in pH 6.0 citrate-buffer, and sections were blocked against endogenous mouse IgG and secondary antibody host-serum affinity utilizing commercially available blocking reagents (Vector Mouse on Mouse “MOM” Kit and Normal Goat Serum, Vector Laboratories). Sections were subjected to primary antibodies (Ki-67 at 1:150 and p53 at 1:250) and incubated overnight at 4°C. Subsequent biotin/streptavidin-fluorescein detection of bound Ki-67 primary was conducted the following day according to MOM kit instructions and detection of bound p53 primary was done with Alexa fluor 647 goat anti-rabbit (Cell Signaling # 4414). Nuclei were counterstained with Hoechst 33342 (Fisher), and coverslipping performed using Vectashield (Vector Laboratories). Slides were imaged using 60x oil objective on Nikon CSU-W1 spinning disk confocal microscope at UTSW Quantitative Light Microscopy Core. Hoechst 33342, Alexa fluor 647 conjugated p53, and fluorescein conjugated Ki-67 were visualized using DAPI, Cy5, and GFP settings, respectively. The same image acquisition settings, such as laser power and exposure time, were used for all samples. Representative images were magnified 3x, pseudo-colored according to the acquisition color channel, and the same linear brightness and contrast adjustment was used for all conditions using Fiji v2.16 (ImageJ2). p53 positive, Ki-67 positive, and double positive cell percentages were quantified using a custom script in QuPath (v0.5.1) named “p53_Ki67_quantitation.groovy” (Data S6). In brief, the number of cells was quantified by the number of nuclei defined by DAPI fluorescence. Cells stained positive for p53 and/or Ki-67 were manually classified on two representative training images from each condition, and the classifier setting was recorded in “Classifier.json” (Data S6). The classifier was then used to quantify the entire set of images. For each tumor, at least seven images (2 to 3 tissue sections, 3763 to 8170 cells in total) were quantified.

### Efficacy studies

#### HCT116 xenograft models

Five million parental HCT116 cells or HCT116 NVL R403W^CRISPR^ in a volume of 0.1 mL were implanted in the right flank of NSG mice. Tumor volumes were measured every 2 to 4 days with calipers. Volume is calculated as [(L * W^2^)*3.14]/6. Ten days after tumor implantation when volumes reached ∼110 mm^3^, treatment was started with 35 mg/kg MM927 or vehicle (10% DMSO/10% Kolliphor EL/10% Solutal/70% D5W) (0.2 mL/mouse) twice a day for 21 days.

Mice were euthanized at the end of the 21-day treatment period or when any tumor reached 2 cm in its largest diameter, in agreement with the approved animal protocol. At the end of treatment, blood and tumor tissue were collected for pharmacokinetic analysis, measurement of complete blood counts, and blood chemistry measurements as described. Complete blood count was analyzed by UTSW Animal Resource Center. Serum AST, ALT, and BUN were analyzed by UTSW Metabolic Phenotyping core using VITROS MicroSlide technology.

#### MOLM-13 xenograft models

MOLM-13 cells were engineered to stably express firefly luciferase, enabling the quantitation of leukemic burden by bioluminescence imaging in live mice. 1 x 10^6^ MOLM-13-luc cells were injected intravenously into the tail vein of NSG mice. Mice were treated with 35 mg/kg MM927 or vehicle control (0.2 mL/mouse) twice a day for 14 days starting 15 days after transplantation at which point all mice had detectable disease by bioluminescence imaging. To monitor tumor burden, five minutes before imaging, mice were injected intraperitoneally with 100 μL of PBS containing D-luciferin monopotassium salt (40 mg/mL). Mice were anesthetized with isoflurane 5 min before imaging. All mice were imaged using an IVIS Lumina III (Revvity Perkin-Elmer) with Living Image software (Revvity). To measure the background luminescence, a negative control mouse not transplanted with MOLM-13-luc cells was imaged. The bioluminescence signal (total photon flux) was quantified with region of interest measurement tools in Living Image software. Leukemic disease burden was calculated as observed total photon flux in xenografted mice minus background total photon flux in negative control mice. Negative values were set to 1 for purposes of statistical analysis. Endpoint analysis was performed 29 or 30 days after transplantation and leukemic disease burden quantified by bioluminescence imaging (day 29) or flow cytometry analysis of leukemic cells present in the peripheral blood or bone marrow (day 30). The bone marrow was flushed from tibias and femurs using staining medium (Hanks Buffered Saline Solution supplemented with 2% bovine serum) and was dissociated into a single-cell suspension by gently triturating with a 23-gauge needle and then filtering through a 40 μm cell strainer. Blood was collected by cardiac puncture and subjected to ammonium chloride–potassium chloride red cell lysis. The cells were counted and subsequently stained with Alexa Fluor 488 mouse anti-human CD45 (BD Pharmingen) at 4°C for 30 minutes. Dead cells were identified by staining with DAPI (4′,6-diamidino-2-phenylindole) after antibody staining. Cells were analyzed using a FACSAria Fusion (BD Biosciences) cytometer and the percentage of MOLM13-luc cells were analyzed using FlowJo v10.10 (BD) as the percentage of DAPI negative (live) cells that were human CD45 positive within each sample.

#### Analysis of PRISM Screen Cell Line Response Data

MM17 was evaluated by The PRISM Lab at the Broad Institute. The log2-fold change in cell viability (compared to vehicle) at an MM17 concentration of 10 µM was employed for all analyses. the analysis of cell line response to MM17 with respect to p53 genotype, we used the Broad Institute DepMap 22Q2 release data files (CCLE_gene_cn.csv & CCLE_mutations_bool_hotspot.cvs). Two p53 analyses were performed, using gene copy number and the presence of hotspot p53 mutations. For the copy number analysis, the DepMap gene copy number was categorized as either deleted or not based on whether the copy number was less than the arbitrary cutoff of 0.8. For comparison, the Broad PRISM Repurposing 20Q2 data set was used (secondary-screen-logfold-change.csv). Because many compounds are represented in the secondary screen data set at more than one concentration, we selected a single treatment for each compound, preferentially choosing concentrations close to 10 µM. The correlation between the log2-fold change of the series of cell lines in the presence of MM17 was calculated vs the log2-fold change of the same series of cell lines in the presence of the compounds in the secondary screen using a R source file (Data S5).

#### Overexpression and purification of NVL and NVL^dEQ^

The D1/D2 core of human NVL (residues 254-857) was codon optimized for *E. coli* expression and cloned into a custom pET28-based vector with an N-terminal 14xHis tag (MSKHHHHSGHHHTGHHHHSGSHHHTG), followed by a NEDD8 tag from *B. distachyon* (*67*). Catalytically inactive NVL^dEQ^ was generated by introducing point mutations in the Walker B motifs of D1 (E306Q) and D2 (E634Q) via Gibson assembly. All constructs were sequence-verified. Plasmids were transformed into BL21(αDE3) LOBSTR cells (Kerafast). Cultures were grown overnight at 37 °C and used to inoculate 12 L LB medium at 1:200 dilution. At OD_600_ of 0.6 – 0.8, the temperature was reduced to 18° C, and protein expression was with 200 µM IPTG for 16 h. Cells were harvested by centrifuging (4000 x *g*, 30 min) and resuspended in buffer B1 (50 mM HEPES, pH 8.0, 800 mM KOAc, 5 mM MgOAc_2_, 40 mM imidazole, pH 7.0, 1 mM TCEP, 10% Glycerol) supplemented with PMSF and DNaseI. Cells were lysed by sonication (Branson; three 5s pulses at 30 % with 30s pauses) and cleared by centrifugation (18,000*g*, 30 min). Proteins were purified on an AKTA Pure system using a tandem affinity-ion exchange protocol. Cleared lysate was applied to a 5 mL Ni-NTA column HisTrapFF; Cytiva), washed with 5 column volumes (CV) each of buffer B1 followed and buffer A1 (identical to B1 but with 50mM KOAc). The Ni elution was directly applied to a HiTrapQ ion-exchange column (Cytiva) with buffer B2 (same as A1 but with 400mM imidazole). Protein was then eluted from the Q-column with a linear gradient buffer A1 to B1. Peak fractions were pooled, avoiding later fractions containing DnaK (identified via mass spectrometry), and concentrated to 1 mL using a 30 kDa cutoff filter (Amicon ultra, Millipore). The protein was further purified by size exclusion chromatography on an Superdex 200 Increase 10/300 column (Cytiva) equilibrated in gel filtration buffer (50 mM HEPES, pH 8.0, 200 mM KOAc, 5 mM MgOAc_2_, 1 mM TCEP pH 7.0, 0.01 % NP-40). Peak fractions were pooled, concentrated to ∼20 mg/mL, and flash frozen in liquid nitrogen.

#### Mass photometry

Mass photometry (MP) measurements were performed on purified NVL2^dEQ^ in the presence or absence of 10 µM MM17. Protein samples were diluted to 100 nM in filtered MP buffer (50 mM HEPES, pH 8.0, 200 mM KOAc, 5 mM MgOAc_2_, 1 mM TCEP, pH 7.0). Where indicated, 10 µM MM17 was included. All measurements were performed on a Refeyn TwoMP instrument.

Calibration was conducted using standards ranging from 66 kDa (bovine serum albumen) to 660 kDa (thyroglobulin). Microscopy wells were prepared with freshly cleaned glass slides and 8-well silicone gaskets. For each acquisition, 16.2 µL of MP buffer was added to bring the sample into focus under native settings. Then, 1.8 µL of protein sample was mixed into the droplet and a 1-minute video was acquired using AcquireMP software. Videos were processed and analyzed with DiscoverMP (Refeyn).

#### Cryo-EM Grid preparation

Purified NVL and NVL^dEQ^ were diluted to 1 mg/mL in gel-filtration buffer prior to vitrification. For apo NVL^dEQ^, the bdNEDD8 tag was cleaved using bdNED protease at a 1:1000 (w/w) ratio for 1 hour at 4 °C to mitigate preferred particle orientation. For NVL^dEQ^ in complex with MM17 or MM927, compounds were prepared as 10 mM stocks in DMSO and diluted 10-fold into GF buffer for a 2.5% final DMSO concentration. Compound stocks were then added at a 1:10 ratio (100 µM final concentration). In the presence of compound, preferred orientation was not observed, and the bdNEDD8 tag was not removed. For vitrification, 3.5 µL of sample was applied to glow-discharged (30V, 30 s) Quantifoil R2/1 300-mesh copper grids coated with continuous-carbon. Grids were blotted with a 15 s wait time, blot force of 3, and blot time 4.5 s at 4 °C and 100 % using a Mark IV Vitrobot (Fisher), then plunge-frozen into liquid ethane.

#### Cryo-EM Data collection

Micrographs for apo-NVL^dEQ^, NVL^dEQ^-MM17, and NVL^dEQ^-MM927 were acquired on a Titan Krios transmission elecron microscope (Fisher) operating at 300 kV and equipped with a Gatan K3 direct electron detector using a GIF-Quantum energy filter (slit width, 20 eV, apo-NVL^dEQ^, NVL^dEQ^-MM17 or 10 eV, NVL^dEQ^-MM927). Automated data collection was performed using SerialEM (*68*) with a defocus range of 0.6 – 1.9 µm. For apo-NVL^dEQ^ and NVL^dEQ^-MM17, micrographs were collected at a nominal magnification of 105,000x (physical pixel size, 0.83 Å) and dose fractionated into 40 frames of 0.05 s per frame, with a dose rate of 20.1 e^-^/pixel/s. The total exposure time was 2 s, resulting in a cumulative dose of 40.2 e^-^/pixel. For NVL^dEQ^-MM297, micrographs were acquired at the same nominal magnification (pixel size, 0.827 Å) and dose fractionated into 30 frames of 0.067 s pre frame, with a dose rate of 21.6 e^-^/pixel/s. The total exposure time was 2 s, corresponding to a cumulative dose of 43.1 e^-^/pixel.

#### Cryo-EM data processing

All datasets were acquired as LZW compressed .tif files and processed using RELION 4.0 (*69*, *70*). Motion correction was performed with MotionCor2 (*71*), and contrast transfer function (CTF) parameters were estimated using GCTF (*72*). CTF power spectra were manually inspected for clear Thon rings and absence of ice contamination, and only those meeting these criteria were selected for further processing. Approximately 1,000 particles were manually picked, ensuring representative top and side views, and used to generate initial 2D templates for reference-based autopicking. Particles were then extracted and binned fourfold for 2D classification using the EM algorithm. Classes displaying clear secondary structure features in both top and side views were selected for 3D classification. Reference maps for 3D classification were low-pass filtered to 20 Å; initially the 2.8Å Rix7 map (EMD-25474 (*41*)) was used, and subsequently, NVL maps generated in this study served as references. Selected 3D classes were inspected in Chimera, and particles were extracted at the full resolution (un-binned) for 3D refinement with no imposed symmetry (C_1_). To enhance local map quality for subunits E and F, several rounds of local classification were performed using various masks around subunits D/E and F to subtract densities outside the mask. 3D classification without particle alignment was applied to these densities to isolated particle sets with the clearest features in subunits E and F. These subsets were unmasked, 3D refined, followed by CTF refinement, particle polishing, a second round of CTF refinement and a final round of 3D refinement and postprocessing in RELION 4.0 (*69*, *70*). The final resolutions were estimated using the gold-standard Fourier shell correlation (FSC = 0.143): NVL ^dEQ^-MM17 at 3.05 Å from 149 567 particles; apo-NVL^dEQ^ at 2.83 Å from 146,455 particles and NVL^dEQ^-MM297 at 2.86 Å from 193,335 particles. Local resolution maps were generated using the ResMap wrapper within RELION (*73*).

#### Model building and refinement

An AlphaFold 2 (*74*) -predicted structure of NVL was trimmed to remove the N-terminal domain and unresolved loops, in agreement with the construct boundaries used for structural studies.

Individual D1 and D2 domains for each of the 6 subunits were docked into the cryo-EM density as rigid bodies using ChimeraX (*75*). Further model building and adjustments were performed manually using Coot (*76*). Clear polypeptide chain density was observed within the axial pore of NVL, consistent with prior Rix7 structures. Although bulky side chains were resolved, a specific sequence could not be assigned. Because Rix7 is known to capture the N-terminus of one of its subunits, we interpret the peptide as belonging to one of the N-terminal 14x His-tag and modeled the density as poly-histidine assuming multiple averaged registers. Waters surrounding the magnesium in the ATP binding pockets were added manually while bulk waters in the core of apo-NVL were added using phenix.douse with default settings. Final models were refined using phenix.real_space_refine (*77*). Molecular graphics were generated using Chimera X (*75*) and PyMOL (Version 2.0, Schrödinger).

#### Synthesis of MM17, probe analog MM514, probe binding competitor analogs MM524 and MM691, *in vivo* study analog MM927

Reagents and conditions: (a) 2-methoxyethanamine, HATU, DIPEA, DMF, rt; (b) 1,2-difluoro-4-nitrobenzene, K_2_CO_3_, DMF, 70 °C; (c) Fe, NH_4_Cl, MeOH/H_2_O, reflux; (d) aryl carboxylic acid, HATU, DIPEA, DMF, rt; (e) BBr_3_, DCM, −78 to 0 °C; (f) 3-bromoprop-1-yne, NaH, DMF, 0 °C to rt; (g) step 1: HCl, NaNO_2_, H_2_O, 0 °C; step 2: Na_2_S_2_, H_2_O, 0 °C; (h) but-2-yn-1-amine, HATU, DIPEA, DMF, rt; (i) NaBH_4_, MeOH, 0 °C – rt. DCM, dichloromethane; DIPEA, diisopropylethylamine; DMF, dimethylformamide; HATU, 1-[bis(dimethylamino)methylene]-1*H*-1,2,3-triazolo[4,5-*b*]pyridinium 3-oxide hexafluorophosphate.

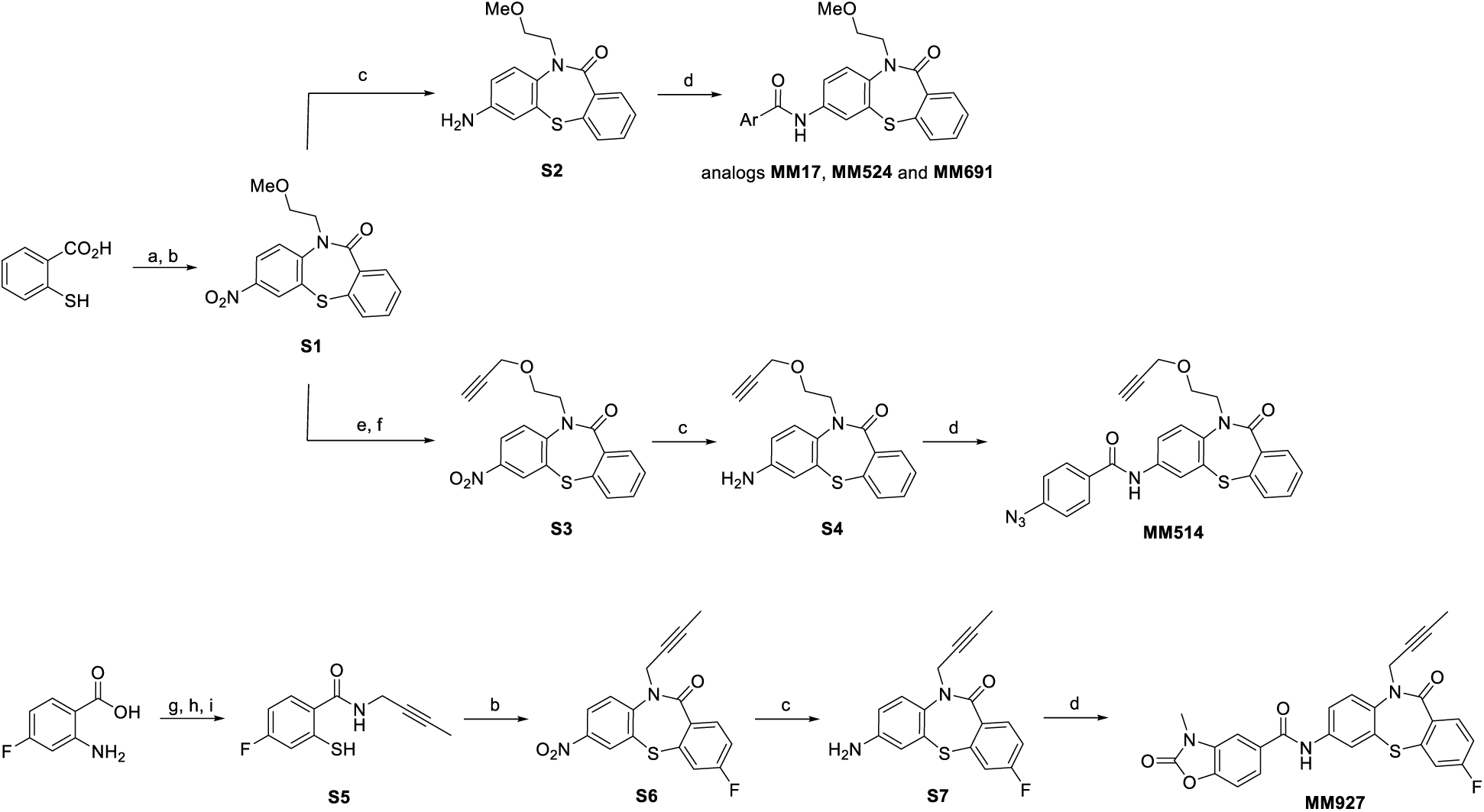

#### 10-(2-Methoxyethyl)-7-nitrodibenzo[b,f][1,4]thiazepin-11(10H)-one (**S1**)

A mixture of 2-mercaptobenzoic acid (771 mg, 5.0 mmol), HATU (2.85 g, 7.5 mmol) and DIPEA (1.30 mL, 7.5 mmol) in DMF (20 mL) was stirred at room temperature for 15 min. 2-Methoxyethanamine (0.65 mL, 7.5 mmol) was added in one portion and the resulting mixture was stirred at room temperature overnight. 1N aq. HCl was added to quench the reaction, followed by extraction with ethyl acetate (3 times). The combined organic phase was dried over Na_2_SO_4_, filtered and concentrated under reduced pressure to give a crude yellow oil used as such in the next step. 1,2-Difluoro-4-nitrobenzene (0.23 mL, 2.08 mmol) and K_2_CO_3_ (717 mg, 5.19 mmol) were added portion-wise with constant stirring to a solution of the above yellow oil (366 mg, 1.73 mmol) in anhydrous DMF (5 mL). The reaction mixture was stirred for 12 h at 75 °C. Then the reaction mixture was diluted with 100 mL of water and extracted with ethyl acetate. The organic layers were washed with saturated brine and dried over Na_2_SO_4_. The organic phase was evaporated under reduced pressure to give the crude product, which was purified by flash chromatography on silica gel (hexane/EtOAc = 20:80) to afford the product **S1** (239 mg, 80% yield over two steps) as a yellow solid. ^1^H NMR (400 MHz, Chloroform-d) δ 8.47 (d, *J* = 2.6 Hz, 1H), 8.17 (dd, *J* = 8.9, 2.7 Hz, 1H), 7.78 (d, *J* = 8.9 Hz, 1H), 7.72 – 7.63 (m, 1H), 7.50 – 7.43 (m, 1H), 7.41 – 7.29 (m, 2H), 4.44 – 4.33 (m, 1H), 4.17 – 4.07 (m, 1H), 3.89 (ddd, *J* = 10.6, 6.8, 3.9 Hz, 1H), 3.75 – 3.65 (m, 1H), 3.34 (s, 3H). ^13^C NMR (100 MHz, Chloroform-d) δ 168.5, 150.2, 144.6, 137.9, 137.4, 137.0, 131.4, 131.3, 131.1, 129.2, 128.2, 127.0, 124.7, 70.5, 59.0, 52.7.

#### 7-Amino-10-(2-methoxyethyl)dibenzo[b,f][1,4]thiazepin-11(10H)-one (**S2**)

To a solution of *S1* (204 mg, 0.618 mmol) in MeOH/H_2_O (8:2 mL) was added NH_4_Cl (562 mg, 10.5 mmol) and Fe (102 mg, 1.854 mmol). After reflux for 2 h, the reaction mixture was filtered through celite and washed with MeOH. The collected filtrate was evaporated under vacuum to give the crude yellow solid product which was pure enough to be used in the next step.

#### 7-Nitro-10-(2-(prop-2-yn-1-yloxy)ethyl)dibenzo[b,f][1,4]thiazepin-11(10H)-one (**S3**)

To a stirred solution of *S1* (85 mg, 0.26 mmol) in DCM (5 mL) under N_2_ at −78 °C, was added boron tribromide (1 M in DCM, 0.64 mL, 0.64 mmol). The reaction was allowed to slowly warm to RT and stirred for 12 h. The reaction mixture was quenched with H_2_O and extracted with ethyl acetate (3 times). The combined organic phase was dried over Na_2_SO_4_ and concentrated under reduced pressure to give the crude product that was used as such in the next step. To an ice cooled solution of the above oil (11 mg, 0.035 mmol) in dry THF (1 mL) was added sodium hydride portionwise (60% in mineral oil, 2.8 mg, 0.07 mmol). The reaction mixture was stirred at 0 °C for 30 mins after which 3-bromoprop-1-yne (80% in toluene, 6.0 µL, 0.053 mmol) was added dropwise. The temperature was slowly raised from 0 °C to room temperature overnight.

The reaction mixture was quenched with saturated aqueous NH_4_Cl and extracted with EtOAc. The combined organic phase was dried over Na_2_SO_4_ and concentrated under reduced pressure to give a crude product that was purified by flash chromatography on silica gel using a mixture of EtOAc in hexanes (10 to 20% gradient) to afford the title compound as a yellow oil (71% yield). ^1^H NMR (400 MHz, Chloroform-d) ^1^H NMR (600 MHz, Chloroform-d) δ 8.47 (d, *J* = 2.6 Hz, 1H), 8.17 (dd, *J* = 8.9, 2.7 Hz, 1H), 7.76 (d, *J* = 8.9 Hz, 1H), 7.69 – 7.65 (m, 1H), 7.49 – 7.44 (m, 1H), 7.39 – 7.32 (m, 2H), 4.45 (ddd, *J* = 14.3, 6.6, 3.8 Hz, 1H), 4.18 – 4.13 (m, 3H), 4.04 (ddd, *J* = 10.3, 6.6, 3.8 Hz, 1H), 3.86 (ddd, *J* = 10.1, 6.1, 3.9 Hz, 1H), 2.41 (t, *J* = 2.4 Hz, 1H). ^13^C NMR (150 MHz, Chloroform-d) δ 168.5, 150.1, 144.6, 137.8, 137.5, 137.1, 131.4, 131.3, 131.1, 129.2, 128.2, 126.9, 124.7, 79.2, 74.8, 67.9, 58.5, 52.5.

#### 7-Amino-10-(2-methoxyethyl)dibenzo[b,f][1,4]thiazepin-11(10H)-one (**S4**)

To a solution of *S3* (7.2 mg, 0.02 mmol) in MeOH/H_2_O (4:1 mL) was added NH_4_Cl (18.5 mg, 0.345 mmol) and Fe (3.4 mg, 0.061 mmol). After reflux for 2 h, the mixture was filtered through celite and washed with MeOH. The collected filtrate was evaporated under vacuum to give the crude product which was pure enough to be used as such in the next step.

#### 10-(But-2-yn-1-yl)-3-fluoro-7-nitrodibenzo[b,f][1,4]thiazepin-11(10H)-one (**S6**)

2-Amino-3-fluoro-benzoic acid (20.0 g, 129.0 mmol) in water (60 mL) with concentated HCl (24 mL) was cooled to 5 °C. A cold solution of sodium nitrite (9.8 g, 129.0 mmol) in water (40 mL) was added dropwise at 5 °C. After addition, the mixture was stirred for 30 min at same temperature. A solution of disodium disulfide prepared with boiled water (40 mL), Na_2_S•9H_2_O (34.4 g, 143.0 mmol), sulfur (4.6 g, 143.0 mmol) and a solution of NaOH (5.42 g, 135.0 mmol) was cooled to 5°C and added dropwise into the above mixture at 5 °C. After being stirred at room temperature for 2 h, the mixture was acidified with HCl. The corresponding disulfide derivative was filtered, washed with water and dried in vacuum. The crude product was purified by Prep-HPLC to give product as yellow solid.

A mixture of above yellow solid (3.0g, 17.4 mmol), HATU (9.93 g, 26.2 mmol) and DIPEA (9.6 g, 26.2 mmol) in DMF (50 mL) was stirred at room temperature for 15 min, but-2-yn-1-amine (2.8 g, 26.2 mmol) was added in one portion. The resulting mixture was stirred at room temperature overnight. 1N aq. HCl was added to quench the reaction and the residue was partitioned between ethyl acetate and H_2_O. The organic phase was dried over Na_2_SO_4_, filtered, concentrated afford crude used for next step directly.

To a suspension of above crude oil (19.4 mmol) in MeOH (100 ml) was added NaBH_4_ (1.47 g 38.7 mmol) at 0 °C, then the mixture was stirred at room temperature for 30 min, the solvent was removed and acidified with 2 N HCl to pH = 4. The residue was diluted with ethyl acetate, the aqueous phase was lyophilized to give 4.25 g crude product, which was used as such for the next step.

To a solution of the above intermediate (4.25 g, 19.1mmol) in anhydrous DMF (80 mL) Cwas added .1,2-Difluoro-4-nitrobenzene (3.05 g, 19.1 mmol) and K_2_CO_3_ (7.89 g, 57.2 mmol). The reaction mixture was stirred for 12 h at 75 °C. Then the reaction mixture was diluted with 100 mL of water and extracted with ethyl acetate. The organic layer was washed with saturated brine, dried over Na_2_SO_4_, filtered, concentrated and purified by flash chromatography on silica gel (Hexane/EtOAc = 1:3) to afford product ***S6*** (2.0 g, 33.6 % yield over three steps) as a yellow solid. ^1^H NMR (600 MHz, Chloroform-d) ^1^H NMR (600 MHz, Chloroform-d) δ 8.48 (d, *J* = 2.6 Hz, 1H), 8.24 (dd, *J* = 8.9, 2.6 Hz, 1H), 7.92 (d, *J* = 8.9 Hz, 1H), 7.81 (dd, *J* = 8.7, 5.7 Hz, 1H), 7.21 (dd, *J* = 8.0, 2.3 Hz, 1H), 7.08 (td, *J* = 8.8, 8.4, 2.6 Hz, 1H), 5.00 (dd, *J* = 17.1, 2.3 Hz, 1H), 4.45 (dd, *J* = 17.2, 2.4 Hz, 1H), 1.86 (t, *J* = 2.4 Hz, 3H). ^13^C NMR (151 MHz, Chloroform-d) δ 166.7, 164.5, 162.8, 149.3, 144.8, 139.2 (d, *J* = 8.6 Hz), 135.0, 133.2 (d, *J* = 3.4 Hz), 128.3, 124.9, 118.3, 118.1, 116.7, 116.6, 81.3, 73.9, 41.4, 3.8.

#### 7-Amino-10-(but-2-yn-1-yl)-3-fluorodibenzo[b,f][1,4]thiazepin-11(10H)-one (**S7**)

To a solution of S6 (2.0 g, 5.85 mmol) in MeOH/H_2_O (40:10 mL) was added NH_4_Cl (5.37g, 99.4 mmol) and Fe (1.64 g, 29.2 mmol). After reflux for 2 h, the reaction mixture was filtered through celite and washed with MeOH. The collected filtrate was evaporated in vacuum to give a crude yellow solid which is pure enough to be used in the next step.

### General procedure for the synthesis of amides (step d)

To a stirred suspension of aniline *S2* or *S4* or *S7* (0.067 mmol) in dry DMF (1 mL) was added HATU (38 mg, 0.1 mmol), DIPEA (17.4 µL, 0.1 mmol) and the corresponding aryl carboxylic acid (0.1 mmol). After stirring for 12 h at room temperature, the reaction mixture was diluted with H_2_O and the aqueous phase was extracted with DCM. The combined organic phase was washed with brine, dried over Na_2_SO_4_, and concentrated under vacuum. The residue was purified by flash chromatography on silica gel (hexane/EtOAc = 30:70) to afford the corresponding amide products.

#### N-(10-(2-Methoxyethyl)-11-oxo-10,11-dihydrodibenzo[b,f][1,4]thiazepin-7-yl)furan-3-carboxamide (MM17)

Prepared from **S2** according to the general procedure, *MM17* was obtained as a yellow solid (72% yield). ^1^H NMR (400 MHz, Chloroform-d). δ 8.05 (s, 1H), 7.96 (s, 2H), 7.85 (d, *J* = 2.5 Hz, 1H), 7.63 (dd, *J* = 7.0, 2.2 Hz, 1H), 7.57 (dd, *J* = 8.8, 2.5 Hz, 1H), 7.44 (d, *J* = 8.5 Hz, 2H), 7.36 (dd, *J* = 7.4, 1.6 Hz, 1H), 7.24 (dd, *J* = 7.3, 2.0 Hz, 2H), 6.73 (d, *J* = 1.0 Hz, 1H), 4.54 (ddd, *J* = 13.7, 6.7, 5.5 Hz, 1H), 3.92 (dt, *J* = 13.8, 5.4 Hz, 1H), 3.85 – 3.75 (m, 1H), 3.68 – 3.58 (m, 1H), 3.33 (s, 3H). ^13^C NMR (101 MHz, Chloroform-d) δ 169.3, 160.8, 145.4, 144.1, 139.8, 138.9, 138.2, 137.0, 135.7, 131.1, 131.0, 130.7, 128.6, 126.7, 124.1, 122.7, 121.3, 108.4, 70.1, 58.9, 51.6. HRMS (ESI) calcd for C_21_H_19_N_2_O_4_S [M+H]^+^ 395.1060, found 395.1072. Purity = 96.8%.

#### 4-Azido-N-(11-oxo-10-(2-(prop-2-yn-1-yloxy)ethyl)-10,11-dihydro-dibenzo[b,f][1,4]thiazepin-7-yl)benzamide (MM514)

Prepared from **S4** according to the general procedure, probe *MM514* was obtained as a yellow solid (82% yield). ^1^H NMR (400 MHz, Chloroform-d) δ 7.92 (d, *J* = 2.5 Hz, 1H), 7.84 (d, *J* = 8.6 Hz, 2H), 7.77 (s, 1H), 7.67 (dd, *J* = 7.6, 1.7 Hz, 1H), 7.60 (dd, *J* = 8.8, 2.5 Hz, 1H), 7.50 (d, *J* = 8.7 Hz, 1H), 7.42 (dd, *J* = 7.5, 1.5 Hz, 1H), 7.35 – 7.25 (m, 2H), 7.12 (d, *J* = 8.7 Hz, 2H), 4.57 (qd, *J* = 8.9, 8.1, 5.1 Hz, 1H), 4.16 (d, *J* = 2.4 Hz, 2H), 3.97 (dq, *J* = 9.0, 5.0 Hz, 2H), 3.81 (dt, *J* = 9.5, 4.2 Hz, 1H), 2.41 (t, *J* = 2.3 Hz, 1H). ^13^C NMR (101 MHz, Chloroform-d) δ 169.2, 164.8, 144.0, 139.9, 138.9, 138.2, 137.1, 135.9, 131.1, 131.0, 130.8, 130.7, 129.0, 128.7, 126.7, 124.2, 121.4, 119.2, 79.5, 74.7, 67.6, 58.4, 51.6. HRMS (ESI) calcd for C_25_H_20_N_5_O_3_S [M+H]^+^ 470.1281, found 470.1290. Purity = 94.1%.

#### 4-Methoxy-N-(10-(2-methoxyethyl)-11-oxo-10,11-dihydrodibenzo[b,f][1,4]thiazepin-7-yl)benzamide (MM524)

Prepared from **S2** according to the general procedure, analog *MM524* was obtained as a yellow solid (74% yield). ^1^H NMR (400 MHz, Chloroform-d) δ 7.92 (d, *J* = 2.5 Hz, 1H), 7.87 (s, 1H), 7.81 (d, *J* = 8.8 Hz, 2H), 7.67 (dd, *J* = 7.3, 2.0 Hz, 1H), 7.60 (dd, *J* = 8.8, 2.5 Hz, 1H), 7.47 (d, *J* = 8.8 Hz, 1H), 7.41 (dd, *J* = 7.3, 1.7 Hz, 1H), 7.34 – 7.20 (m, 2H), 6.96 (d, *J* = 8.8 Hz, 2H), 4.61 – 4.50 (m, 1H), 3.98 – 3.88 (m, 1H), 3.87 (s, 3H), 3.85 – 3.74 (m, 1H), 3.65 (dd, *J* = 10.0, 5.6 Hz, 1H), 3.34 (s, 3H). ^13^C NMR (101 MHz, Chloroform-d) δ 169.1, 165.2, 162.7, 139.8, 138.9, 138.4, 137.1, 136.0, 131.1, 131.0, 130.7, 129.0, 128.6, 126.7, 126.5, 124.0, 121.2, 114.1, 70.2, 58.9, 55.5, 51.5. HRMS (ESI) calcd for C_24_H_23_N_2_O_4_S [M+H]^+^ 435.1373, found 435.1387. Purity = 96.1%.

#### 2-Methoxy-N-(10-(2-methoxyethyl)-11-oxo-10,11-dihydrodibenzo[b,f][1,4]thiazepin-7-yl)benzamide (MM691)

Prepared from **S2** according to the general procedure, analog *MM691* was obtained as a yellow solid (50% yield). ^1^H NMR (400 MHz, Chloroform-d) δ 8.05 (s, 1H), 7.93 (d, *J* = 2.5 Hz, 1H), 7.68 – 7.57 (m, 2H), 7.47 (d, *J* = 8.8 Hz, 1H), 7.42 – 7.37 (m, 2H), 7.37 – 7.33 (m, 2H), 7.31 – 7.22 (m, 2H), 7.12 – 7.03 (m, 1H), 4.54 (ddd, *J* = 13.8, 6.6, 5.5 Hz, 1H), 3.92 (dt, *J* = 13.8, 5.5 Hz, 1H), 3.85 (s, 3H), 3.83 – 3.77 (m, 1H), 3.68 – 3.59 (m, 1H), 3.34 (s, 3H). ^13^C NMR (101 MHz, Chloroform-d) δ 169.1, 165.6, 160.0, 140.0, 138.9, 138.3, 137.1, 135.9, 135.8, 131.1, 131.0, 130.7, 129.8, 128.6, 126.7, 124.1, 121.2, 118.7, 118.2, 112.6, 70.2, 58.9, 55.5, 51.6. HRMS (ESI) calcd for C_24_H_23_N_2_O_4_S [M+H]^+^ 435.1373, found 435.1388. Purity = 96.5%.

#### N-(10-(but-2-yn-1-yl)-3-fluoro-11-oxo-10,11-dihydrodibenzo[b,f][1,4]thiazepin-7-yl)-3-methyl-2-oxo-2,3-dihydrobenzo[d]oxazole-5-carboxamide (MM927)

Prepared from **S7** according to the general procedure, analog *MM927* was obtained as a white solid (56% yield). ^1^H NMR (400 MHz, Chloroform-d): δ 7.93 (d, *J* = 2.8 Hz, 1H), 7.80-7.88 (m, 1H), 7.68-7.74 (m, 2H), 7.66-7.67 (m, 1H), 7.56-7.58 (m, 2H), 7.26-7.29 (m, 1H), 7.14-7.16 (m, 1H), 7.00-7.05 (m, 1H), 4.90-4.95 (m, 1H), 4.42-4.47 (m, 1H), 3.47 (s, 3H), 1.86 (t, *J* = 4.4 Hz, 3H). ^13^C NMR (151 MHz, Chloroform-d) δ 167.6, 164.8, 164.0, 162.3, 154.4, 145.2, 140.6 (d, *J* = 8.2 Hz), 139.9, 136.1, 134.9, 134.0 (d, *J* = 9.1 Hz), 133.7 (d, *J* = 3.3 Hz), 132.5, 130.7, 125.3, 124.2, 121.5, 121.3, 118.0, 117.9, 116.1, 116.0, 109.7, 108.1, 80.4, 74.6, 41.3, 28.4, 3.8. HRMS (ESI) calcd for C_26_H_19_FN_3_O_4_S [M + H]^+^ 488.1075, found 488.1106. Purity = 99.9%.

---

**Figure S1.**
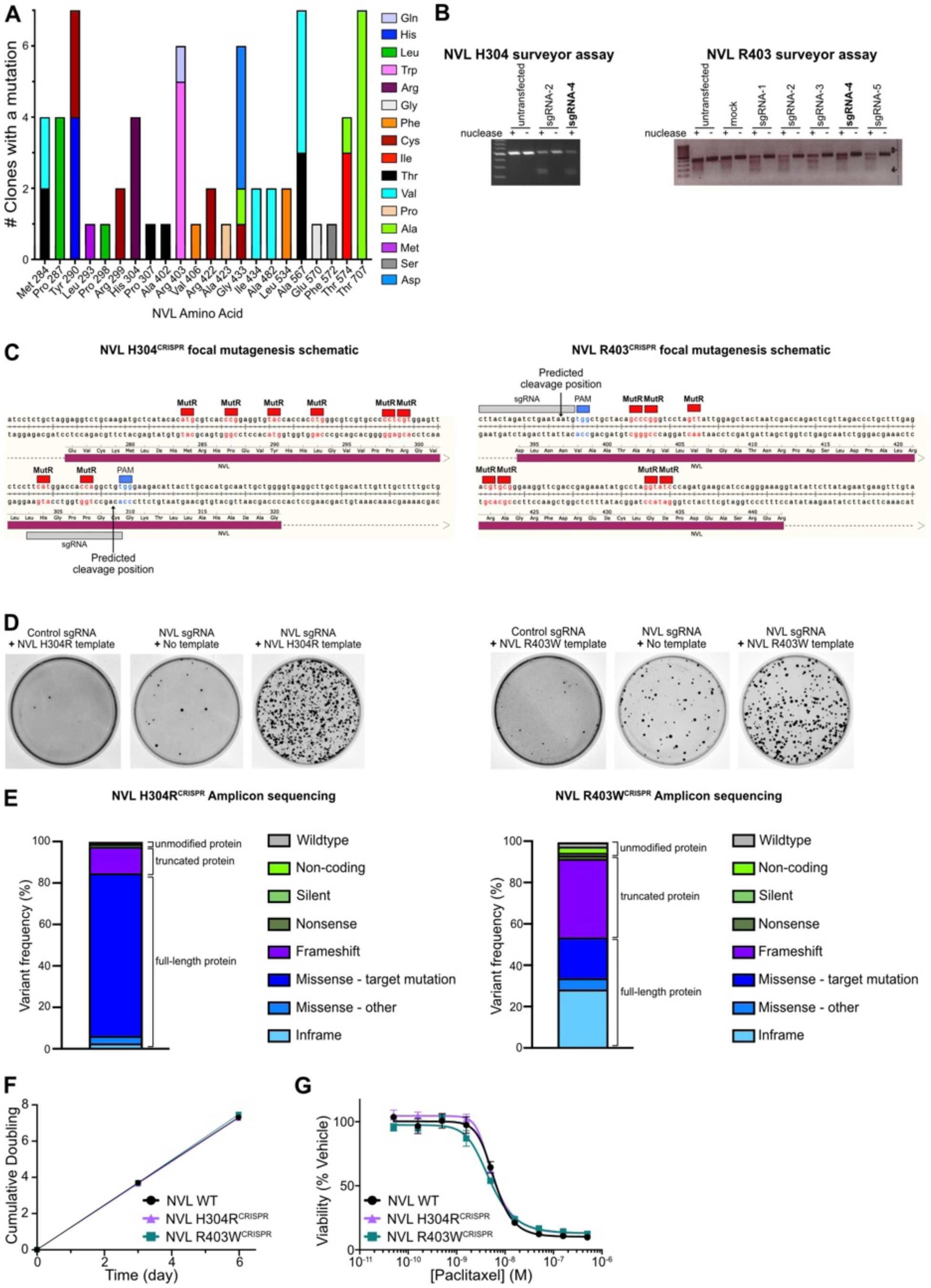
Analysis of MM17 resistant clones and characterization of NVL CRISPR focal mutagenesis. **A.** Complete set of NVL mutations in resistant clones identified by PCR based amplicon sequencing of four exons (9, 12, 14, 18) with mutations found in (Fig. 1F, Table S2). **B.** T7 endonuclease surveyor assay to determine the appropriate guide that cut the target region. The selected guide is bolded. **C.** Schematics of CRISPR focal mutagenesis at MM17 resistant hotspot regions of NVL. **D.** Cresyl violet staining of transfected HCT116 cells after selection with 1 µM MM17 for 2 weeks (NVL H304R^CRISPR^) or 7.5 µM MM17 for 10 days (R403W^CRISPR^). **E.** Amplicon sequencing results of NVL H304R^CRISPR^ and R403W^CRISPR^. **F.** Growth curves of HCT116 NVL WT, NVL H304R^CRISPR^, and R403W^CRISPR^ cell lines. (n = 3 biological replicates, mean ± sem). **G.** Viability assay of cell lines in (F) treated with paclitaxel for 72 hours. (n = 3 biological replicates, mean ± sem).

**Figure S2.**
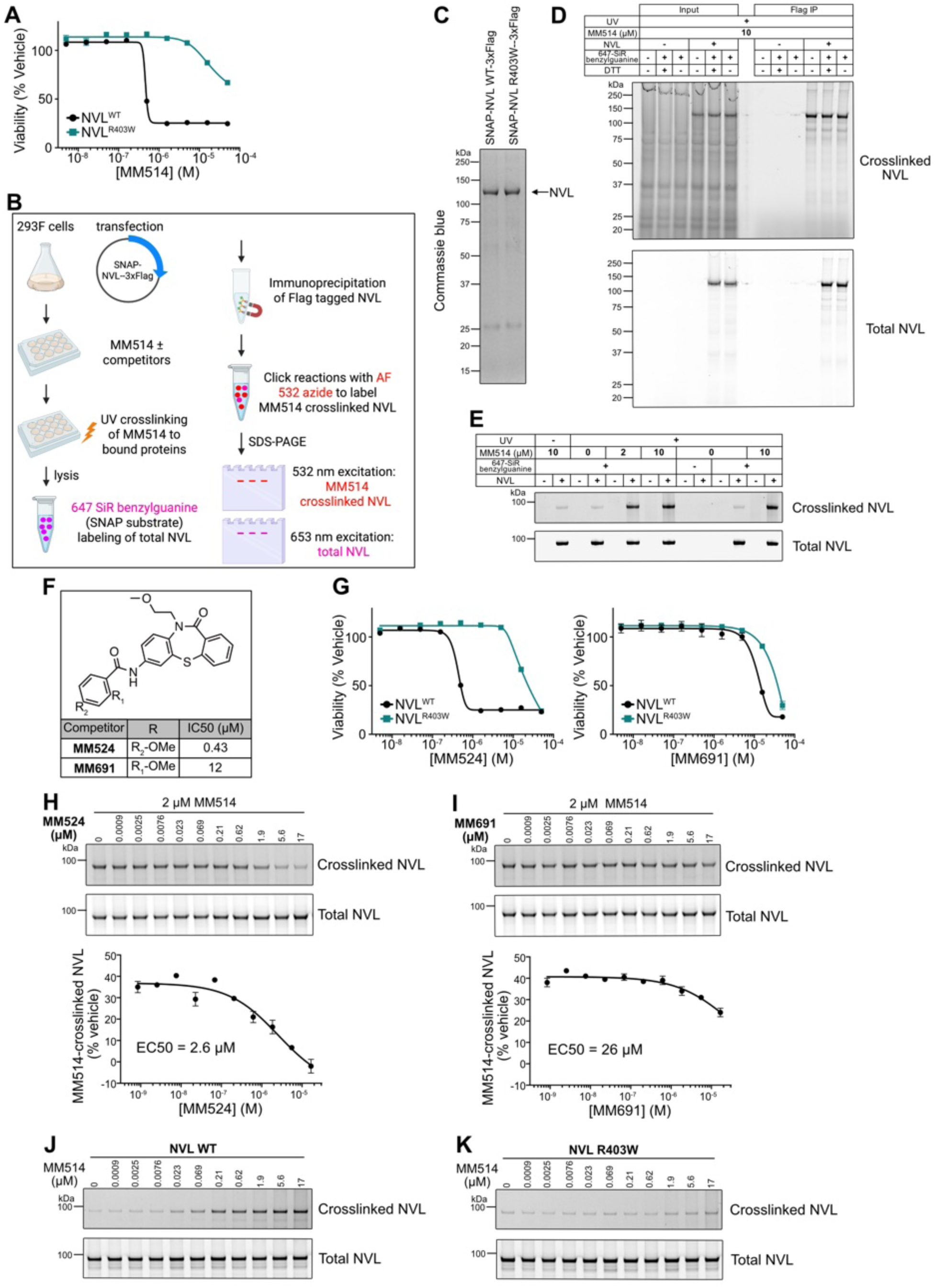
Chemical probe crosslinking assay. A. Viability assay of HCT116 NVL WT and NVL R403W^CRISPR^ cells treated with MM514 for 72 hours. (n = 3 biological replicates, mean ± sem). B. Schematics of MM514 UV-crosslinking of recombinant NVL. C. Coomassie blue stain of recombinant NVL WT and R403W following Flag immunoprecipitation from 293F cells. D. Crosslinked and total NVL in input or Flag-immunoprecipitated from 293F cells treated with 10 µM MM514 and UV light. E. Crosslinked and total NVL WT-SNAP purified from 293F cells treated MM514, with or without UV light. Note the spectral overlap between Alexa fluor 532 azide (crosslinked NVL) and Alexa fluor 647 benzylguanine (total NVL) (lane 12 and 14), resulting in background fluorescence when excited at 532 nm despite the lack of MM514. F. Chemical structures of competitor compounds MM524 and MM691 and associated HCT116 anti-proliferation IC50 values calculated from G. Viability assay of HCT116 NVL WT and NVL R403W^CRISPR^ cells treated with MM524 and MM691 for 72 hours. (n = 3 biological replicates, mean ± sem H. Fluorescence scans and quantitation of recombinant NVL WT 293F cells treated with MM514, and MM524 (H) or MM691 (I). J, K. Recombinant NVL purified from 293F cells treated with MM514 and UV light, related to Fig. 1I, J.

**Figure S3.**
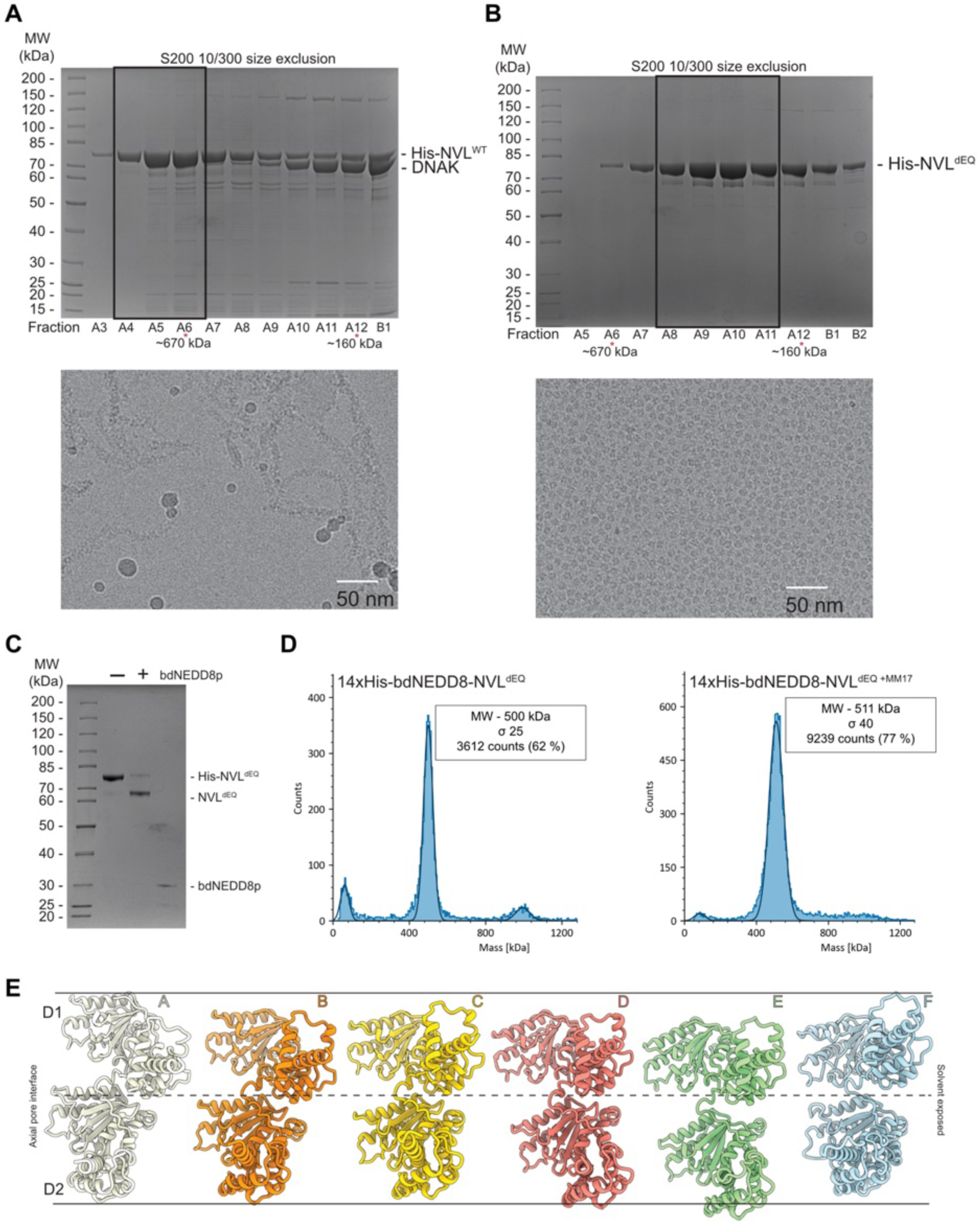
Purification and biophysical characterization of NVL. **A.** Coomassie Blue-stained SDS-PAGE of peak fractions from a size exclusion chromatography (SEC) run of NVL WT on a Superdex 200 10/300 column (top) and corresponding cryo-EM micrograph of the boxed fractions showing filamentous assemblies (bottom). Approximate molecular weights (red asterisks) are estimated from protein standards run on the same column. DnaK was identified as the major contaminant by mass spectrometry. **B.** Coomassie Blue-stained SDS-PAGE of peak fractions from NVL^dEQ^ SEC (top) and corresponding cryo-EM micrograph of boxed fractions showing hexamer formation (bottom). Approximate molecular weights are indicated as in A. **C.** Coomassie Blue-stained SDS-PAGE comparing cleaved and uncleaved NVL^dEQ^. Uncleaved protein was used for MM17-bound reconstruction; cleaved protein was used to make the apo-NVL^dEQ^ grids to reduce preferred orientation. **D.** Mass photometry analysis of apo (left) or MM17-bound (right) NVL^dEQ^. Histograms show particle counts fitted to gaussian curves. Peak positions and estimated molecular weights are indicated. **E.** Schematic of NVL subunits A-E in staircase arrangement. Individual subunits are depicted relative to a fixed position on the y-axis (dotted line), highlighting the staircase-like offset between subunits A-E and the dynamic position of subunit F.

**Figure S4.**
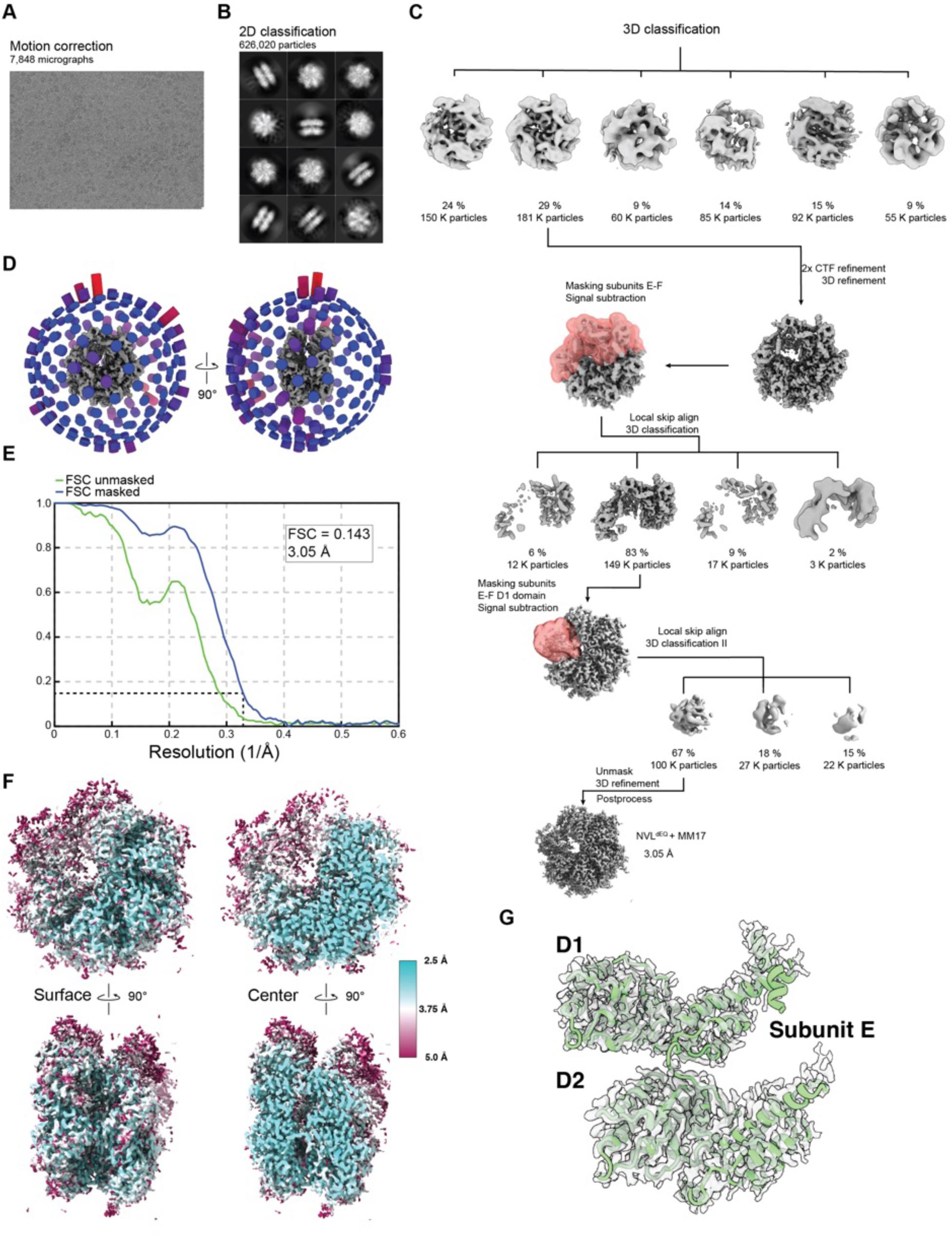
Cryo-EM data processing and reconstruction of NVL^dEQ^-MM17. A. Representative motion-corrected micrograph of NVL^dEQ^-MM17. B. Representative 2D class average showing top and side views of the NVL hexamer. C. Overview of the 3D classification and refinement workflow. Initial classification was based on global map quality. Two rounds of 3D classification without alignment using focused masks around the D/E subunits and F subunits (red envelopes) to improve local resolution. D. Angular distribution plot showing particle orientation coverage. E. Gold-standard Fourier shell correlation (FSC) curves used to estimate the overall resolution, with a cutoff of FSC = 0.143. F. Final maps colored by local resolution as determined by ResMap. Top down (top) and side (bottom) views are shown for both surface renderings (left) and center slices (right). G. Model of subunit E (D1 and D2 domains) docked into the corresponding cryo-EM density.

**Figure S5.**
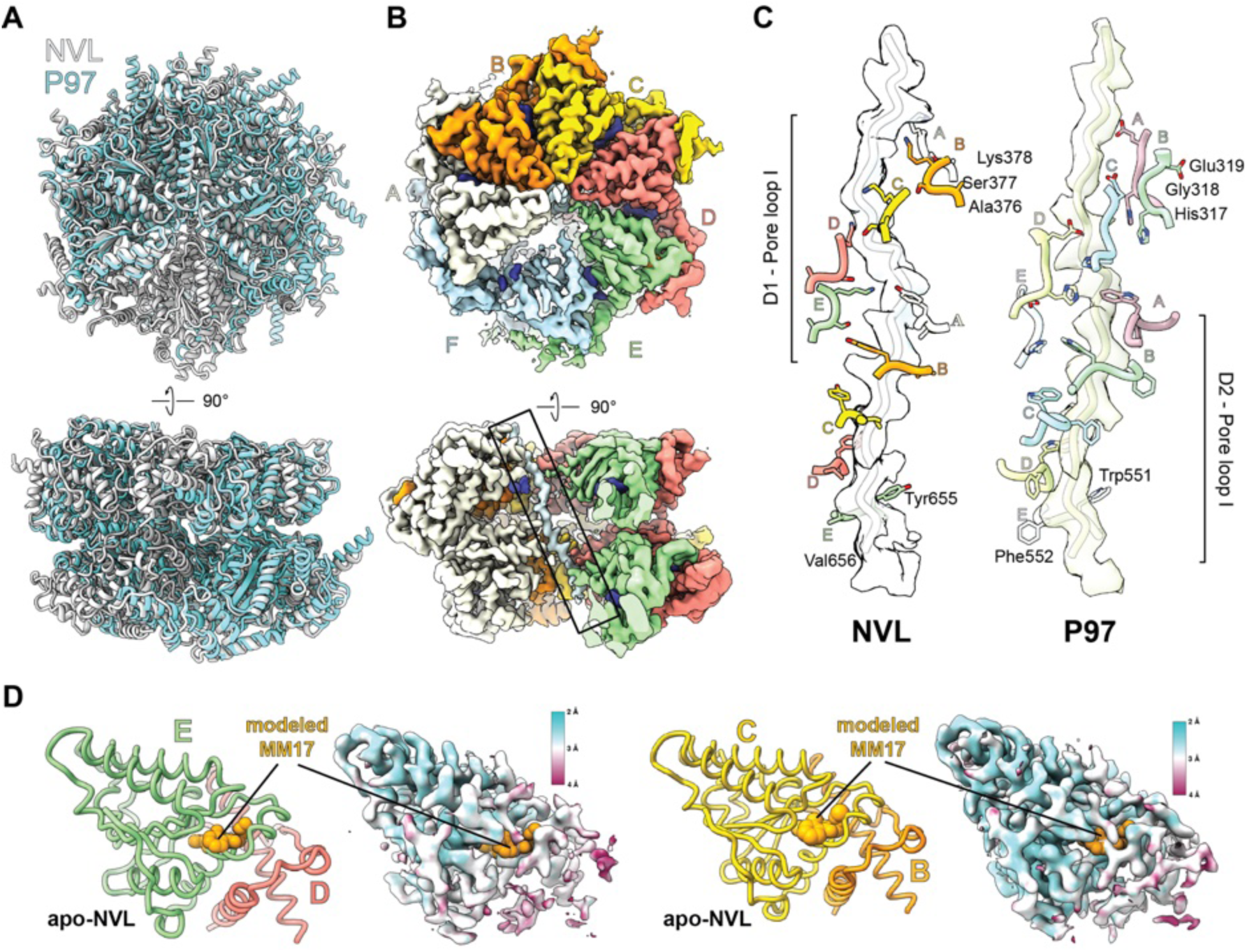
Structural comparison between NVL and P97 hexamers and insights into MM17 engagement. **A.** Structural alignment between NVL^dEQ^ (gray) and p97 (light blue; PDB 7MHS (*43*)) hexamers reveals conserved architecture. Subunit F of P97 is not modeled due to weak local density. **B.** Substrate mimic in the NVL^dEQ^-MM17 structure. Top-down (top) and side (bottom) views of the cryo-EM map shows a peptide-like density (boxed) spanning the axial pore. Subunits are colored as in Fig. 2; subunit F is omitted in the side view for clarity. **C.** Pore-loop engagement of the axial peptide. Comparison of D1/D2 pore-loop interactions with the axial peptide in NVL^dEQ^-MM17 (left) and p97 (right; PDB 7MHS; EMDB 23835 (*43*)). Cryo-EM density is shown as a transparent surface; pore loops and substrate peptides are shown as cartoons with side chains as sticks. **D**. Accessibility of the MM17 binding pocket in the apo-structure. Cartoon models and cryo-EM maps of the D1 domain interface between subunits D/E (left) and B/C (right), with maps colored by local resolution. MM17 from the NVL^dEQ^-MM17 structure is overlayed as orange spheres, highlighting better binding pocket accessibility in subunit E compared to subunit C.

**Figure S6.**
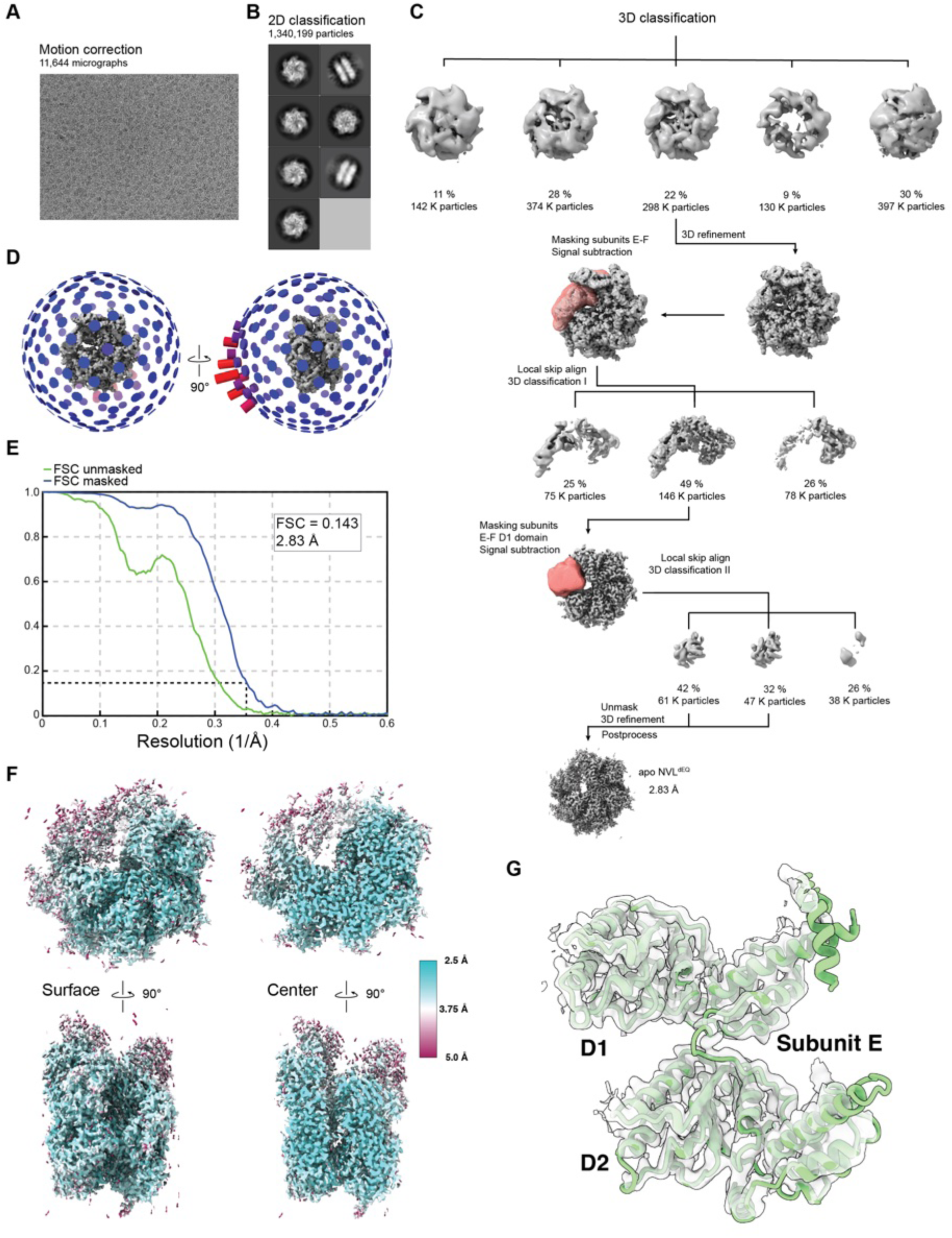
Cryo-EM data processing of apo-NVL^dEQ^. A. Representative motion-corrected micrograph of apo-NVL^dEQ^. B. Representative 2D class average showing top and side views of the NVL hexamer. C. Overview of the 3D classification and refinement workflow. Initial classification was based on global map quality. Two rounds of 3D classification without alignment using focused masks around the D/E subunits and F subunits (red envelopes) to improve local resolution. D. Angular distribution plot showing particle orientation coverage. E. Gold-standard Fourier shell correlation (FSC) curves used to estimate the overall resolution, with a cutoff of FSC = 0.143. F. Final maps colored by local resolution as determined by ResMap. Top down (top) and side (bottom) views are shown for both surface renderings (left) and center slices (right). G. Model of subunit E (D1 and D2 domains) docked into the corresponding cryo-EM density.

**Figure S7.**
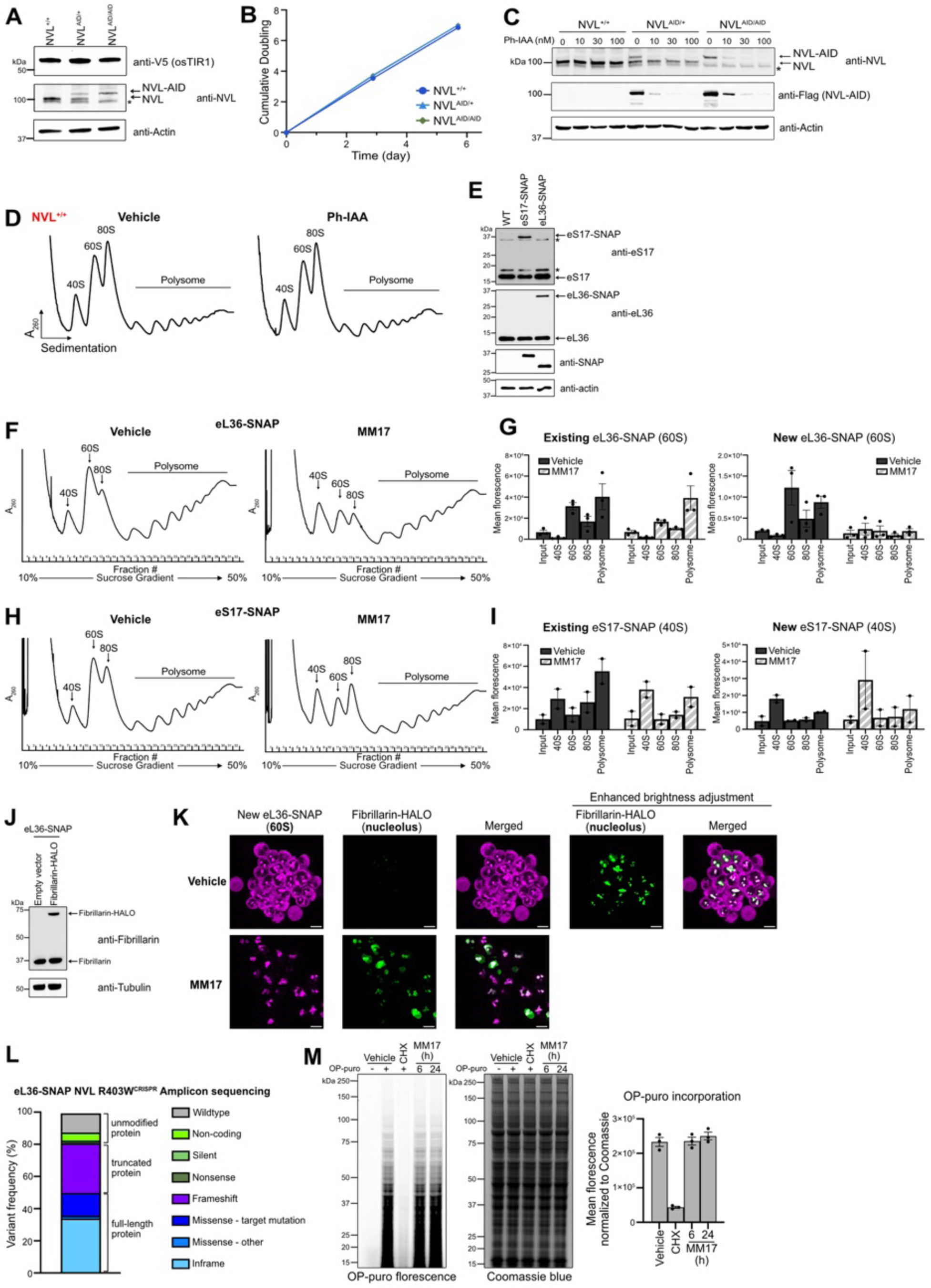
Characterization of cell lines with AID tagged NVL and SNAP tagged ribosomal proteins. **A.** Immunoblots of V5 (osTIR1), NVL and actin (loading control) in HCT116 NVL^+/+^, NVL^AID/+^, NVL^AID/AID^ cells. * non-specific bands. **B.** Growth curves of cell lines in (A). (n = 3 biological replicates, mean ± sem). **C.** Immunoblots of NVL, Flag (NVL-AID) and actin (loading control) of HCT116 NVL^+/+^, NVL^AID/+^, NVL^AID/AID^ cells treated for 6 hours with Ph-IAA. * non-specific bands. **D.** Polysome profiling of HCT116 NVL^+/+^ treated with 100 nM Ph-IAA for 24 hours. **E**. Immunoblots of eS17, eL36, SNAP and actin (loading control) in HCT116 WT, eS17-SNAP, and eL36-SNAP cells. * non-specific bands. **F.** Polysome profiling to separate ribosomal species in HCT116 eL36-SNAP cells treated with 3 µM MM17 for 6 hours. The ribosome-associated fractions (7 to 29) are subsequently analyzed in Fig. 3E. **G**. Quantitation of existing and newly synthesized eL36-SNAP in sucrose gradient fractions corresponding to different ribosomal species, related to Fig. 3E. (n=3 biological replicates, mean ± sem). **H.** Polysome profiling to separate ribosomal species in HCT116 eL36-SNAP cells treated with 3 µM MM17 for 6 hours. The ribosome-associated fractions (7 to 29) are subsequently analyzed in Fig. 3F. **I**. Quantitation of existing and newly synthesized eS17-SNAP in sucrose gradient fractions corresponding to different ribosomal species, related to Fig. 3F. (n=2 biological replicates, mean ± sem). **J.** Immunoblots of fibrillarin and tubulin (loading control) in HCT116 eL36-SNAP cells expressing empty vector or fibrillarin-HALO. **K.** Microscopy images of newly synthesized eL36-SNAP (magenta) and nucleolar marker fibrillarin-HALO (green) treated for 24 hours with 3 µM MM17. Enhanced brightness adjustment of fibrillarin-HALO is shown for vehicle treatment. (Scale bar = 10 µm). **L.** Amplicon sequencing result of HCT116 eL36-SNAP NVL R403W^CRISPR^ cell line. **M.** O-propargyl-puromycin (OP-puro) incorporation assay to assess protein translation in HCT116 cells treated with 10 µg/mL cycloheximide for 30 minutes, or 3 µM MM17. Fluorescence scans and quantitation showing Alexa fluor 532 azide conjugated OP-puro and Coomassie blue staining (loading control). (n = 3 biological replicates, mean ± sem).

**Figure S8.**
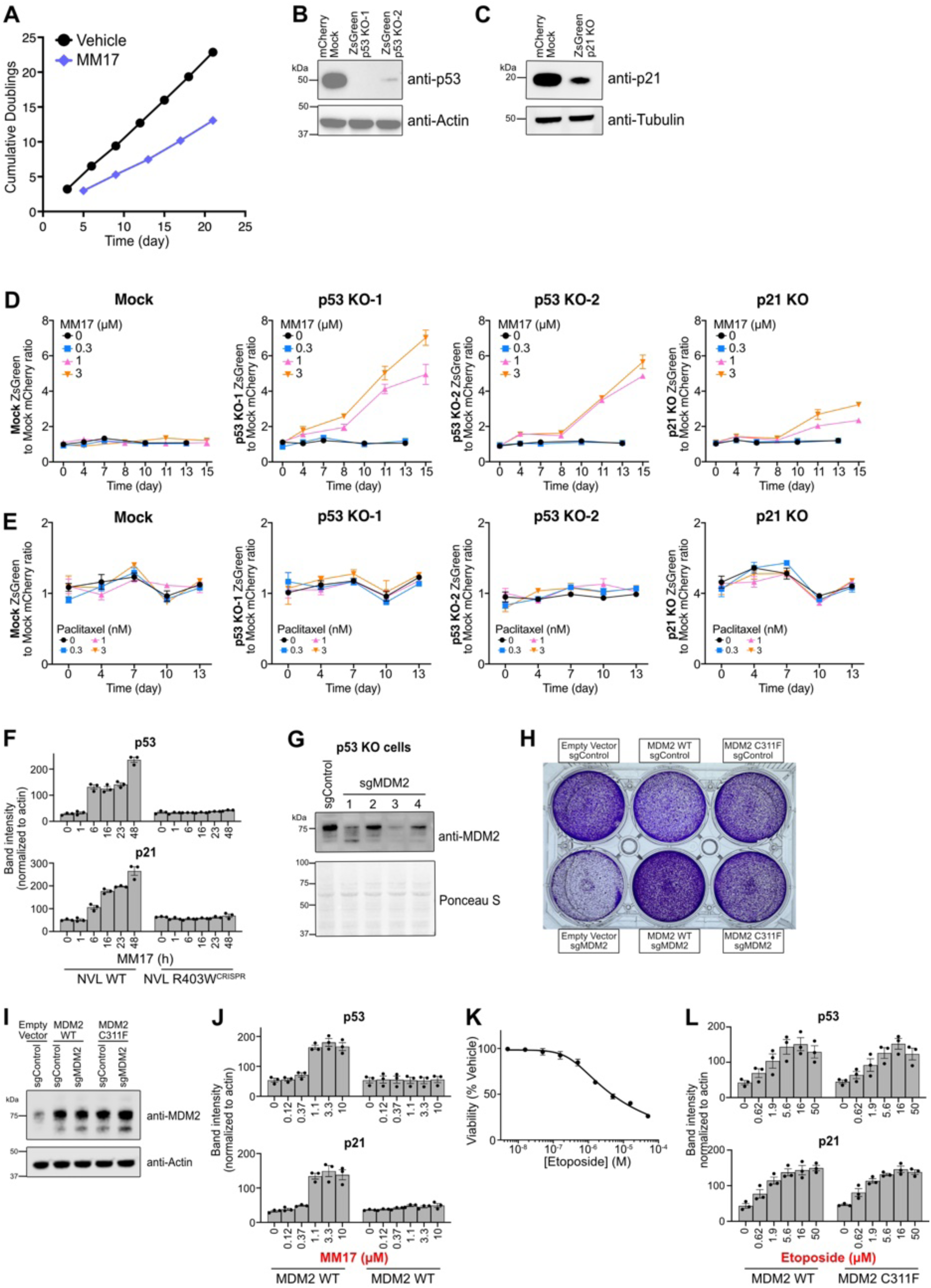
Characterization of MDM2 mediated p53 stabilization. **A.** Growth curves of Cas9-expressing HCT116 in the genome wide CRISPR/Cas9 knockout screen shown in Fig. 4A. At each plotted point, cells were treated for 24 hours with vehicle or 1 µM MM17 and allowed to recover without treatment until 80% confluency (n=1). **B.** Immunoblots of p53 and actin (loading control) in HCT116 mCherry mock, ZsGreen p53 KO guide 1 (p53 KO-1), and ZsGreen p53 KO guide 2 (p53 KO-2) cells. **C.** Immunoblots of p21 and tubulin (loading control) in HCT116 mCherry mock, and ZsGreen p21 KO cells. **D, E.** Growth competition assay between mCherry mock and ZsGreen mock, ZsGreen p53 KO-1, ZsGreen p53 KO-2, or ZsGreen p21 KO cells. At each plotted point, cells were treated for 24 hours with MM17 (**D**) or paclitaxel (**E**) and allowed to recover without compound until confluency. (n=3 biological replicates, mean ± sem). **F.** Quantitation of p53 and p21 immunoblots, related to Fig. 4B. (n = 3 biological replicates, mean ± sem). **G.** Immunoblot of MDM2 and Ponceau S staining (loading control) in HCT116 p53 KO cells transduced with non-target control or four sgRNAs targeting MDM2. sgMDM2 guide 3 was selected for subsequent experiments. **H.** Cresyl violet staining after 1 week of antibiotics selection of HCT116 cells expressing empty vector, ectopic MDM2 WT, or MDM2 C311F transduced with mock or MDM2 guide. **I.** Immunoblots of MDM2 and actin (loading control) of cell lines in (H). **J.** Quantitation of p53 and p21 immunoblots, related to Fig. 4D. (n = 3 biological replicates, mean ± sem). **K.** Viability assay of HCT116 cells treated with etoposide for 72 hours. (n = 3 biological replicates, mean ± sem). **L.** Quantitation of p53 and p21 immunoblots, related to Fig. 4E. (n = 3 biological replicates, mean ± sem).

**Figure S9.**
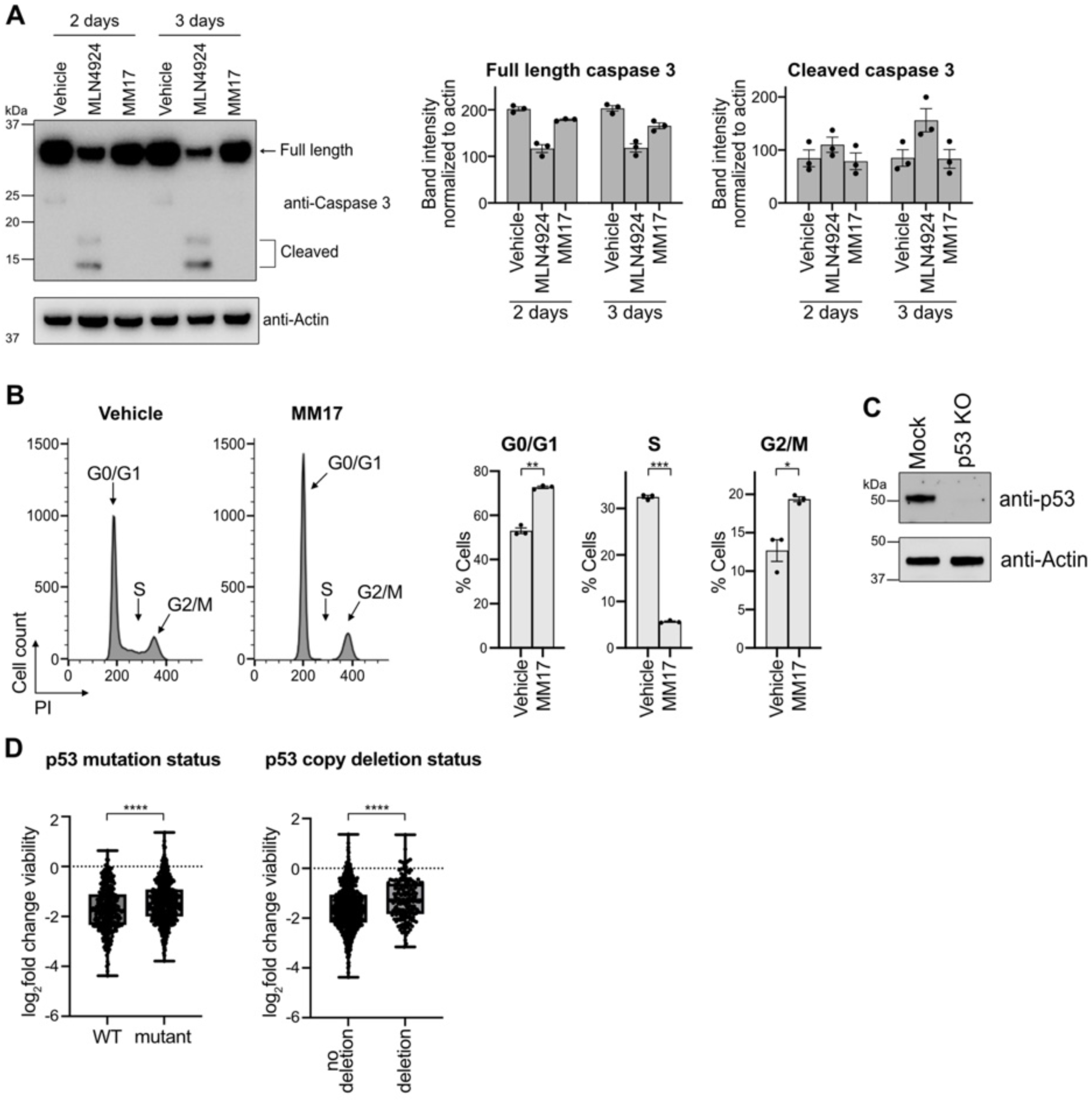
MM17 leads to cell cycle arrest. **A.** Immunoblots and quantitation of caspase-3 and actin (loading control) in HCT116 cells treated with 1 µM MLN4924 or 3 µM MM17. (n = 3 biological replicates, mean ± sem). **B.** Flow cytometry-based cell cycle analysis in HCT116 cells treated with vehicle or 3 µM MM17 for 3 days. (n = 3 biological replicates, mean ± sem; two-tailed unpaired t-test, ***p<0.001, **p<0.01, *p<0.05). **C.** Immunoblots of p53 and actin (loading control) in HCT116 mock engineered or p53 knockout (KO) cells. **D.** Viability (log_2_fold change relative to vehicle) of 855 cancer cell lines treated with 10 µM MM17, grouped by their p53 mutation or copy number deletion status. Each dot represents a cell line. (two-tailed paired t-test, ****p<0.0001).

**Figure S10.**
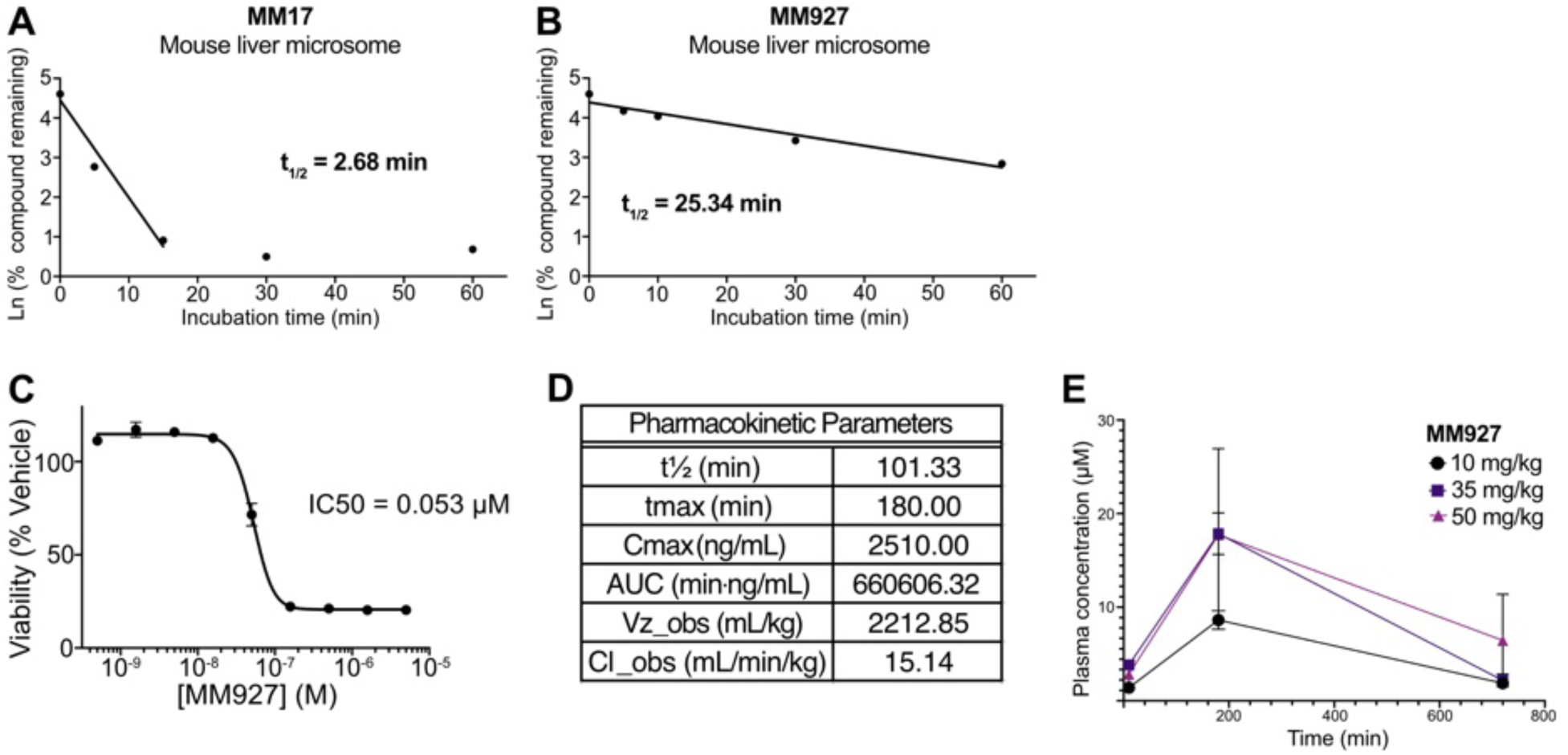
Pharmacokinetic properties of MM927. A, B. Compound stability in mouse liver microsomes incubated in vitro with MM17 (A) or MM927 (B) over time. (MM17: n = 1; MM927: n = 2 biological replicates, mean ± sem). C. Viability assay of HCT116 cells treated with MM927 for 72 hours. (n = 3 biological replicates, mean ± sem). D. Pharmacokinetic parameters of MM927, related to Fig. 5B. E. MM927 concentrations in mouse plasma following an IP injection (n = 3 biological replicates, mean ± sem).

**Figure S11.**
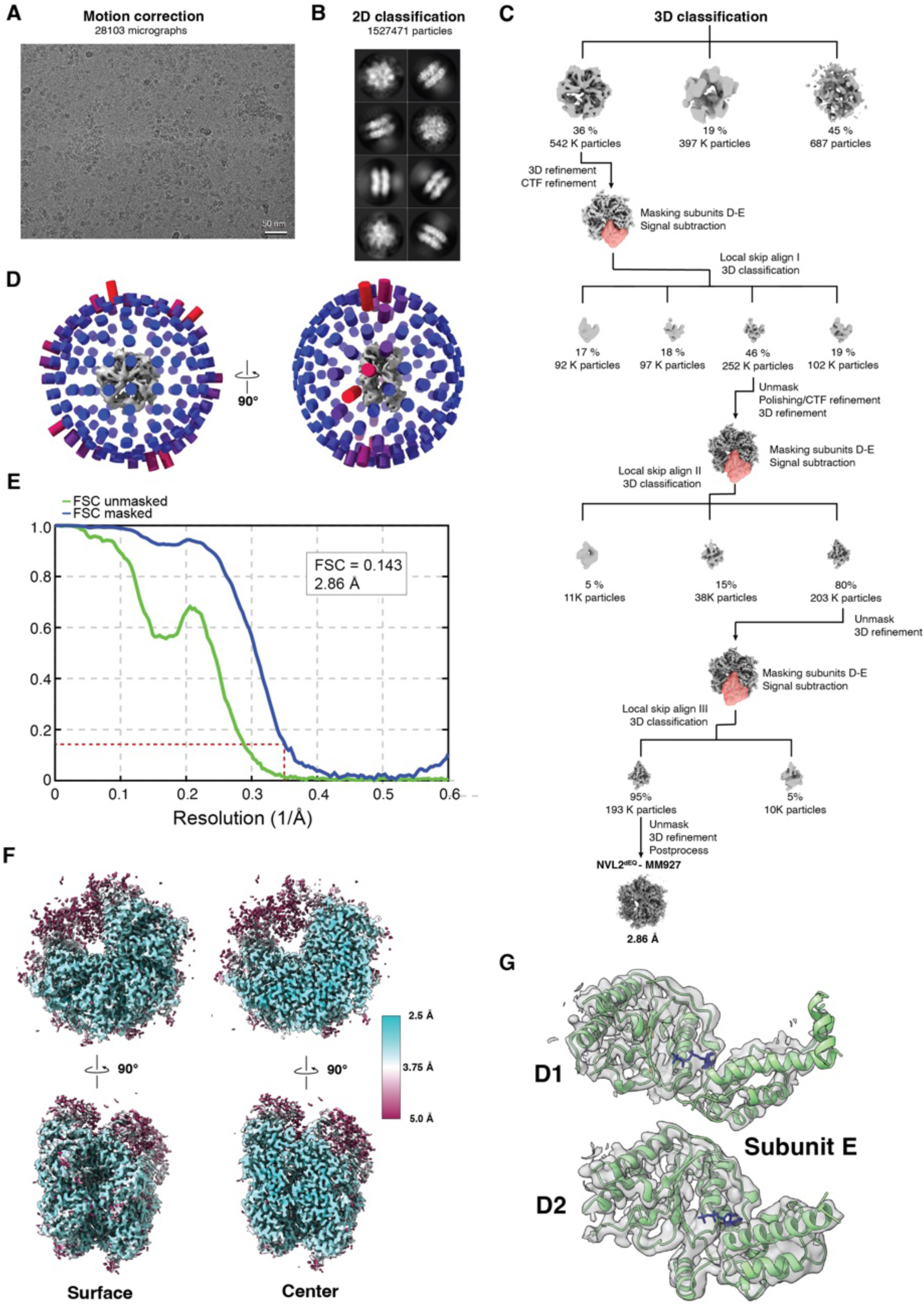
Cryo-EM data processing of NVL^dEQ^-MM927. A. Representative motion-corrected micrograph of NVL^dEQ^-MM927. B. Representative 2D class average showing top and side views of the NVL hexamer. C. Overview of the 3D classification and refinement workflow. Initial classification was based on global map quality. Three rounds of 3D classification without alignment using focused masks around the D/E subunits and F subunits (red envelopes) to improve local resolution. D. Angular distribution plot showing particle orientation coverage. E. Gold-standard Fourier shell correlation (FSC) curves used to estimate the overall resolution, with a cutoff of FSC = 0.143. F. Final maps colored by local resolution as determined by ResMap. Top down (top) and side (bottom) views are shown for both surface renderings (left) and center slices (right). G. Model of subunit E (D1 and D2 domains) docked into the corresponding cryo-EM density.

**Figure S12.**
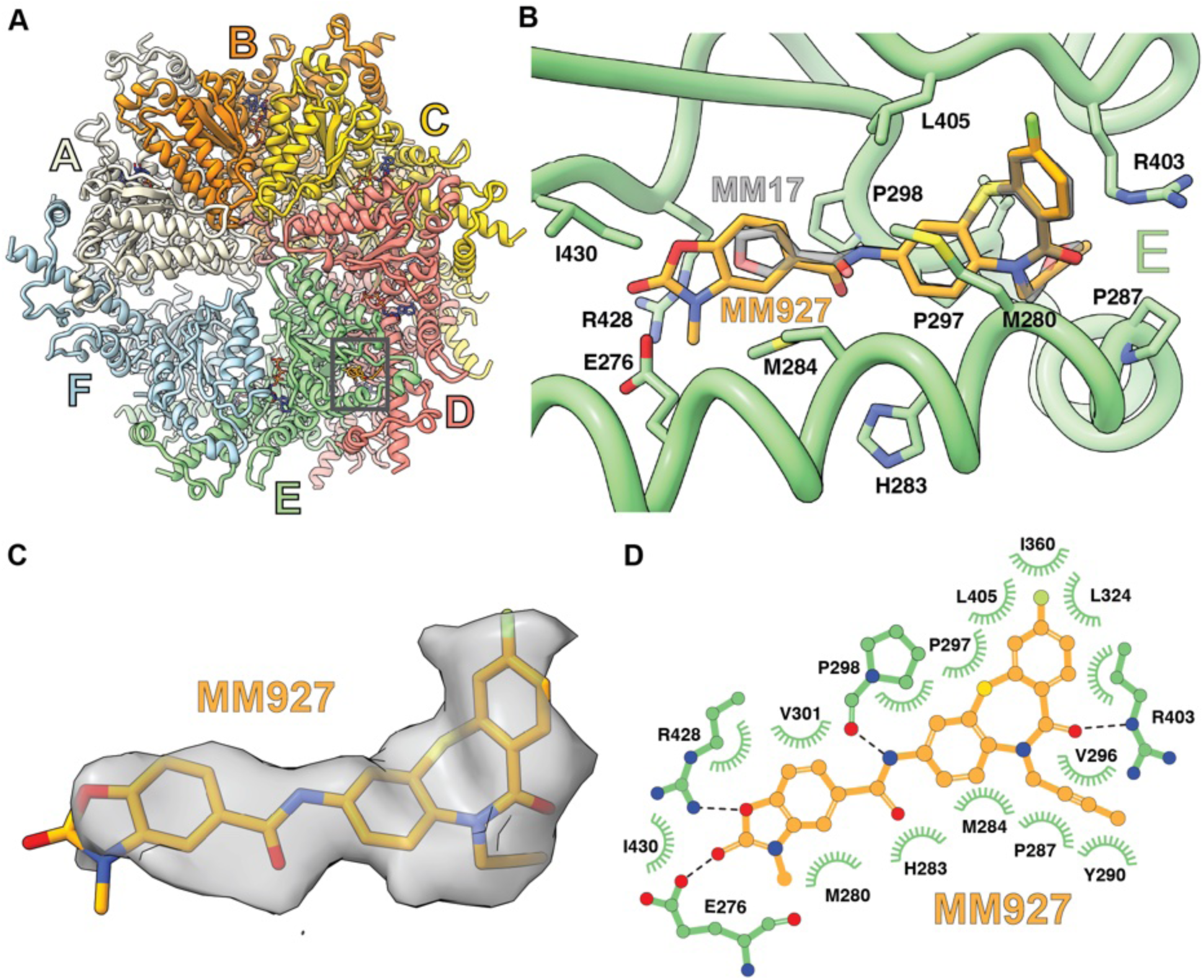
Cryo-EM structure of NVL^dEQ^ in complex with MM927. **A.** Overview of the NVL^dEQ^ hexamer in complex with MM927, shown as cartoons with subunits colored as in Fig. 2. **B.** Binding site comparison showing MM927 (orange) and MM17 (transparent gray) overlaid in the same ligand pocket of subunit E. **C.** Cryo-EM density for MM927 in subunit E, shown as a transparent surface **D.** Atomic diagram of MM927 (orange) and its interaction network with NVL residues (green). Hydrophobic interactions are displayed as fans and hydrogen bonds shown as dashed lines.

**Figure S13.**
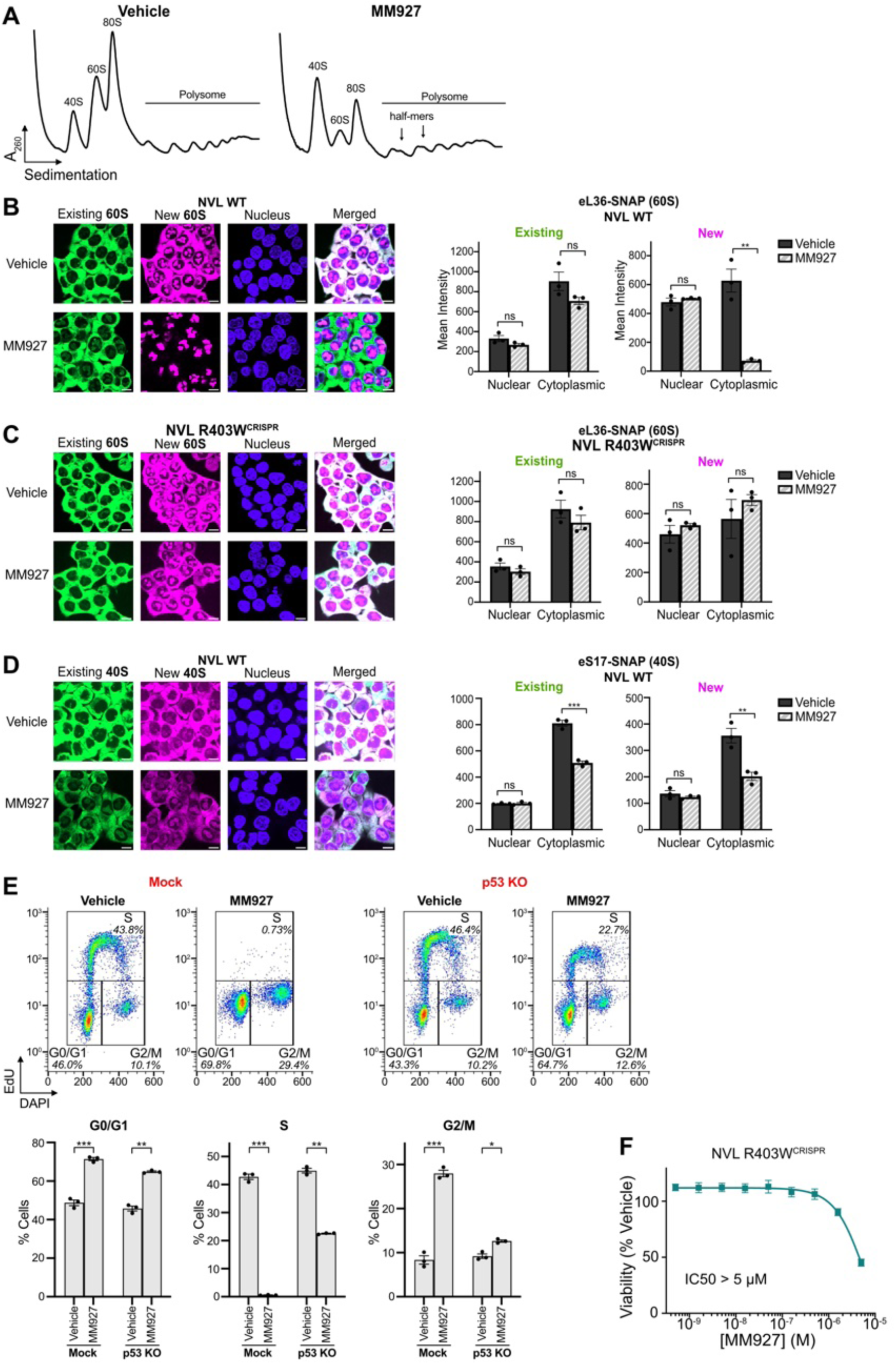
MM927 mechanism of action studies. **A.** Polysome profiling of HCT116 NVL WT (**A**) and NVL R403W^CRISPR^ (**B**) cells treated with 0.5 µM MM927 for 24 hours. **B, C.** Representative microscopy images and quantitation of existing (green) and newly synthesized (magenta) eL36-SNAP in NVL WT (**B**) and NVL R403W^CRISPR^ (**C**) cells treated with 0.5 µM MM927 for 24 hours. (Scale bar = 10 µm; n = 3 biological replicates, mean ± sem; two-tailed unpaired t-test, **p<0.01, ns = non-significant). Data from the vehicle group is re-used from Fig. 3G, H, as these are done in the same experiment. **D.** Representative microscopy images and quantitation of existing (green) and newly synthesized (magenta) eS17-SNAP in HCT116 cells treated with 0.5 µM MM927 for 24 hours. (Scale bar = 10 µm; n = 3 biological replicates, mean ± sem; two-tailed unpaired t-test, **p<0.01, ns = non-significant). Data from the vehicle group is re-used from Fig. 3I, as these are done in the same experiment. **E.** Flow cytometry-based analysis of cell cycle profile in HCT116 mock and p53 KO cells treated with 0.5 µM MM927. (n = 3 biological replicates, mean ± sem; two-tailed unpaired t-test, ***p<0.001, **p<0.01, *p<0.05, ns = non-significant). **F.** Viability assay of HCT116 NVL R403W^CRISPR^ cells treated with MM927 for 72 hours. (N = 3 biological replicates, mean ± sem).

**Figure S14.**
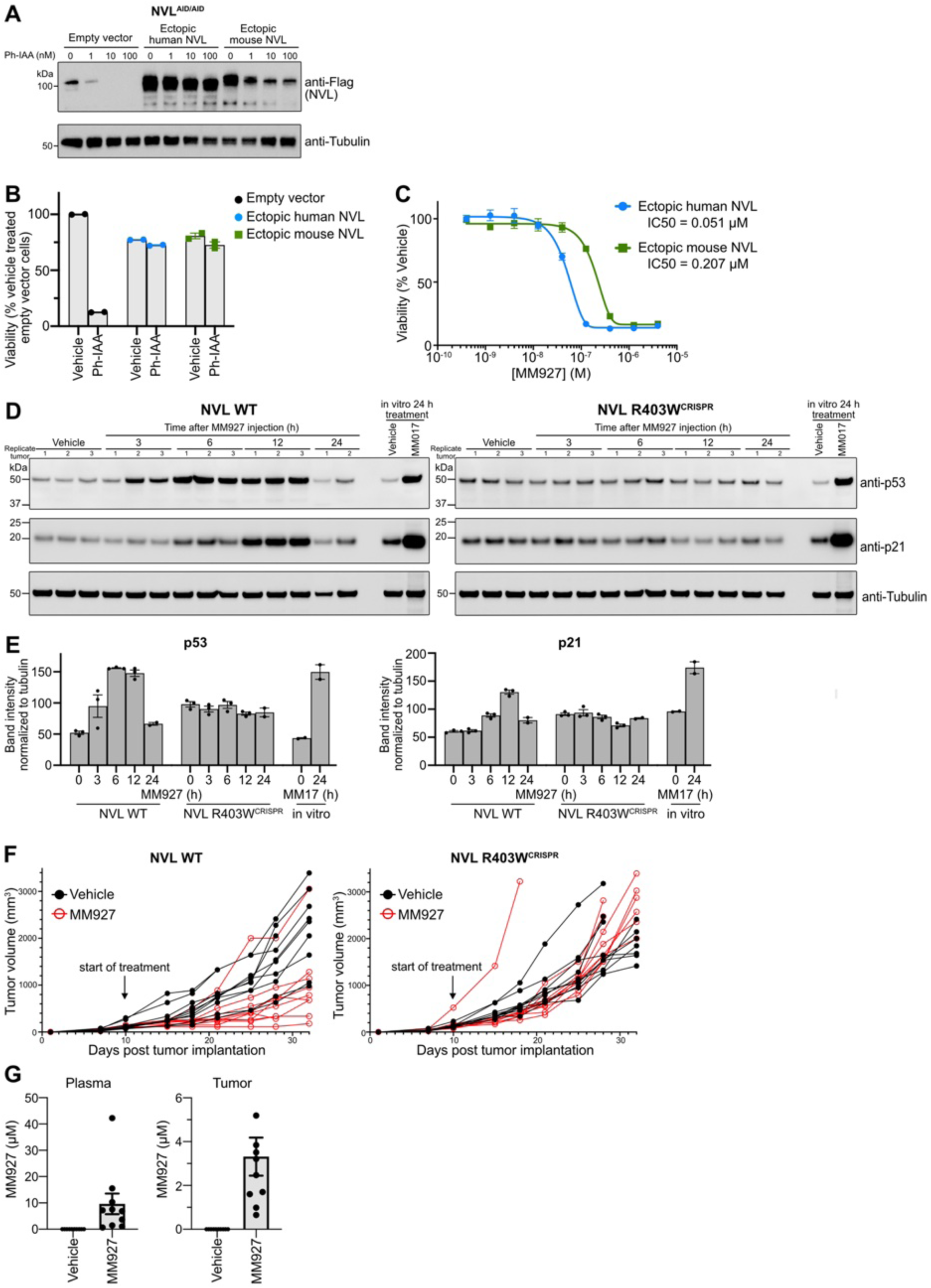
MM927 studies in HCT116 tumor xenografts. **A.** Immunoblots of Flag (NVL) and tubulin (loading control) in HCT116 NVL^AID/AID^ cells ectopically expressing empty vector, human NVL, or mouse NVL treated with Ph-IAA for 6 hours. **B.** Viability assay of cell lines in (A) treated with 0.03 µM Ph-IAA for 72 hours (n = 2 biological replicates, mean ± sem). **C.** Viability assay of HCT116 NVL^AID/AID^ cells ectopically expressing human NVL or mouse NVL in treated with MM927 and 0.03 µM Ph-IAA for 72 hours (n = 2 biological replicates; mean ± sem). **D, E.** Immunoblots (**D**) and quantitation (**E**) of p53, p21, and tubulin (loading control) of HCT116 NVL WT and NVL R403W^CRISPR^ tumor xenografts implanted in either flank of the same mouse treated with one 35 mg/kg MM927 IP injection. Right two lanes are *in vitro* samples of HCT116 cells treated with 10 µM MM17 for 24 hours. (n=2 to 3 biological replicates, mean ± sem). **F.** Individual tumor volume measurements in Fig. 5E. Each line represents a mouse. **G.** Pharmacokinetic quantitation of MM927 3 hours after the last dose in the mouse plasma and tumors (n = 10 mice/group, mean ± sem).

**Figure S15.**
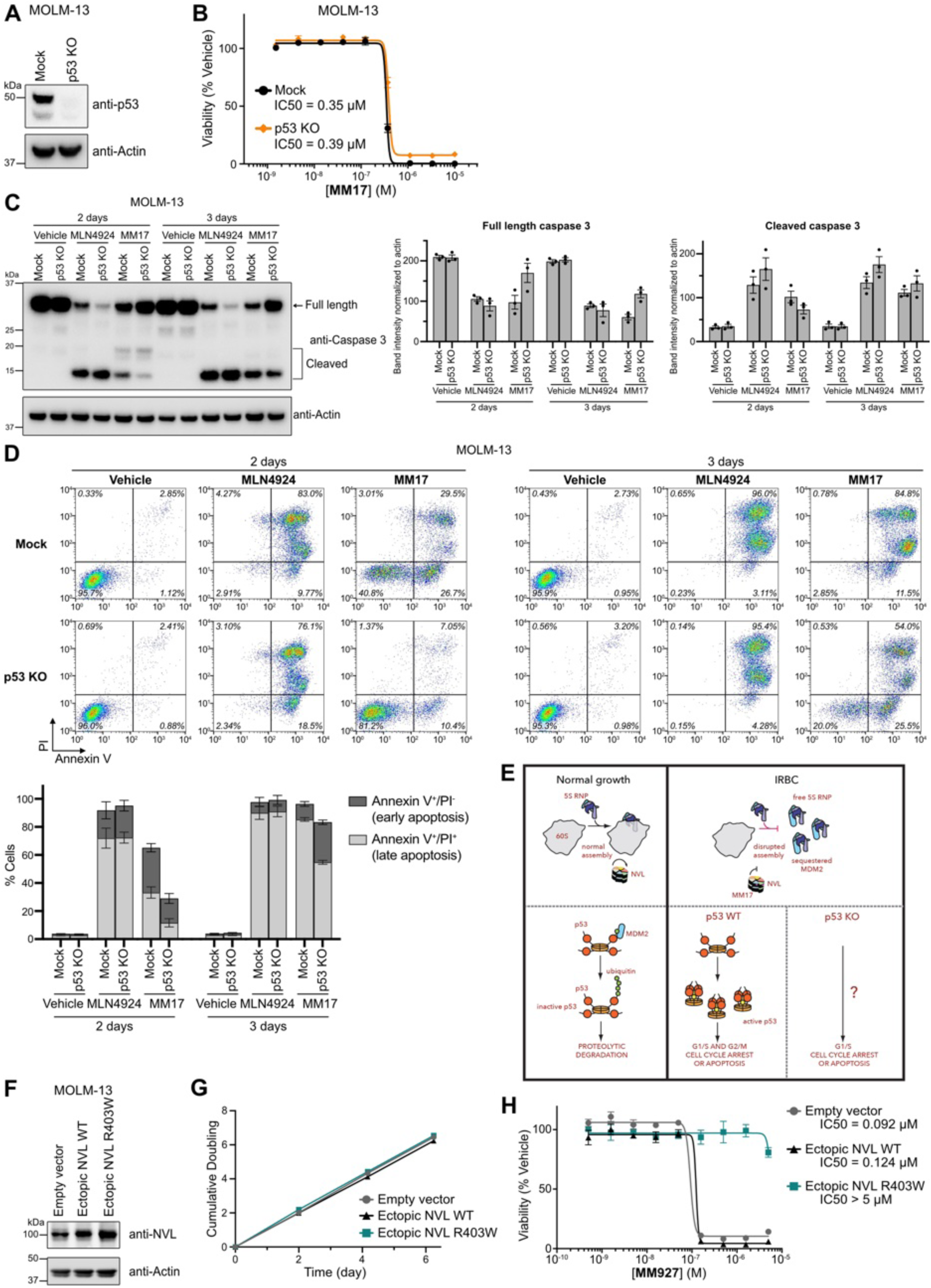
In vitro studies of NVL inhibitors in MOLM-13. **A.** Immunoblots of p53 and actin (loading control) in MOLM-13 mock engineered or p53 knockout (KO) cells. **B.** Viability assay of cell lines in (A) treated with MM17 for 72 hours. (n = 3 biological replicates, mean ± sem). **C.** Immunoblots and quantitation of caspase-3 and actin (loading control) in MOLM-13 mock and p53 KO cells treated with 1 µM MLN4924 or 3 µM MM17. (n = 3 biological replicates, mean ± sem). **D.** Flow cytometry-based apoptosis analysis and quantitation in MOLM-13 mock and p53 KO cells treated with 1 µM MLN4924 or 3 µM MM17. (n = 3 biological replicates, mean ± sem). **E.** Schematic illustrating the proposed mechanism of cellular response to MM17. **F.** Immunoblots of NVL and actin (loading control) in MOLM-13 cells expressing empty vector, NVL WT, or NVL R403W. **G.** Growth curves of cell lines in (F). n = 3 biological replicates, mean ± sem). **H.** Viability assay of cell lines in (F) treated with MM927 for 72 hours. (n = 2 biological replicates, mean ± sem).

**Figure S16.**
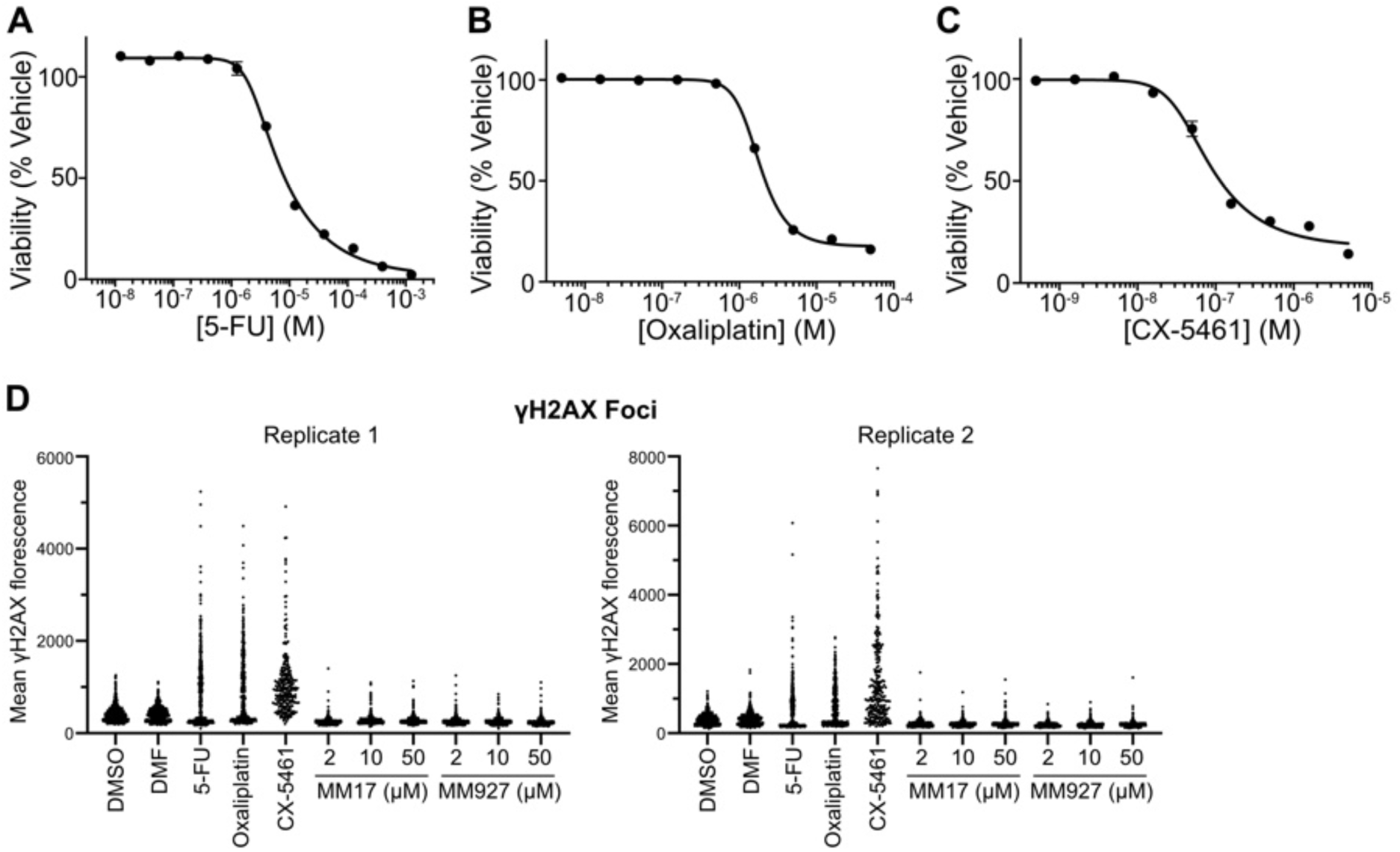
NVL inhibitors do not cause DNA damage. A, B, **C.** Viability assay of HCT116 cells treated with 5-FU (**A**), oxaliplatin (**B**), or CX-5461 (**C**) for 72 hours. (n = 3 biological replicates, mean ± sem). **C.** Quantitation of 2 additional biological replicates of γH2AX immunofluorescence staining, related to Fig. 6A. DMF is the vehicle control for oxaliplatin. DMSO is the vehicle control for all other compounds.

**Table S1.** Summary of previously published compounds that influence ribosome biogenesis.

**Table S2.** Sequencing analysis of compound resistant clones.

**Table S3.** Cryo-EM data collection, refinement, and validation statistics.

**Table S4.** CRISPR/Cas9 knockout screen gene comparison.

**Table S5.** p53 hotspot mutation and copy number analysis of MM17 activity in the PRISM screen.

**Table S6.** Compound viability profile correlation of MM17 in the PRISM screen.

**Data S1.** SI NMR LC for MM17, MM514, MM524, MM691, and MM927.

**Data S2.** PDB EM validation report for the Cryo-EM structure of apo NVL.

**Data S3.** PDB EM validation report for the Cryo-EM structure of NVL bound to MM17.

**Data S4.** PDB EM validation report for the Cryo-EM structure of NVL bound to MM927.

**Data S5.** R source file for the PRISM screen.

**Data S6.** Fiji and QuPath scripts for quantification of microscopy images.

**Data S7.** Full gel and microscopy images.

**Movie S1.** Allosteric disruption of the NVL D/E subunit interface by MM17. Morph animation of the subunit D/E interface within the D1 AAA+ domain, illustrating the conformational transition from apo-NVL^dEQ^ to the MM17-bound structure. Structural changes were visualized using PyMOL and highlight the ligand-induced rearrangements of the adjacent nucleotide binding pocket.

