## Supplementary material for "A small molecule inhibitor of NVL suppresses tumor growth by blocking ribosome biogenesis": Table S1

**Table S1 - List of compounds targeting ribosome biogenesis.**

| Inhibitors of rRNA Synthesis (RNA Polymerase I – Mediated rRNA Transcription) | | | | | |
|---|---|---|---|---|---|
| **Compound** | **Structure** | **Molecular target** | **Proposed mechanism of action** | **Status** | **References** |
| **CX-5461** (Pidnarulex) | 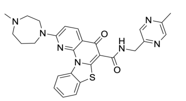 | **Unknown (DNA).** | Inhibitor of RNA polymerase I transcription; proposed to stabilize rDNA G-quadruplex structures and prevent binding of the SL1 transcription factor to the rRNA gene promoter, thereby blocking formation of the Pol I pre-initiation complex. Induces nucleolar stress and p53 activation, and at higher concentrations, can poison topoisomerase II. | **Investigational (Phase I/II clinical trials)** | *(19) (20) (21) (22) (23) (24) (25)* |
| **CX-3543** (Quarfloxin) | 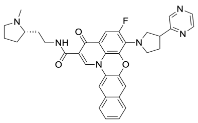 | **Unknown (DNA).** | G-quadruplex DNA stabilizer; binds G-rich sequences in rDNA and displaces nucleolin, thereby inhibiting rRNA transcription by RNA polymerase I. Induces nucleolar apoptosis in cancer cells. | **Investigational (completed Phase II trial; not FDA-approved)** | *(18)* |
| **BMH-21** | 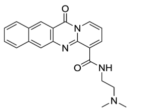 | **Unknown (DNA).** | 9-Aminoacridine derivative that intercalates into GC-rich rDNA and inhibits RNA polymerase I transcription elongation. | **Investigational compound (preclinical)** | *(17)* |
| **Ellipticine** | 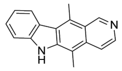 | **Unknown (DNA).** | Planar polycyclic alkaloid that intercalates into GC-rich rDNA and selectively blocks RNA polymerase I–mediated rRNA synthesis by preventing SL1 binding at the promoter. Functions independently of ATM/ATR signaling and inhibits topoisomerase II at higher doses. | **Experimental (ellipticine derivatives were tested in trials but halted due to toxicity)** | *(14)* |
| **Hernandonine** | 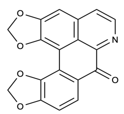 | **Unknown (DNA).** | DNA-intercalating natural polycyclic alkaloid that inhibits rRNA synthesis, induces nucleolar stress, and promotes POLR1A degradation. Triggers apoptosis in tumor cells at low micromolar doses. | **Experimental (preclinical)** | *(16)* |
| **Sempervirine** | 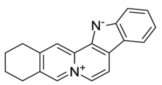 | **Unknown.** | Natural alkaloid from Gelsemium that disrupts ribosome biogenesis by inducing nucleolar stress and degrading the Pol I subunit RPA194. Inhibits rRNA synthesis and remains active in p53-deficient cells via E2F1 suppression. | **Experimental (preclinical)** | *(15)* |

| Inhibitors of Ribosome Assembly (Ribosomal Subunit Maturation) | | | | | |
|---|---|---|---|---|---|
| **Compound** | **Structure** | **Molecular target** | **Proposed mechanism of action** | **Status** | **References** |
| **Diazaborine** | 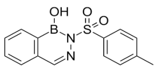 | **Drg1 AAA+ ATPase** | Benzoyl naphthyridinone that inhibits the AAA+ ATPase Drg1, blocking 60S subunit assembly by preventing ATP-dependent release of Rlp24 from cytoplasmic pre-60S particles. | **Research tool (YEAST - S.cerevisiae)** | *(36)* |
| **Ribozinoindole (Rbin-1)** | 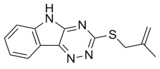 | **Mdn1 AAA+ ATPase.** | Triazinoindole small molecule that inhibits the nuclear AAA+ ATPase Mdn1, blocking pre-60S maturation in the nucleolus by preventing release of assembly factors. | **Research tool (YEAST - S.pombe)** | *(38)* |
| **RBI-1/RBI-2** | 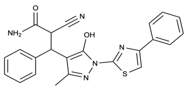 | **Unknown.** | Small molecules identified through phenotypic screening. RBI-2 has been reported to induce polyadenylation and exosome-mediated degradation of pre-rRNA transcripts without affecting RNA polymerase I binding or occupancy at rDNA loci. The molecular target remains unknown. | **Experimental (preclinical)** | *(27)* |
| **Haemanthamine** | 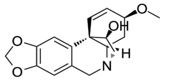 | **60S ribosomal A-site - link to biogenesis unknown.** | Amaryllidaceae alkaloid that binds the 60S ribosomal subunit A-site (inhibiting translation elongation). In human cancer cells, haemanthamine causes activation of the impaired ribosome biogenesis checkpoint (IRBC) and nucleolar accumulation of ribosomal proteins via an unknown mechanism. | **Experimental natural product (preclinical)** | *(26)* |
| **Portimines** | 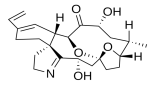 | **60S export factor NMD3.** | Natural product cytotoxin. Evidence that it binds to an essential ribosome biogenesis factero required for 60S export. Mechanism unknown. | **Experimental natural product (preclinical)** | *(28)* |
