## Supplementary material for "A small molecule inhibitor of NVL suppresses tumor growth by blocking ribosome biogenesis": Table S3

**Table S3 - Cryo-EM data collection, refinement, and validation statistics**

| | NVL - apo | NVL – MM17 | NVL – MM297 |
|---|---|---|---|
| **Data collection and processing** | | | |
| Magnification | 105,000x | 105,000x | 105,000x |
| Voltage (kV) | 300 | 300 | 300 |
| Electron exposure ($e^-$/Å$^2$) | 48.4 | 48.4 | 30 |
| Defocus range (μm) | -0.6-(-1.9) | -0.6-(-1.9) | -0.6-(-1.9) |
| Pixel size (Å) | 0.83 | 0.83 | 0.827 |
| Symmetry imposed | C1 | C1 | C1 |
| Initial particle images (no.) | 1,340,199 | 626,020 | 1527471 |
| Final particle images (no.) | 144,455 | 149,567 | 193335 |
| Map resolution (Å) | 2.83 | 3.05 | 2.86 |
| FSC threshold | 0.143 | 0.143 | 0.143 |
| Map resolution range (Å) | 1.66-10 | 1.66-10 | 1.66-10 |
| **Refinement** | | | |
| Model resolution (Å) | 2.94 | 3.05 | 2.8 |
| FSC threshold | 0.5 | 0.5 | 0.5 |
| Map sharpening *B* factor (Å$^2$) | 69.1 | 71.1 | 84.7 |
| Model composition | | | |
| Non-hydrogen atoms | 27178 | 27068 | 26022 |
| Protein residues | 3461 | 3461 | 3327 |
| Ligands | 20 | 20 | 19 |
| Waters | 138 | 0 | 20 |
| *B* factors (Å$^2$) | | | |
| Protein | 30 | 66 | 46.9 |
| Ligand | 65.13 | 53 | 39.6 |
| R.m.s. deviations | | | |
| Bond lengths (Å) | 0.002 | 0.002 | 0.003 |
| Bond angles (°) | 0.494 | 0.523 | 0.551 |
| Validation | | | |
| MolProbity score | 1.49 | 1.45 | 1.04 |
| Clashscore | 9.13 | 8.17 | 2.58 |
| Poor rotamers (%) | 0.03 | 0 | 0 |
| Ramachandran plot | | | |
| Favored (%) | 98.1 | 98.0 | 98.1 |
| Allowed (%) | 1.9 | 2 | 1.9 |
| Disallowed (%) | 0 | 0 | 0 |
