## Supplementary material for "A small molecule inhibitor of NVL suppresses tumor growth by blocking ribosome biogenesis": Data S1

Copies of $^{1}H$ and $^{13}C$ Spectra

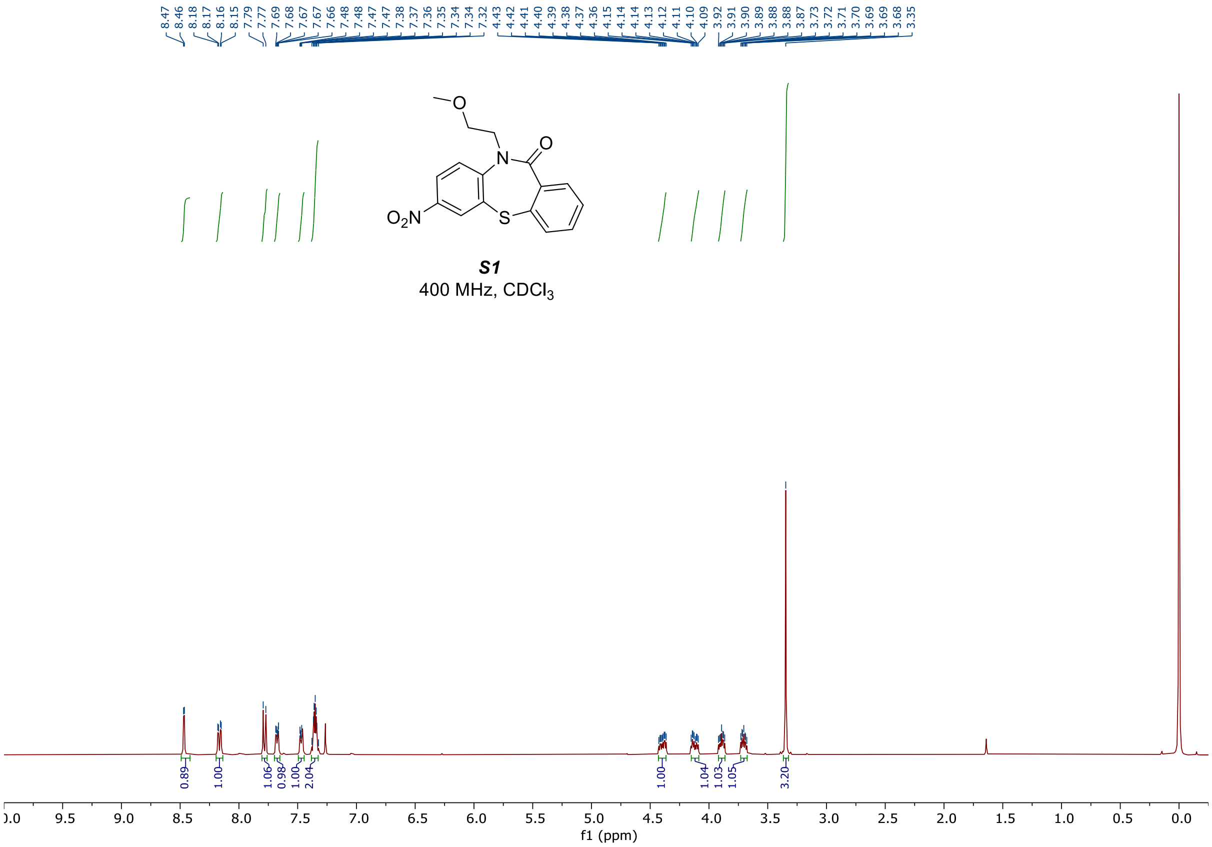
8.47 8.46 8.18 8.17 8.16 8.15 7.79 7.77 7.69 7.68 7.67 7.67 7.66 7.48 7.48 7.47 7.47 7.38 7.37 7.36 7.35 7.34 7.34 7.32 4.43 4.42 4.41 4.40 4.39 4.38 4.37 4.36 4.15 4.14 4.14 4.13 4.12 4.11 4.10 4.09 3.92 3.91 3.90 3.89 3.88 3.88 3.87 3.73 3.72 3.71 3.70 3.69 3.69 3.68 3.35
O
N
O
$O_2N$
S
S1
400 MHz, $CDCl_3$
0.89 1.00 1.06 0.98 1.00 2.04 1.00 1.04 1.03 1.05 3.20
f1 (ppm)

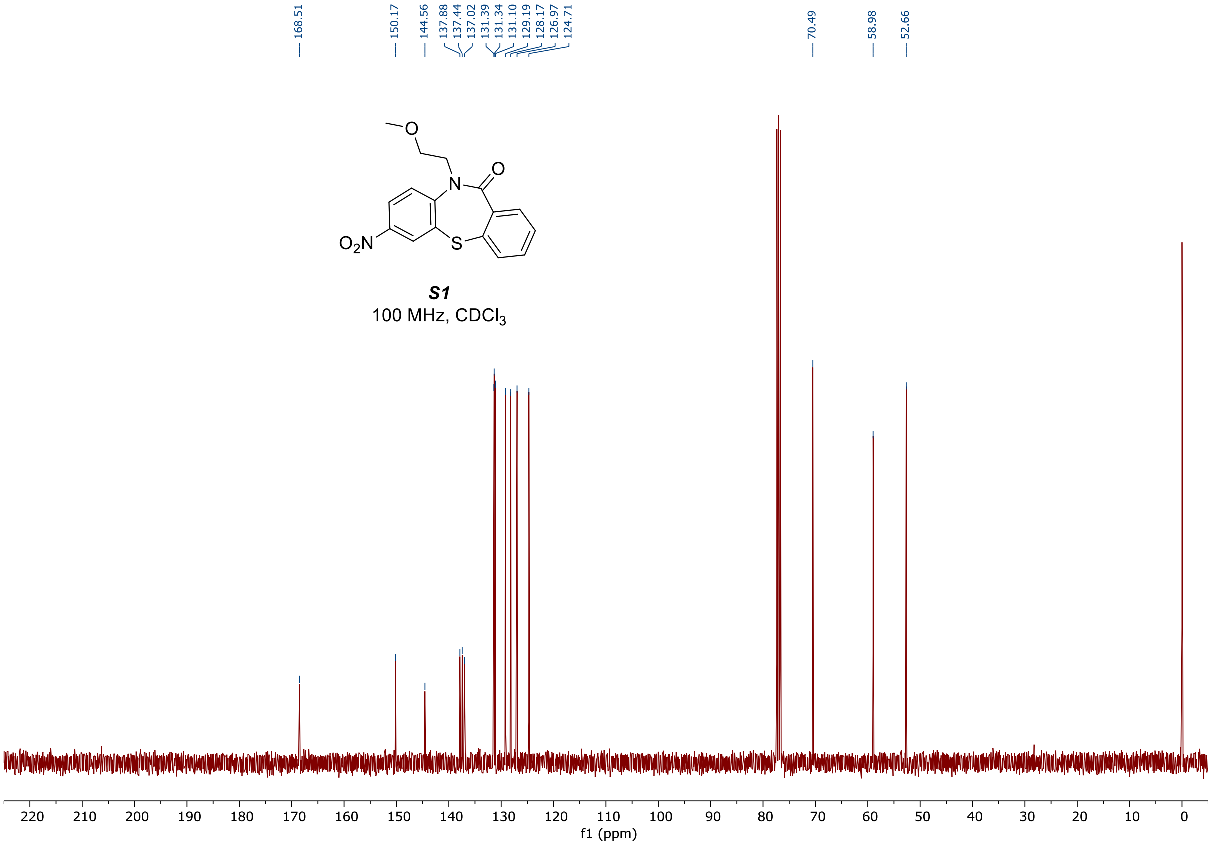
168.51 150.17 144.56 137.88 137.44 137.02 131.39 131.34 131.10 129.19 128.17 126.97 124.71 70.49 58.98 52.66
O
N
O
$O_2N$
S
S1
100 MHz, $CDCl_3$
f1 (ppm)

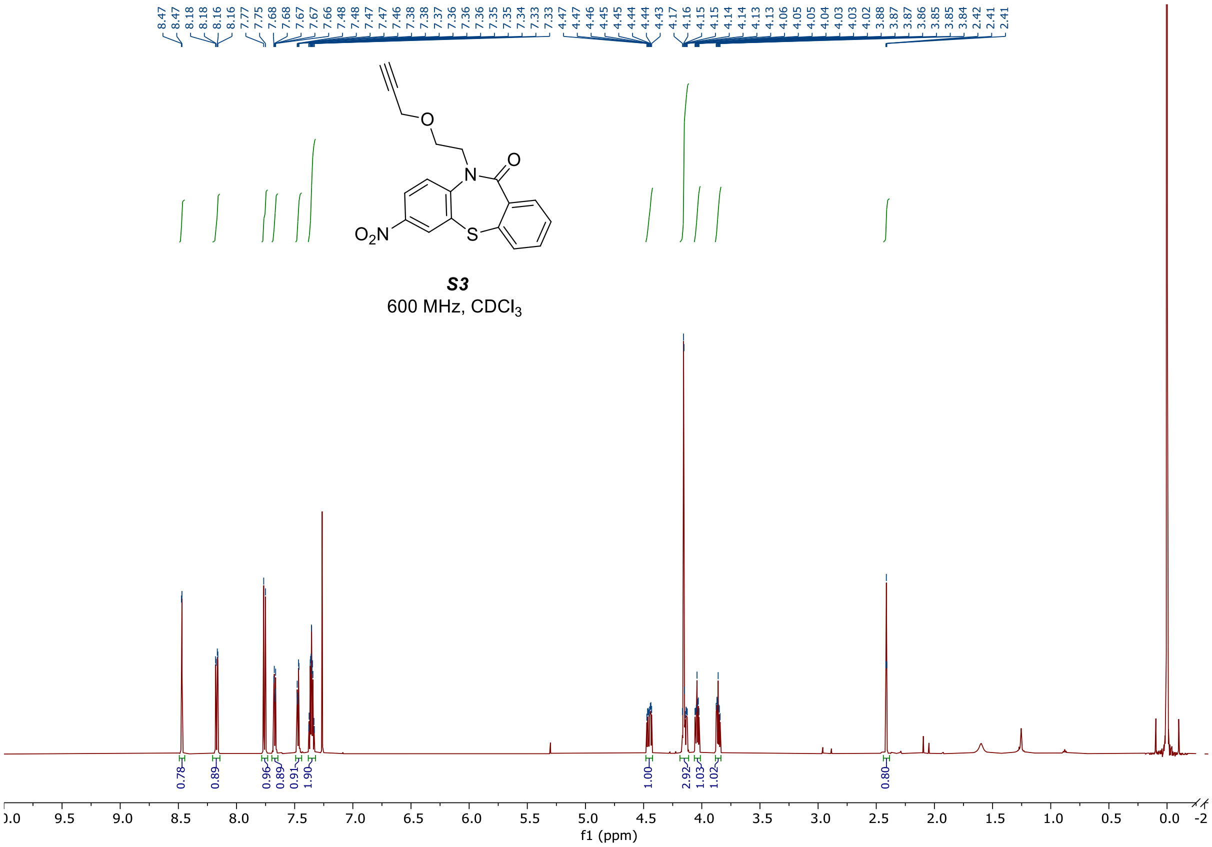

*S3*
600 MHz, $CDCl_3$

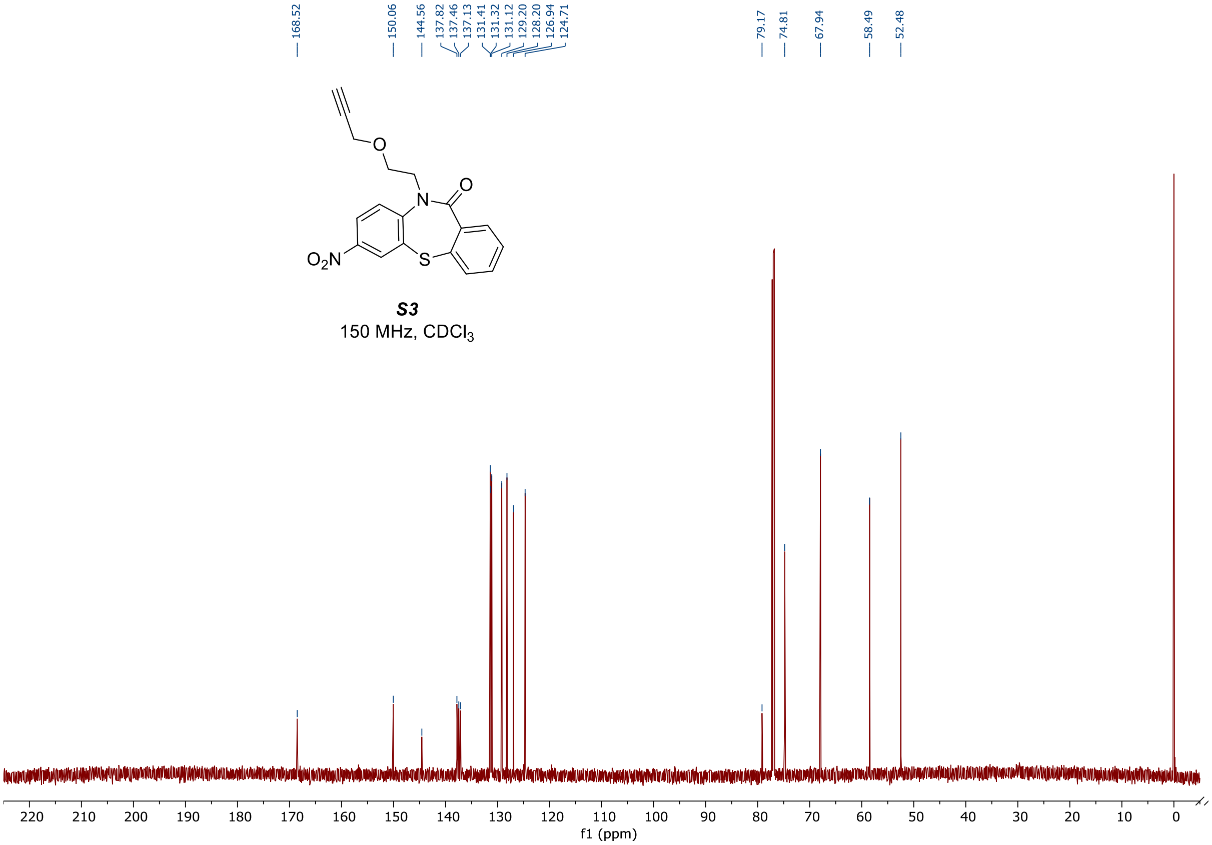

*S3*
150 MHz, $CDCl_3$

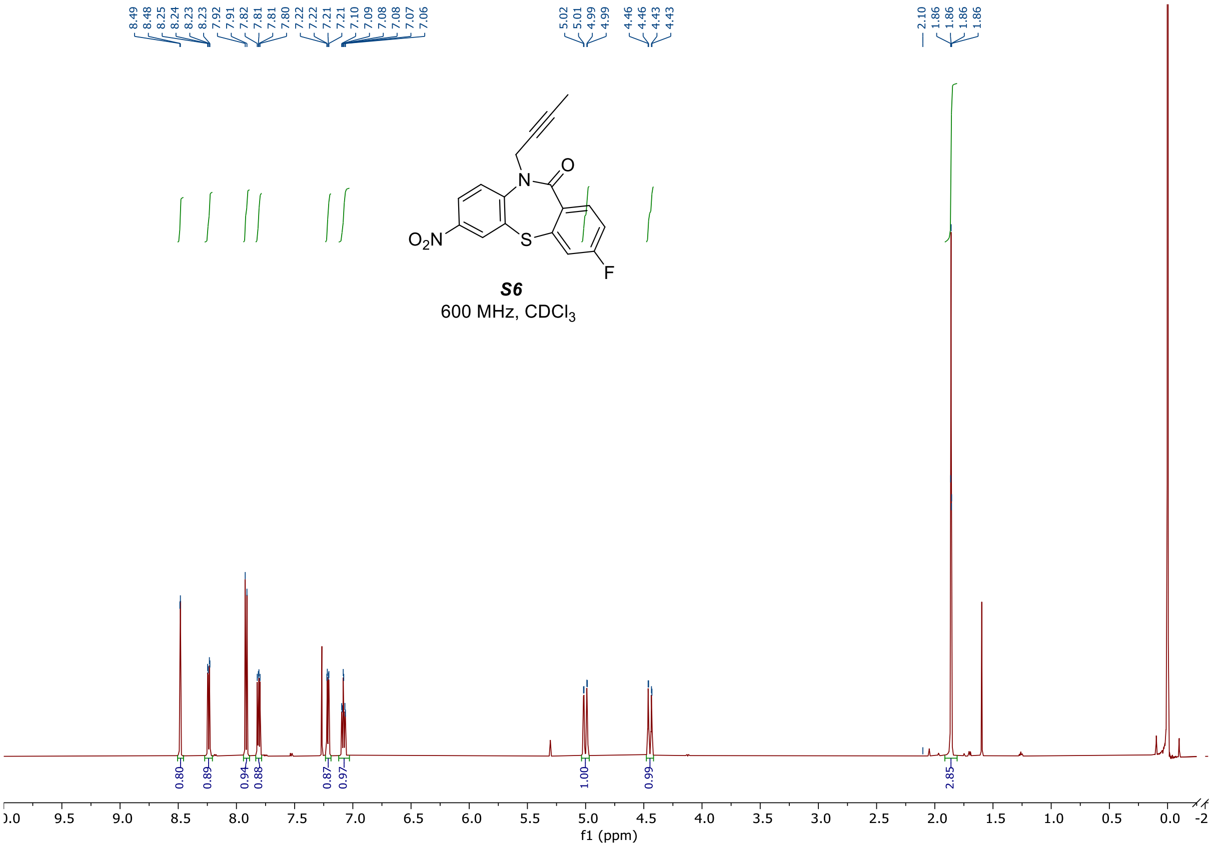
8.49
8.48
8.25
8.24
8.23
8.23
7.92
7.91
7.82
7.81
7.81
7.80
7.22
7.22
7.21
7.21
7.10
7.09
7.08
7.08
7.07
7.06
5.02
5.01
4.99
4.99
4.46
4.46
4.43
4.43
2.10
1.86
1.86
1.86
1.86
O2N
N
O
S
F
S6
600 MHz, CDCl3
0.80
0.89
0.94
0.88
0.87
0.97
1.00
0.99
2.85
f1 (ppm)

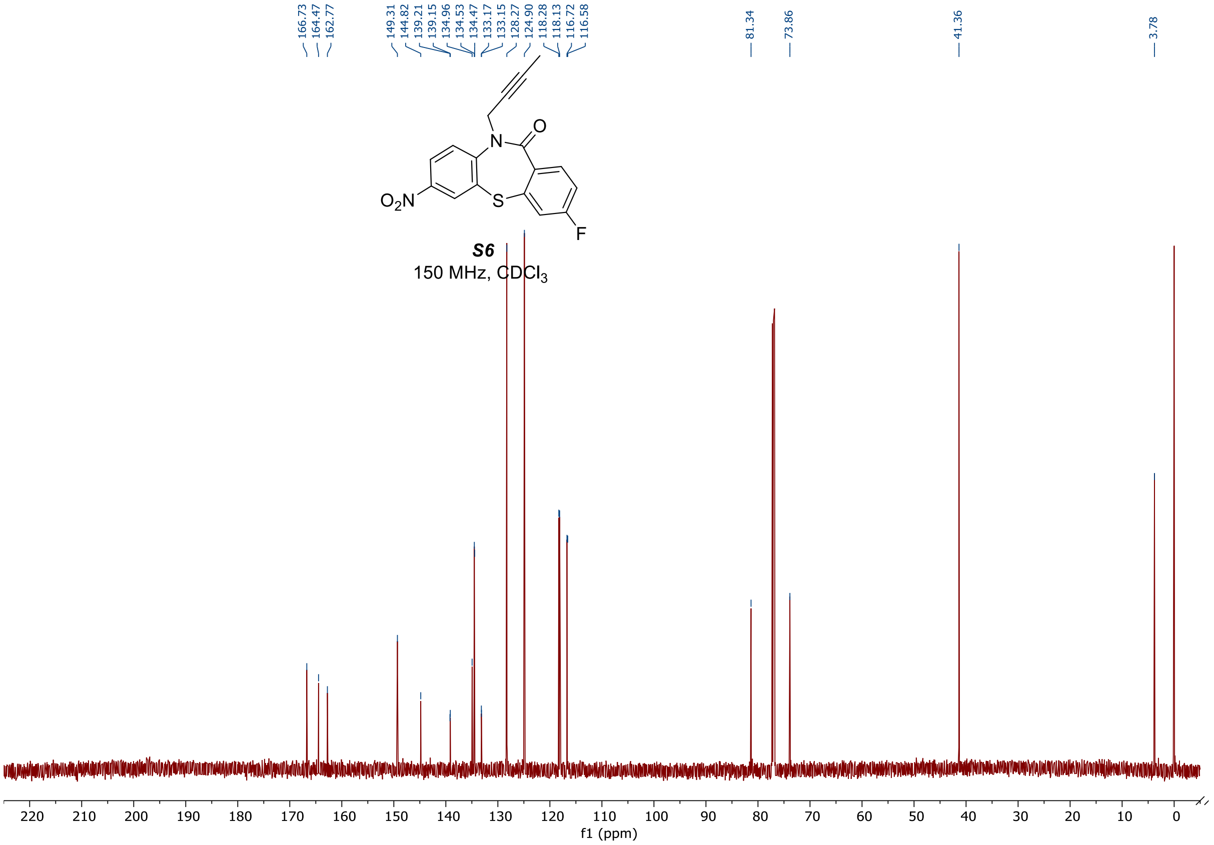
166.73
164.47
162.77
149.31
144.82
139.21
139.15
134.96
134.53
134.47
133.17
133.15
128.27
124.90
118.28
118.13
116.72
116.58
81.34
73.86
41.36
3.78
O2N
N
O
S
F
S6
150 MHz, CDCl3
f1 (ppm)

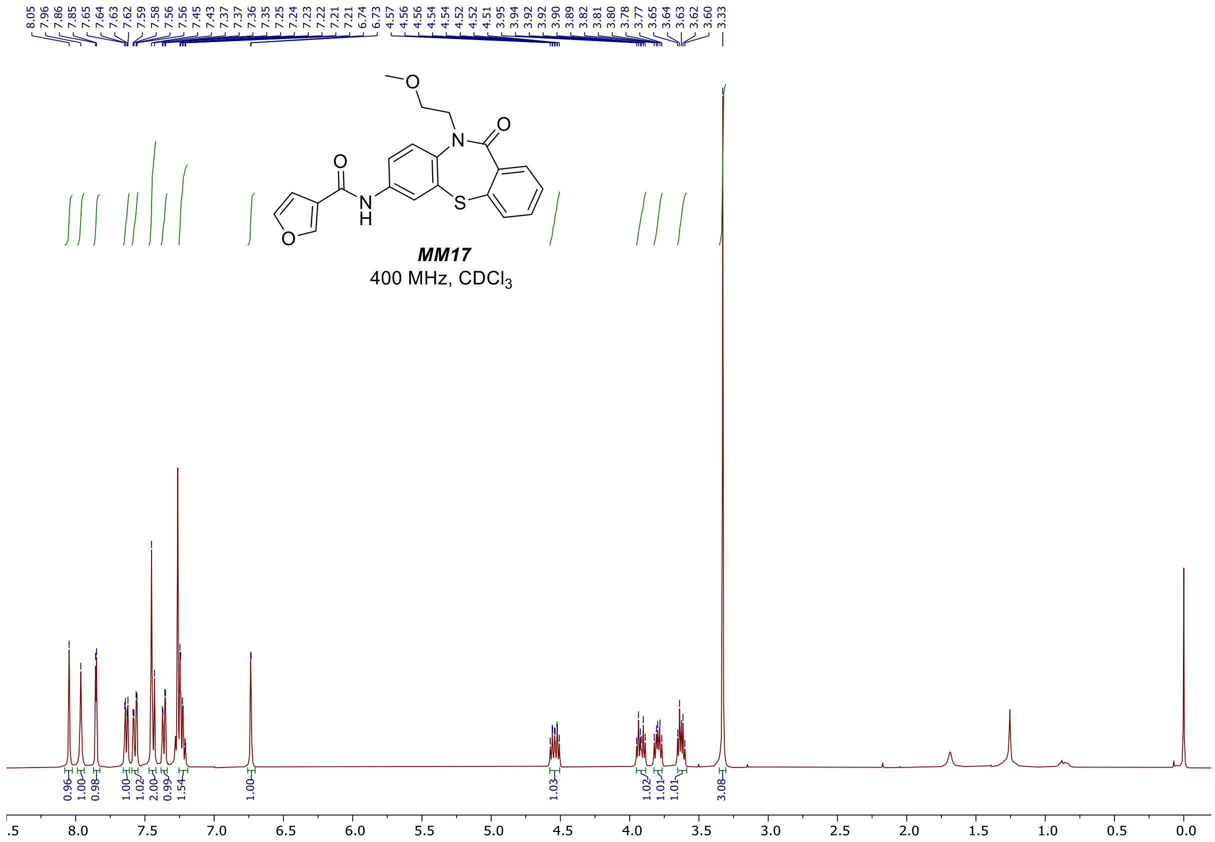

8.05 7.96 7.86 7.85 7.65 7.64 7.63 7.62 7.59 7.58 7.56 7.56 7.45 7.43 7.37 7.36 7.35 7.25 7.24 7.23 7.22 7.22 7.21 7.21 6.74 6.73 4.57 4.56 4.56 4.54 4.54 4.52 4.52 4.51 3.95 3.94 3.92 3.92 3.90 3.89 3.82 3.81 3.80 3.78 3.77 3.65 3.64 3.63 3.62 3.60 3.33
O
N
O
O
N
H
S
O
MM17
400 MHz, CDCl3
0.96 1.00 0.98 1.00 1.02 2.00 0.99 1.54 1.00 1.03 1.02 1.01 1.01 3.08
8.0 7.5 7.0 6.5 6.0 5.5 5.0 4.5 4.0 3.5 3.0 2.5 2.0 1.5 1.0 0.5 0.0


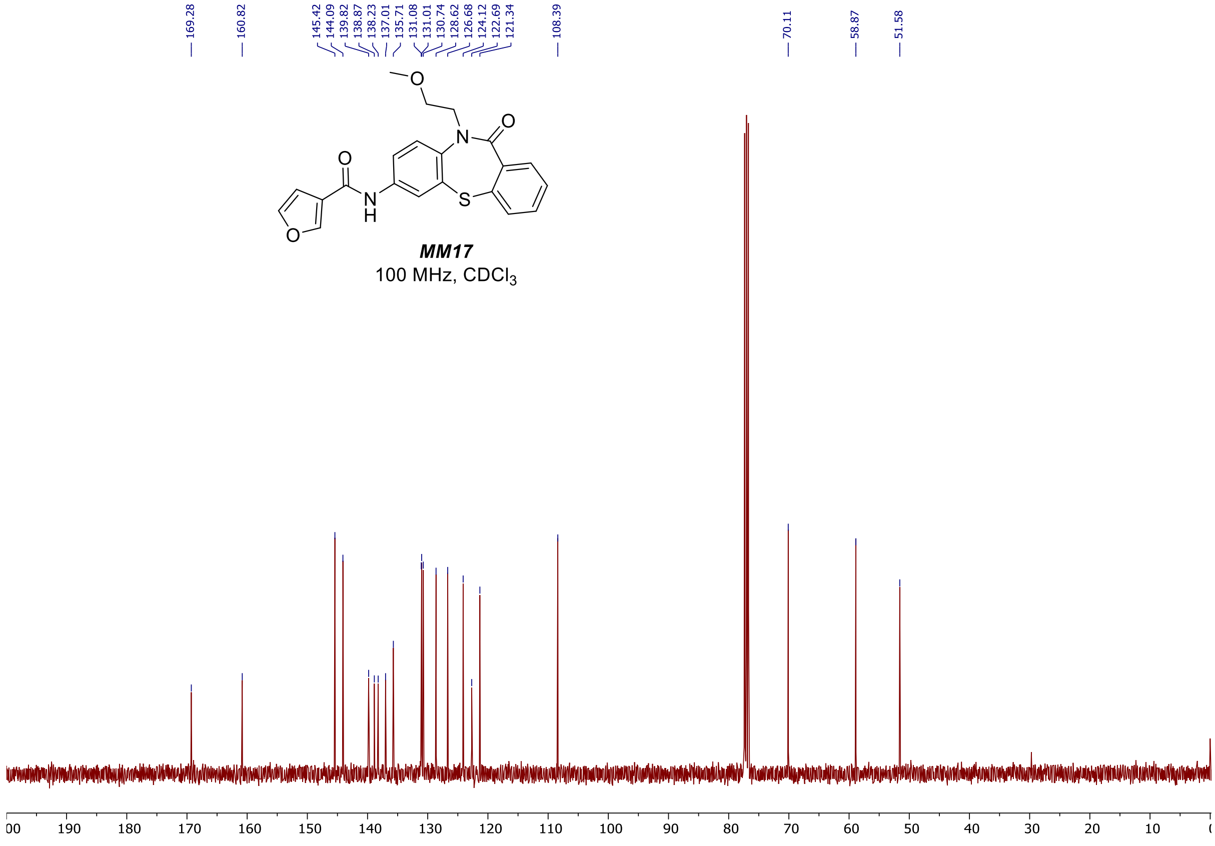

169.28 160.82 145.42 144.09 139.82 138.87 138.23 137.01 135.71 131.08 131.01 130.74 128.62 126.68 124.12 122.69 121.34 108.39 70.11 58.87 51.58
O
N
O
O
N
H
S
O
MM17
100 MHz, CDCl3
190 180 170 160 150 140 130 120 110 100 90 80 70 60 50 40 30 20 10

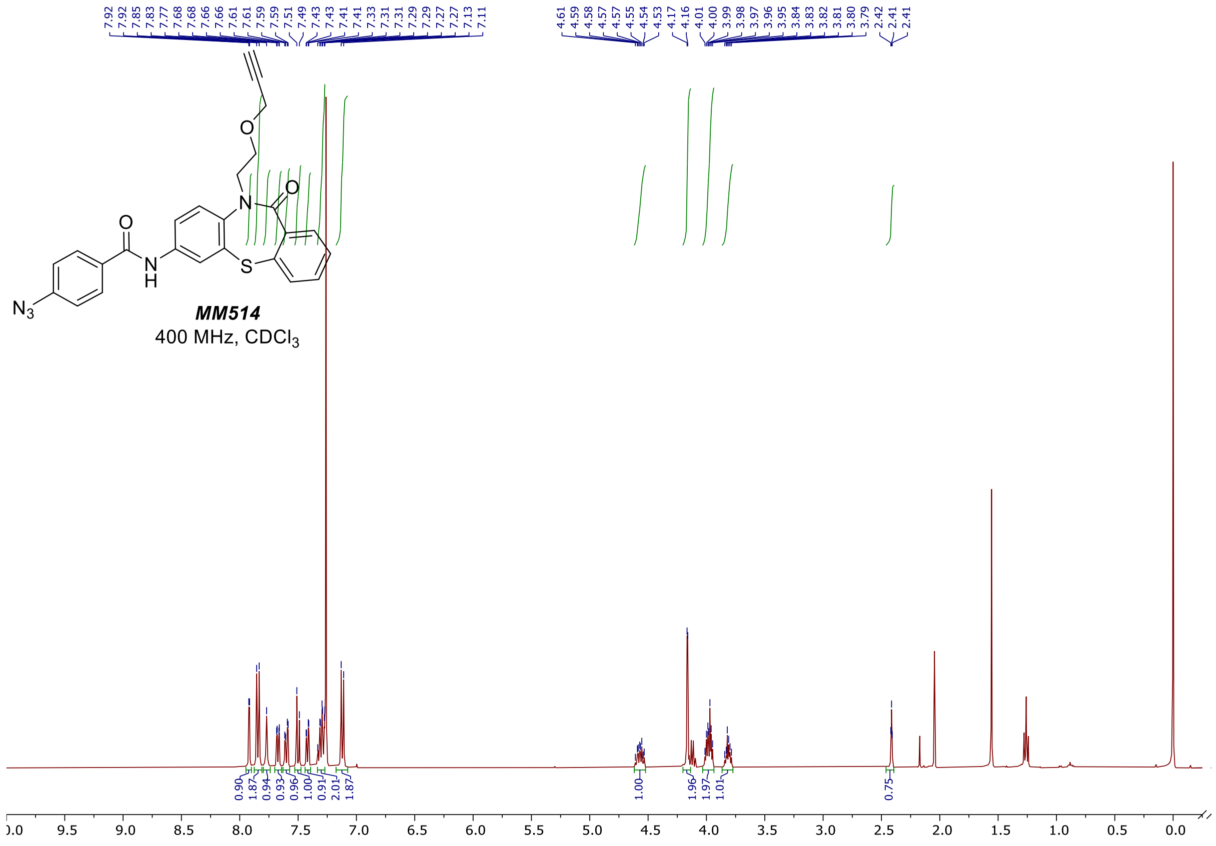
MM514
400 MHz, CDCl3

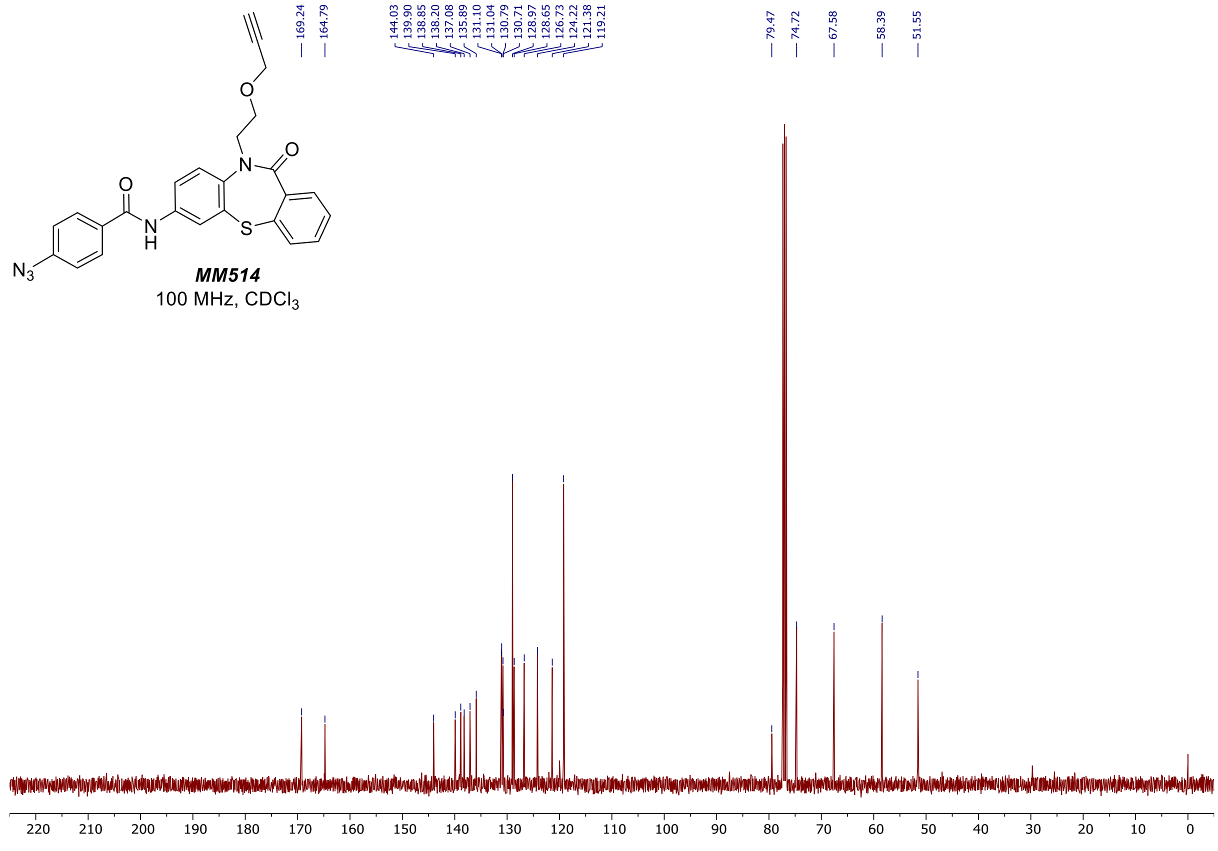
MM514
100 MHz, CDCl3
169.24
164.79
144.03
139.90
138.85
138.20
137.08
135.89
131.10
131.04
130.79
130.71
128.97
128.65
126.73
124.22
121.38
119.21
79.47
74.72
67.58
58.39
51.55

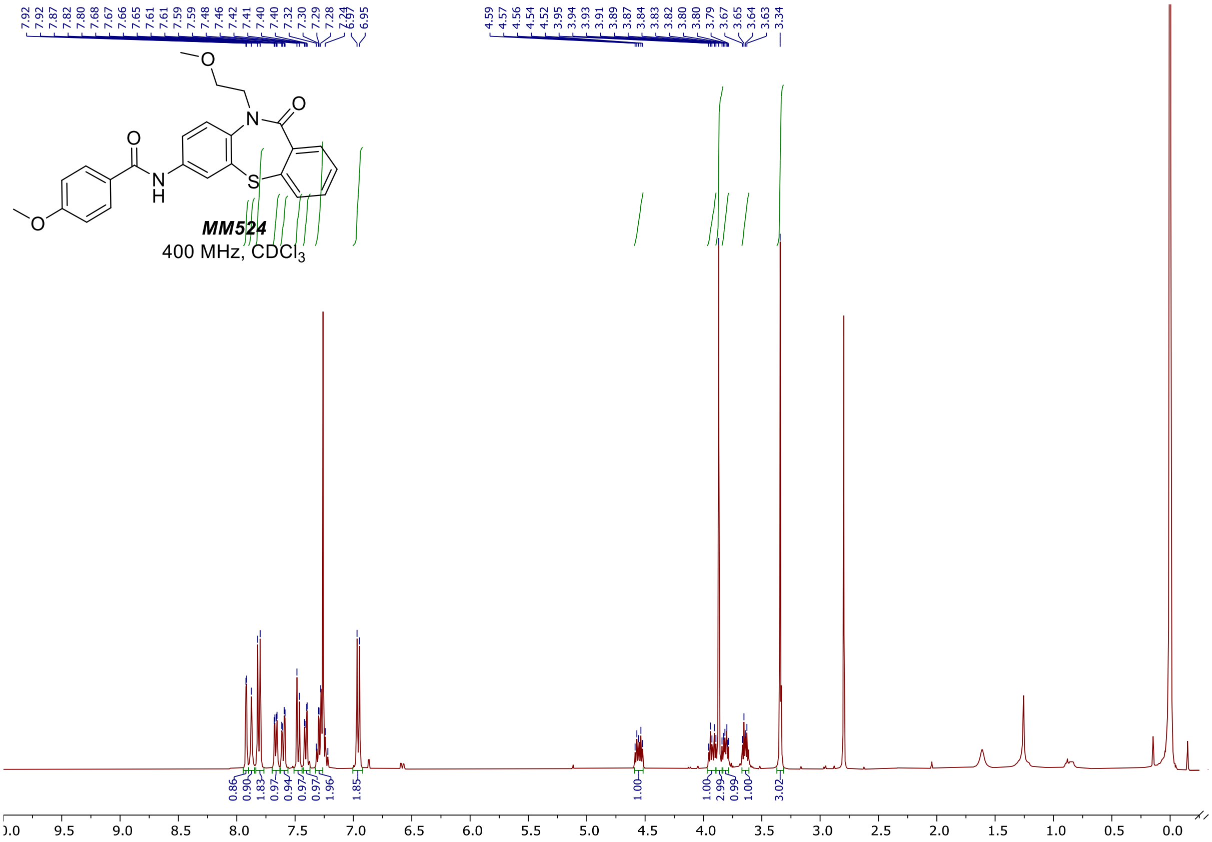
7.92 7.92 7.87 7.82 7.80 7.68 7.67 7.66 7.65 7.61 7.61 7.59 7.59 7.48 7.46 7.42 7.41 7.40 7.40 7.32 7.30 7.29 7.28 7.24 6.97 6.95
4.59 4.57 4.56 4.54 4.52 3.95 3.94 3.93 3.91 3.89 3.87 3.84 3.83 3.82 3.80 3.80 3.79 3.67 3.65 3.64 3.63 3.34
MM524
400 MHz, CDCl3
0.86 0.90 1.83 0.97 0.94 0.97 0.97 1.96 1.85 1.00 1.00 2.99 0.99 1.00 3.02
10.0 9.5 9.0 8.5 8.0 7.5 7.0 6.5 6.0 5.5 5.0 4.5 4.0 3.5 3.0 2.5 2.0 1.5 1.0 0.5 0.0

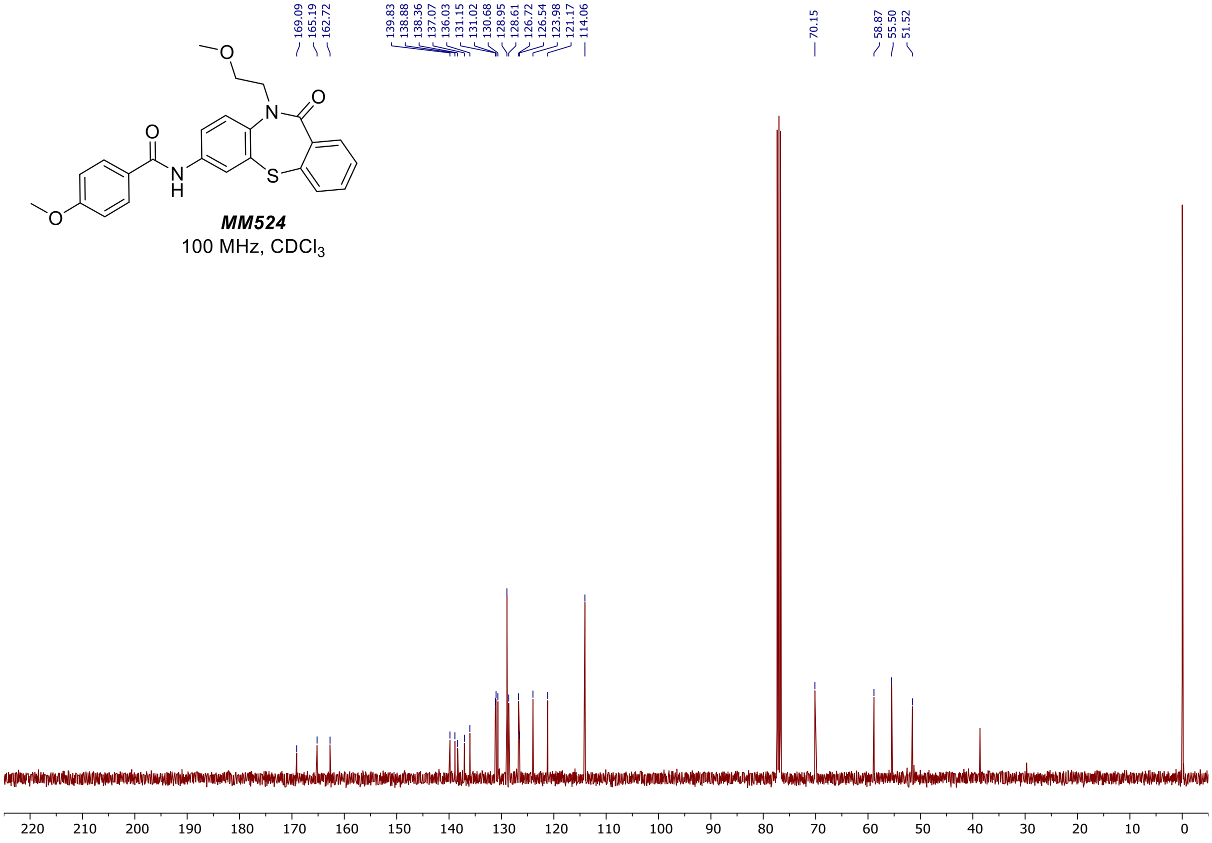
169.09 165.19 162.72
139.83 138.88 138.36 137.07 136.03 131.15 131.02 130.68 128.95 128.61 126.72 126.54 123.98 121.17 114.06
70.15
58.87 55.50 51.52
MM524
100 MHz, CDCl3
220 210 200 190 180 170 160 150 140 130 120 110 100 90 80 70 60 50 40 30 20 10 0

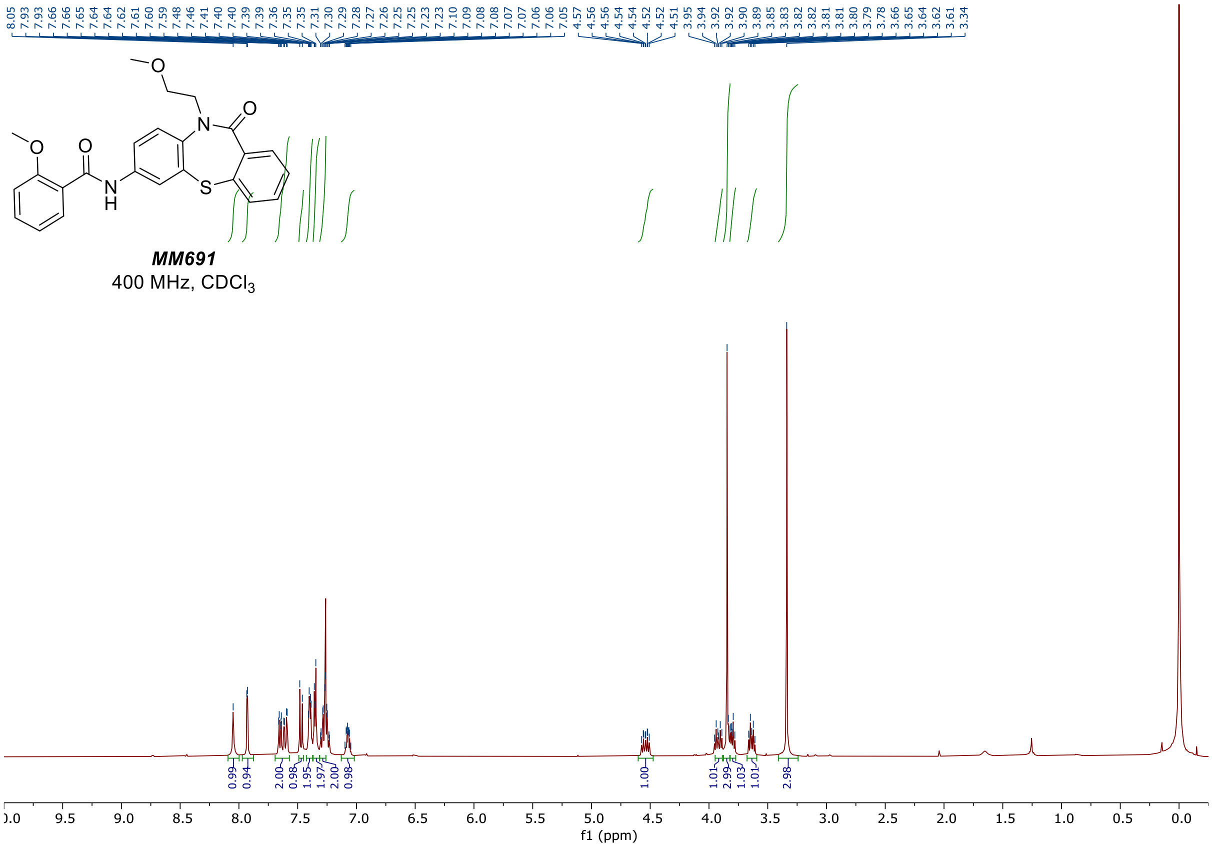
8.05 7.93 7.93 7.66 7.65 7.64 7.64 7.62 7.61 7.60 7.59 7.48 7.46 7.41 7.40 7.39 7.39 7.36 7.35 7.35 7.31 7.30 7.29 7.28 7.27 7.26 7.25 7.25 7.23 7.23 7.10 7.09 7.08 7.08 7.07 7.07 7.06 7.06 7.05 4.57 4.56 4.56 4.54 4.54 4.52 4.52 4.51 3.95 3.94 3.92 3.92 3.90 3.89 3.85 3.83 3.82 3.82 3.81 3.81 3.80 3.79 3.78 3.66 3.65 3.64 3.62 3.61 3.34
MM691
400 MHz, CDCl3
0.99 0.94 2.00 0.98 1.95 1.97 2.00 0.98 1.00 1.01 2.99 1.03 1.01 2.98
f1 (ppm)

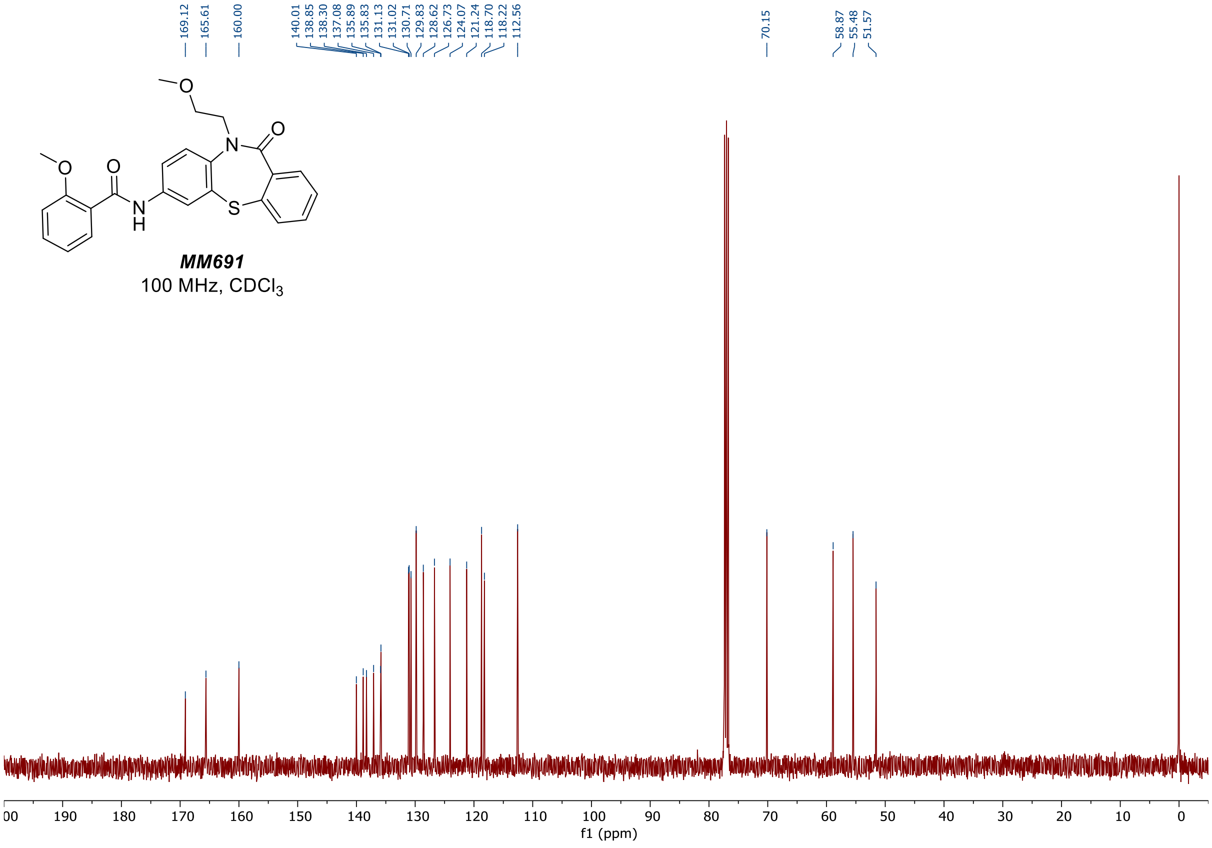
169.12 165.61 160.00 140.01 138.85 138.30 137.08 135.89 135.83 131.13 131.02 130.71 129.83 128.62 126.73 124.07 121.24 118.70 118.22 112.56 70.15 58.87 55.48 51.57
MM691
100 MHz, CDCl3
f1 (ppm)

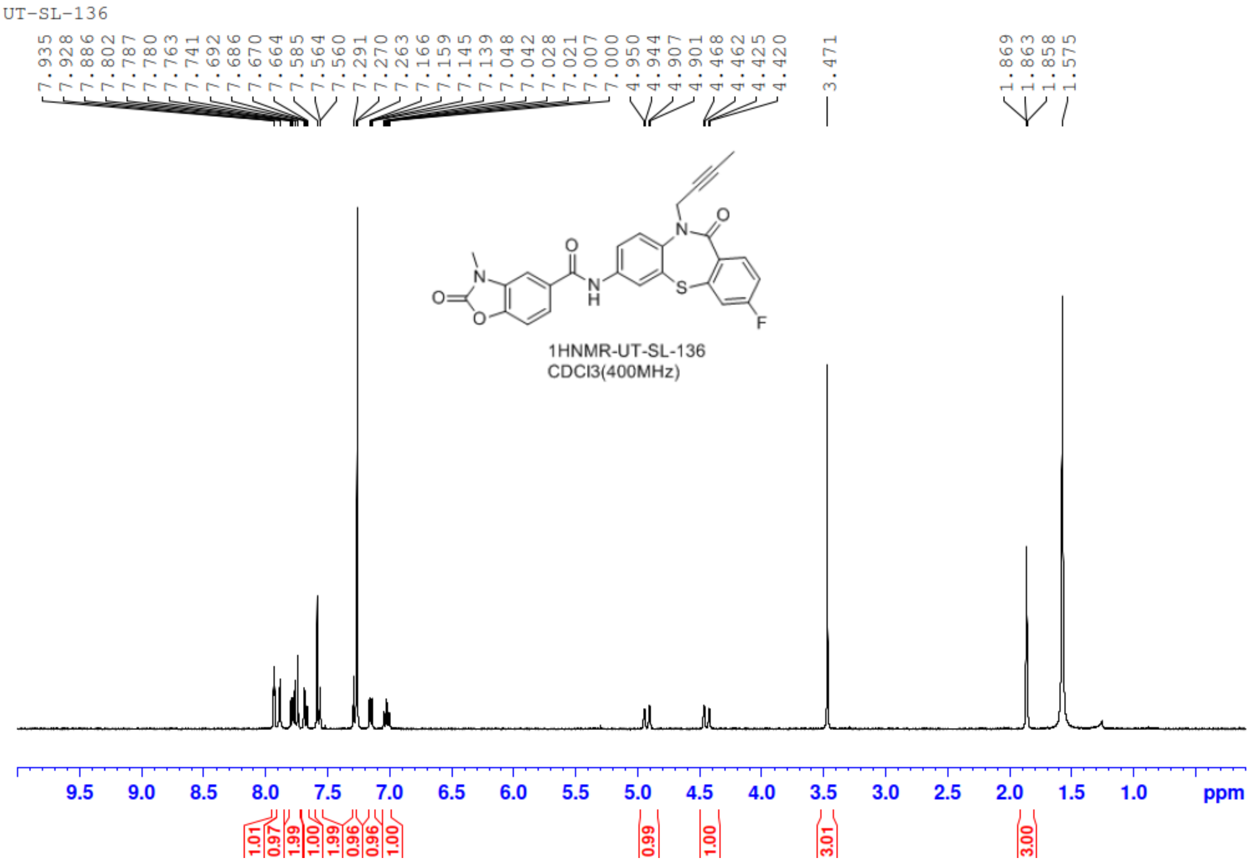
UT-SL-136
7.935 7.928 7.886 7.802 7.787 7.780 7.763 7.741 7.692 7.686 7.670 7.664 7.585 7.564 7.560 7.291 7.270 7.263 7.166 7.159 7.145 7.139 7.048 7.042 7.028 7.021 7.007 7.000 4.950 4.944 4.907 4.901 4.468 4.462 4.425 4.420
3.471
1.869 1.863 1.858 1.575
1HNMR-UT-SL-136
CDCl3(400MHz)
1.01 0.97 1.99 1.00 1.99 0.96 0.96 1.00 0.99 1.00 3.01 3.00
ppm

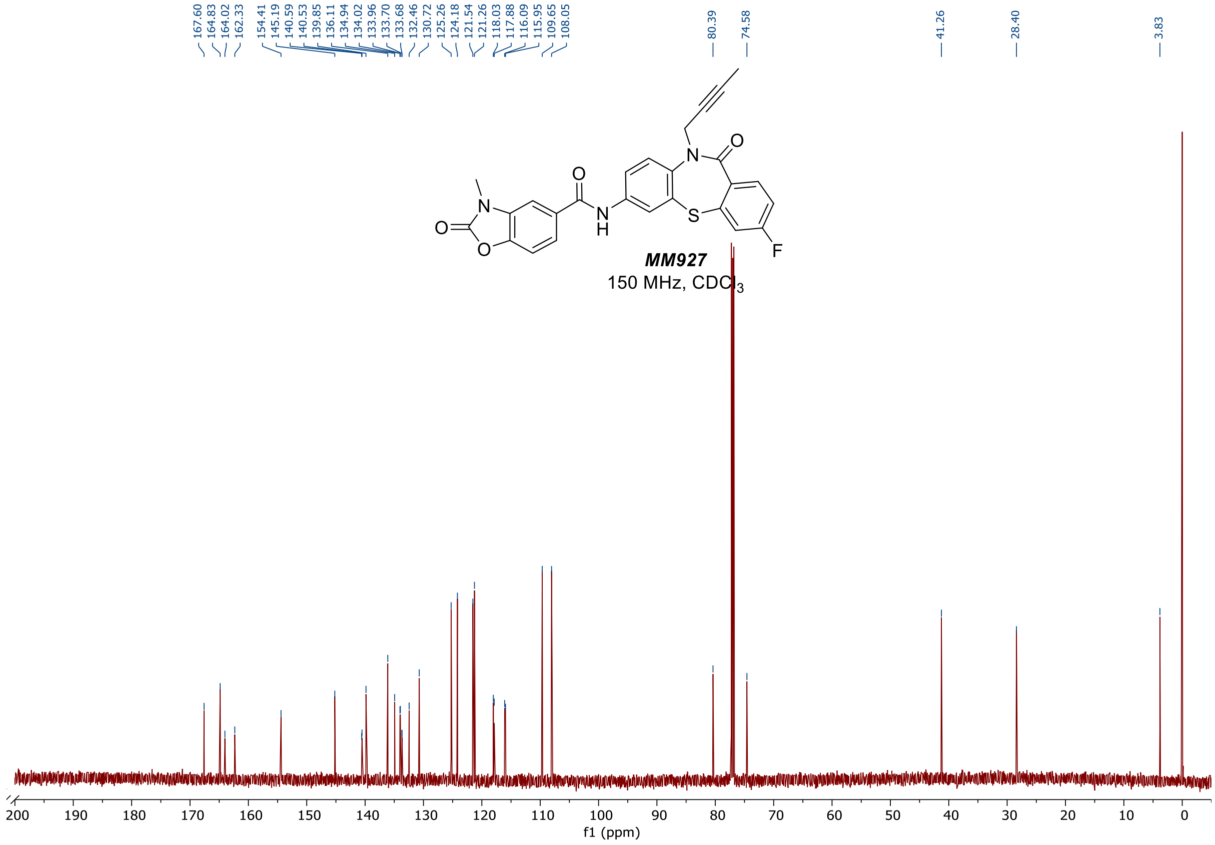
167.60 164.83 164.02 162.33 154.41 145.19 140.59 140.53 139.85 136.11 134.94 134.02 133.96 133.70 133.68 132.46 130.72 125.26 124.18 121.54 121.26 118.03 117.88 116.09 115.95 109.65 108.05
80.39
74.58
41.26
28.40
3.83
MM927
150 MHz, CDCl3
f1 (ppm)

HPLC Chromatograms (254 nM)

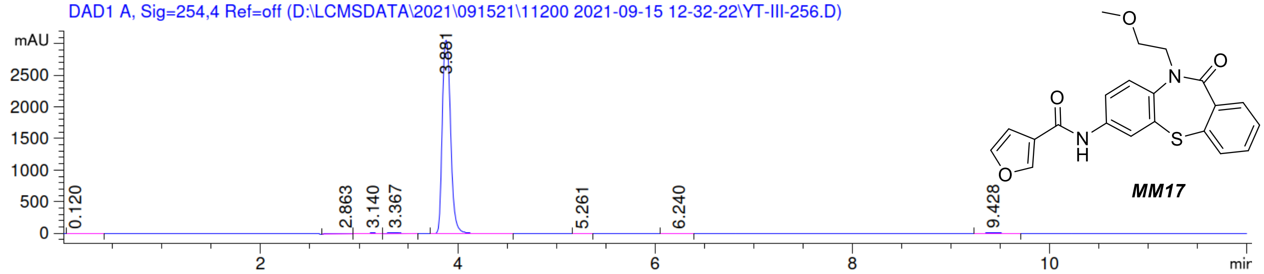


***MM17***

==================================================================

Area Percent Report

==================================================================

Sorted By : Signal
Multiplier: : 1.0000
Dilution: : 1.0000
Use Multiplier & Dilution Factor with ISTDs

Signal 1: DAD1 A, Sig=254,4 Ref=off

| Peak # | RetTime [min] | Type | Width [min] | Area [mAU*s] | Height [mAU] | Area % |
|---|---|---|---|---|---|---|
| 1 | 0.120 | BB | 0.1190 | 51.49717 | 5.95045 | 0.2786 |
| 2 | 2.863 | BV | 0.1641 | 125.99622 | 9.92660 | 0.6816 |
| 3 | 3.140 | VV | 0.2029 | 168.10851 | 11.08994 | 0.9094 |
| 4 | 3.367 | VB | 0.1425 | 154.29399 | 16.90509 | 0.8347 |
| 5 | 3.881 | BB | 0.0943 | 1.78957e4 | 3056.65723 | 96.8086 |
| 6 | 5.261 | BB | 0.0844 | 7.09199 | 1.32688 | 0.0384 |
| 7 | 6.240 | BB | 0.1067 | 11.18461 | 1.57743 | 0.0605 |
| 8 | 9.428 | BB | 0.1226 | 71.77771 | 9.04416 | 0.3883 |
| Totals : | | | | 1.84857e4 | 3112.47779 | |

DAD1 A, Sig=254,4 Ref=off (D:\LCMSDATA\2022\030722\102621 2022-03-07 10-22-30\YT-IV-054.D)

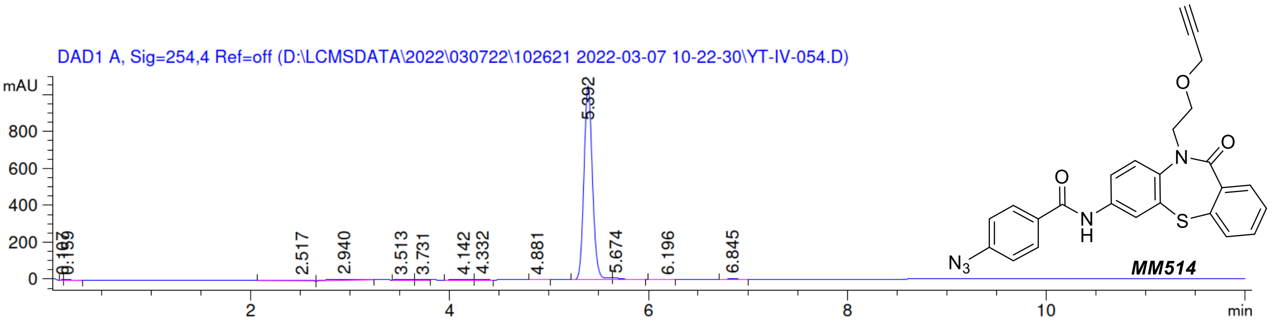


**MM514**

=====================================================================

Area Percent Report

=====================================================================

Sorted By : Signal
Multiplier: : 1.0000
Dilution: : 1.0000
Use Multiplier & Dilution Factor with ISTDs

Signal 1: DAD1 A, Sig=254,4 Ref=off

| Peak # | RetTime [min] | Type | Width [min] | Area [mAU*s] | Height [mAU] | Area % |
|---|---|---|---|---|---|---|
| 1 | 0.107 | BV | 0.0279 | 6.68075 | 3.55866 | 0.1013 |
| 2 | 0.159 | VB | 0.0801 | 22.70214 | 4.13269 | 0.3441 |
| 3 | 2.517 | BB | 0.2653 | 65.14112 | 3.43489 | 0.9873 |
| 4 | 2.940 | BB | 0.2525 | 115.80000 | 5.71145 | 1.7550 |
| 5 | 3.513 | BV | 0.1046 | 13.70055 | 1.88894 | 0.2076 |
| 6 | 3.731 | VV | 0.0864 | 8.42569 | 1.48035 | 0.1277 |
| 7 | 4.142 | BV | 0.1474 | 16.81437 | 1.67087 | 0.2548 |
| 8 | 4.332 | VB | 0.0798 | 16.68235 | 3.14884 | 0.2528 |
| 9 | 4.881 | BB | 0.0832 | 7.49947 | 1.38489 | 0.1137 |
| 10 | 5.392 | VV | 0.0909 | 6210.52637 | 1050.99951 | 94.1249 |
| 11 | 5.674 | VB | 0.1069 | 63.34982 | 8.31243 | 0.9601 |
| 12 | 6.196 | BV | 0.0908 | 9.72139 | 1.55848 | 0.1473 |
| 13 | 6.845 | BB | 0.0927 | 41.12897 | 6.78571 | 0.6233 |
| Totals : | | | | 6598.17300 | 1094.06770 | |

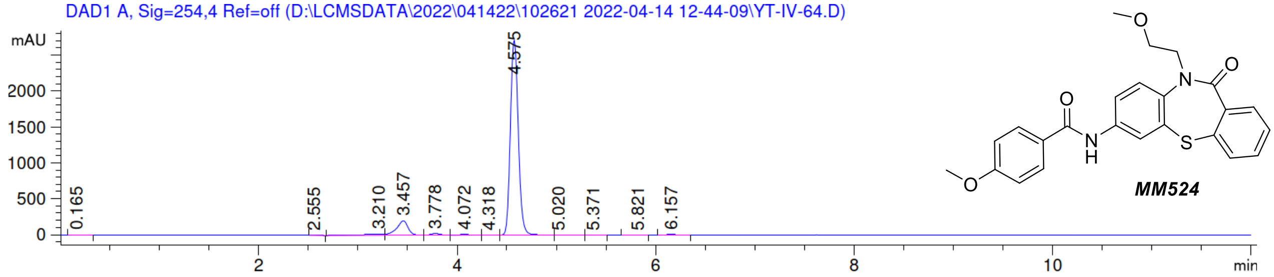


***MM524***

```
=====================================================================
                         Area Percent Report
=====================================================================

Sorted By             :      Signal
Multiplier:                   :      1.0000
Dilution:                     :      1.0000
Use Multiplier & Dilution Factor with ISTDs


Signal 1: DAD1 A, Sig=254,4 Ref=off
```

| Peak # | RetTime [min] | Type | Width [min] | Area [mAU*s] | Height [mAU] | Area % |
|---|---|---|---|---|---|---|
| 1 | 0.130 | BB | 0.1077 | 48.10815 | 6.25833 | 1.7328 |
| 2 | 3.677 | VB | 0.0978 | 22.97034 | 3.44646 | 0.8274 |
| 3 | 3.961 | BB | 0.1063 | 11.43055 | 1.51053 | 0.4117 |
| 4 | 4.463 | BB | 0.0888 | 2669.33838 | 465.92749 | 96.1453 |
| 5 | 5.970 | BB | 0.1150 | 8.83433 | 1.10836 | 0.3182 |
| 6 | 11.512 | BB | 0.1523 | 15.67762 | 1.59986 | 0.5647 |

Totals : 2776.35937 479.85103

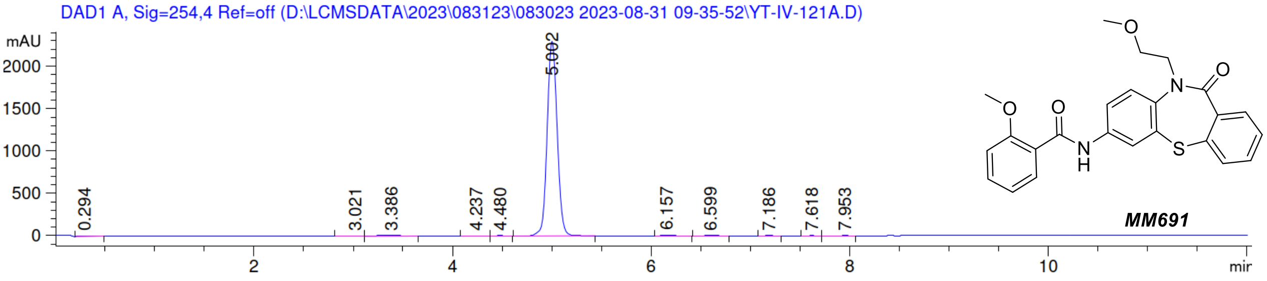


***MM691***

```
=====================================================================
                         Area Percent Report
=====================================================================

Sorted By             :      Signal
Multiplier:                  :      1.0000
Dilution:                    :      1.0000
Use Multiplier & Dilution Factor with ISTDs


Signal 1: DAD1 A, Sig=254,4 Ref=off
```

| Peak # | RetTime [min] | Type | Width [min] | Area [mAU*s] | Height [mAU] | Area % |
|---|---|---|---|---|---|---|
| 1 | 0.294 | BB | 0.1248 | 63.97485 | 6.98596 | 0.3829 |
| 2 | 3.021 | BV | 0.1788 | 80.69483 | 5.71116 | 0.4830 |
| 3 | 3.386 | VB | 0.2404 | 196.90923 | 11.46367 | 1.1785 |
| 4 | 4.237 | BV | 0.1230 | 38.17231 | 4.49836 | 0.2285 |
| 5 | 4.480 | VB | 0.1025 | 32.77188 | 4.99892 | 0.1961 |
| 6 | 5.002 | BV | 0.1099 | 1.61313e4 | 2299.16431 | 96.5462 |
| 7 | 6.157 | VB | 0.1160 | 86.33553 | 11.19411 | 0.5167 |
| 8 | 6.599 | BB | 0.1199 | 41.06409 | 5.10288 | 0.2458 |
| 9 | 7.186 | BB | 0.1018 | 15.79751 | 2.49885 | 0.0945 |
| 10 | 7.618 | BV | 0.0982 | 10.78207 | 1.74137 | 0.0645 |
| 11 | 7.953 | VB | 0.1144 | 10.57028 | 1.36405 | 0.0633 |
| Totals : | | | | 1.67084e4 | 2354.72365 | |

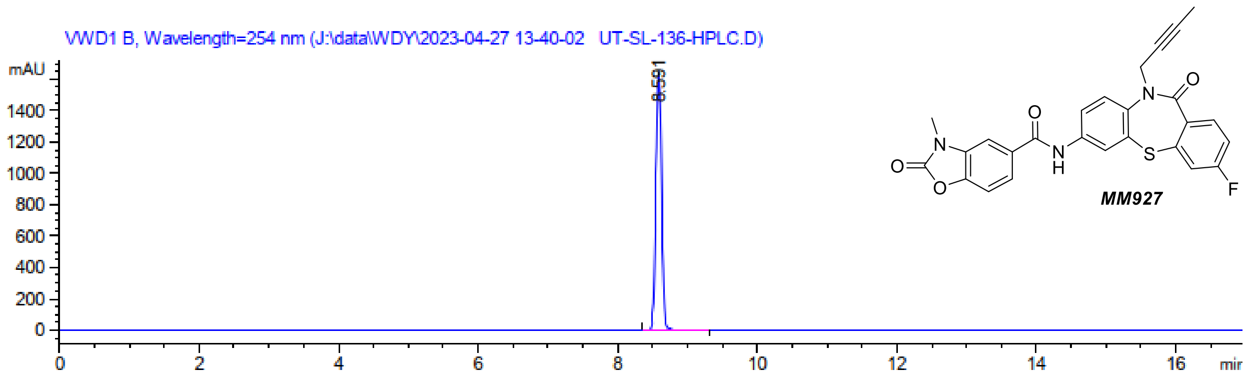


=====================================================================

Area Percent Report

=====================================================================

Sorted By : Signal
Multiplier : 1.0000
Dilution : 1.0000
Sample Amount: : 1.00000 [ng/ul] (not used in calc.)
Use Multiplier & Dilution Factor with ISTDs

Signal 2: VWD1 B, Wavelength=254 nm

| Peak # | RetTime [min] | Type | Width [min] | Area [mAU*s] | Height [mAU] | Area % |
|---|---|---|---|---|---|---|
| 1 | 8.591 | BV R | 0.0890 | 9187.68457 | 1649.84424 | 100.0000 |

Totals : 9187.68457 1649.84424
