## Supplementary material for "A small molecule inhibitor of NVL suppresses tumor growth by blocking ribosome biogenesis": Data S3

### Full wwPDB EM Validation Report (i)

Apr 26, 2024 – 02:50 PM EDT

| | | |
|---|---|---|
| PDB ID | : | 9BJJ |
| EMDB ID | : | EMD-44634 |
| Title | : | Cryo-EM structure of NVL bound the the MM017 inhibitor |
| Deposited on | : | 2024-04-25 |
| Resolution | : | 3.06 Å(reported) |

**This wwPDB validation report is for manuscript review**

This is a Full wwPDB EM Validation Report.

This report is produced by the wwPDB biocuration pipeline after annotation of the structure.

We welcome your comments at **
A user guide is available at
https://www.wwpdb.org/validation/2017/EMValidationReportHelp
with specific help available everywhere you see the (i) symbol.

The types of validation reports are described at
http://www.wwpdb.org/validation/2017/FAQs#types.

The following versions of software and data (see references (i)) were used in the production of this report:

| | | |
|---|---|---|
| EMDB validation analysis | : | 0.0.1.dev92 |
| Mogul | : | 1.8.5 (274361), CSD as541be (2020) |
| MolProbity | : | 4.02b-467 |
| buster-report | : | 1.1.7 (2018) |
| Percentile statistics | : | 20191225.v01 (using entries in the PDB archive December 25th 2019) |
| MapQ | : | 1.9.13 |
| Ideal geometry (proteins) | : | Engh & Huber (2001) |
| Ideal geometry (DNA, RNA) | : | Parkinson et al. (1996) |
| Validation Pipeline (wwPDB-VP) | : | 2.36.2 |

For Manuscript Review

### 1 Overall quality at a glance (i)

The following experimental techniques were used to determine the structure:
*ELECTRON MICROSCOPY*

The reported resolution of this entry is 3.06 Å.

Percentile scores (ranging between 0-100) for global validation metrics of the entry are shown in the following graphic. The table shows the number of entries on which the scores are based.

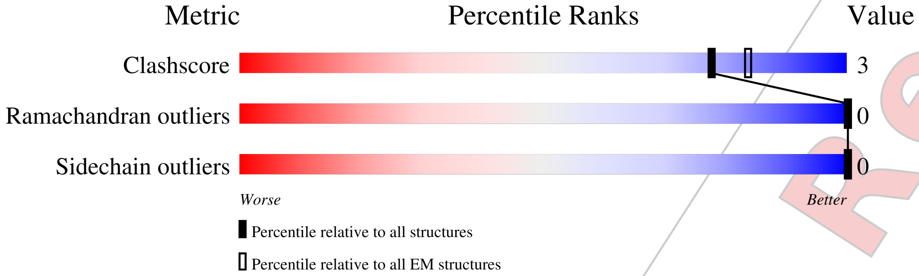


| Metric | Whole archive (#Entries) | EM structures (#Entries) |
|---|---|---|
| Clashscore | 158937 | 4297 |
| Ramachandran outliers | 154571 | 4023 |
| Sidechain outliers | 154315 | 3826 |

The table below summarises the geometric issues observed across the polymeric chains and their fit to the map. The red, orange, yellow and green segments of the bar indicate the fraction of residues that contain outliers for >=3, 2, 1 and 0 types of geometric quality criteria respectively. A grey segment represents the fraction of residues that are not modelled. The numeric value for each fraction is indicated below the corresponding segment, with a dot representing fractions <=5% The upper red bar (where present) indicates the fraction of residues that have poor fit to the EM map (all-atom inclusion < 40%). The numeric value is given above the bar.

| Mol | Chain | Length | Quality of chain |
|---|---|---|---|
| 1 | A | 856 | 10% / 63% • 33% |
| 1 | B | 856 | • / 63% • 33% |
| 1 | C | 856 | • / 62% 5% 33% |
| 1 | D | 856 | 9% / 63% • 34% |
| 1 | E | 856 | 36% / 55% 6% 38% |
| 1 | F | 856 | 46% / 48% • 48% |
| 2 | P | 24 | 46% / 88% 12% |

For Manuscript Review

### 2 Entry composition (i)

There are 5 unique types of molecules in this entry. The entry contains 25639 atoms, of which 0 are hydrogens and 0 are deuteriums.

In the tables below, the AltConf column contains the number of residues with at least one atom in alternate conformation and the Trace column contains the number of residues modelled with at most 2 atoms.

- Molecule 1 is a protein called Nuclear valosin-containing protein-like.

| Mol | Chain | Residues | Atoms | | | | | AltConf | Trace |
|---|---|---|---|---|---|---|---|---|---|
| 1 | A | 572 | Total 4409 | C 2764 | N 791 | O 823 | S 31 | 0 | 0 |
| 1 | B | 571 | Total 4400 | C 2759 | N 790 | O 820 | S 31 | 0 | 0 |
| 1 | C | 571 | Total 4391 | C 2754 | N 787 | O 819 | S 31 | 0 | 0 |
| 1 | D | 569 | Total 4379 | C 2747 | N 784 | O 817 | S 31 | 0 | 0 |
| 1 | E | 527 | Total 4047 | C 2547 | N 722 | O 747 | S 31 | 0 | 0 |
| 1 | F | 444 | Total 3430 | C 2160 | N 615 | O 626 | S 29 | 0 | 0 |

There are 12 discrepancies between the modelled and reference sequences:

| Chain | Residue | Modelled | Actual | Comment | Reference |
|---|---|---|---|---|---|
| A | 365 | GLN | GLU | engineered mutation | UNP O15381 |
| A | 682 | GLN | GLU | engineered mutation | UNP O15381 |
| B | 365 | GLN | GLU | engineered mutation | UNP O15381 |
| B | 682 | GLN | GLU | engineered mutation | UNP O15381 |
| C | 365 | GLN | GLU | engineered mutation | UNP O15381 |
| C | 682 | GLN | GLU | engineered mutation | UNP O15381 |
| D | 365 | GLN | GLU | engineered mutation | UNP O15381 |
| D | 682 | GLN | GLU | engineered mutation | UNP O15381 |
| E | 365 | GLN | GLU | engineered mutation | UNP O15381 |
| E | 682 | GLN | GLU | engineered mutation | UNP O15381 |
| F | 365 | GLN | GLU | engineered mutation | UNP O15381 |
| F | 682 | GLN | GLU | engineered mutation | UNP O15381 |

- Molecule 2 is a protein called Peptide substrate mimic.

For Manuscript Review

| Mol | Chain | Residues | Atoms | | | | AltConf | Trace |
|---|---|---|---|---|---|---|---|---|
| | | | Total | C | N | O | | |
| 2 | P | 24 | 240 | 144 | 72 | 24 | 0 | 0 |

- Molecule 3 is ADENOSINE-5'-TRIPHOSPHATE (three-letter code: ATP) (formula: $C_{10}H_{16}N_5O_{13}P_3$).

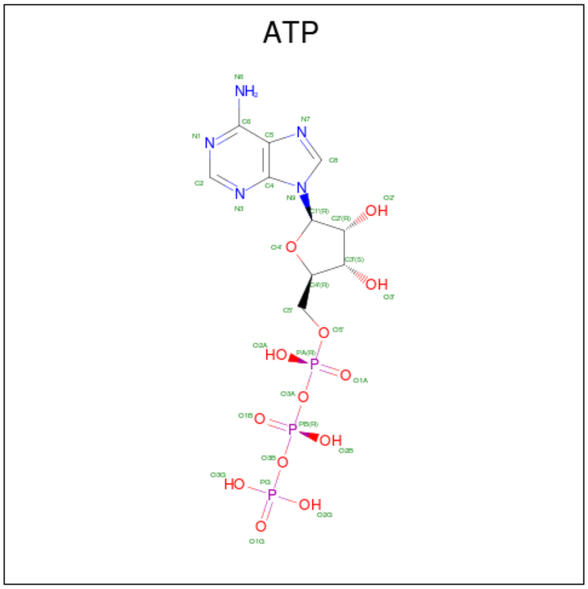


| Mol | Chain | Residues | Atoms | | | | | AltConf |
|---|---|---|---|---|---|---|---|---|
| | | | Total | C | N | O | P | |
| 3 | A | 1 | 31 | 10 | 5 | 13 | 3 | 0 |
| 3 | A | 1 | 31 | 10 | 5 | 13 | 3 | 0 |
| 3 | B | 1 | 31 | 10 | 5 | 13 | 3 | 0 |
| 3 | B | 1 | 31 | 10 | 5 | 13 | 3 | 0 |
| 3 | C | 1 | 31 | 10 | 5 | 13 | 3 | 0 |
| 3 | C | 1 | 31 | 10 | 5 | 13 | 3 | 0 |
| 3 | D | 1 | 31 | 10 | 5 | 13 | 3 | 0 |
| 3 | D | 1 | 31 | 10 | 5 | 13 | 3 | 0 |
| 3 | E | 1 | 31 | 10 | 5 | 13 | 3 | 0 |

- Molecule 4 is MAGNESIUM ION (three-letter code: MG) (formula: Mg).

For Manuscript Review

| Mol | Chain | Residues | Atoms | | AltConf |
|---|---|---|---|---|---|
| | | | Total | Mg | |
| 4 | A | 2 | 2 | 2 | 0 |
| 4 | B | 2 | 2 | 2 | 0 |
| 4 | C | 2 | 2 | 2 | 0 |
| 4 | D | 2 | 2 | 2 | 0 |

- Molecule 5 is N-[10-(2-methoxyethyl)-11-oxo-10,11-dihydrodibenzo[b,f][1,4]thiazepin-7-yl]furan-3-carboxamide (three-letter code: A1AP0) (formula: $C_{21}H_{18}N_2O_4S$) (labeled as "Ligand of Interest" by depositor).

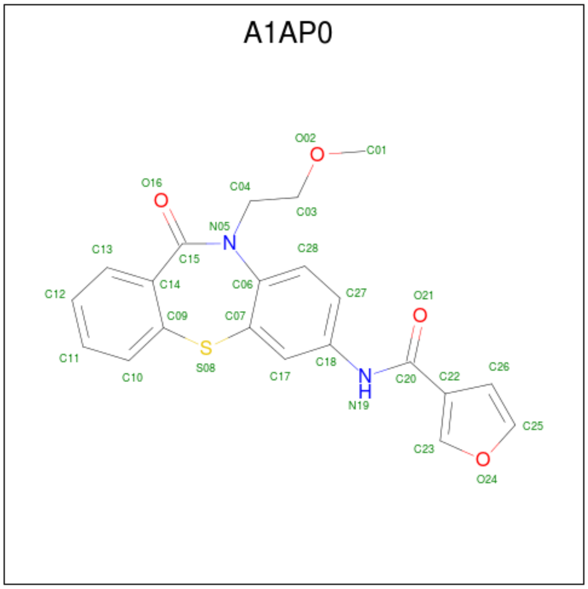


| Mol | Chain | Residues | Atoms | | | | | AltConf |
|---|---|---|---|---|---|---|---|---|
| | | | Total | C | N | O | S | |
| 5 | E | 1 | 28 | 21 | 2 | 4 | 1 | 0 |
| 5 | F | 1 | 28 | 21 | 2 | 4 | 1 | 0 |

For Manuscript Review

### 3 Residue-property plots (i)

These plots are drawn for all protein, RNA, DNA and oligosaccharide chains in the entry. The first graphic for a chain summarises the proportions of the various outlier classes displayed in the second graphic. The second graphic shows the sequence view annotated by issues in geometry and atom inclusion in map density. Residues are color-coded according to the number of geometric quality criteria for which they contain at least one outlier: green = 0, yellow = 1, orange = 2 and red = 3 or more. A red diamond above a residue indicates a poor fit to the EM map for this residue (all-atom inclusion < 40%). Stretches of 2 or more consecutive residues without any outlier are shown as a green connector. Residues present in the sample, but not in the model, are shown in grey.

- Molecule 1: Nuclear valosin-containing protein-like

Chain A:

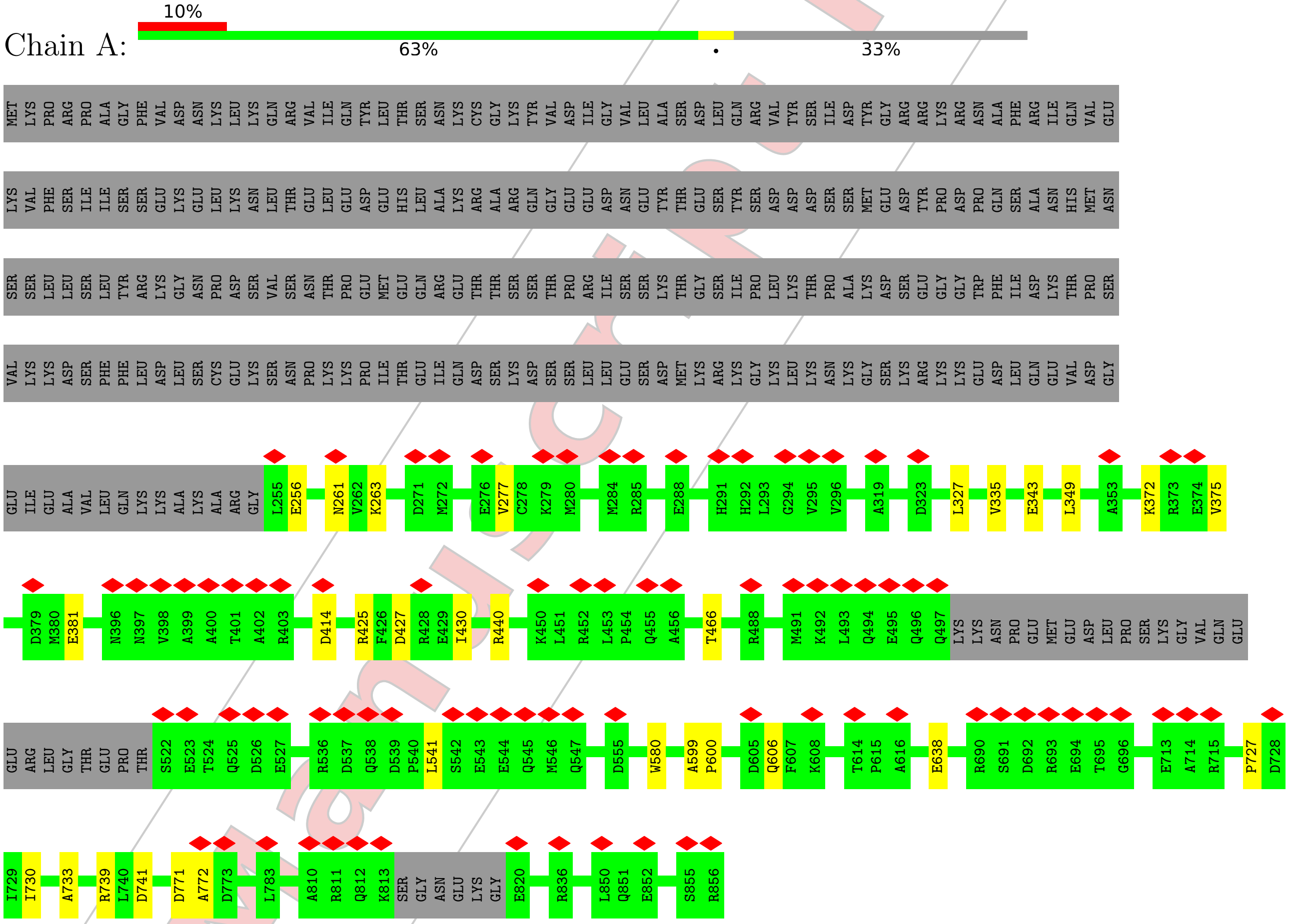


- Molecule 1: Nuclear valosin-containing protein-like

Chain B:

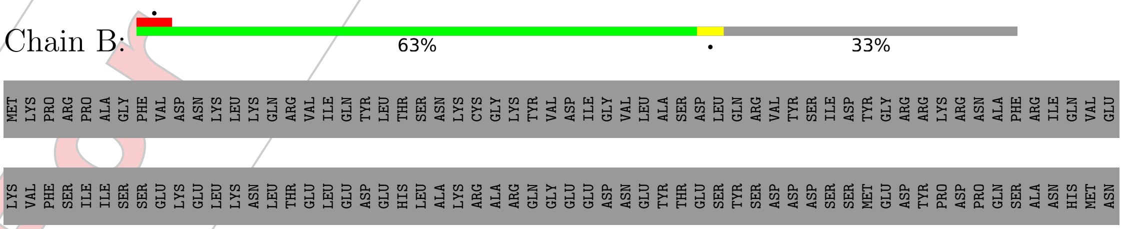


For Manuscript Review

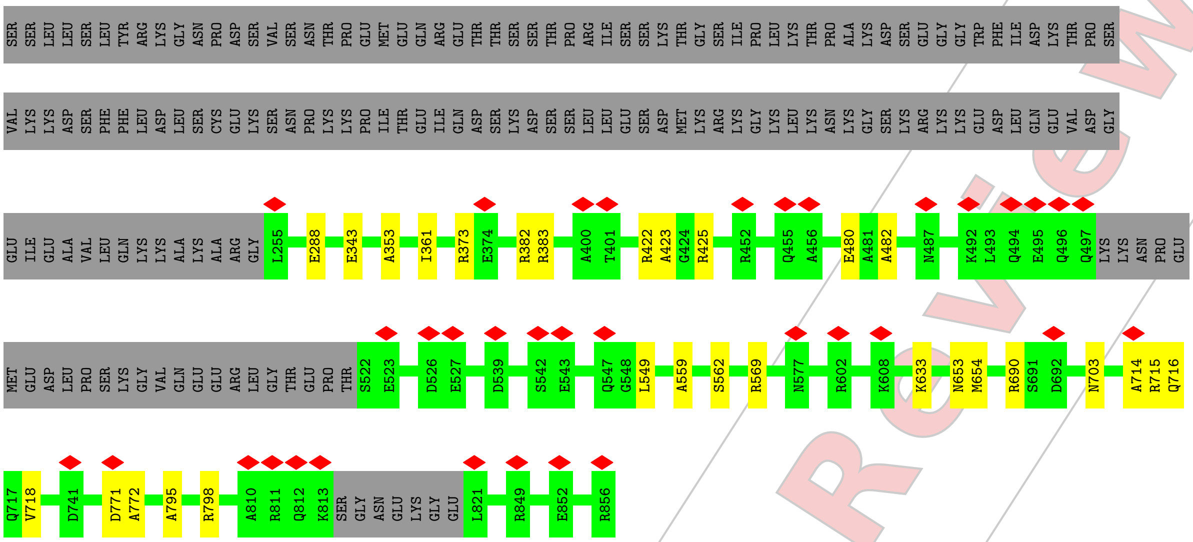


- Molecule 1: Nuclear valosin-containing protein-like

Chain C:

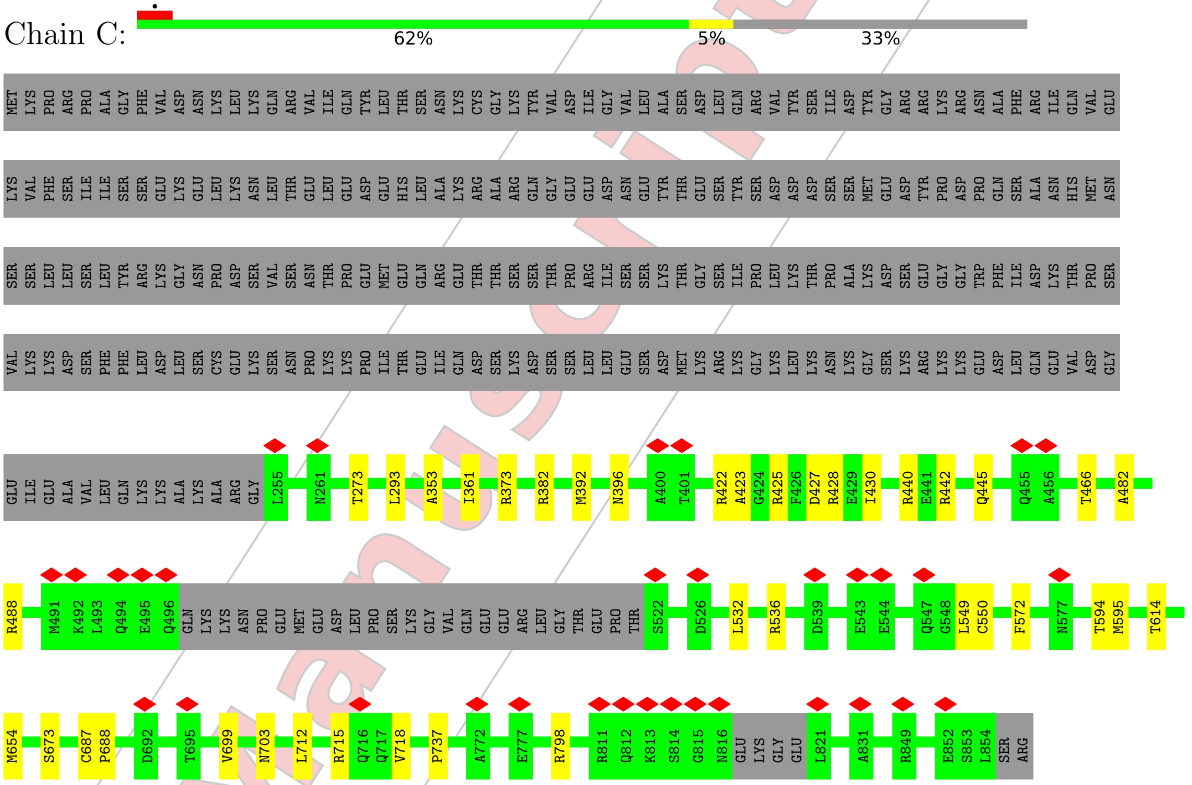


- Molecule 1: Nuclear valosin-containing protein-like

Chain D:

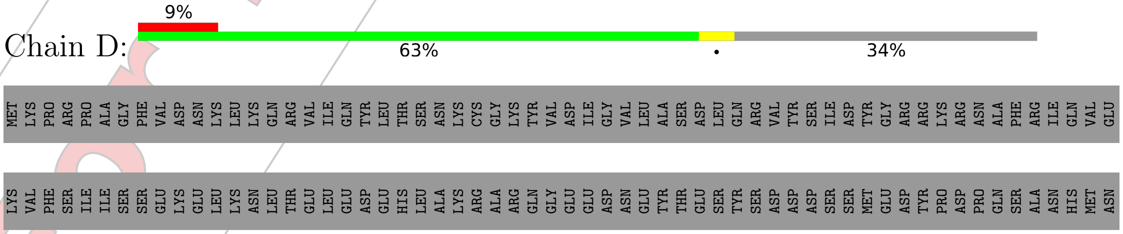


For Manuscript Review

SER SER LEU LEU SER LEU TYR ARG LYS GLY ASN PRO ASP SER VAL SER ASN THR PRO GLU MET GLU GLN ARG GLU THR THR SER SER THR PRO ARG ILE SER SER LYS THR GLY SER ILE PRO LEU LYS THR PRO ALA LYS ASP SER GLU GLY GLY TRP PHE ILE ASP LYS THR PRO SER

VAL LYS LYS ASP SER PHE PHE LEU ASP LEU SER CYS GLU LYS SER ASN PRO LYS LYS PRO ILE THR GLU ILE GLN ASP SER LYS ASP SER SER LEU LEU GLU SER ASP MET LYS ARG LYS GLY LYS LEU LYS ASN LYS GLY SER LYS ARG LYS LYS GLU ASP LEU GLN GLU VAL ASP GLY

GLU ILE GLU ALA VAL LEU GLN LYS LYS ALA LYS ALA ARG GLY L255 I259 S260 N261 L293 A353 I361 K372 V375 M380 I384 A400 T401 A402 R422 A423 R440 Q455 A456 T466 Q494 E495 Q496 Q497 LYS LYS ASN PRO GLU MET GLU ASP

LEU PRO SER LYS GLY VAL GLN GLU GLU ARG LEU GLY THR GLU PRO THR S522 E523 T524 Q525 D526 E527 L528 Q529 R530 L531 L532 G533 L534 L535 R536 D537 Q538 D539 P540 L541 S542 E543 E544 Q545 M546 Q547 G548 L549 E552 D555 S562 V563 Q564 A567 K568 R569 E570 G571 F572 V573

T574 V575 P576 N577 E592 T614 G625 R693 E694 T695 V699 N703 L706 E713 A714 R715 Q716 I734 G738 K742 T743 L744 F745 V746 D753 N764 L770 D771 A772 D773 E777 A780 G781 D782 L783 R784 E799 R806 Q807 E808 M809 A810

R811 Q812 LYS SER GLY ASN GLU LYS GLY GLU L821 K822 V823 K826 A831 F832 K833 K834 V835 R836 K841 K842 R849 S853 L854 S855 ARG

• Molecule 1: Nuclear valosin-containing protein-like

Chain E: 36% | 55% | 6% | 38%

MET LYS PRO ARG PRO ALA GLY PHE VAL ASP ASN LYS LEU LYS GLN ARG VAL ILE GLN TYR LEU THR SER ASN LYS CYS GLY LYS TYR VAL ASP ILE GLY VAL LEU ALA SER ASP LEU GLN ARG VAL TYR SER ILE ASP TYR GLY ARG ARG LYS ARG ASN ALA PHE ARG ILE GLN VAL GLU

LYS VAL PHE SER ILE ILE SER SER GLU LYS GLU LEU LYS ASN LEU THR GLU LEU GLU ASP GLU HIS LEU ALA LYS ARG ALA ARG GLN GLY GLU GLU ASP ASN GLU TYR THR GLU SER TYR SER ASP ASP SER SER MET GLU ASP TYR PRO ASP PRO GLN SER ALA ASN HIS MET ASN

SER SER LEU LEU SER LEU TYR ARG LYS GLY ASN PRO ASP SER VAL SER ASN THR PRO GLU MET GLU GLN ARG GLU THR THR SER SER THR PRO ARG ILE SER SER LYS THR GLY SER ILE PRO LEU LYS THR PRO ALA LYS ASP SER GLU GLY GLY TRP PHE ILE ASP LYS THR PRO SER

VAL LYS LYS ASP SER PHE PHE LEU ASP LEU SER CYS GLU LYS SER ASN PRO LYS LYS PRO ILE THR GLU ILE GLN ASP SER LYS ASP SER SER LEU LEU GLU SER ASP MET LYS ARG LYS GLY LYS LEU LYS ASN LYS GLY SER LYS ARG LYS LYS GLU ASP LEU GLN GLU VAL ASP GLY

GLU ILE GLU ALA VAL LEU GLN LYS LYS ALA LYS ALA ARG GLY LEU GLU F257 Q258 I259 S260 N261 F264 E265 D266 V267 G268 G269 N270 D271 M272 T273 L274 K275 K279 M280 L281 I282 R285 E288 H292 L293 G305 K311 A319 G320 E321 L322 D323 I326 L327 K328

V329 A330 A331 P332 E333 I334 V335 S336 G337 V338 S339 G340 E341 S342 E343 Q344 K345 L346 R347 E348 L349 F350 E351 V354 S355 N356 A357 Q365 I366 D367 A368 I369 T370 P371 LYS ARG GLU VAL ALA SER K378 D379 M380 E381 R382 R383 I384 V385 D393 N397 V398 A399 A400 L405

N411 R412 P413 D414 S415 L416 D417 R421 G433 I434 P435 D436 E437 A438 S439 R440 I443 L444 Q445 T446 R449 K450 L451 R452 L453 P454 Q455 A456 F457 D458 F459 C460 H461 L462 A463 H464 L465 T466 P467 G468 F469 V470 G471 A472 D473 L474 M475 A476 R479 E480 A481 A482 M483 C484 A485

V486 N487 R488 V489 L490 M491 K492 L493 Q494 E495 Q496 Q497 LYS LYS ASN PRO GLU MET GLU ASP LEU PRO SER LYS GLY VAL GLN GLU GLU ARG LEU GLY THR GLU PRO THR SER GLU THR GLN ASP GLU LEU GLN ARG LEU L532 G533 L534 L535 R536 D537 Q538 ASP PRO L541 S542 E543 E544 Q545

For Manuscript Review

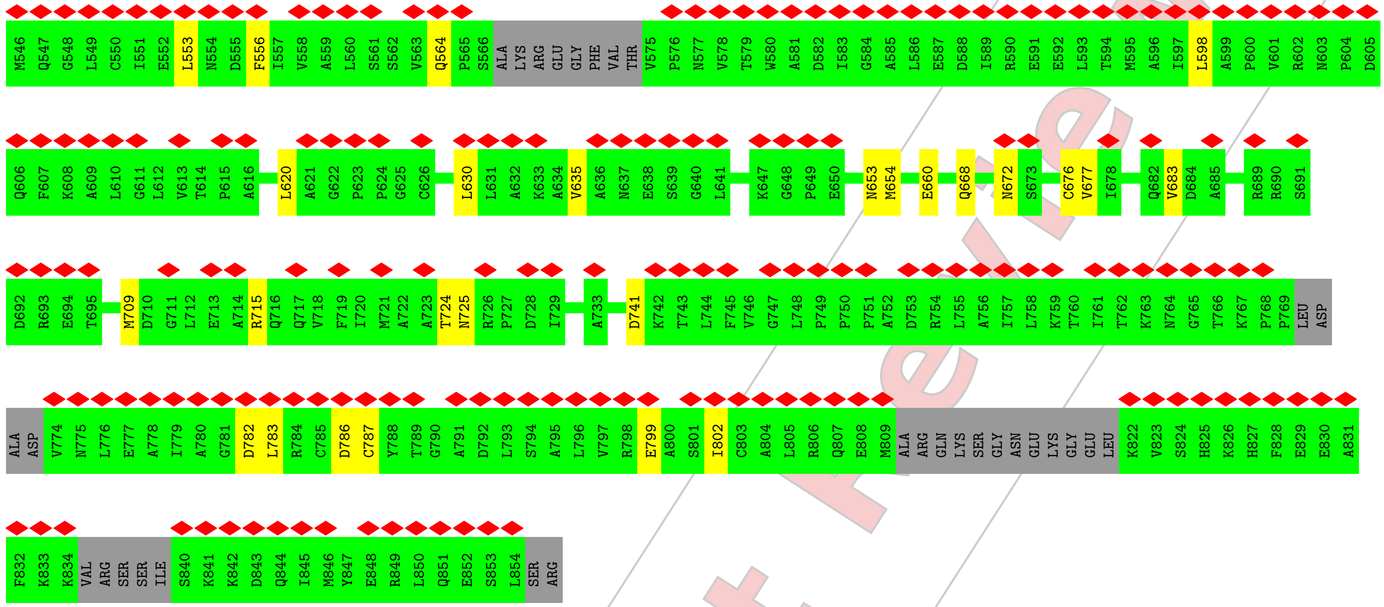


• Molecule 1: Nuclear valosin-containing protein-like

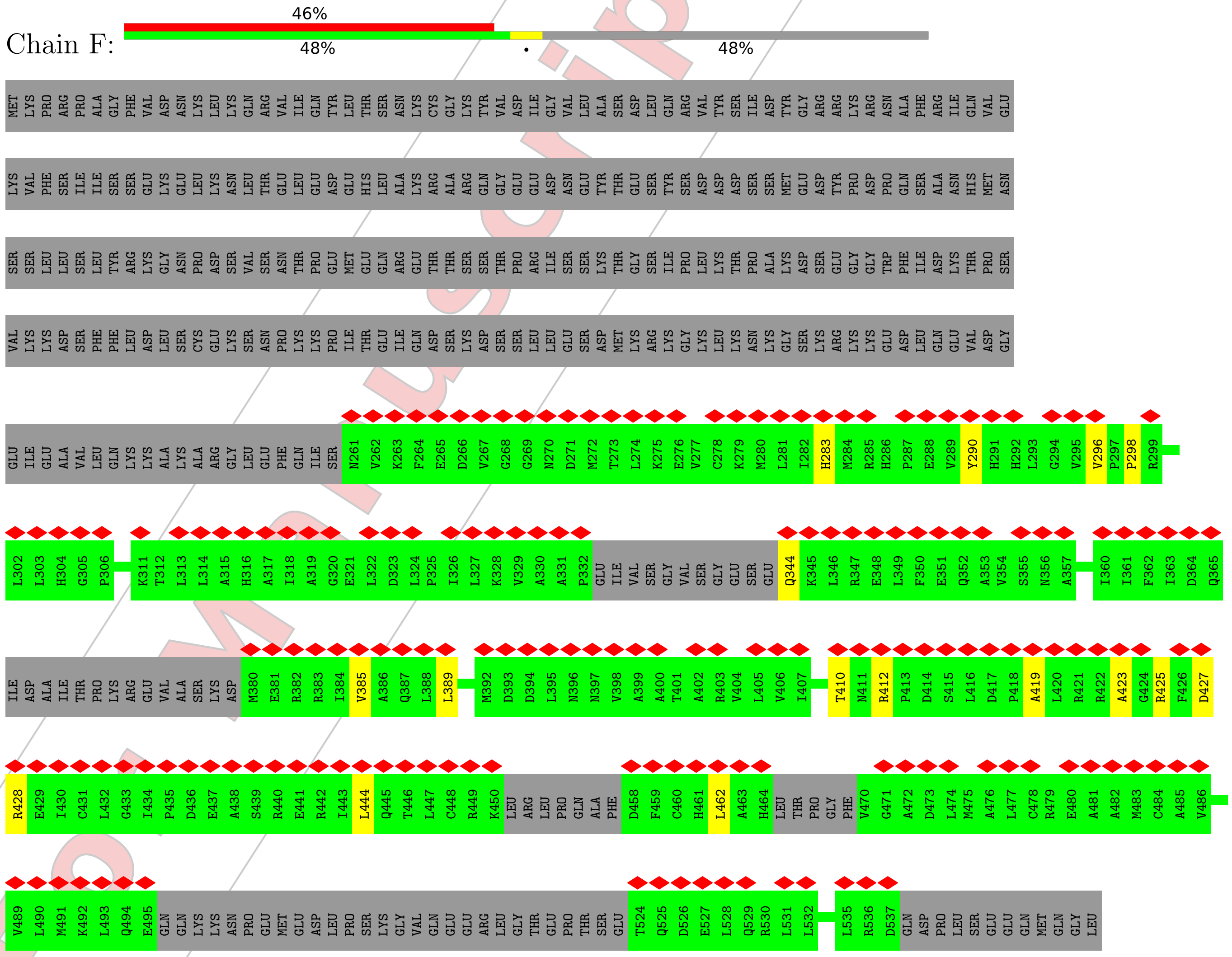


For Manuscript Review

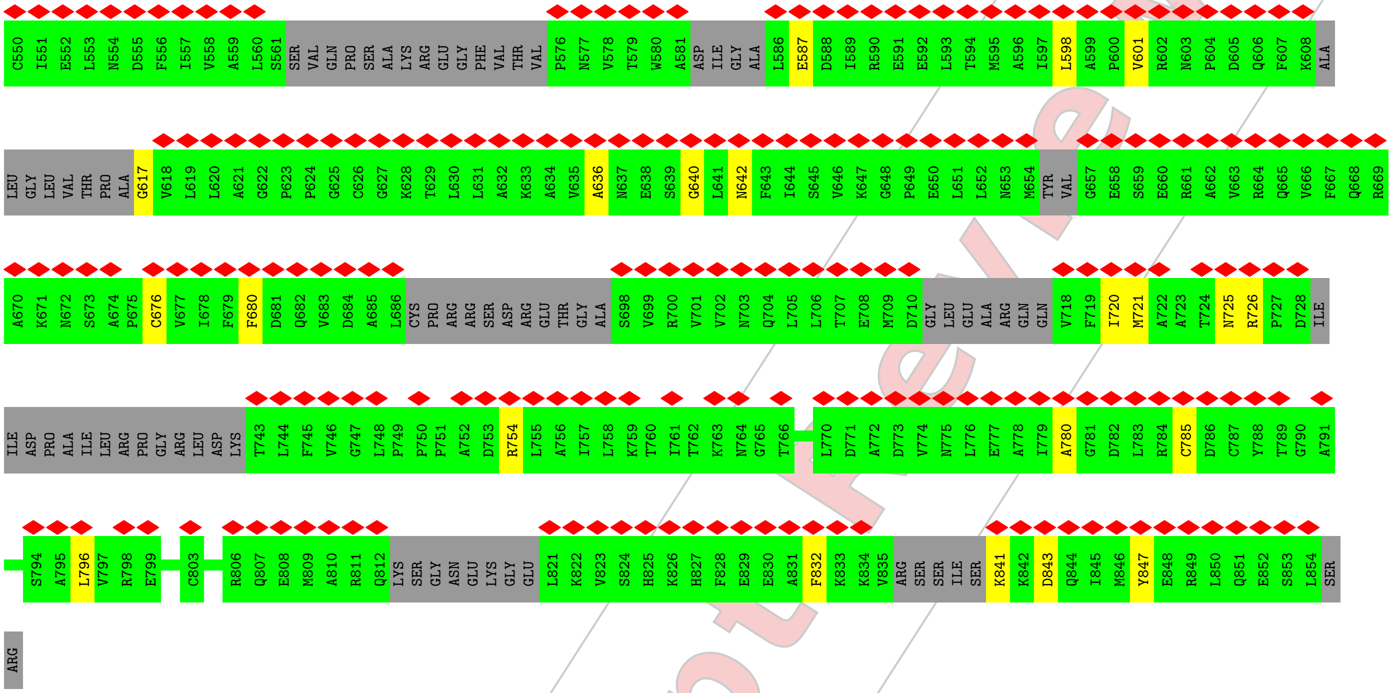


• Molecule 2: Peptide substrate mimic

Chain P:

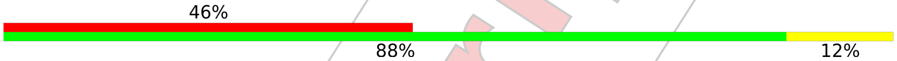


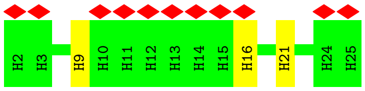


For Manuscript Review

### 4 Experimental information ⓘ

| Property | Value | Source |
|---|---|---|
| EM reconstruction method | SINGLE PARTICLE | Depositor |
| Imposed symmetry | POINT, Not provided | |
| Number of particles used | 100118 | Depositor |
| Resolution determination method | FSC 0.143 CUT-OFF | Depositor |
| CTF correction method | PHASE FLIPPING AND AMPLITUDE CORRECTION | Depositor |
| Microscope | TFS KRIOS | Depositor |
| Voltage (kV) | 300 | Depositor |
| Electron dose $(e^-/\text{Å}^2)$ | 1.4 | Depositor |
| Minimum defocus (nm) | 600 | Depositor |
| Maximum defocus (nm) | 1900 | Depositor |
| Magnification | Not provided | |
| Image detector | GATAN K3 (6k x 4k) | Depositor |
| Maximum map value | 0.120 | Depositor |
| Minimum map value | -0.057 | Depositor |
| Average map value | 0.000 | Depositor |
| Map value standard deviation | 0.004 | Depositor |
| Recommended contour level | 0.012 | Depositor |
| Map size (Å) | 232.4, 232.4, 232.4 | wwPDB |
| Map dimensions | 280, 280, 280 | wwPDB |
| Map angles (°) | 90.0, 90.0, 90.0 | wwPDB |
| Pixel spacing (Å) | 0.83, 0.83, 0.83 | Depositor |

For Manuscript Review

### 5 Model quality (i)

#### 5.1 Standard geometry (i)

Bond lengths and bond angles in the following residue types are not validated in this section: A1AP0, ATP, MG

The Z score for a bond length (or angle) is the number of standard deviations the observed value is removed from the expected value. A bond length (or angle) with $|Z| > 5$ is considered an outlier worth inspection. RMSZ is the root-mean-square of all Z scores of the bond lengths (or angles).

| Mol | Chain | Bond lengths | | Bond angles | |
|---|---|---|---|---|---|
| | | RMSZ | #\|Z\| >5 | RMSZ | #\|Z\| >5 |
| 1 | A | 0.24 | 0/4474 | 0.48 | 0/6048 |
| 1 | B | 0.24 | 0/4465 | 0.49 | 0/6036 |
| 1 | C | 0.24 | 0/4456 | 0.48 | 0/6026 |
| 1 | D | 0.24 | 0/4444 | 0.48 | 0/6011 |
| 1 | E | 0.24 | 0/4105 | 0.48 | 0/5547 |
| 1 | F | 0.23 | 0/3468 | 0.46 | 0/4668 |
| 2 | P | 0.18 | 0/263 | 0.32 | 0/357 |
| All | All | 0.24 | 0/25675 | 0.48 | 0/34693 |

There are no bond length outliers.

There are no bond angle outliers.

There are no chirality outliers.

There are no planarity outliers.

#### 5.2 Too-close contacts (i)

In the following table, the Non-H and H(model) columns list the number of non-hydrogen atoms and hydrogen atoms in the chain respectively. The H(added) column lists the number of hydrogen atoms added and optimized by MolProbity. The Clashes column lists the number of clashes within the asymmetric unit, whereas Symm-Clashes lists symmetry-related clashes.

| Mol | Chain | Non-H | H(model) | H(added) | Clashes | Symm-Clashes |
|---|---|---|---|---|---|---|
| 1 | A | 4409 | 0 | 4515 | 19 | 0 |
| 1 | B | 4400 | 0 | 4509 | 20 | 0 |
| 1 | C | 4391 | 0 | 4497 | 28 | 0 |
| 1 | D | 4379 | 0 | 4484 | 24 | 0 |
| 1 | E | 4047 | 0 | 4148 | 52 | 0 |
| 1 | F | 3430 | 0 | 3518 | 25 | 0 |

*Continued on next page...*

For Manuscript Review

For Manuscript Review

*Continued from previous page...*

| Mol | Chain | Non-H | H(model) | H(added) | Clashes | Symm-Clashes |
|---|---|---|---|---|---|---|
| 2 | P | 240 | 0 | 167 | 3 | 0 |
| 3 | A | 62 | 0 | 24 | 1 | 0 |
| 3 | B | 62 | 0 | 24 | 2 | 0 |
| 3 | C | 62 | 0 | 24 | 0 | 0 |
| 3 | D | 62 | 0 | 24 | 1 | 0 |
| 3 | E | 31 | 0 | 12 | 0 | 0 |
| 4 | A | 2 | 0 | 0 | 0 | 0 |
| 4 | B | 2 | 0 | 0 | 0 | 0 |
| 4 | C | 2 | 0 | 0 | 0 | 0 |
| 4 | D | 2 | 0 | 0 | 0 | 0 |
| 5 | E | 28 | 0 | 0 | 0 | 0 |
| 5 | F | 28 | 0 | 0 | 1 | 0 |
| All | All | 25639 | 0 | 25946 | 150 | 0 |

The all-atom clashscore is defined as the number of clashes found per 1000 atoms (including hydrogen atoms). The all-atom clashscore for this structure is 3.

All (150) close contacts within the same asymmetric unit are listed below, sorted by their clash magnitude.

| Atom-1 | Atom-2 | Interatomic distance (Å) | Clash overlap (Å) |
|---|---|---|---|
| 1:E:481:ALA:HB1 | 1:E:556:PHE:CD1 | 1.58 | 1.37 |
| 1:E:481:ALA:HB1 | 1:E:556:PHE:CE1 | 1.64 | 1.31 |
| 1:E:481:ALA:CB | 1:E:556:PHE:CD1 | 2.24 | 1.20 |
| 1:E:481:ALA:CB | 1:E:556:PHE:CE1 | 2.24 | 1.19 |
| 1:E:481:ALA:CB | 1:E:556:PHE:HE1 | 1.78 | 0.94 |
| 1:E:481:ALA:CB | 1:E:556:PHE:HD1 | 1.77 | 0.86 |
| 1:E:457:PHE:HE2 | 1:E:556:PHE:HE2 | 1.25 | 0.85 |
| 1:F:642:ASN:HB2 | 1:F:676:CYS:SG | 2.22 | 0.80 |
| 1:E:481:ALA:HB1 | 1:E:556:PHE:HD1 | 1.27 | 0.80 |
| 1:E:457:PHE:CE2 | 1:E:556:PHE:HE2 | 2.02 | 0.76 |
| 1:E:380:MET:SD | 1:E:383:ARG:NH1 | 2.58 | 0.76 |
| 1:E:481:ALA:HB3 | 1:E:556:PHE:HE1 | 1.52 | 0.74 |
| 1:D:380:MET:SD | 2:P:9:HIS:ND1 | 2.61 | 0.73 |
| 1:E:481:ALA:HB3 | 1:E:556:PHE:CE1 | 2.19 | 0.72 |
| 3:A:901:ATP:O1G | 1:B:425:ARG:NH1 | 2.23 | 0.72 |
| 1:B:633:LYS:NZ | 1:C:712:LEU:O | 2.24 | 0.70 |
| 1:C:422:ARG:NH1 | 1:C:423:ALA:O | 2.23 | 0.70 |
| 1:E:481:ALA:HB2 | 1:E:556:PHE:CD1 | 2.21 | 0.70 |
| 1:E:564:GLN:N | 1:E:564:GLN:OE1 | 2.25 | 0.69 |
| 1:A:541:LEU:N | 1:B:288:GLU:OE2 | 2.28 | 0.66 |
| 1:F:617:GLY:N | 1:F:720:ILE:O | 2.29 | 0.66 |

*Continued on next page...*

*Continued from previous page...*

| Atom-1 | Atom-2 | Interatomic distance (Å) | Clash overlap (Å) |
|---|---|---|---|
| 1:D:422:ARG:NH1 | 1:D:423:ALA:O | 2.30 | 0.64 |
| 1:A:771:ASP:OD1 | 1:A:772:ALA:N | 2.31 | 0.63 |
| 1:F:754:ARG:NH1 | 1:F:780:ALA:O | 2.31 | 0.63 |
| 1:B:343:GLU:N | 1:B:343:GLU:OE1 | 2.32 | 0.61 |
| 1:E:487:ASN:OD1 | 1:F:283:HIS:NE2 | 2.33 | 0.61 |
| 1:C:488:ARG:NH2 | 1:C:550:CYS:O | 2.34 | 0.60 |
| 1:E:343:GLU:OE2 | 1:E:383:ARG:NH1 | 2.34 | 0.60 |
| 1:E:457:PHE:CE2 | 1:E:556:PHE:CE2 | 2.88 | 0.60 |
| 1:A:343:GLU:OE1 | 1:A:343:GLU:N | 2.34 | 0.59 |
| 1:D:715:ARG:NH2 | 1:D:738:GLY:O | 2.35 | 0.59 |
| 1:B:422:ARG:NH1 | 1:B:423:ALA:O | 2.35 | 0.59 |
| 1:A:372:LYS:NZ | 1:A:414:ASP:O | 2.36 | 0.58 |
| 1:C:392:MET:O | 1:C:396:ASN:ND2 | 2.35 | 0.58 |
| 1:E:380:MET:HA | 1:E:383:ARG:CZ | 2.33 | 0.58 |
| 1:D:592:GLU:OE2 | 1:D:742:LYS:NZ | 2.29 | 0.58 |
| 3:B:901:ATP:O2G | 1:C:425:ARG:NH1 | 2.37 | 0.57 |
| 1:B:373:ARG:O | 1:B:382:ARG:NH1 | 2.38 | 0.57 |
| 1:E:397:ASN:OD1 | 1:E:398:VAL:N | 2.37 | 0.57 |
| 1:C:373:ARG:O | 1:C:382:ARG:NH1 | 2.36 | 0.56 |
| 1:E:653:ASN:OD1 | 1:E:654:MET:N | 2.39 | 0.56 |
| 1:E:311:LYS:HE3 | 1:E:411:ASN:OD1 | 2.06 | 0.55 |
| 1:A:727:PRO:O | 1:A:730:ILE:HG22 | 2.08 | 0.54 |
| 1:D:440:ARG:NE | 1:D:466:THR:O | 2.41 | 0.54 |
| 1:A:580:TRP:NE1 | 1:A:638:GLU:OE2 | 2.37 | 0.54 |
| 1:C:549:LEU:HG | 1:D:293:LEU:HD21 | 1.91 | 0.53 |
| 1:D:552:GLU:N | 1:D:555:ASP:OD2 | 2.40 | 0.53 |
| 1:E:709:MET:O | 1:E:715:ARG:NH1 | 2.41 | 0.53 |
| 1:D:546:MET:O | 1:E:292:HIS:NE2 | 2.42 | 0.52 |
| 1:E:311:LYS:CE | 1:E:411:ASN:OD1 | 2.57 | 0.52 |
| 1:E:455:GLN:OE1 | 1:E:455:GLN:N | 2.41 | 0.51 |
| 1:E:481:ALA:HB2 | 1:E:556:PHE:HD1 | 1.63 | 0.51 |
| 1:F:598:LEU:O | 1:F:601:VAL:HG12 | 2.10 | 0.51 |
| 1:A:375:VAL:HG12 | 1:A:375:VAL:O | 2.10 | 0.50 |
| 1:B:353:ALA:CB | 1:B:361:ILE:HD11 | 2.42 | 0.50 |
| 1:C:715:ARG:NH1 | 1:C:718:VAL:O | 2.44 | 0.50 |
| 1:F:419:ALA:O | 1:F:425:ARG:NH1 | 2.40 | 0.50 |
| 1:E:553:LEU:HA | 1:E:556:PHE:HB2 | 1.92 | 0.50 |
| 1:F:298:PRO:O | 5:F:1001:A1AP0:N19 | 2.44 | 0.50 |
| 1:F:841:LYS:N | 1:F:843:ASP:OD1 | 2.45 | 0.50 |
| 1:B:798:ARG:NH2 | 1:C:614:THR:O | 2.42 | 0.49 |
| 1:C:798:ARG:NH2 | 1:D:614:THR:O | 2.44 | 0.49 |

*Continued on next page...*

For Manuscript Review

*Continued from previous page...*

| Atom-1 | Atom-2 | Interatomic distance (Å) | Clash overlap (Å) |
|---|---|---|---|
| 1:E:380:MET:HA | 1:E:383:ARG:NH2 | 2.28 | 0.49 |
| 1:E:367:ASP:OD2 | 1:E:412:ARG:NH1 | 2.46 | 0.48 |
| 1:A:261:ASN:O | 1:A:263:LYS:NZ | 2.43 | 0.48 |
| 1:F:385:VAL:O | 1:F:389:LEU:HD13 | 2.12 | 0.48 |
| 1:F:754:ARG:NH2 | 1:F:785:CYS:O | 2.46 | 0.48 |
| 1:A:381:GLU:OE1 | 1:A:381:GLU:N | 2.43 | 0.48 |
| 1:A:606:GLN:OE1 | 1:A:606:GLN:N | 2.44 | 0.48 |
| 1:D:746:VAL:HG13 | 1:D:746:VAL:O | 2.13 | 0.48 |
| 1:D:782:ASP:OD1 | 1:D:783:LEU:N | 2.47 | 0.48 |
| 1:E:470:VAL:HG12 | 1:E:471:GLY:N | 2.29 | 0.48 |
| 1:F:427:ASP:OD1 | 1:F:428:ARG:N | 2.46 | 0.48 |
| 1:F:587:GLU:OE1 | 1:F:587:GLU:N | 2.43 | 0.47 |
| 1:B:562:SER:O | 1:C:428:ARG:NH2 | 2.42 | 0.47 |
| 1:F:636:ALA:O | 1:F:640:GLY:N | 2.47 | 0.47 |
| 1:F:642:ASN:CB | 1:F:676:CYS:SG | 3.00 | 0.47 |
| 1:B:653:ASN:OD1 | 1:B:654:MET:N | 2.47 | 0.47 |
| 1:D:782:ASP:OD2 | 1:D:784:ARG:NH2 | 2.48 | 0.47 |
| 1:D:771:ASP:OD1 | 1:D:772:ALA:N | 2.44 | 0.47 |
| 1:A:425:ARG:O | 1:A:427:ASP:N | 2.47 | 0.46 |
| 1:E:668:GLN:O | 1:E:672:ASN:ND2 | 2.45 | 0.46 |
| 1:D:531:LEU:O | 1:E:282:ILE:HD11 | 2.15 | 0.46 |
| 1:F:726:ARG:NH1 | 1:F:847:TYR:OH | 2.49 | 0.46 |
| 1:E:786:ASP:OD1 | 1:E:787:CYS:N | 2.48 | 0.46 |
| 1:E:676:CYS:SG | 1:E:677:VAL:N | 2.88 | 0.46 |
| 1:A:277:VAL:HG22 | 1:A:430:ILE:HD13 | 1.96 | 0.46 |
| 1:C:353:ALA:CB | 1:C:361:ILE:HD11 | 2.46 | 0.46 |
| 1:E:472:ALA:HB1 | 1:F:423:ALA:HB3 | 1.99 | 0.46 |
| 1:F:725:ASN:OD1 | 1:F:726:ARG:N | 2.49 | 0.46 |
| 3:B:901:ATP:PG | 1:C:425:ARG:HH12 | 2.39 | 0.45 |
| 1:C:442:ARG:NH1 | 1:C:445:GLN:OE1 | 2.49 | 0.45 |
| 1:A:256:GLU:N | 1:A:256:GLU:OE1 | 2.49 | 0.45 |
| 1:A:739:ARG:O | 1:A:741:ASP:N | 2.49 | 0.45 |
| 1:B:690:ARG:NH1 | 1:B:703:ASN:OD1 | 2.50 | 0.45 |
| 1:B:549:LEU:HG | 1:C:293:LEU:HD21 | 2.00 | 0.44 |
| 1:B:714:ALA:O | 1:B:716:GLN:NE2 | 2.50 | 0.44 |
| 1:C:482:ALA:HB1 | 1:D:293:LEU:HD13 | 1.99 | 0.44 |
| 1:D:625:GLY:HA2 | 3:D:902:ATP:H5'1 | 1.99 | 0.44 |
| 1:D:380:MET:O | 1:D:384:ILE:HG12 | 2.18 | 0.44 |
| 2:P:16:HIS:O | 2:P:16:HIS:ND1 | 2.50 | 0.44 |
| 1:B:771:ASP:OD1 | 1:B:772:ALA:N | 2.49 | 0.44 |
| 1:D:699:VAL:O | 1:D:703:ASN:ND2 | 2.47 | 0.44 |

*Continued on next page...*

For Manuscript Review

Continued from previous page...

| Atom-1 | Atom-2 | Interatomic distance (Å) | Clash overlap (Å) |
|---|---|---|---|
| 1:A:599:ALA:HB3 | 1:A:600:PRO:HD3 | 1.99 | 0.44 |
| 1:E:782:ASP:OD1 | 1:E:783:LEU:N | 2.51 | 0.44 |
| 1:F:680:PHE:N | 1:F:721:MET:O | 2.48 | 0.44 |
| 1:D:353:ALA:CB | 1:D:361:ILE:HD11 | 2.47 | 0.44 |
| 1:E:660:GLU:OE1 | 1:E:660:GLU:N | 2.39 | 0.44 |
| 1:E:535:LEU:HD22 | 1:E:535:LEU:H | 1.83 | 0.43 |
| 1:B:480:GLU:HB3 | 1:B:559:ALA:HB1 | 2.00 | 0.43 |
| 1:C:687:CYS:N | 1:C:688:PRO:HD3 | 2.33 | 0.43 |
| 1:C:798:ARG:NH2 | 1:D:614:THR:OG1 | 2.51 | 0.43 |
| 1:D:592:GLU:HG2 | 1:D:744:LEU:HD21 | 2.01 | 0.43 |
| 1:E:443:ILE:HG21 | 1:E:474:LEU:HD12 | 2.00 | 0.43 |
| 1:E:683:VAL:HG22 | 1:E:683:VAL:O | 2.18 | 0.43 |
| 1:E:329:VAL:HG21 | 1:E:349:LEU:HD11 | 2.00 | 0.43 |
| 1:E:724:THR:OG1 | 1:E:725:ASN:N | 2.50 | 0.42 |
| 1:C:440:ARG:NE | 1:C:466:THR:O | 2.52 | 0.42 |
| 1:C:594:THR:HG23 | 1:C:595:MET:N | 2.33 | 0.42 |
| 1:C:654:MET:O | 2:P:21:HIS:N | 2.47 | 0.42 |
| 1:E:486:VAL:HG21 | 1:F:290:TYR:CE1 | 2.53 | 0.42 |
| 1:F:344:GLN:N | 1:F:344:GLN:OE1 | 2.52 | 0.42 |
| 1:B:482:ALA:HB1 | 1:C:293:LEU:HD13 | 2.01 | 0.42 |
| 1:A:440:ARG:NE | 1:A:466:THR:O | 2.53 | 0.42 |
| 1:B:795:ALA:HB2 | 1:C:737:PRO:HB3 | 2.02 | 0.42 |
| 1:F:296:VAL:HG23 | 1:F:296:VAL:O | 2.20 | 0.42 |
| 1:C:699:VAL:O | 1:C:703:ASN:ND2 | 2.46 | 0.42 |
| 1:A:335:VAL:HG21 | 1:B:383:ARG:CD | 2.50 | 0.42 |
| 1:E:741:ASP:OD1 | 1:E:741:ASP:N | 2.52 | 0.42 |
| 1:E:799:GLU:HA | 1:E:802:ILE:HG22 | 2.02 | 0.42 |
| 1:D:549:LEU:HG | 1:E:293:LEU:HD21 | 2.02 | 0.41 |
| 1:D:706:LEU:HD23 | 1:D:734:ILE:HD13 | 2.02 | 0.41 |
| 1:E:630:LEU:HD12 | 1:E:630:LEU:N | 2.35 | 0.41 |
| 1:C:532:LEU:O | 1:C:536:ARG:HG2 | 2.20 | 0.41 |
| 1:F:425:ARG:O | 1:F:427:ASP:N | 2.49 | 0.41 |
| 1:C:273:THR:HG23 | 1:C:430:ILE:HG21 | 2.02 | 0.41 |
| 1:D:372:LYS:HB2 | 1:D:375:VAL:HG22 | 2.03 | 0.41 |
| 1:F:410:THR:HG22 | 1:F:412:ARG:H | 1.86 | 0.41 |
| 1:F:444:LEU:HD21 | 1:F:462:LEU:HD23 | 2.01 | 0.41 |
| 1:A:733:ALA:O | 1:A:739:ARG:NH1 | 2.46 | 0.41 |
| 1:E:413:PRO:O | 1:E:421:ARG:NH2 | 2.54 | 0.41 |
| 1:A:327:LEU:HD21 | 1:A:349:LEU:HD12 | 2.03 | 0.41 |
| 1:E:457:PHE:HZ | 1:E:556:PHE:HD2 | 1.68 | 0.41 |
| 1:E:311:LYS:NZ | 1:E:411:ASN:OD1 | 2.53 | 0.40 |

Continued on next page...

For Manuscript Review

*Continued from previous page...*

| Atom-1 | Atom-2 | Interatomic distance (Å) | Clash overlap (Å) |
|---|---|---|---|
| 1:F:796:LEU:HD11 | 1:F:832:PHE:HD1 | 1.86 | 0.40 |
| 1:B:715:ARG:NH1 | 1:B:718:VAL:O | 2.52 | 0.40 |
| 1:C:572:PHE:HZ | 1:C:673:SER:HG | 1.66 | 0.40 |
| 1:E:598:LEU:HD12 | 1:E:635:VAL:HG13 | 2.03 | 0.40 |
| 1:E:620:LEU:HD12 | 1:E:620:LEU:N | 2.36 | 0.40 |
| 1:B:569:ARG:NH2 | 1:C:427:ASP:O | 2.54 | 0.40 |

There are no symmetry-related clashes.

#### 5.3 Torsion angles (i)

##### 5.3.1 Protein backbone (i)

In the following table, the Percentiles column shows the percent Ramachandran outliers of the chain as a percentile score with respect to all PDB entries followed by that with respect to all EM entries.

The Analysed column shows the number of residues for which the backbone conformation was analysed, and the total number of residues.

| Mol | Chain | Analysed | Favoured | Allowed | Outliers | Percentiles | |
|---|---|---|---|---|---|---|---|
| 1 | A | 566/856 (66%) | 555 (98%) | 11 (2%) | 0 | 100 | 100 |
| 1 | B | 565/856 (66%) | 561 (99%) | 4 (1%) | 0 | 100 | 100 |
| 1 | C | 565/856 (66%) | 560 (99%) | 5 (1%) | 0 | 100 | 100 |
| 1 | D | 563/856 (66%) | 554 (98%) | 9 (2%) | 0 | 100 | 100 |
| 1 | E | 511/856 (60%) | 501 (98%) | 10 (2%) | 0 | 100 | 100 |
| 1 | F | 412/856 (48%) | 403 (98%) | 9 (2%) | 0 | 100 | 100 |
| 2 | P | 22/24 (92%) | 21 (96%) | 1 (4%) | 0 | 100 | 100 |
| All | All | 3204/5160 (62%) | 3155 (98%) | 49 (2%) | 0 | 100 | 100 |

There are no Ramachandran outliers to report.

##### 5.3.2 Protein sidechains (i)

In the following table, the Percentiles column shows the percent sidechain outliers of the chain as a percentile score with respect to all PDB entries followed by that with respect to all EM entries.

For Manuscript Review

The Analysed column shows the number of residues for which the sidechain conformation was analysed, and the total number of residues.

| Mol | Chain | Analysed | Rotameric | Outliers | Percentiles | |
|---|---|---|---|---|---|---|
| 1 | A | 482/739 (65%) | 482 (100%) | 0 | 100 | 100 |
| 1 | B | 481/739 (65%) | 481 (100%) | 0 | 100 | 100 |
| 1 | C | 480/739 (65%) | 480 (100%) | 0 | 100 | 100 |
| 1 | D | 479/739 (65%) | 479 (100%) | 0 | 100 | 100 |
| 1 | E | 442/739 (60%) | 442 (100%) | 0 | 100 | 100 |
| 1 | F | 374/739 (51%) | 374 (100%) | 0 | 100 | 100 |
| 2 | P | 24/24 (100%) | 24 (100%) | 0 | 100 | 100 |
| All | All | 2762/4458 (62%) | 2762 (100%) | 0 | 100 | 100 |

There are no protein residues with a non-rotameric sidechain to report.

Sometimes sidechains can be flipped to improve hydrogen bonding and reduce clashes. All (3) such sidechains are listed below:

| Mol | Chain | Res | Type |
|---|---|---|---|
| 1 | B | 725 | ASN |
| 1 | C | 725 | ASN |
| 1 | E | 775 | ASN |

##### 5.3.3 RNA (i)

There are no RNA molecules in this entry.

#### 5.4 Non-standard residues in protein, DNA, RNA chains (i)

There are no non-standard protein/DNA/RNA residues in this entry.

#### 5.5 Carbohydrates (i)

There are no monosaccharides in this entry.

#### 5.6 Ligand geometry (i)

Of 19 ligands modelled in this entry, 8 are monoatomic - leaving 11 for Mogul analysis.

For Manuscript Review

In the following table, the Counts columns list the number of bonds (or angles) for which Mogul statistics could be retrieved, the number of bonds (or angles) that are observed in the model and the number of bonds (or angles) that are defined in the Chemical Component Dictionary. The Link column lists molecule types, if any, to which the group is linked. The Z score for a bond length (or angle) is the number of standard deviations the observed value is removed from the expected value. A bond length (or angle) with $|Z| > 2$ is considered an outlier worth inspection. RMSZ is the root-mean-square of all Z scores of the bond lengths (or angles).

| Mol | Type | Chain | Res | Link | Bond lengths | | | Bond angles | | |
|---|---|---|---|---|---|---|---|---|---|---|
| | | | | | Counts | RMSZ | $\#\|Z\| > 2$ | Counts | RMSZ | $\#\|Z\| > 2$ |
| 5 | A1AP0 | F | 1001 | - | 28,31,31 | 2.59 | 12 (42%) | 35,43,43 | 3.55 | 16 (45%) |
| 3 | ATP | B | 901 | 4 | 26,33,33 | 0.67 | 0 | 31,52,52 | 0.74 | 1 (3%) |
| 3 | ATP | D | 901 | 4 | 26,33,33 | 0.63 | 0 | 31,52,52 | 0.69 | 1 (3%) |
| 3 | ATP | D | 902 | 4 | 26,33,33 | 0.62 | 0 | 31,52,52 | 0.67 | 0 |
| 5 | A1AP0 | E | 902 | - | 28,31,31 | 2.42 | 11 (39%) | 35,43,43 | 3.54 | 15 (42%) |
| 3 | ATP | A | 902 | 4 | 26,33,33 | 0.63 | 0 | 31,52,52 | 0.70 | 1 (3%) |
| 3 | ATP | C | 902 | 4 | 26,33,33 | 0.64 | 0 | 31,52,52 | 0.70 | 1 (3%) |
| 3 | ATP | C | 901 | 4 | 26,33,33 | 0.65 | 0 | 31,52,52 | 0.71 | 1 (3%) |
| 3 | ATP | A | 901 | 4 | 26,33,33 | 0.62 | 0 | 31,52,52 | 0.70 | 1 (3%) |
| 3 | ATP | E | 901 | - | 26,33,33 | 0.61 | 0 | 31,52,52 | 0.66 | 0 |
| 3 | ATP | B | 902 | 4 | 26,33,33 | 0.66 | 0 | 31,52,52 | 0.72 | 1 (3%) |

In the following table, the Chirals column lists the number of chiral outliers, the number of chiral centers analysed, the number of these observed in the model and the number defined in the Chemical Component Dictionary. Similar counts are reported in the Torsion and Rings columns. '-' means no outliers of that kind were identified.

| Mol | Type | Chain | Res | Link | Chirals | Torsions | Rings |
|---|---|---|---|---|---|---|---|
| 5 | A1AP0 | F | 1001 | - | - | 8/12/28/28 | 0/3/4/4 |
| 3 | ATP | B | 901 | 4 | - | 2/18/38/38 | 0/3/3/3 |
| 3 | ATP | D | 901 | 4 | - | 5/18/38/38 | 0/3/3/3 |
| 3 | ATP | D | 902 | 4 | - | 4/18/38/38 | 0/3/3/3 |
| 5 | A1AP0 | E | 902 | - | - | 7/12/28/28 | 0/3/4/4 |
| 3 | ATP | A | 902 | 4 | - | 0/18/38/38 | 0/3/3/3 |
| 3 | ATP | C | 902 | 4 | - | 2/18/38/38 | 0/3/3/3 |
| 3 | ATP | C | 901 | 4 | - | 3/18/38/38 | 0/3/3/3 |
| 3 | ATP | A | 901 | 4 | - | 5/18/38/38 | 0/3/3/3 |
| 3 | ATP | E | 901 | - | - | 4/18/38/38 | 0/3/3/3 |
| 3 | ATP | B | 902 | 4 | - | 2/18/38/38 | 0/3/3/3 |

All (23) bond length outliers are listed below:

For Manuscript Review

| Mol | Chain | Res | Type | Atoms | Z | Observed(Å) | Ideal(Å) |
|---|---|---|---|---|---|---|---|
| 5 | F | 1001 | A1AP0 | C15-N05 | 8.19 | 1.47 | 1.37 |
| 5 | E | 902 | A1AP0 | C15-N05 | 7.95 | 1.47 | 1.37 |
| 5 | F | 1001 | A1AP0 | C06-N05 | 5.14 | 1.48 | 1.43 |
| 5 | E | 902 | A1AP0 | C06-N05 | 4.21 | 1.47 | 1.43 |
| 5 | F | 1001 | A1AP0 | C18-N19 | 3.72 | 1.49 | 1.41 |
| 5 | F | 1001 | A1AP0 | C07-S08 | 3.62 | 1.81 | 1.78 |
| 5 | E | 902 | A1AP0 | C07-S08 | 3.45 | 1.81 | 1.78 |
| 5 | F | 1001 | A1AP0 | C09-S08 | 3.29 | 1.81 | 1.78 |
| 5 | E | 902 | A1AP0 | C18-N19 | 3.23 | 1.48 | 1.41 |
| 5 | E | 902 | A1AP0 | C09-S08 | 3.19 | 1.81 | 1.78 |
| 5 | E | 902 | A1AP0 | C14-C15 | 2.94 | 1.54 | 1.50 |
| 5 | F | 1001 | A1AP0 | C14-C15 | 2.84 | 1.54 | 1.50 |
| 5 | F | 1001 | A1AP0 | C04-N05 | 2.67 | 1.54 | 1.47 |
| 5 | F | 1001 | A1AP0 | C20-N19 | 2.43 | 1.42 | 1.35 |
| 5 | E | 902 | A1AP0 | C04-N05 | 2.42 | 1.53 | 1.47 |
| 5 | F | 1001 | A1AP0 | C28-C06 | 2.30 | 1.43 | 1.39 |
| 5 | E | 902 | A1AP0 | C28-C06 | 2.26 | 1.43 | 1.39 |
| 5 | E | 902 | A1AP0 | C20-N19 | 2.19 | 1.41 | 1.35 |
| 5 | F | 1001 | A1AP0 | C04-C03 | 2.18 | 1.57 | 1.50 |
| 5 | F | 1001 | A1AP0 | C17-C07 | 2.16 | 1.43 | 1.39 |
| 5 | E | 902 | A1AP0 | C04-C03 | 2.14 | 1.57 | 1.50 |
| 5 | F | 1001 | A1AP0 | C26-C22 | 2.12 | 1.45 | 1.42 |
| 5 | E | 902 | A1AP0 | C17-C07 | 2.07 | 1.42 | 1.39 |

All (38) bond angle outliers are listed below:

| Mol | Chain | Res | Type | Atoms | Z | Observed($^o$) | Ideal($^o$) |
|---|---|---|---|---|---|---|---|
| 5 | E | 902 | A1AP0 | C14-C15-N05 | 7.95 | 131.54 | 119.63 |
| 5 | F | 1001 | A1AP0 | C14-C15-N05 | 7.92 | 131.49 | 119.63 |
| 5 | E | 902 | A1AP0 | C06-N05-C15 | 7.13 | 132.57 | 124.96 |
| 5 | F | 1001 | A1AP0 | C06-N05-C15 | 6.98 | 132.42 | 124.96 |
| 5 | F | 1001 | A1AP0 | C09-S08-C07 | 6.92 | 117.29 | 99.13 |
| 5 | E | 902 | A1AP0 | C09-S08-C07 | 6.88 | 117.18 | 99.13 |
| 5 | F | 1001 | A1AP0 | C07-C06-N05 | 6.09 | 125.55 | 121.07 |
| 5 | E | 902 | A1AP0 | C22-C20-N19 | 6.08 | 129.31 | 115.92 |
| 5 | F | 1001 | A1AP0 | C27-C18-C17 | -5.90 | 112.66 | 119.65 |
| 5 | E | 902 | A1AP0 | C04-N05-C06 | -5.83 | 111.94 | 118.41 |
| 5 | E | 902 | A1AP0 | C07-C06-N05 | 5.81 | 125.34 | 121.07 |
| 5 | F | 1001 | A1AP0 | C04-N05-C06 | -5.78 | 112.00 | 118.41 |
| 5 | E | 902 | A1AP0 | O16-C15-N05 | -5.73 | 115.16 | 121.01 |
| 5 | F | 1001 | A1AP0 | C22-C20-N19 | 5.71 | 128.48 | 115.92 |
| 5 | E | 902 | A1AP0 | C27-C18-C17 | -5.64 | 112.96 | 119.65 |
| 5 | F | 1001 | A1AP0 | O16-C15-N05 | -5.47 | 115.43 | 121.01 |

*Continued on next page...*

For Manuscript Review

*Continued from previous page...*

| Mol | Chain | Res | Type | Atoms | Z | Observed(°) | Ideal(°) |
|---|---|---|---|---|---|---|---|
| 5 | F | 1001 | A1AP0 | C28-C06-C07 | -4.50 | 112.34 | 118.65 |
| 5 | E | 902 | A1AP0 | C28-C06-C07 | -4.37 | 112.53 | 118.65 |
| 5 | F | 1001 | A1AP0 | O16-C15-C14 | -4.12 | 113.09 | 120.01 |
| 5 | E | 902 | A1AP0 | O16-C15-C14 | -3.99 | 113.30 | 120.01 |
| 5 | E | 902 | A1AP0 | O21-C20-C22 | -3.45 | 114.78 | 120.94 |
| 5 | E | 902 | A1AP0 | O21-C20-N19 | -3.42 | 115.91 | 123.71 |
| 5 | F | 1001 | A1AP0 | O21-C20-C22 | -3.23 | 115.17 | 120.94 |
| 5 | F | 1001 | A1AP0 | O21-C20-N19 | -3.22 | 116.35 | 123.71 |
| 5 | F | 1001 | A1AP0 | C27-C28-C06 | 2.74 | 124.83 | 119.19 |
| 5 | E | 902 | A1AP0 | C27-C28-C06 | 2.73 | 124.80 | 119.19 |
| 5 | E | 902 | A1AP0 | C28-C06-N05 | 2.61 | 122.12 | 119.24 |
| 5 | F | 1001 | A1AP0 | C28-C06-N05 | 2.60 | 122.10 | 119.24 |
| 5 | F | 1001 | A1AP0 | C28-C27-C18 | 2.32 | 122.98 | 120.30 |
| 3 | B | 901 | ATP | C5-C6-N6 | 2.21 | 123.72 | 120.35 |
| 5 | E | 902 | A1AP0 | C28-C27-C18 | 2.13 | 122.76 | 120.30 |
| 3 | B | 902 | ATP | PB-O3B-PG | 2.13 | 140.13 | 132.83 |
| 5 | F | 1001 | A1AP0 | C03-C04-N05 | 2.12 | 116.07 | 112.56 |
| 3 | A | 902 | ATP | PB-O3B-PG | 2.12 | 140.09 | 132.83 |
| 3 | A | 901 | ATP | PB-O3B-PG | 2.10 | 140.04 | 132.83 |
| 3 | D | 901 | ATP | PB-O3B-PG | 2.04 | 139.84 | 132.83 |
| 3 | C | 902 | ATP | PB-O3B-PG | 2.02 | 139.75 | 132.83 |
| 3 | C | 901 | ATP | PB-O3B-PG | 2.02 | 139.75 | 132.83 |

There are no chirality outliers.

All (42) torsion outliers are listed below:

| Mol | Chain | Res | Type | Atoms |
|---|---|---|---|---|
| 3 | A | 901 | ATP | C5'-O5'-PA-O2A |
| 3 | D | 901 | ATP | C5'-O5'-PA-O2A |
| 5 | E | 902 | A1AP0 | N19-C20-C22-C23 |
| 5 | E | 902 | A1AP0 | N19-C20-C22-C26 |
| 5 | E | 902 | A1AP0 | O21-C20-C22-C23 |
| 5 | E | 902 | A1AP0 | O21-C20-C22-C26 |
| 5 | F | 1001 | A1AP0 | C04-C03-O02-C01 |
| 5 | F | 1001 | A1AP0 | N19-C20-C22-C23 |
| 5 | F | 1001 | A1AP0 | N19-C20-C22-C26 |
| 5 | F | 1001 | A1AP0 | O21-C20-C22-C23 |
| 5 | F | 1001 | A1AP0 | O21-C20-C22-C26 |
| 5 | F | 1001 | A1AP0 | C17-C18-N19-C20 |
| 5 | F | 1001 | A1AP0 | C27-C18-N19-C20 |
| 5 | F | 1001 | A1AP0 | O02-C03-C04-N05 |
| 5 | E | 902 | A1AP0 | C27-C18-N19-C20 |

*Continued on next page...*

For Manuscript Review

Continued from previous page...

| Mol | Chain | Res | Type | Atoms |
|---|---|---|---|---|
| 5 | E | 902 | A1AP0 | C17-C18-N19-C20 |
| 3 | B | 901 | ATP | O4'-C4'-C5'-O5' |
| 3 | E | 901 | ATP | O4'-C4'-C5'-O5' |
| 3 | E | 901 | ATP | C3'-C4'-C5'-O5' |
| 3 | B | 902 | ATP | PA-O3A-PB-O1B |
| 3 | B | 901 | ATP | C3'-C4'-C5'-O5' |
| 5 | E | 902 | A1AP0 | C04-C03-O02-C01 |
| 3 | A | 901 | ATP | C5'-O5'-PA-O3A |
| 3 | D | 901 | ATP | C5'-O5'-PA-O3A |
| 3 | A | 901 | ATP | PA-O3A-PB-O1B |
| 3 | C | 901 | ATP | PA-O3A-PB-O1B |
| 3 | D | 901 | ATP | PA-O3A-PB-O1B |
| 3 | D | 902 | ATP | PG-O3B-PB-O1B |
| 3 | A | 901 | ATP | C5'-O5'-PA-O1A |
| 3 | D | 901 | ATP | C5'-O5'-PA-O1A |
| 3 | C | 902 | ATP | PA-O3A-PB-O2B |
| 3 | D | 902 | ATP | PB-O3A-PA-O2A |
| 3 | E | 901 | ATP | C4'-C5'-O5'-PA |
| 3 | C | 902 | ATP | PA-O3A-PB-O1B |
| 3 | D | 902 | ATP | PB-O3A-PA-O1A |
| 3 | A | 901 | ATP | PA-O3A-PB-O2B |
| 3 | B | 902 | ATP | PA-O3A-PB-O2B |
| 3 | C | 901 | ATP | PA-O3A-PB-O2B |
| 3 | D | 901 | ATP | PA-O3A-PB-O2B |
| 3 | D | 902 | ATP | PG-O3B-PB-O2B |
| 3 | C | 901 | ATP | C5'-O5'-PA-O1A |
| 3 | E | 901 | ATP | C5'-O5'-PA-O1A |

There are no ring outliers.

4 monomers are involved in 5 short contacts:

| Mol | Chain | Res | Type | Clashes | Symm-Clashes |
|---|---|---|---|---|---|
| 5 | F | 1001 | A1AP0 | 1 | 0 |
| 3 | B | 901 | ATP | 2 | 0 |
| 3 | D | 902 | ATP | 1 | 0 |
| 3 | A | 901 | ATP | 1 | 0 |

The following is a two-dimensional graphical depiction of Mogul quality analysis of bond lengths, bond angles, torsion angles, and ring geometry for all instances of the Ligand of Interest. In addition, ligands with molecular weight > 250 and outliers as shown on the validation Tables will also be included. For torsion angles, if less then 5% of the Mogul distribution of torsion angles is within 10 degrees of the torsion angle in question, then that torsion angle is considered an outlier.

For Manuscript Review

Any bond that is central to one or more torsion angles identified as an outlier by Mogul will be highlighted in the graph. For rings, the root-mean-square deviation (RMSD) between the ring in question and similar rings identified by Mogul is calculated over all ring torsion angles. If the average RMSD is greater than 60 degrees and the minimal RMSD between the ring in question and any Mogul-identified rings is also greater than 60 degrees, then that ring is considered an outlier. The outliers are highlighted in purple. The color gray indicates Mogul did not find sufficient equivalents in the CSD to analyse the geometry.

**Ligand A1AP0 F 1001**

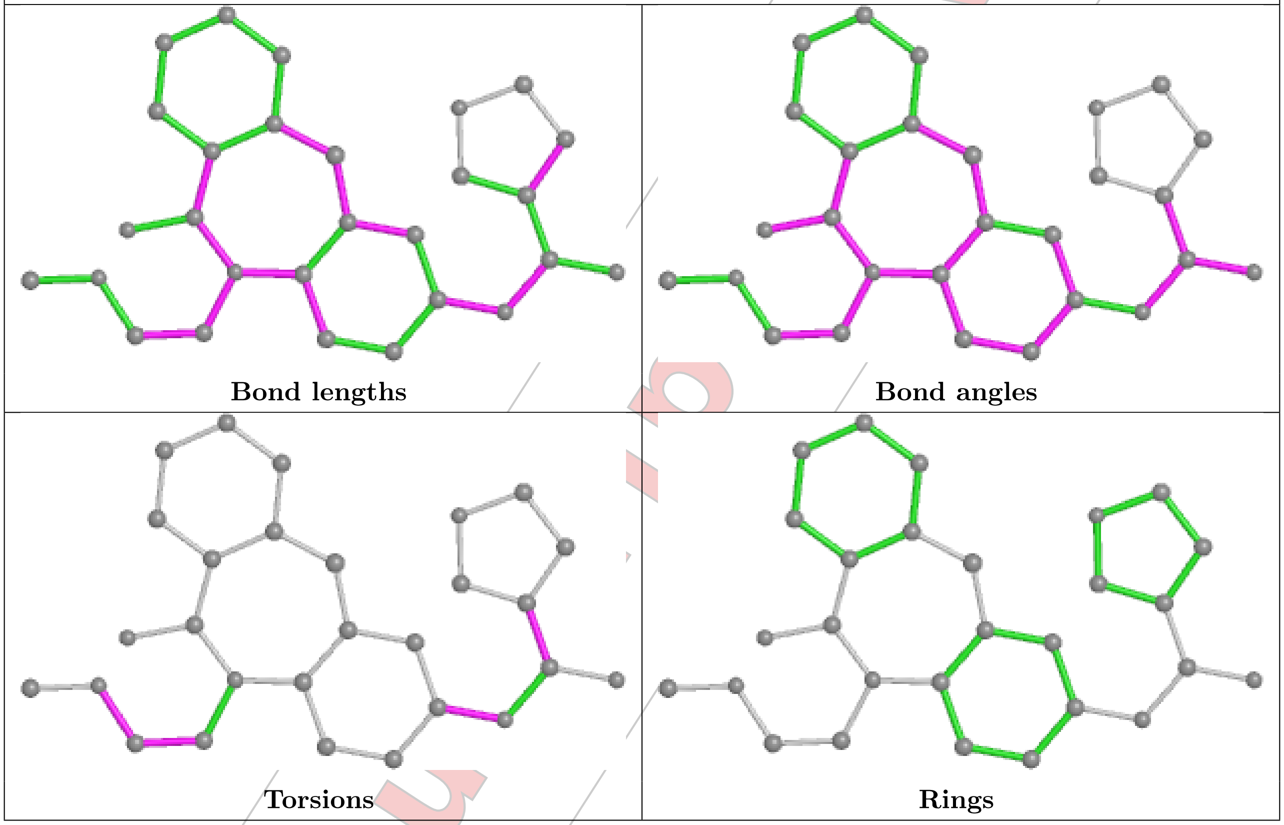


For Manuscript Review

**Ligand ATP B 901**

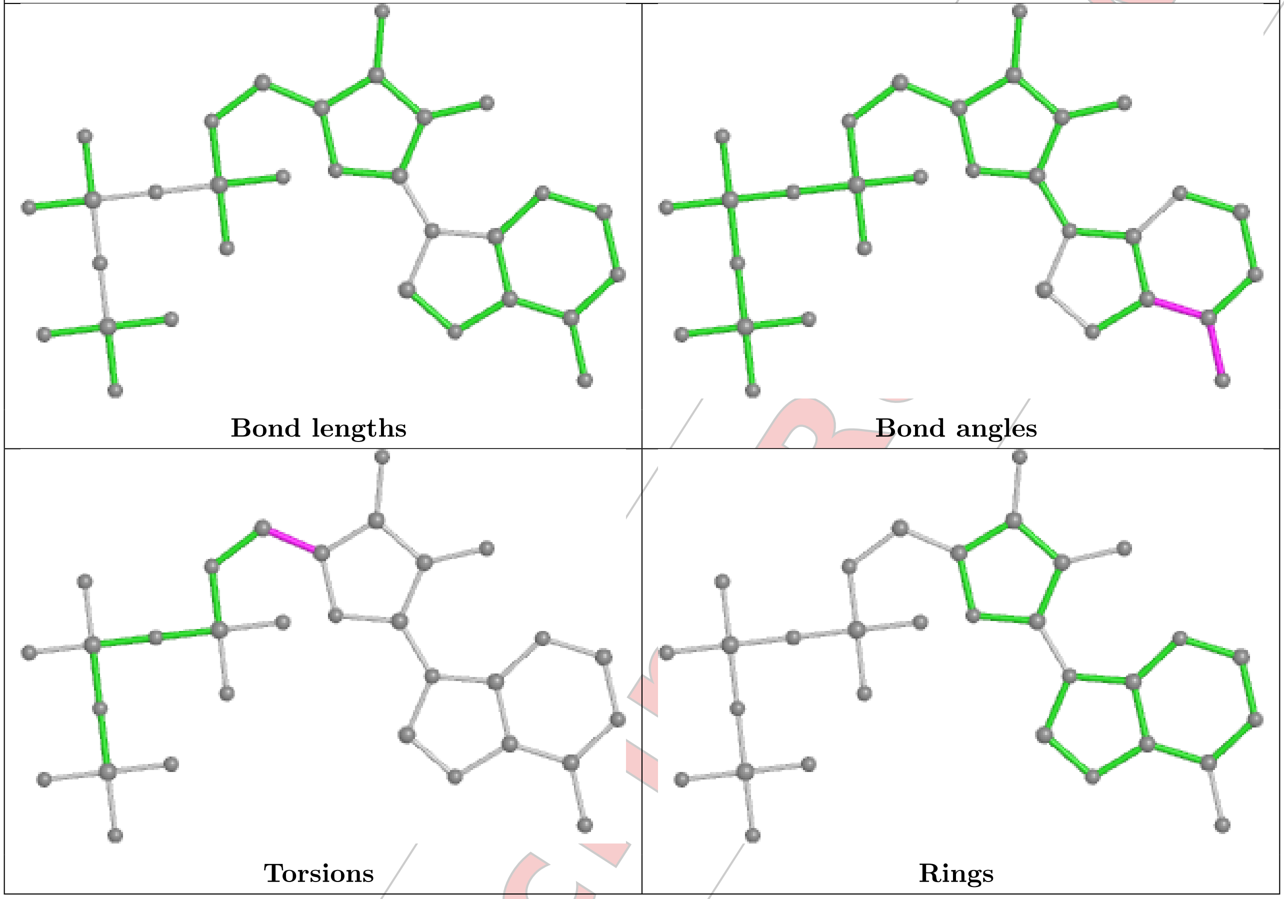


For Manuscript Review

Ligand ATP D 901

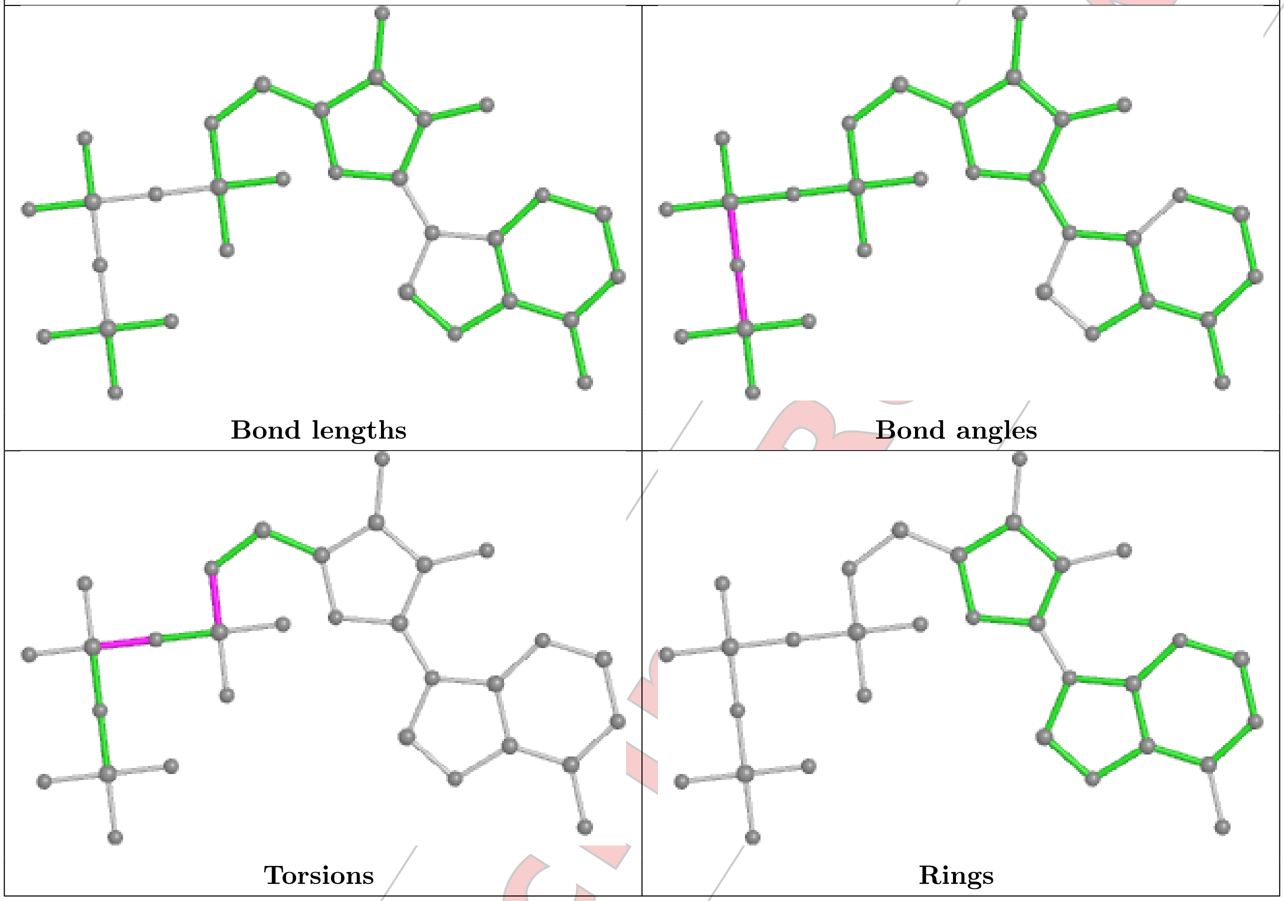


For Manuscript Review

#### Ligand ATP D 902

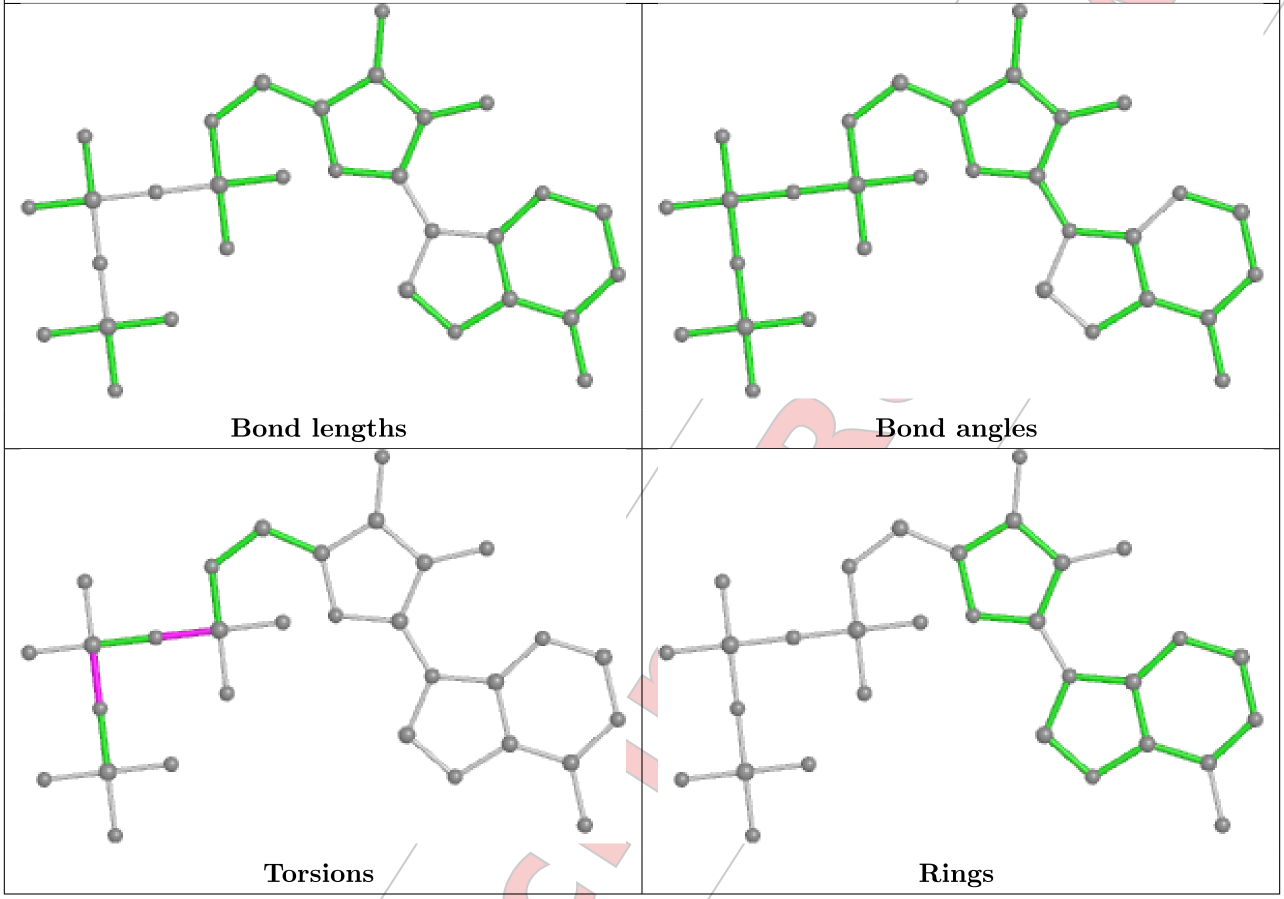


#### Ligand A1AP0 E 902

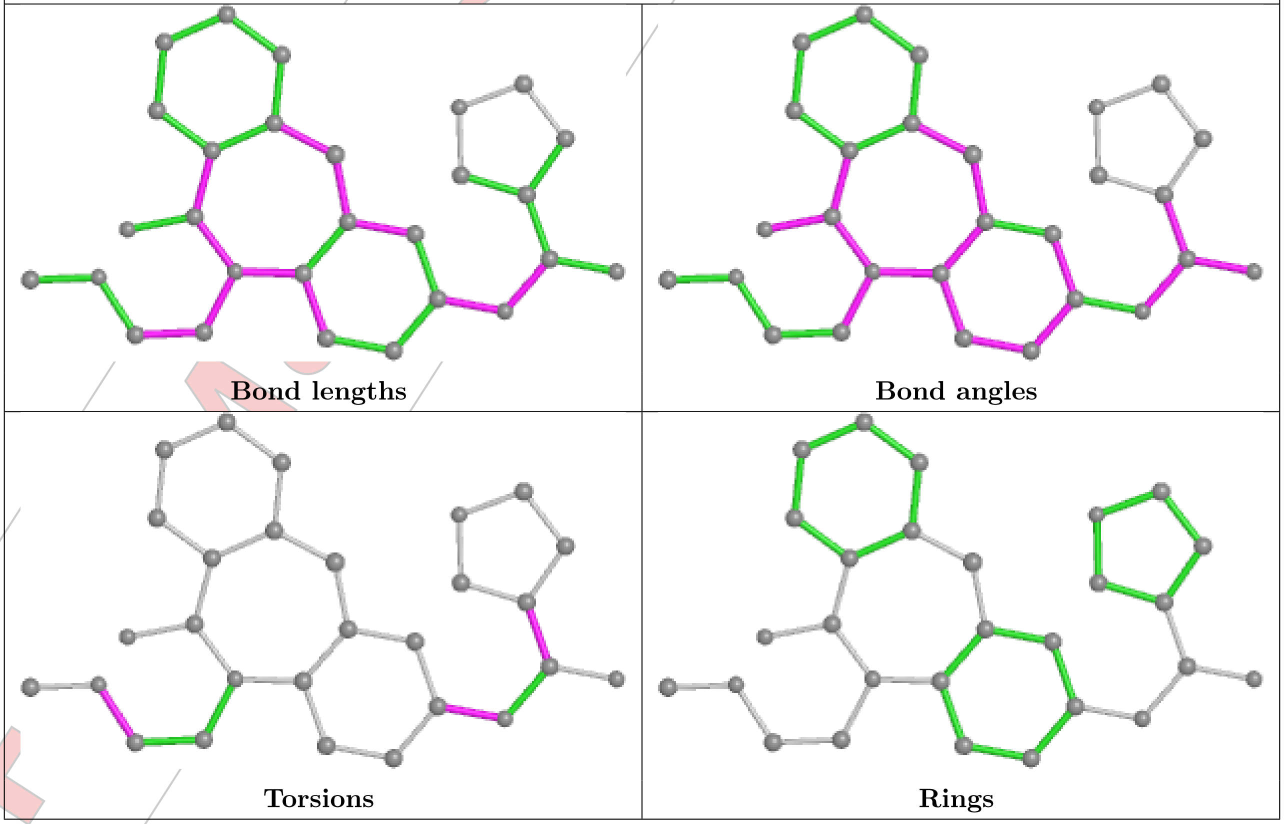

**Ligand ATP A 902**

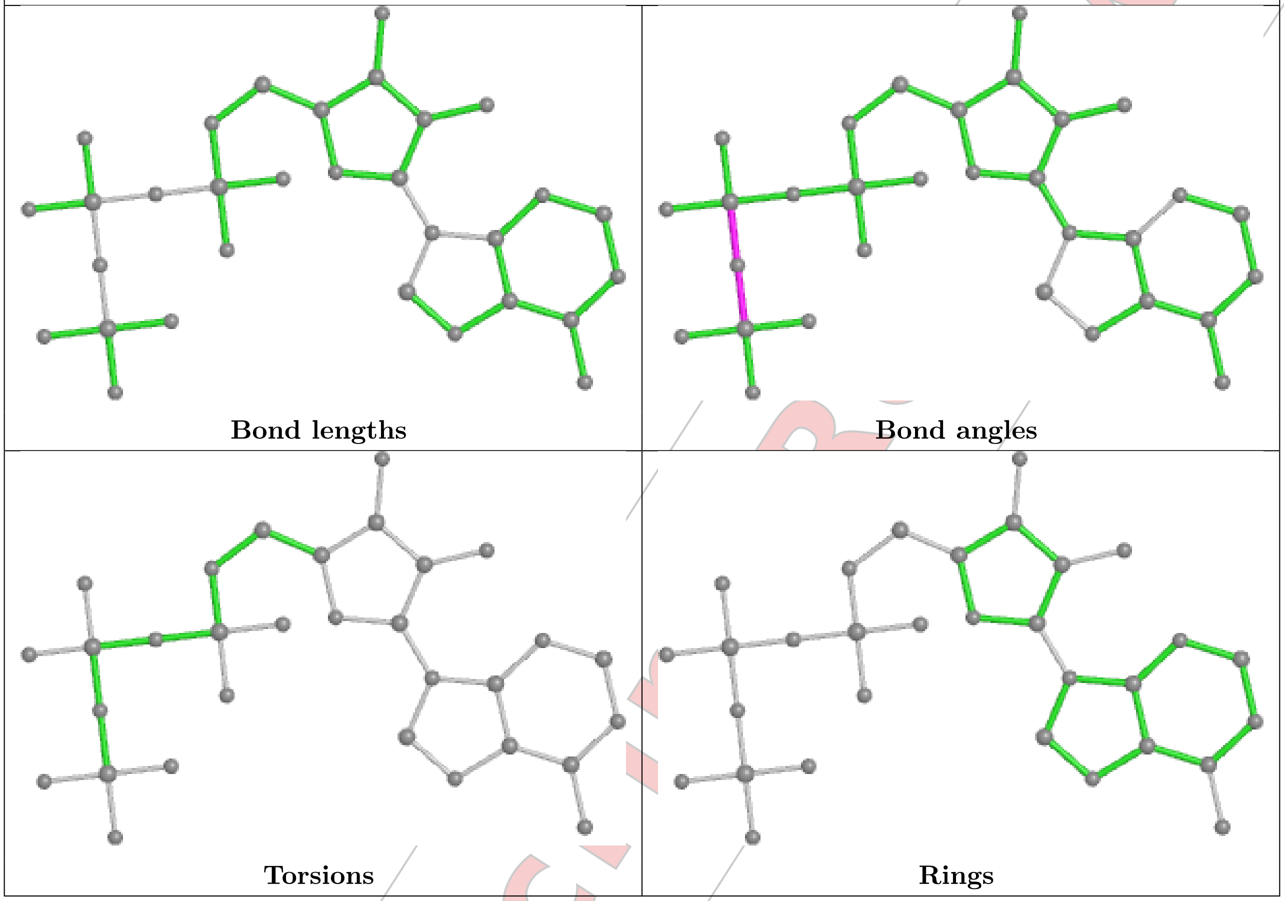


For Manuscript Review

**Ligand ATP C 902**

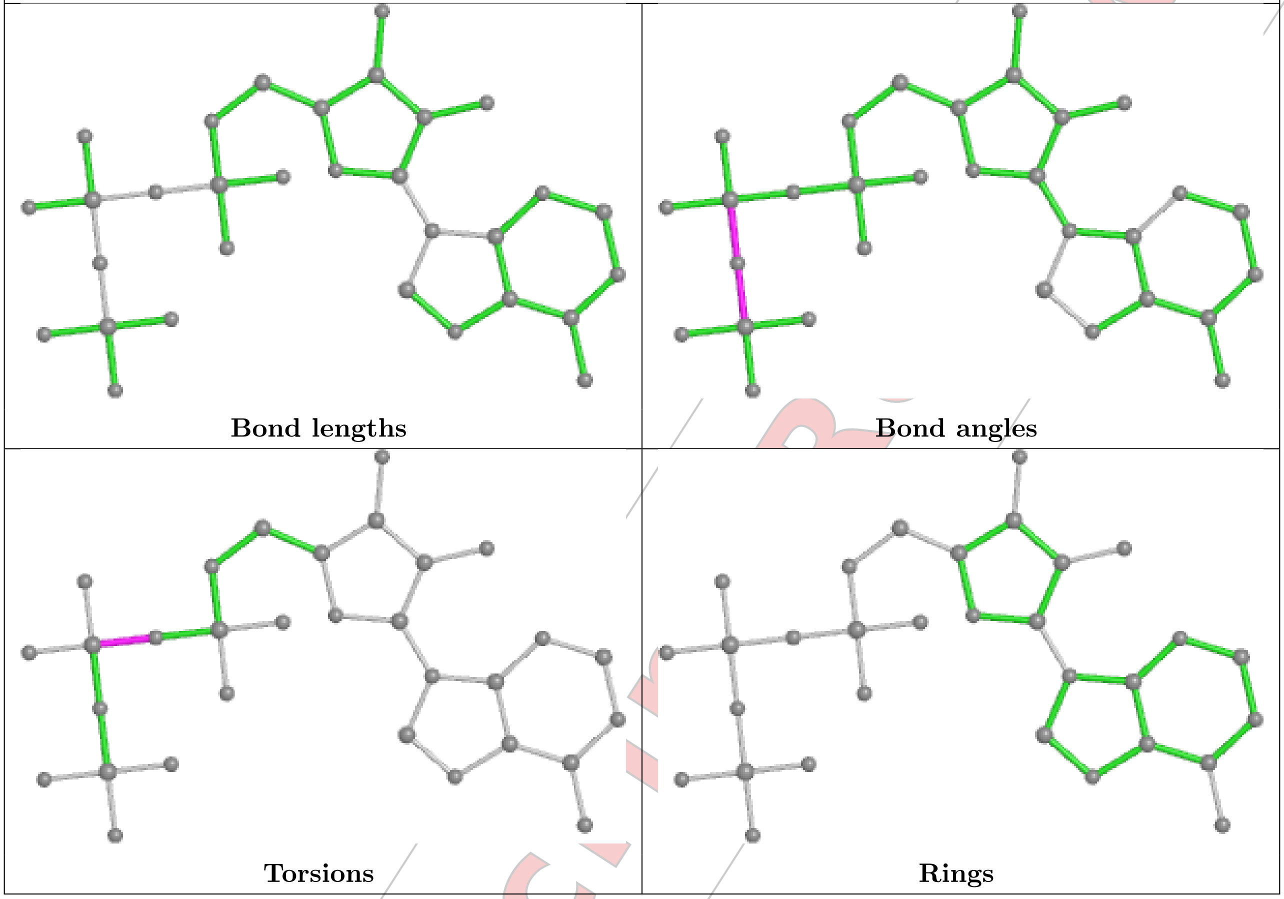


For Manuscript Review

**Ligand ATP C 901**

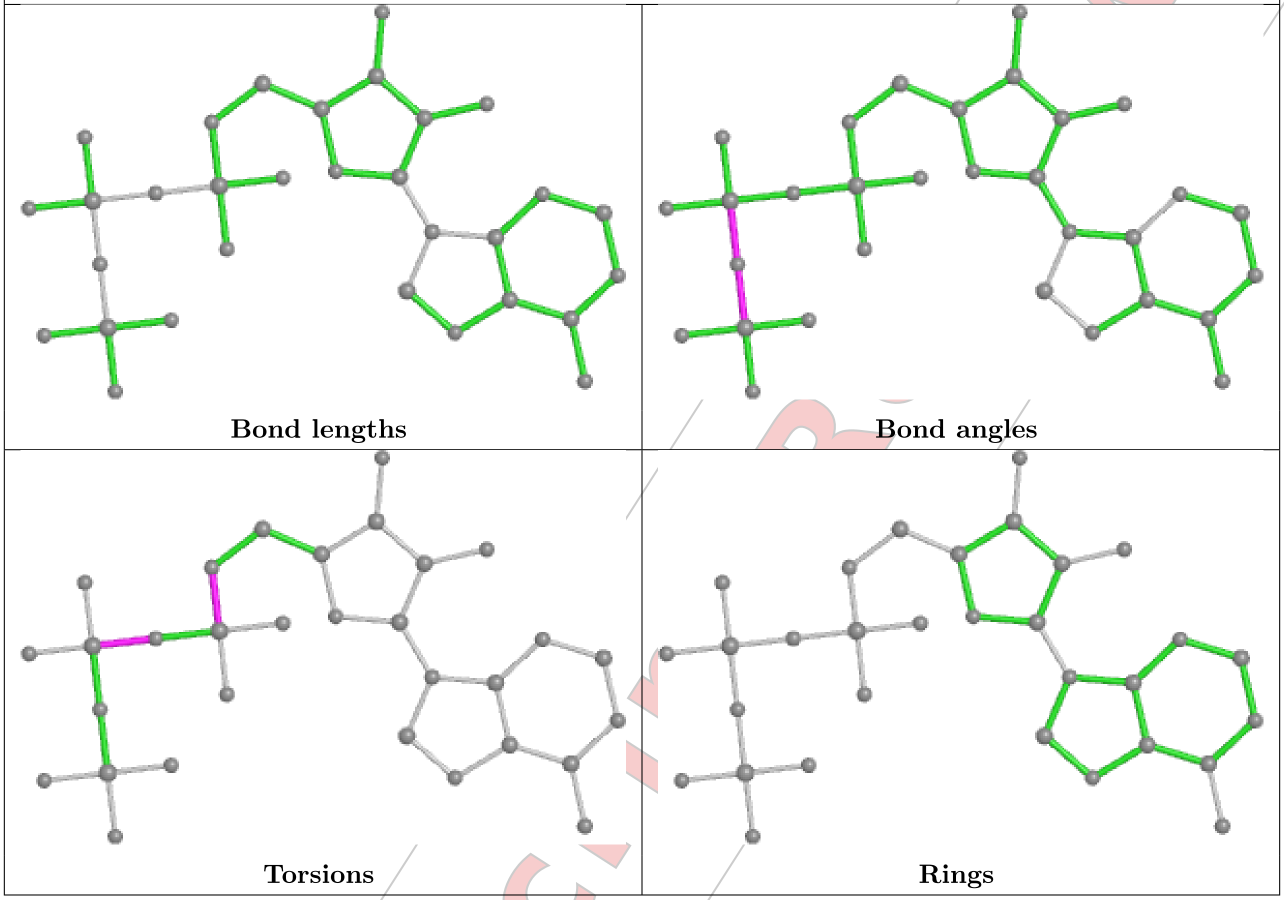


For Manuscript Review

**Ligand ATP A 901**

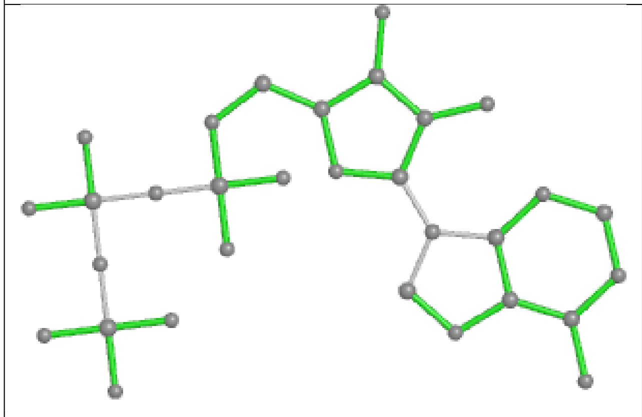

**Bond lengths**

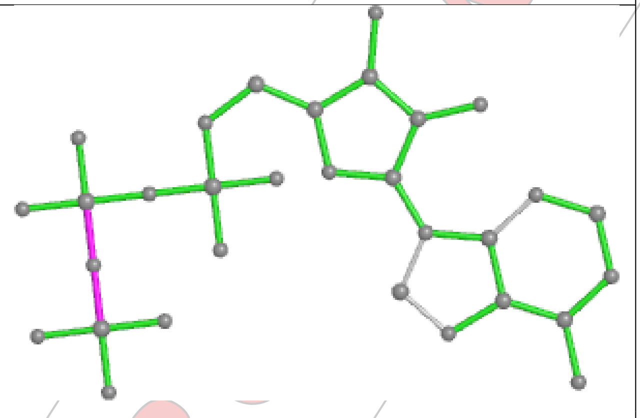

**Bond angles**

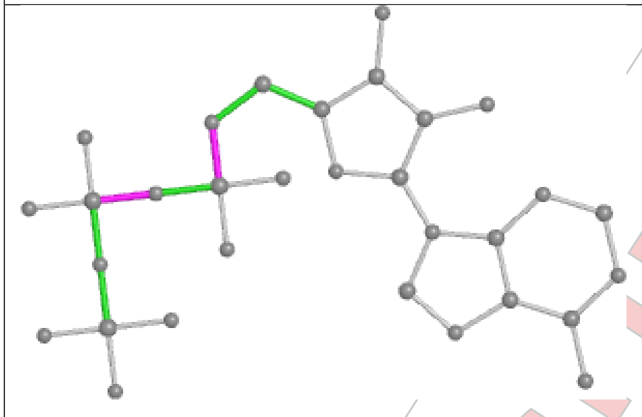

**Torsions**

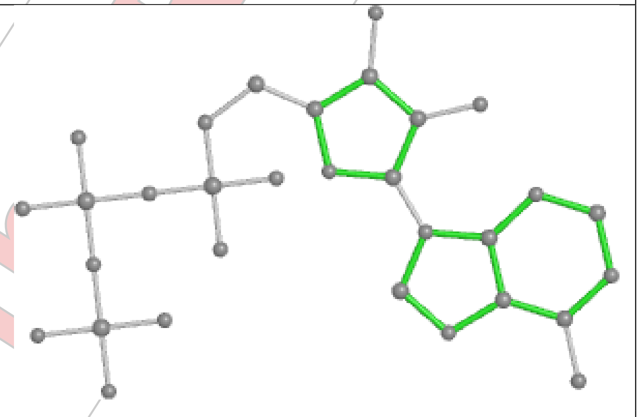

**Rings**

For Manuscript Review

**Ligand ATP E 901**

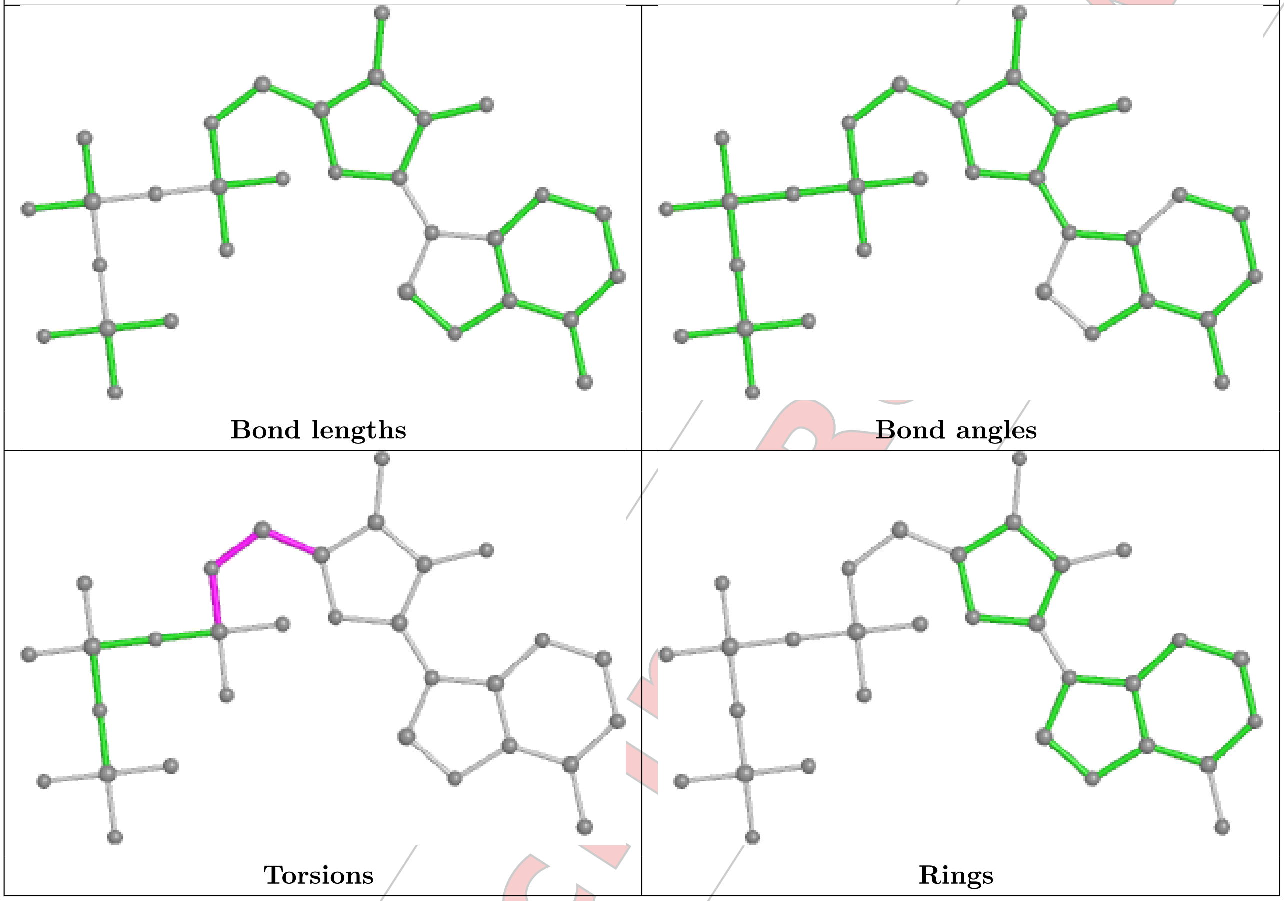


For Manuscript Review

**Ligand ATP B 902**

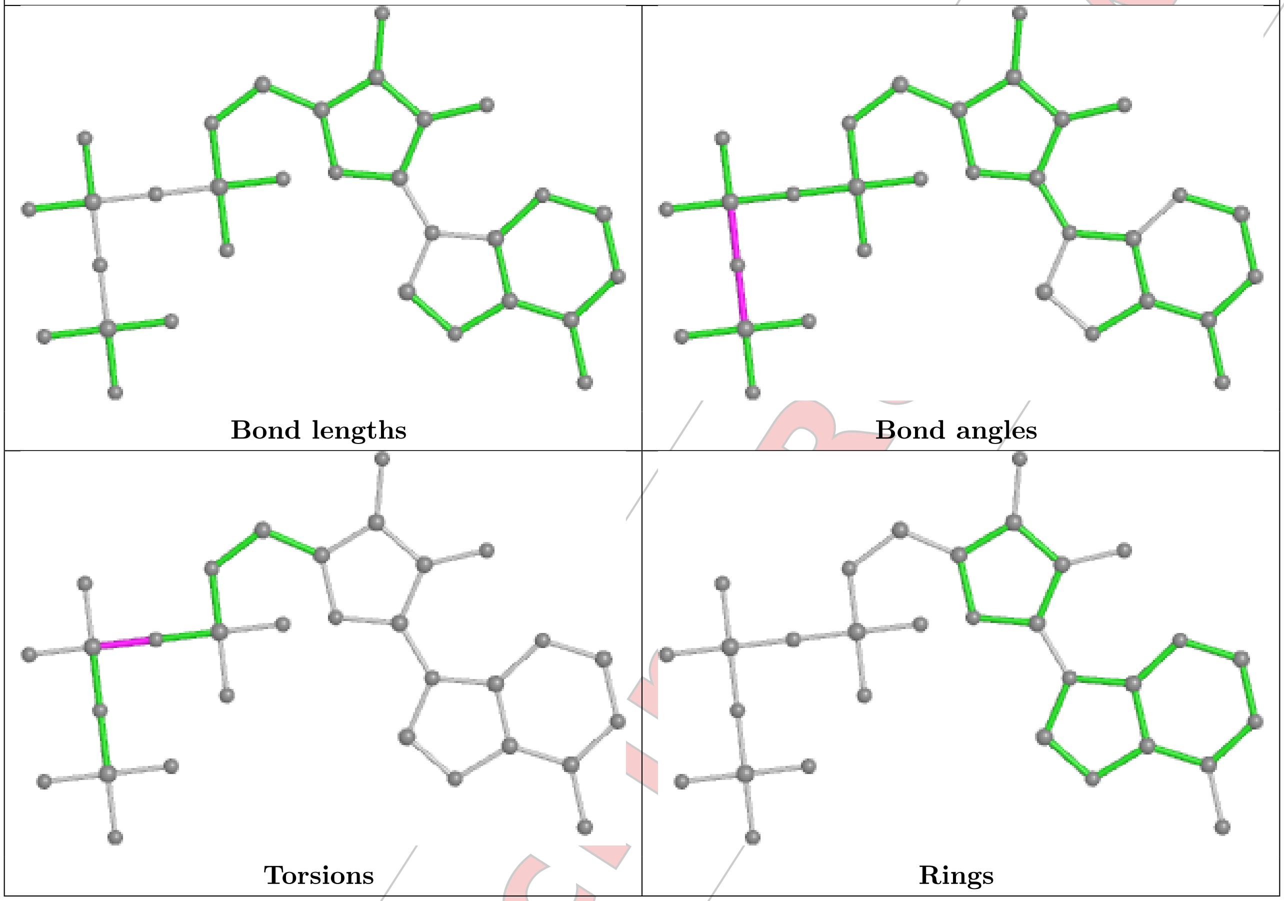


#### 5.7 Other polymers (i)

There are no such residues in this entry.

#### 5.8 Polymer linkage issues (i)

There are no chain breaks in this entry.

For Manuscript Review

### 6 Map visualisation (i)

This section contains visualisations of the EMDB entry EMD-44634. These allow visual inspection of the internal detail of the map and identification of artifacts.

Images derived from a raw map, generated by summing the deposited half-maps, are presented below the corresponding image components of the primary map to allow further visual inspection and comparison with those of the primary map.

#### 6.1 Orthogonal projections (i)

##### 6.1.1 Primary map

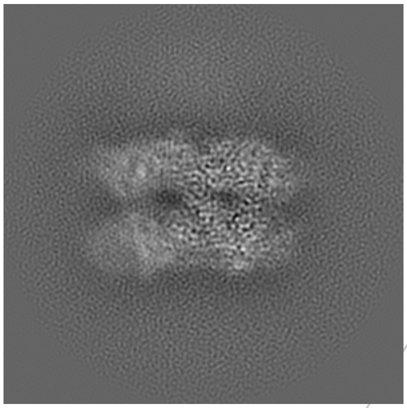

X

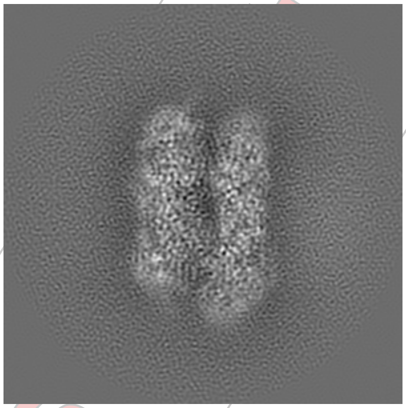

Y

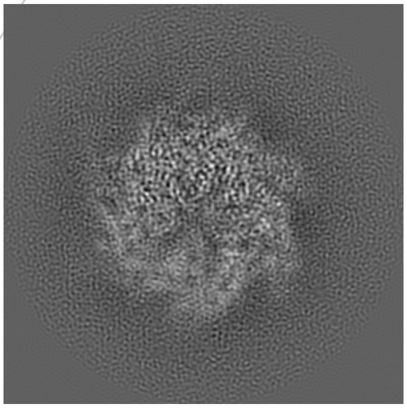

Z

##### 6.1.2 Raw map

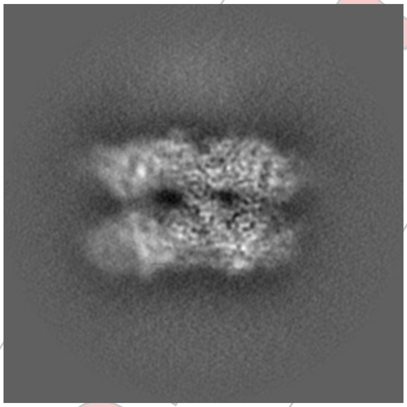

X

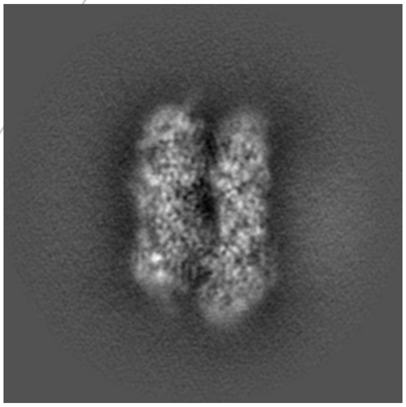

Y

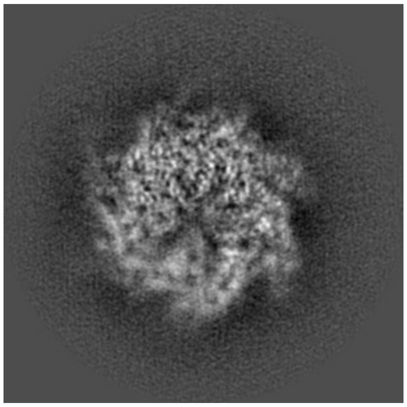

Z

The images above show the map projected in three orthogonal directions.

For Manuscript Review

#### 6.2 Central slices (i)

##### 6.2.1 Primary map

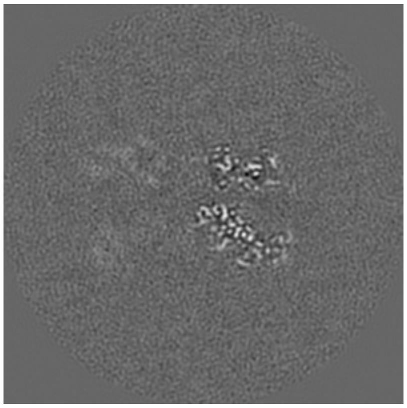
X Index: 140

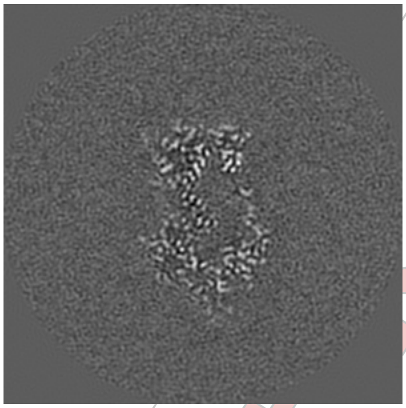
Y Index: 140

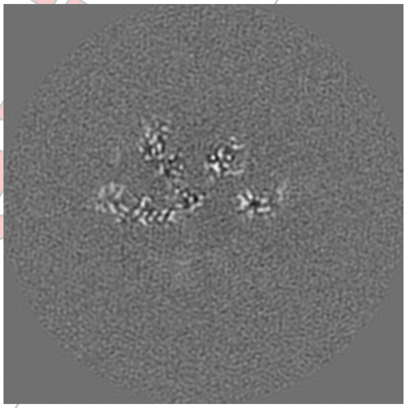
Z Index: 140

##### 6.2.2 Raw map

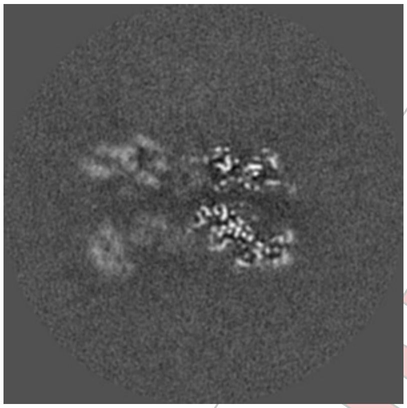
X Index: 140

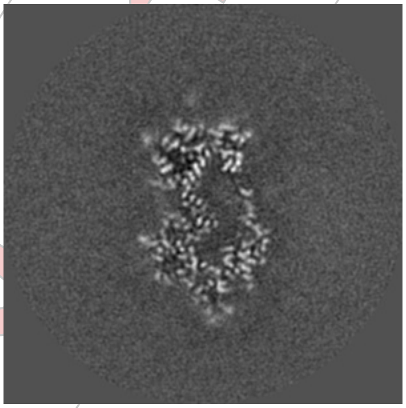
Y Index: 140

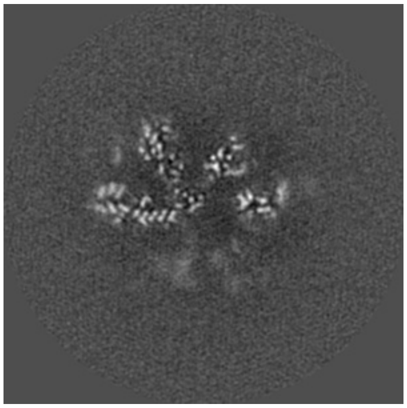
Z Index: 140

The images above show central slices of the map in three orthogonal directions.

For Manuscript Review

#### 6.3 Largest variance slices ⓘ

##### 6.3.1 Primary map

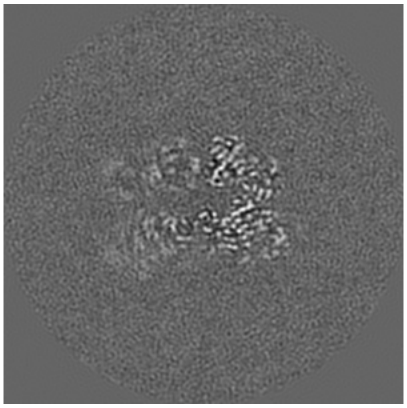
X Index: 158

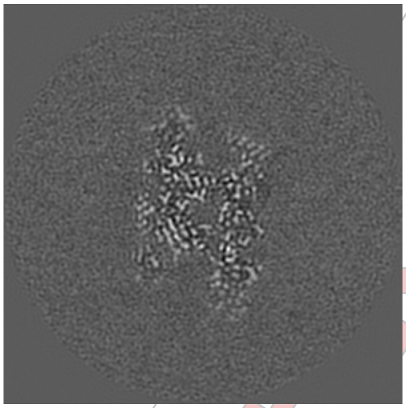
Y Index: 165

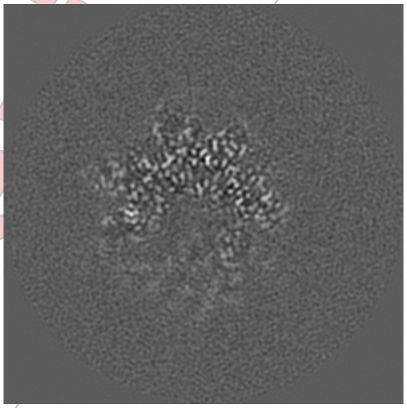
Z Index: 161

##### 6.3.2 Raw map

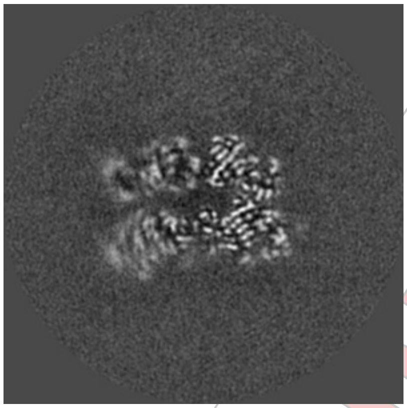
X Index: 158

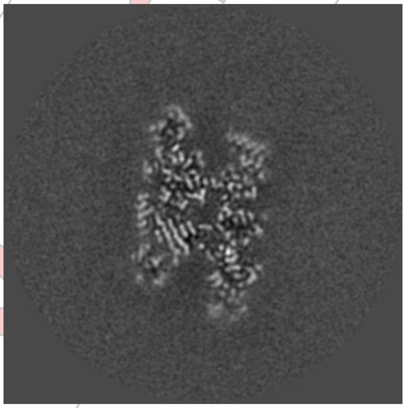
Y Index: 166

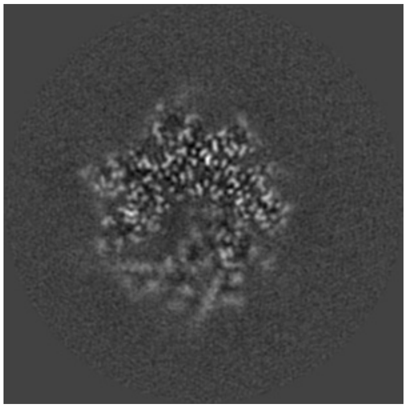
Z Index: 161

The images above show the largest variance slices of the map in three orthogonal directions.

For Manuscript Review

#### 6.4 Orthogonal standard-deviation projections (False-color) (i)

##### 6.4.1 Primary map

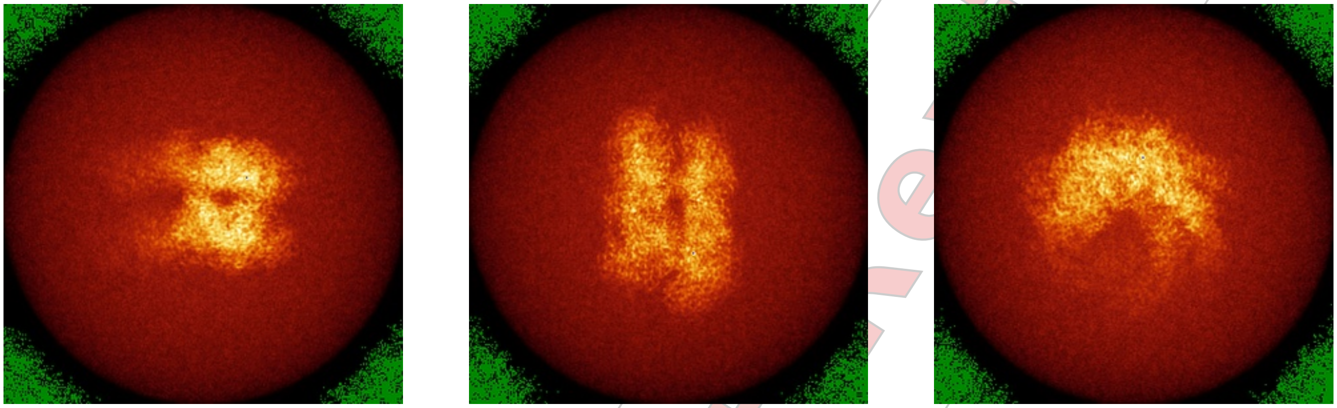

X Y Z

##### 6.4.2 Raw map

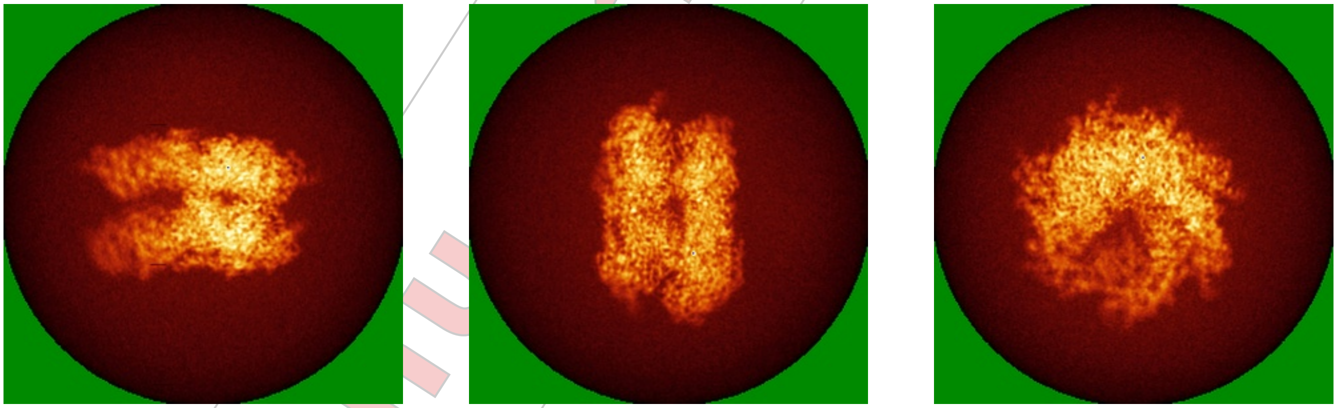

X Y Z

The images above show the map standard deviation projections with false color in three orthogonal directions. Minimum values are shown in green, max in blue, and dark to light orange shades represent small to large values respectively.

For Manuscript Review

#### 6.5 Orthogonal surface views

##### 6.5.1 Primary map

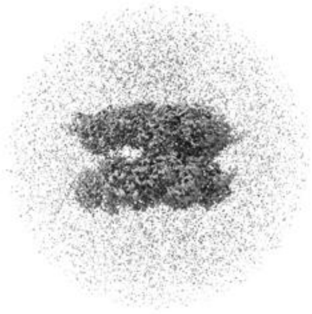

X

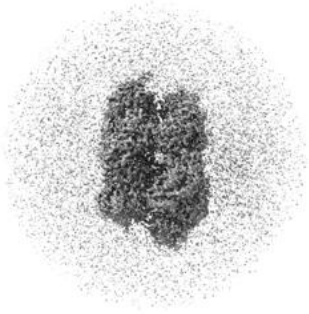

Y

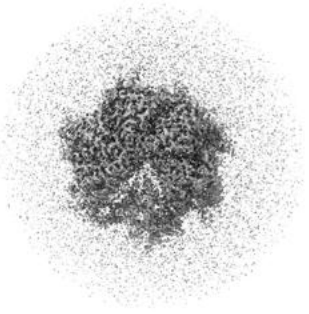

Z

The images above show the 3D surface view of the map at the recommended contour level 0.012. These images, in conjunction with the slice images, may facilitate assessment of whether an appropriate contour level has been provided.

##### 6.5.2 Raw map

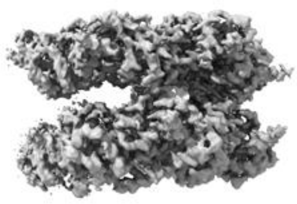

X

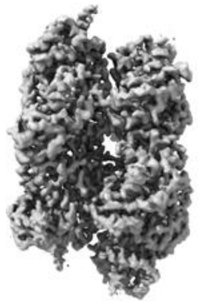

Y

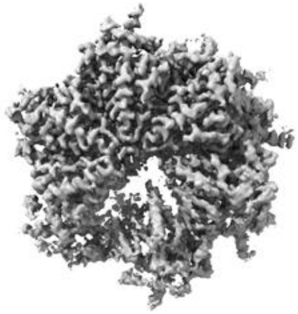

Z

These images show the 3D surface of the raw map. The raw map's contour level was selected so that its surface encloses the same volume as the primary map does at its recommended contour level.

#### 6.6 Mask visualisation

This section was not generated. No masks/segmentation were deposited.

For Manuscript Review

### 7 Map analysis (i)

This section contains the results of statistical analysis of the map.

#### 7.1 Map-value distribution (i)

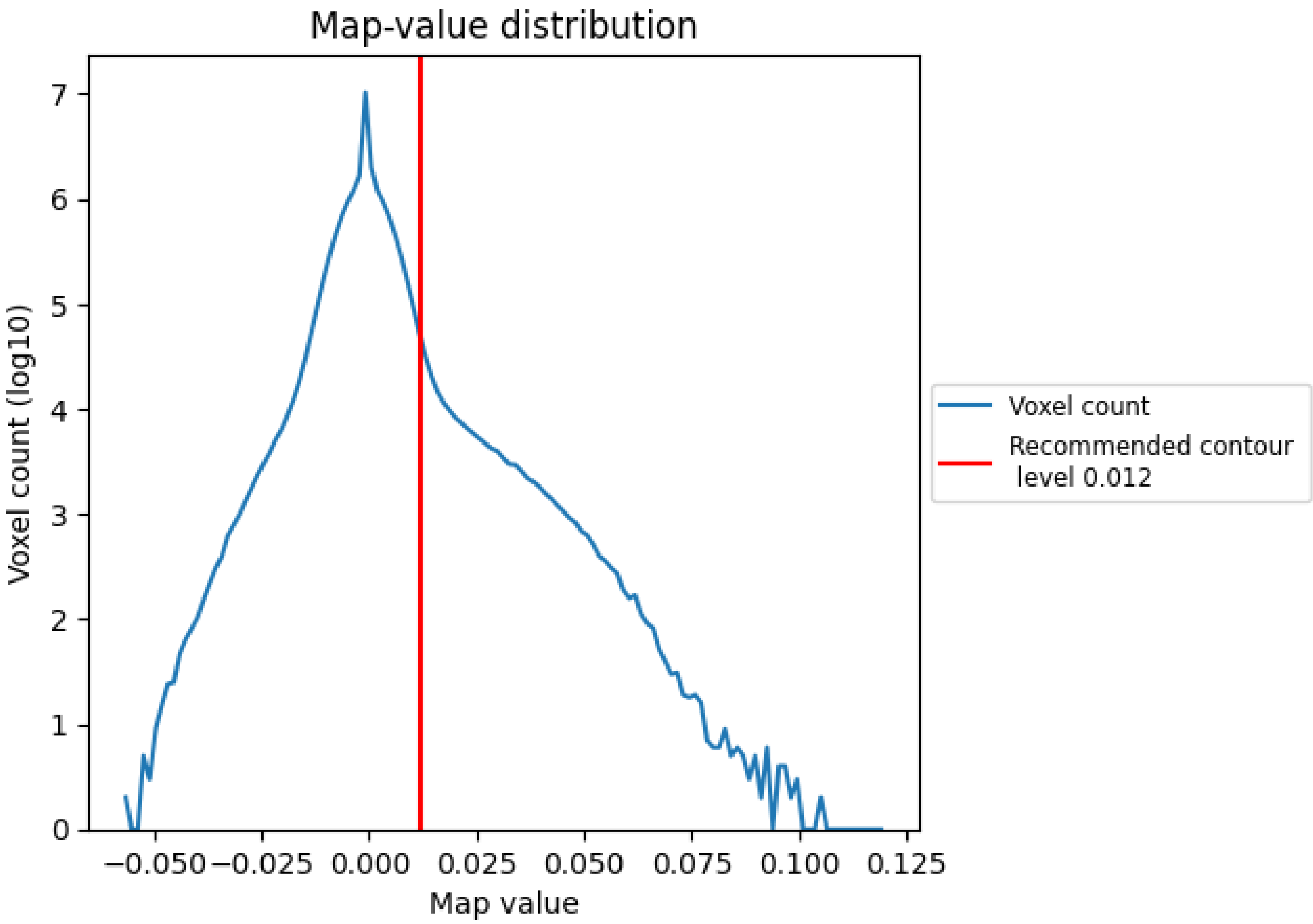


The map-value distribution is plotted in 128 intervals along the x-axis. The y-axis is logarithmic. A spike in this graph at zero usually indicates that the volume has been masked.

For Manuscript Review

#### 7.2 Volume estimate (i)

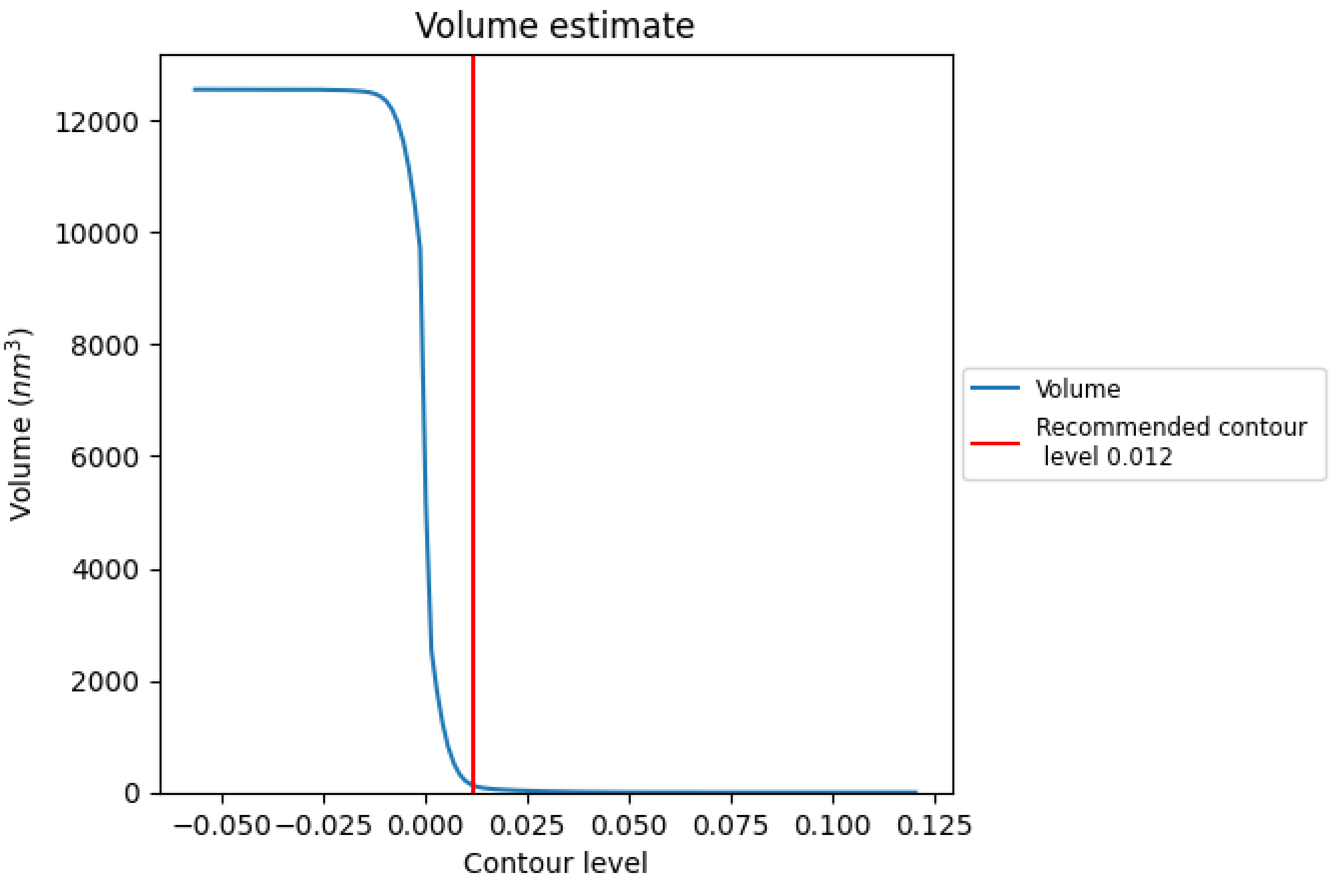


The volume at the recommended contour level is 120 $nm^3$; this corresponds to an approximate mass of 108 kDa.

The volume estimate graph shows how the enclosed volume varies with the contour level. The recommended contour level is shown as a vertical line and the intersection between the line and the curve gives the volume of the enclosed surface at the given level.

For Manuscript Review

#### 7.3 Rotationally averaged power spectrum (i)

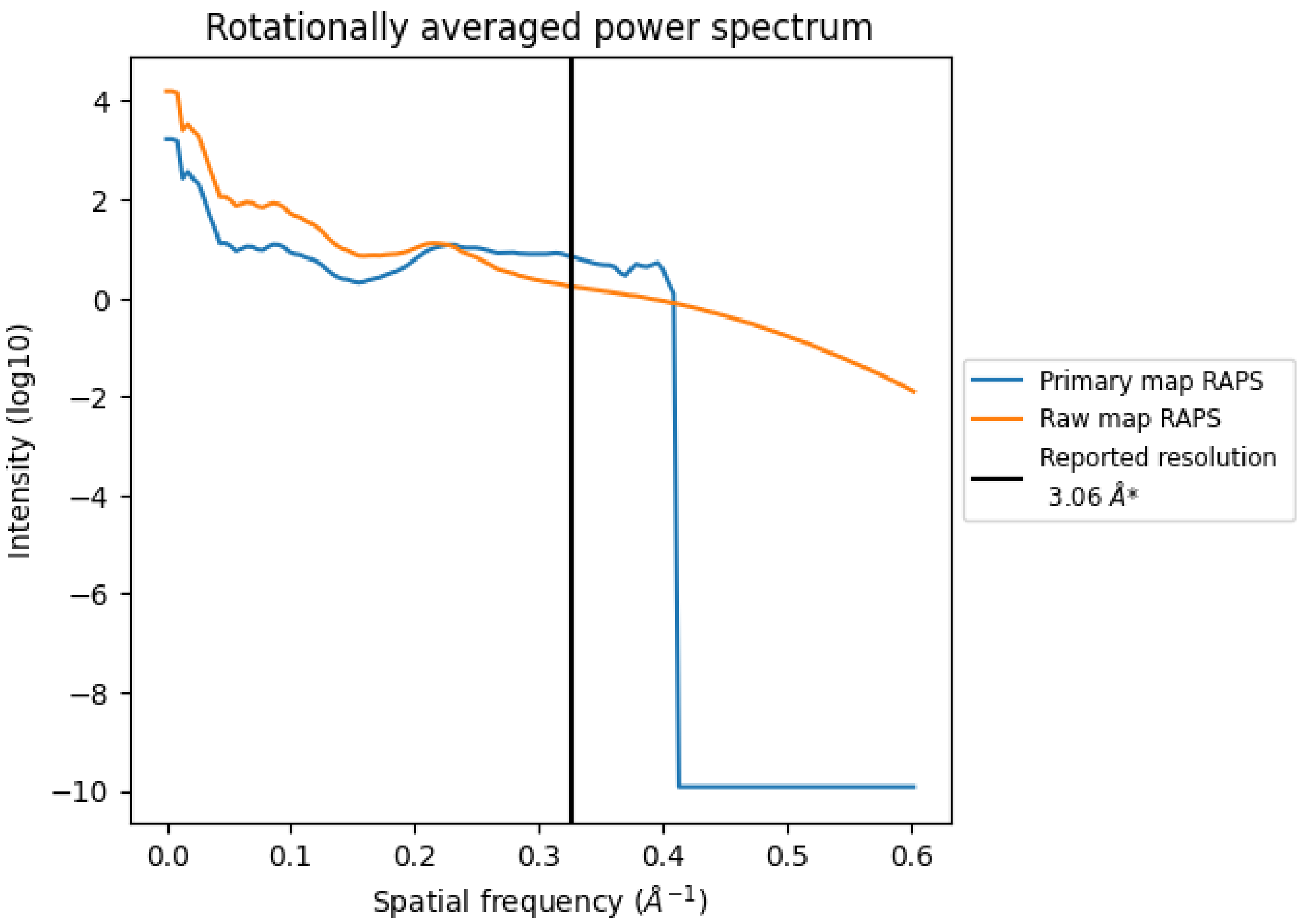


*Reported resolution corresponds to spatial frequency of 0.327 Å$^{-1}$

For Manuscript Review

### 8 Fourier-Shell correlation (i)

Fourier-Shell Correlation (FSC) is the most commonly used method to estimate the resolution of single-particle and subtomogram-averaged maps. The shape of the curve depends on the imposed symmetry, mask and whether or not the two 3D reconstructions used were processed from a common reference. The reported resolution is shown as a black line. A curve is displayed for the half-bit criterion in addition to lines showing the 0.143 gold standard cut-off and 0.5 cut-off.

#### 8.1 FSC (i)

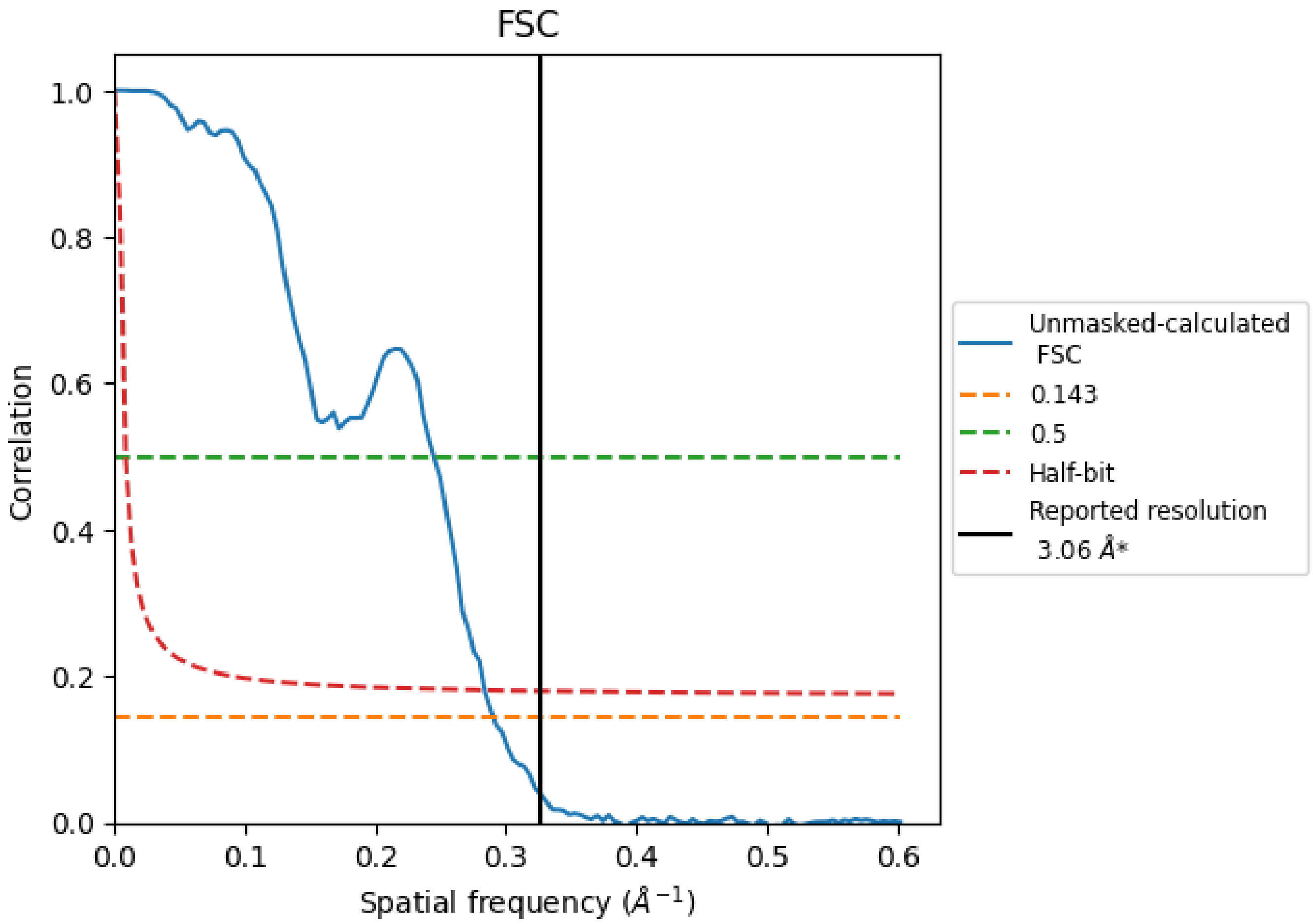


*Reported resolution corresponds to spatial frequency of 0.327 Å$^{-1}$

For Manuscript Review

#### 8.2 Resolution estimates (i)

| Resolution estimate (Å) | Estimation criterion (FSC cut-off) | | |
|---|---|---|---|
| | 0.143 | 0.5 | Half-bit |
| Reported by author | 3.06 | - | - |
| Author-provided FSC curve | - | - | - |
| Unmasked-calculated* | 3.44 | 4.09 | 3.52 |

*Resolution estimate based on FSC curve calculated by comparison of deposited half-maps. The value from deposited half-maps intersecting FSC 0.143 CUT-OFF 3.44 differs from the reported value 3.06 by more than 10 %

For Manuscript Review

### 9 Map-model fit (i)

This section contains information regarding the fit between EMDB map EMD-44634 and PDB model 9BJJ. Per-residue inclusion information can be found in section 3 on page 6.

#### 9.1 Map-model overlay (i)

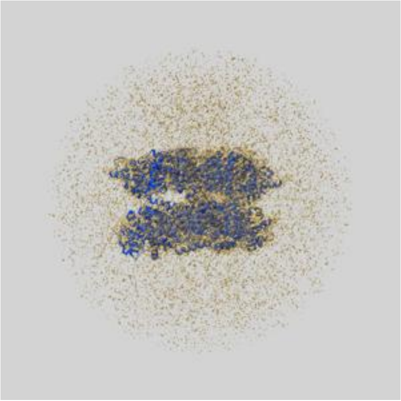

X

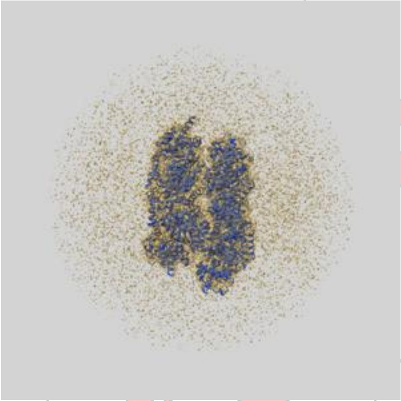

Y

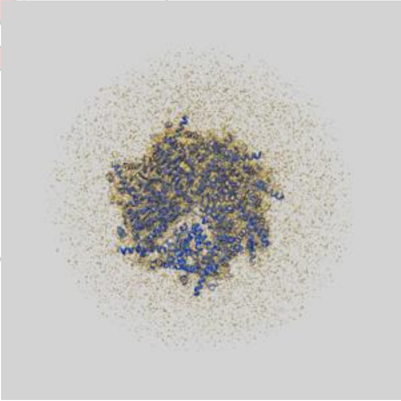

Z

The images above show the 3D surface view of the map at the recommended contour level 0.012 at 50% transparency in yellow overlaid with a ribbon representation of the model coloured in blue. These images allow for the visual assessment of the quality of fit between the atomic model and the map.

For Manuscript Review

#### 9.2 Q-score mapped to coordinate model (i)

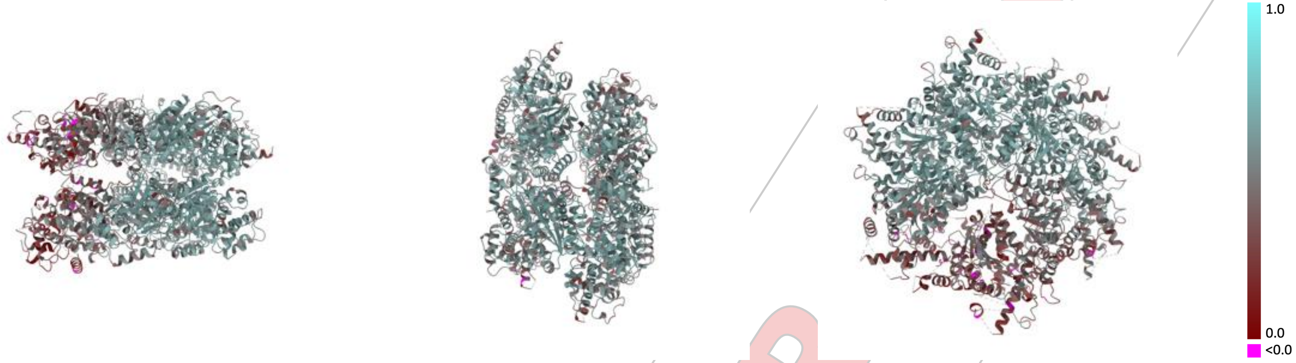


X Y Z

The images above show the model with each residue coloured according its Q-score. This shows their resolvability in the map with higher Q-score values reflecting better resolvability. Please note: Q-score is calculating the resolvability of atoms, and thus high values are only expected at resolutions at which atoms can be resolved. Low Q-score values may therefore be expected for many entries.

#### 9.3 Atom inclusion mapped to coordinate model (i)

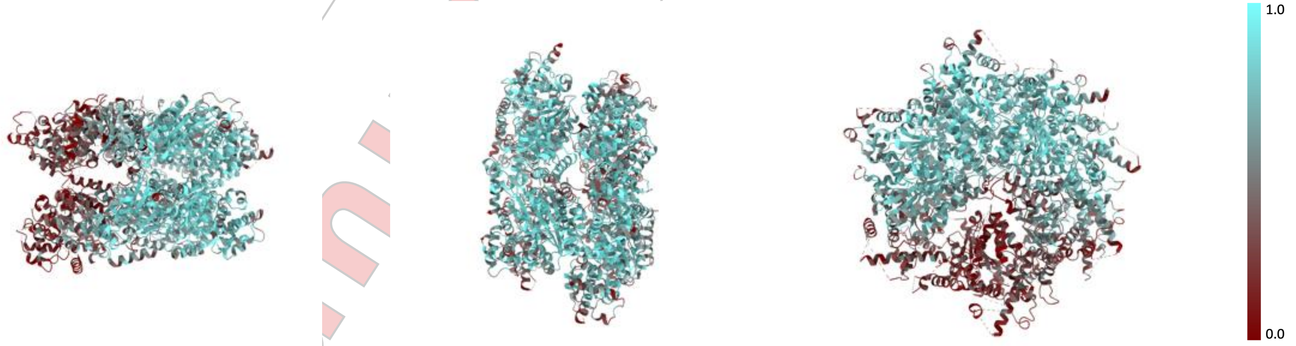


X Y Z

The images above show the model with each residue coloured according to its atom inclusion. This shows to what extent they are inside the map at the recommended contour level (0.012).

For Manuscript Review

#### 9.4 Atom inclusion (i)

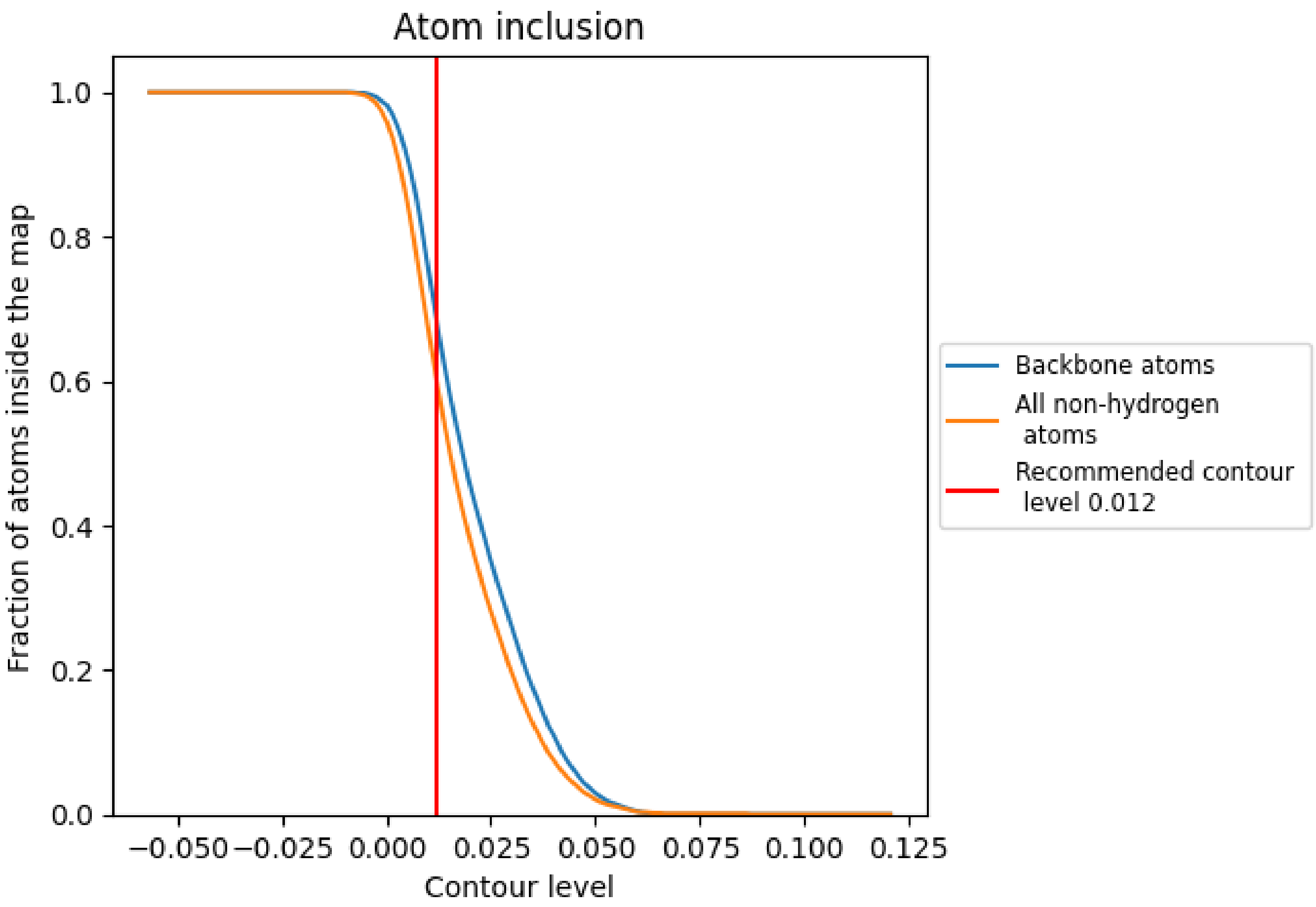


At the recommended contour level, 68% of all backbone atoms, 60% of all non-hydrogen atoms, are inside the map.

For Manuscript Review

#### 9.5 Map-model fit summary (i)

The table lists the average atom inclusion at the recommended contour level (0.012) and Q-score for the entire model and for each chain.

| Chain | Atom inclusion | Q-score |
|---|---|---|
| All | 0.6010 | 0.4950 |
| A | 0.6590 | 0.5260 |
| B | 0.7740 | 0.5780 |
| C | 0.7830 | 0.5780 |
| D | 0.7100 | 0.5470 |
| E | 0.3800 | 0.3820 |
| F | 0.2020 | 0.3080 |
| P | 0.4040 | 0.4400 |

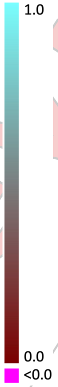


For Manuscript Review
