## Supplementary material for "A small molecule inhibitor of NVL suppresses tumor growth by blocking ribosome biogenesis": Data S5: Data S5_MM17_Treatment_Response.qmd

````
---
title: "MM017 Treatment Response Correlations"
format: 
  pdf: 
    warning: false
    cache: false
    toc: true
    author-meta: "Johann Peterson"
    title-meta: "MM017 Treatment Response Correlations"
    number-sections: true
    fig-format: png
  html:
    code-fold: true
    self-contained: true
execute:
  output-width: 60
knitr:
  opts_chunk:
    fig.path: figures/
    paged.print: false
include-in-header:
  text: |
   \usepackage[auth-sc]{authblk}
editor: visual
---

# Preliminaries

```{r}
#| label: time_it_hook
#| eval: true
#| echo: false
#| output: false
# https://bookdown.org/yihui/rmarkdown-cookbook/time-chunk.html#time-chunk

knitr::knit_hooks$set(time_it = local({
  now <- NULL
  function(before, options) {
    if (before) {
      # record the current time before each chunk
      now <<- Sys.time()
    } else {
      # calculate the time difference after a chunk
      res <- difftime(Sys.time(), now, units = "secs")
      # return a character string to show the time
      paste("Time for this code chunk to run:", round(res,
        2), "seconds")
    }
  }
}))
```

```{r}
#| eval: true
#| echo: false
#| output: false

options(width = 60)

filename <- knitr::current_input()
filename_no_ext <- xfun::with_ext(filename, "qmd")
```

Source file: `{r} filename_no_ext`

Run date: `{r} format(Sys.time(), '%d %B %Y')`

## Required paackages

```{r}
#| label: load_packages
#| eval: true
#| echo: false
#| output: false

require(tidyverse)
# require(tidyr)
# require(dplyr)
# require(ggplot2)
require(ggfortify)
require(ggrepel)
require(GGally)
require(reshape2)
require(stringr)
require(moments)
require(wPerm)
library(xlsx)
```

## File Names

-   [secondary-screen-replicate-collapsed-logfold-change.csv](https://depmap.org/portal/data_page/?tab=allData&releasename=PRISM%20Repurposing%2020Q2&filename=secondary-screen-replicate-collapsed-logfold-change.csv)
-   [secondary-screen-replicate-collapsed-treatment-info.csv](https://depmap.org/portal/data_page/?tab=allData&releasename=PRISM%20Repurposing%2020Q2&filename=secondary-screen-replicate-collapsed-treatment-info.csv)

```{r}
#| label: file_names

data_directory <- "data"
output_directory <- "output"

input_files = list(
    lfc = "secondary-screen-replicate-collapsed-logfold-change.csv",
    treatment = "secondary-screen-replicate-collapsed-treatment-info.csv",
    MM017 = "MTS019_DEEPAK_NIJHAWAN_LFC_1_3_10_uM_MATRIX_hg.csv",
    lfc_matrix_long_file = "lfc_matrix_long.rds"
  )

output_files = list(
    compounds_of_interest_file = "LFC_Compounds_of_Interest.xlsx",
    correlations_table = "LFC_correlations.xlsx"
)
```

```{r}
#| label: prepend_directories

for (fname in names(input_files)) {
  input_files[fname] = file.path(
    data_directory,
    input_files[fname]
  )
}

for (fname in names(output_files)) {
  output_files[fname] = file.path(
    output_directory,
    output_files[fname]
  )
}
```

## Constants

```{r}
#| label: constants

MM017_broad_id <- "BRD-U00103208"
MM017_name <- "MM017"
MM017_dose <- 10.0
MM017_treatment_id <- paste(
  MM017_broad_id, 
  ".", 
  format(MM017_dose, nsmall=1), 
  sep='')

ribosome_drugs <- c(
  "carmofur",
  "5-fluorouracil",
  "CX-5461",
  "oxaliplatin"
)
```

# LFC Analysis

## Load response data for MM017

```{r}
#| label: read_mm017_table

MM017_table <- read.csv(input_files$MM017)
MM017_table <- MM017_table %>% 
  rename_with( ~ gsub("X","LFC_",.x)) %>%
  rename_with( ~ gsub("\\.uM","_uM",.x)) %>%
  select(one_of(c("DepMap_ID","LFC_10.0_uM"))) %>%
  rename(LFC=LFC_10.0_uM) %>%
  mutate(treatment=MM017_treatment_id)
```

## Load Broad treatment data

```{r}
#| label: read_treatments_table

treatments <- read.csv(input_files$treatment, header=TRUE)
treatments <- treatments %>% 
  select(one_of(c(
    "column_name",
    "broad_id",
    "name",
    "dose",
    "screen_id",
    "moa"
    ))) %>%
  rename(treatment=column_name)
  
treatments$treatment <- treatments$treatment %>%
  str_replace_all("[:-]",".")

treatments <- treatments %>%
  add_row(data.frame(treatment = MM017_treatment_id,
                     broad_id = MM017_broad_id,
                     dose = MM017_dose,
                     name = MM017_name))
```

The order of levels here determines the order of preference for choosing a treatment when a single drug is present in multiple screens. I.e. `MTS010` is the first choice if present.

```{r}
treatments$screen_id <- factor(
  treatments$screen_id,
  ordered=TRUE,
  levels=c("MTS010", "MTS006", "MTS005", "HTS002"))
```

### Find drug names with more than 1 treatment

```{r}
treatments %>% 
  group_by(name) %>% 
  summarize(n=n()) %>% 
  arrange(desc(n), .by_group = TRUE) %>% 
  filter(n > 1)
```

```{r}
treatments %>% 
  group_by(name) %>% 
  arrange(screen_id, desc(dose)) %>% 
  slice_head(n=1) %>% 
  summarize(n=n()) %>% 
  arrange(desc(n))
```

```{r}
treatments_1_per_drug <- treatments %>% 
  group_by(name) %>% 
  arrange(screen_id, desc(dose)) %>% 
  slice_head(n=1)
```

## Load Broad cell line response data

```{r}
#| label: read_lfc_matrix

if(file.exists(input_files$lfc_matrix_long_file)) {
  lfc_matrix_long <- readRDS(input_files$lfc_matrix_long_file)
} else {
  lfc_matrix <- read.csv(input_files$lfc, header=TRUE)
  lfc_matrix <- lfc_matrix %>% rename(DepMap_ID=X)
  lfc_matrix_long <- lfc_matrix %>% 
    filter(!str_detect(DepMap_ID, "FAILED")) %>%
    pivot_longer(
      !DepMap_ID, 
      names_to=c("treatment"), 
      values_to=c("LFC")) %>%
    bind_rows(MM017_table)
  saveRDS(lfc_matrix_long, input_files$lfc_matrix_long_file)
}
```

```{r}
lfc_matrix_wide <- lfc_matrix_long %>% 
  pivot_wider(names_from=DepMap_ID, values_from=LFC)
```

```{r}
lfc_matrix_1_per_drug <- treatments_1_per_drug %>%
  left_join(lfc_matrix_wide, by="treatment")
```

```{r}
lfc_matrix_data_only <- lfc_matrix_1_per_drug %>% 
  drop_na(name) %>% 
  column_to_rownames(var="name") %>% 
  select(starts_with("ACH"))

lfc_matrix_data_only <- 
  lfc_matrix_data_only[, !is.na(lfc_matrix_data_only[MM017_name, ]) ]
```

## Examine the `NA` values

in `lfx_matrix_data_only`

```{r}
#| label: NA_counts_per_row

na_per_row <- apply(lfc_matrix_data_only, 1, function(x) sum(is.na(x)))
hist(na_per_row,  main="NA Counts Per Row", xlab="# of NAs")
```

```{r}
#| label: NA_counts_per_col

na_per_col <- apply(lfc_matrix_data_only, 2, function(x) sum(is.na(x)))
hist(na_per_col,  main="NA Counts Per Column", xlab="# of NAs")
```

```{r}
lfc_temp <- lfc_matrix_data_only[,!(20 < na_per_col)]
na_per_row <- apply(lfc_temp, 1, function(x) sum(is.na(x)))
lfc_temp <- lfc_temp[!(20 < na_per_row),]
```

## Tabulate LFCs for compounds of interest

```{r}
#| label: compounds_of_interest

compounds <- c("MM017", ribosome_drugs)

lfc_matrix_1_per_drug %>% 
  filter(name %in% compounds) %>% 
  select(name, dose, screen_id, broad_id)

lfc_compounds_of_interest <- t(
  lfc_matrix_data_only[compounds,]
)

write.xlsx(lfc_compounds_of_interest,
           file = output_files$compounds_of_interest_file,
           sheetName = "LFC",
           col.names = TRUE,
           row.names = TRUE)
```

```{r}
#| label: correlation_grid_plot

ggpairs(lfc_compounds_of_interest
        # mapping = aes(point_size=0.0005)
        )
```

```{r}
df <- as.data.frame(
  treatments_1_per_drug[
  treatments_1_per_drug$name %in% compounds,
  c("name","dose","treatment")]
)

df

write.xlsx(
  df,
  file = output_files$compounds_of_interest_file,
  sheetName = "treatments",
  col.names = TRUE,
  row.names = TRUE,
  append = TRUE)
```

## Correlations

### 5-fluorouracil

```{r}
#| label: correlation_5fu

cor.test(
  lfc_compounds_of_interest[,"MM017"],
  lfc_compounds_of_interest[,"5-fluorouracil"]
)
```

### CX-5461

```{r}
#| label: correlation_cx

cor.test(
  lfc_compounds_of_interest[,"MM017"],
  lfc_compounds_of_interest[,"CX-5461"]
)
```

### oxaliplatin

```{r}
#| label: correlation_oxaliplatin

cor.test(
  lfc_compounds_of_interest[,"MM017"],
  lfc_compounds_of_interest[,"oxaliplatin"]
)
```

### carmofur

```{r}
#| label: correlation_carmofur

cor.test(
  lfc_compounds_of_interest[,"MM017"],
  lfc_compounds_of_interest[,"carmofur"]
)
```

# Full Correlation Table

```{r}
cor_list <- apply(
        lfc_matrix_data_only,
        1,
        function(r) 1 - cor(
          as.numeric(r), 
          as.numeric(lfc_matrix_data_only[MM017_name,]),
          use = "pairwise.complete.obs")
)
correlations_data_frame <- as.data.frame(cor_list)
correlations_data_frame$compound <- row.names(correlations_data_frame)
names(correlations_data_frame) <- c("one.minus.corr","compound")
```

## P-Values by Permutation Test

```{r}
#| label: cor_permutation_test
#| time_it: true

cor_permutation_list <- apply(
  lfc_matrix_data_only,
  1,
  function(r) {
    keep <- !is.na(r)
    pr <- perm.relation(
      as.numeric(r[keep]), 
      as.numeric(lfc_matrix_data_only[MM017_name,keep])
    )
    return( list(r=pr$Observed, p=pr$p.value))
  }
)
```

```{r}
#| message: true
#| time_it: true
cor_permutation_results <- do.call(
  rbind, 
  lapply(cor_permutation_list, as.data.frame))
cor_permutation_results <- rownames_to_column(
  cor_permutation_results, 
  var = "drug")
```

## Benjamini-Hochberg Correction

```{r}
cor_permutation_results <- cor_permutation_results %>% 
  mutate(fdr = p.adjust(p, method = "BH")) %>%
  drop_na()
```

```{r}
#| label: correlation_volcano_plot

ggplot(data = cor_permutation_results, aes(x=r, y=-log10(p))) + 
  geom_point(size=0.5) +
  geom_point(
    size=2, 
    color="red", 
    data = 
      cor_permutation_results[
        cor_permutation_results$drug %in% ribosome_drugs,]) +
  geom_text_repel(
    aes(label=drug, color="red"), 
    segment.color = "grey50",
    data=cor_permutation_results[
      cor_permutation_results$drug %in% ribosome_drugs,],
    show.legend = FALSE,
    min.segment.length = 0,
    hjust=0,
    # xlim=c(0.5, 1.0),
    nudge_x = 0.75 - 
      cor_permutation_results[
        cor_permutation_results$drug %in% ribosome_drugs,]$r,
    nudge_y = -0.75 + 
      cor_permutation_results[
        cor_permutation_results$drug %in% ribosome_drugs,]$r,
    direction = "y",
    max.overlaps = 10,
    point.padding = 0.1,
    box.padding = 1) +
  xlim(-0.4, 1.1)
```

```{r}
#| label: correlation_histogram

labels <- cor_permutation_results[
  cor_permutation_results$drug %in% compounds,]
labels$x <- 1 - labels$r

ggplot(data = cor_permutation_results, aes(x = 1-r)) +
  geom_histogram(fill = "lightblue", color = "black", binwidth=0.02) +
  annotate("segment", x = labels$x, xend = labels$x, 
           y = 5, yend = 0, color = "red", 
           arrow = arrow(length = unit(0.2, "cm"))) +
  annotate("text", 
           x = labels$x, 
           y = 10, 
           label = labels$drug, 
           color = "red", size = 4, angle = 90, hjust = 0) +
  labs(title = "Correlations with MM017",
       x = "1 - correlation",
       y = "count")
  ylim(-50,100)
```

```{r}
write.xlsx(cor_permutation_results, 
          output_files$correlations_table,
          col.names = TRUE,
          row.names = FALSE)
```
````
