## Supplementary material for "A small molecule inhibitor of NVL suppresses tumor growth by blocking ribosome biogenesis": Data S6: gammaH2AX.ijm

```
run("Set Measurements...", "area mean display redirect=None decimal=3");
dir=getDirectory("Image Directory");
images = getFileList(dir);
setBatchMode(true);

for (a = 0; a < images.length; a++) {
	if (endsWith(images[a], ".nd2")==1){
	run("Bio-Formats Importer", "open="+dir+images[a]+" autoscale color_mode=Composite rois_import=[ROI manager] view=Hyperstack stack_order=XYCZT"); //opens the file
	id = getImageID();
	if (isOpen("Quantification")==0) {Table.create("Quantification");}
	selectImage(id);
	title = getTitle();
	titleS = File.nameWithoutExtension;
	run("Split Channels");
	selectImage("C1-"+title);
	idRed = getImageID();
	selectImage("C2-"+title);
	idDapi = getImageID();
	run("Duplicate...", "title=copy");
	run("Pseudo flat field correction", "blurring=250 hide");
	run("Subtract...", "value=300");
	run("Gaussian Blur...", "sigma=2");
	setAutoThreshold("Huang dark");
	setOption("BlackBackground", true);
	run("Convert to Mask");
	run("Fill Holes");
	run("Adjustable Watershed", "tolerance=2");
	run("Analyze Particles...", "size=50-Infinity add composite");
	selectImage(idRed);
	
		n = roiManager("count");
		for (i = 0; i < n; i++) {
		run("Clear Results");
		roiManager("select", i);
		roiManager("Measure");
		area = getResult("Area", 0);
		meanInt = getResult("Mean", 0);
		run("Clear Results");
	    run("Duplicate...", "title=ROI");
	    run("Find Maxima...", "prominence=600 output=[Point Selection]");
	    run("Measure");
	    maxima = nResults;
	    run("Clear Results");
	    selectWindow("Quantification");
		k = Table.size;
		Table.set("image", k, titleS);
		Table.set("Nucleus", k, i+1);
		Table.set("Area", k, area);
		Table.set("# of spots", k, maxima);
		Table.set("MeanIntensity", k, meanInt);
		Table.update;
	    close();
		}
		
	roiManager("Save", dir+titleS+"_ROIs.zip");
	roiManager("reset");

close("*");
	}
	}

selectWindow("Quantification");
saveAs("Results", dir+"Quantification.csv");
close("Quantification.csv");
if(isOpen("Results"));{close("Results");
showMessage("Done!", "<html>"+"<h1><font color=black>Done!</h1>");
```
