## Supplementary material for "A small molecule inhibitor of NVL suppresses tumor growth by blocking ribosome biogenesis": Data S6: p53_Ki67_quantitation.groovy

```
clearAllObjects();
createFullImageAnnotation(true)
// Specify the model name (cyto, nuclei, cyto2, ... or a path to your custom model as a string)
// Other models for Cellpose https://cellpose.readthedocs.io/en/latest/models.html
// And for Omnipose: https://omnipose.readthedocs.io/models.html
def pathModel = 'cyto3'
def cellpose = Cellpose2D.builder( pathModel )
        .pixelSize( 0.2 )                  // Resolution for detection in um
        .channels( 2 )	               // Select detection channel(s)
//        .tempDirectory( new File( '/tmp' ) ) // Temporary directory to export images to. defaults to 'cellpose-temp' inside the QuPath Project
//        .preprocess( ImageOps.Filters.median( 1 ) )                // List of preprocessing ImageOps to run on the images before exporting them
//        .normalizePercentilesGlobal( 0.1, 99.8, 10 ) // Convenience global percentile normalization. arguments are percentileMin, percentileMax, dowsample.
//        .tileSize( 1024 )                  // If your GPU can take it, make larger tiles to process fewer of them. Useful for Omnipose
//        .cellposeChannels( 2 )           // Overwrites the logic of this plugin with these two values. These will be sent directly to --chan and --chan2
//        .cellprobThreshold( 0.0 )          // Threshold for the mask detection, defaults to 0.0
//        .flowThreshold( 0.4 )              // Threshold for the flows, defaults to 0.4
//        .diameter( 5 )                    // Median object diameter. Set to 0.0 for the `bact_omni` model or for automatic computation
//        .useOmnipose()                   // Use omnipose instead
//        .addParameter( "cluster" )         // Any parameter from cellpose or omnipose not available in the builder.
//        .addParameter( "save_flows" )      // Any parameter from cellpose or omnipose not available in the builder.
//        .addParameter( "anisotropy", "3" ) // Any parameter from cellpose or omnipose not available in the builder.
//        .cellExpansion( 10.0 )              // Approximate cells based upon nucleus expansion
//        .cellConstrainScale( 1.5 )         // Constrain cell expansion using nucleus size
//        .classify( "My Detections" )       // PathClass to give newly created objects
//        .measureShape()                  // Add shape measurements
//        .measureIntensity()              // Add cell measurements (in all compartments)
//        .createAnnotations()             // Make annotations instead of detections. This ignores cellExpansion
//        .simplify( 0 )                     // Simplification 1.6 by default, set to 0 to get the cellpose masks as precisely as possible
        .build()

def imageData = getCurrentImageData()
def pathObjects = getSelectedObjects() // To process only selected annotations, useful while testing
// def pathObjects = getAnnotationObjects() // To process all annotations. For working in batch mode
if (pathObjects.isEmpty()) {
    Dialogs.showErrorMessage( "Cellpose", "Please select a parent object!" )
    return
}
cellpose.detectObjects( imageData, pathObjects )
import qupath.ext.biop.cellpose.Cellpose2D
selectDetections();
runPlugin('qupath.lib.algorithms.IntensityFeaturesPlugin', '{"pixelSizeMicrons":0.11,"region":"ROI","tileSizeMicrons":25.0,"channel1":true,"channel2":true,"channel3":false,"doMean":true,"doStdDev":true,"doMinMax":true,"doMedian":true,"doHaralick":true,"haralickMin":0.0,"haralickMax":65535.0,"haralickDistance":1,"haralickBins":32}')
selectAnnotations();
runPlugin('qupath.lib.plugins.objects.SmoothFeaturesPlugin', '{"fwhmMicrons":25.0,"smoothWithinClasses":false}')
resetDetectionClassifications();
runObjectClassifier("Classifier");
```
