## Supplementary material for "A small molecule inhibitor of NVL suppresses tumor growth by blocking ribosome biogenesis": Data S6: ribosomeLocalization.ijm

```
// latest version: 2025-02-27

run("Set Measurements...", "area mean integrated display redirect=None decimal=0");
dir=getDirectory("Image Directory");
images = getFileList(dir);
setBatchMode(true);

for (a = 0; a < images.length; a++) {
	if (endsWith(images[a], ".nd2")==1){
	run("Bio-Formats Importer", "open="+dir+images[a]+" autoscale color_mode=Composite rois_import=[ROI manager] view=Hyperstack stack_order=XYCZT"); //opens the file
	roiManager("reset");
	id = getImageID();
	if (isOpen("Quantification")==0) {Table.create("Quantification");}
	selectImage(id);
	title = getTitle();
	titleS = File.nameWithoutExtension;
	run("Split Channels");
	selectImage("C1-"+title);
	idRed = getImageID();
	selectImage("C2-"+title);
	idGreen = getImageID();
	selectImage("C3-"+title);
	idDapi = getImageID();
	run("Duplicate...", "title=copy");
	run("Pseudo flat field correction", "blurring=250 hide");
	run("Gaussian Blur...", "sigma=2");
	setAutoThreshold("Huang dark");
	setOption("BlackBackground", true);
	run("Convert to Mask");
	run("Fill Holes");
	run("Adjustable Watershed", "tolerance=2");
	roiManager("reset");
	run("Analyze Particles...", "size=50-Infinity add composite");
	roiManager("Save", dir+titleS+"_nuclei_ROIs.zip");
	selectImage(idRed); ////////////////////
	roiManager("Deselect");
	run("Clear Results");
	roiManager("Measure");		
	selectWindow("Results");
	n=nResults;
	nuclei = n;
	run("Summarize");
	NucArea = getResult("Area", n);
	meanMagentaNuc = getResult("Mean", n);
	MagentaNucIntDen = getResult("IntDen", n);
	MagentaNucRawDen = getResult("RawIntDen", n);
	saveAs("Results", dir+titleS+"_Nuclear_Magenta_Quantification.csv");
	run("Clear Results");
selectImage(idGreen); ////////////////////
	roiManager("Deselect");
	run("Clear Results");
	roiManager("Measure");		
	selectWindow("Results");
	run("Summarize");
	meanGreenNuc = getResult("Mean", n);
	GreenNucIntDen = getResult("IntDen", n);
	GreenNucRawDen = getResult("RawIntDen", n);
	saveAs("Results", dir+titleS+"_Nuclear_Green_Quantification.csv");
	run("Clear Results");

selectImage(idGreen);
run("Duplicate...", " ");
run("Gaussian Blur...", "sigma=2");
setAutoThreshold("Huang dark");
setOption("BlackBackground", true);
run("Convert to Mask");
idDuplicateGreen = getImageID();
run("Options...", "iterations=5 count=4 black do=Close");
b = roiManager("count");
for (i = 0; i < b; i++) {
roiManager("select", i);
setBackgroundColor(0, 0, 0);
run("Clear", "slice");
	}
roiManager("reset");
roiManager("show all");
roiManager("show none");
selectImage(idDuplicateGreen);
setAutoThreshold("Huang dark");
roiManager("reset");
run("Analyze Particles...", "size=50-Infinity add composite");
roiManager("Save", dir+titleS+"_cytoplasm_ROIs.zip");
roiManager("Combine");
roiManager("Add");
roiManager("Deselect");
n = roiManager("count");
selectImage(idGreen);////////////////////
run("Clear Results");
roiManager("Select", n-1);
List.setMeasurements;
areacYTOGreen = List.getValue("Area");
meancYTOGreen = List.getValue("Mean");
GreencYTOIntDen = List.getValue("IntDen");
GreenCytoRawDen = List.getValue("RawIntDen");
run("Clear Results");
roiManager("Select", n-1);
roiManager("Measure");		
selectWindow("Results");
saveAs("Results", dir+titleS+"_Cytoplasm_Green_Quantification.csv");
run("Clear Results");
selectImage(idRed);////////////////////
roiManager("Select", n-1);
List.setMeasurements;
meancYTOMag = List.getValue("Mean");
MagcYTOIntDen = List.getValue("IntDen");
MagCytoRawDen = List.getValue("RawIntDen");
ratio = meanMagentaNuc/meancYTOMag;
run("Clear Results");
roiManager("Select", n-1);
roiManager("Measure");		
selectWindow("Results");
saveAs("Results", dir+titleS+"_Cytoplasm_Magenta_Quantification.csv");
run("Clear Results");
selectWindow("Quantification");

k = Table.size;
		Table.set("image", k, titleS);
		Table.set("Number of nuclei", k, nuclei);
		Table.set("Mean nuclear area", k, NucArea);
		Table.set("Total cytoplasmic area", k, areacYTOGreen);
		Table.set("Green mean intensity Nuclei", k, meanGreenNuc);
		Table.set("Magenta mean intensity Nuclei", k, meanMagentaNuc);
		Table.set("Green mean intensity cytopl", k, meancYTOGreen);
		Table.set("Magenta mean intensity cytopl", k, meancYTOMag);
		Table.set("Magenta ratio nucl/cytopl", k, ratio);

close("*");
	}
	}
roiManager("reset");
roiManager("show all");
roiManager("show none");
selectWindow("Quantification");
saveAs("Results", dir+"Quantification.csv");
close("Quantification.csv");

showMessage("Done!", "<html>"+"<h1><font color=black>Done!</h1>");
```
