## Supplementary material for "A small molecule inhibitor of NVL suppresses tumor growth by blocking ribosome biogenesis": Data S7

Full gel images shown in Fig. 3E (**A**) and Fig. 3F (**B**).

**A**

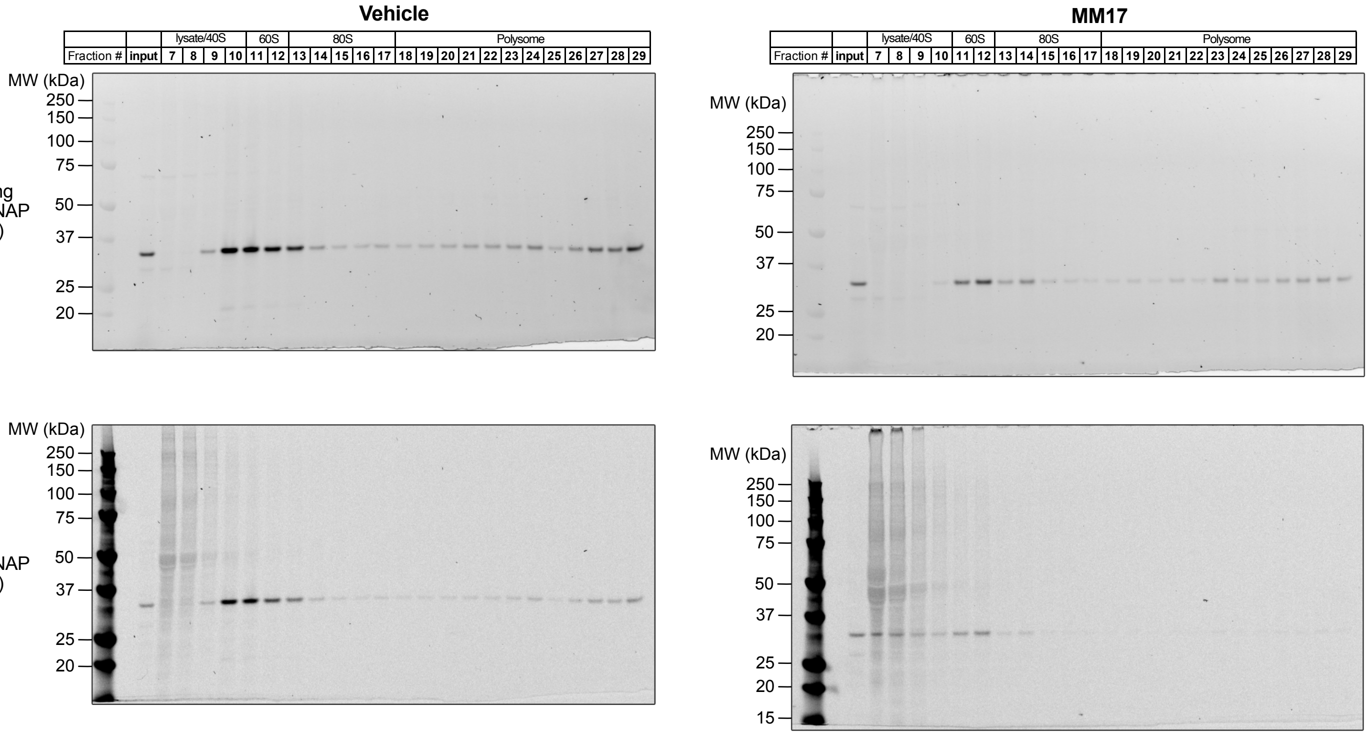


**B**

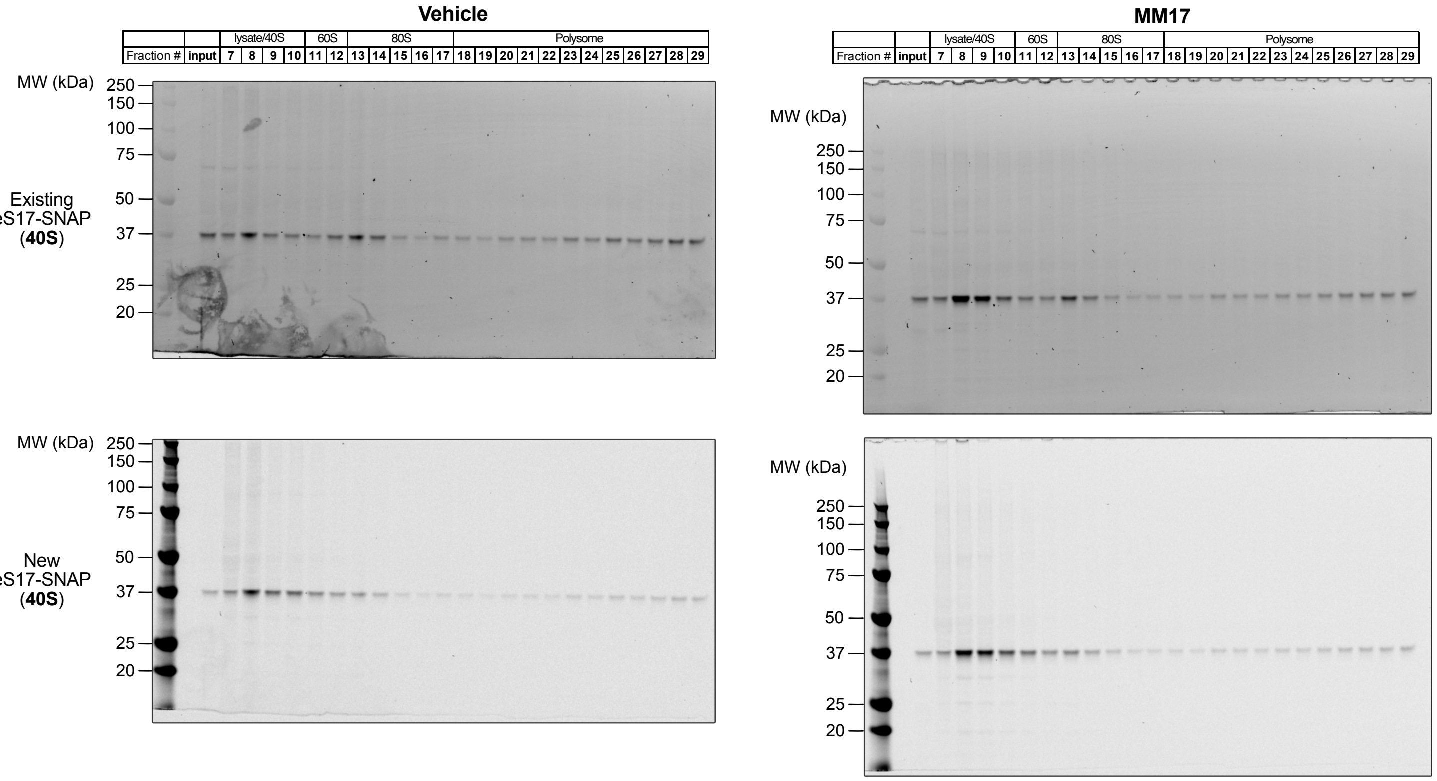

Full microscopy images shown in Fig. 3G and S13B (**A**), Fig. 3H and S13C (**B**), and Fig. 3I and S13D (**C**). Scale bar = 10 μm. White box indicates area used to create figure panels.

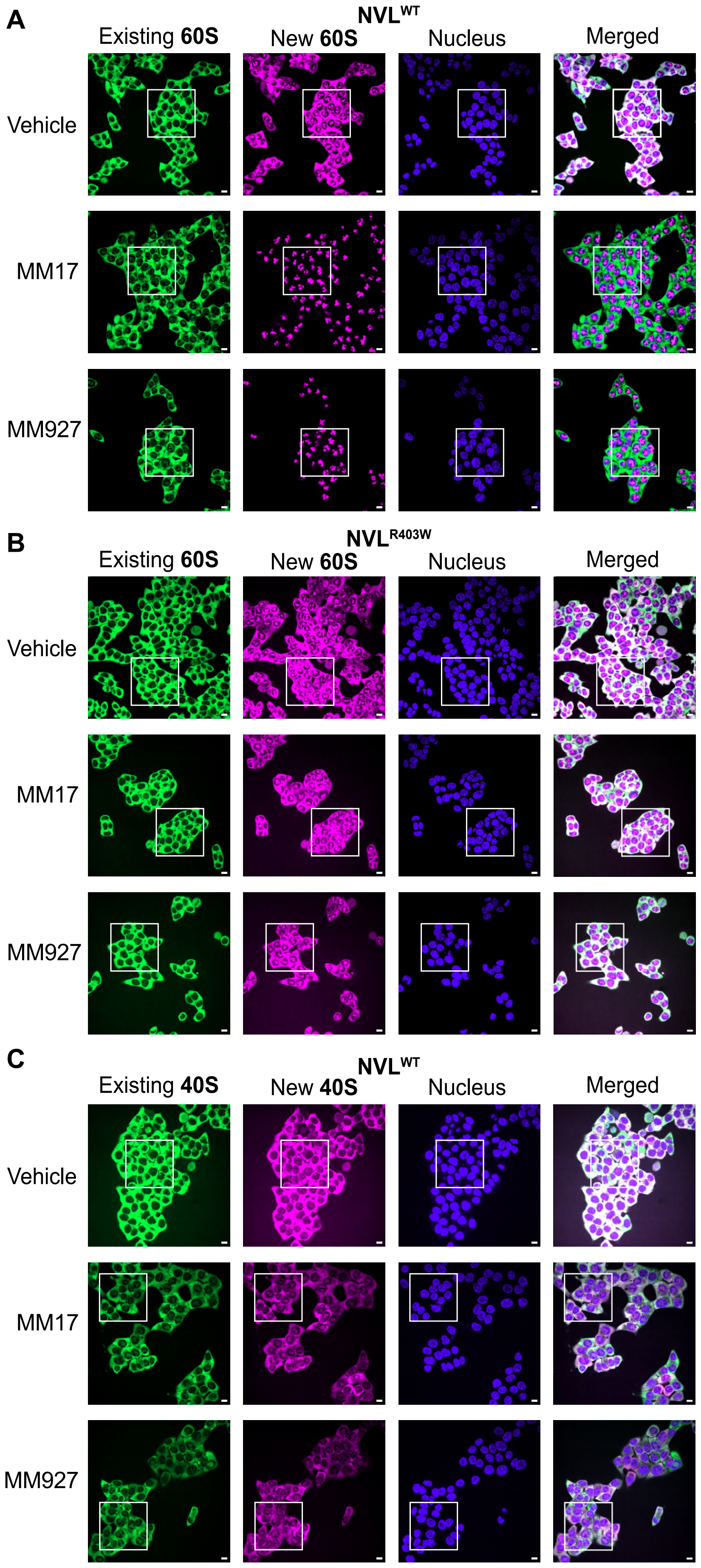

Full Western blot images shown in Fig. 4B.
HRP chemiluminescence images are used in the figure panels.

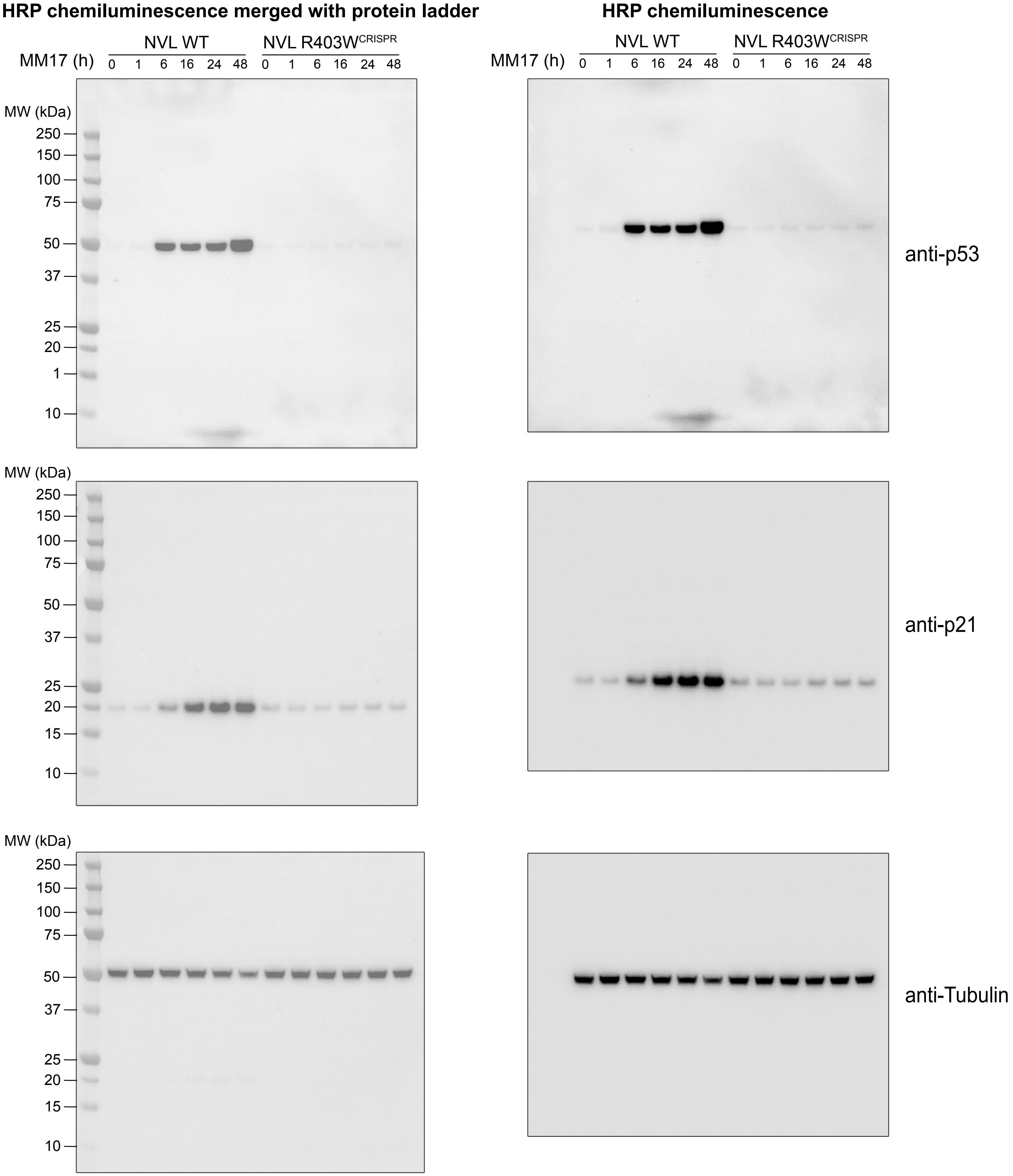

Full Western blot images shown in Fig. 4D.
HRP chemiluminescence images are used in the figure panels.

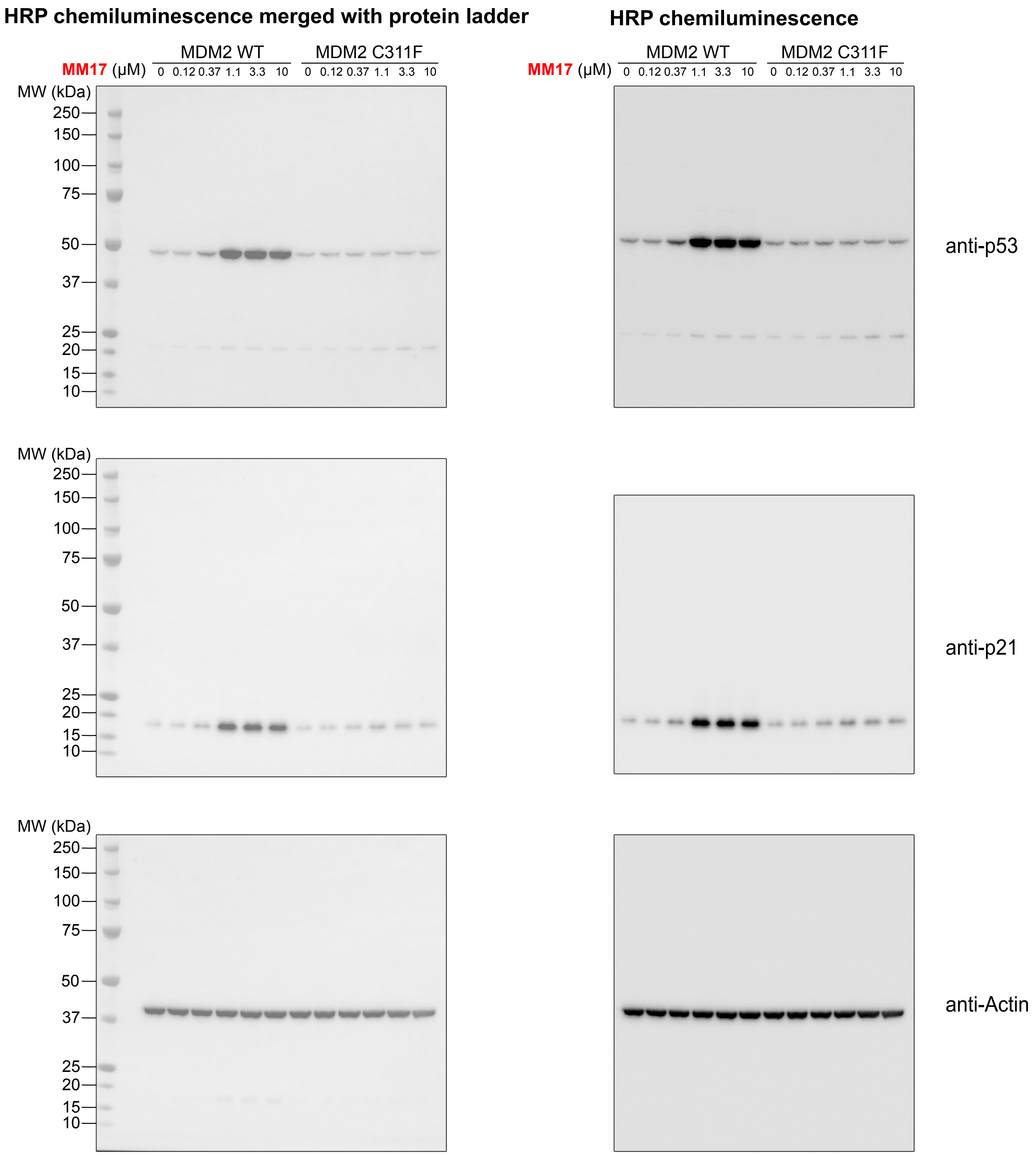

Full Western blot images shown in Fig. 4E.
HRP chemiluminescence images are used in the figure panels.

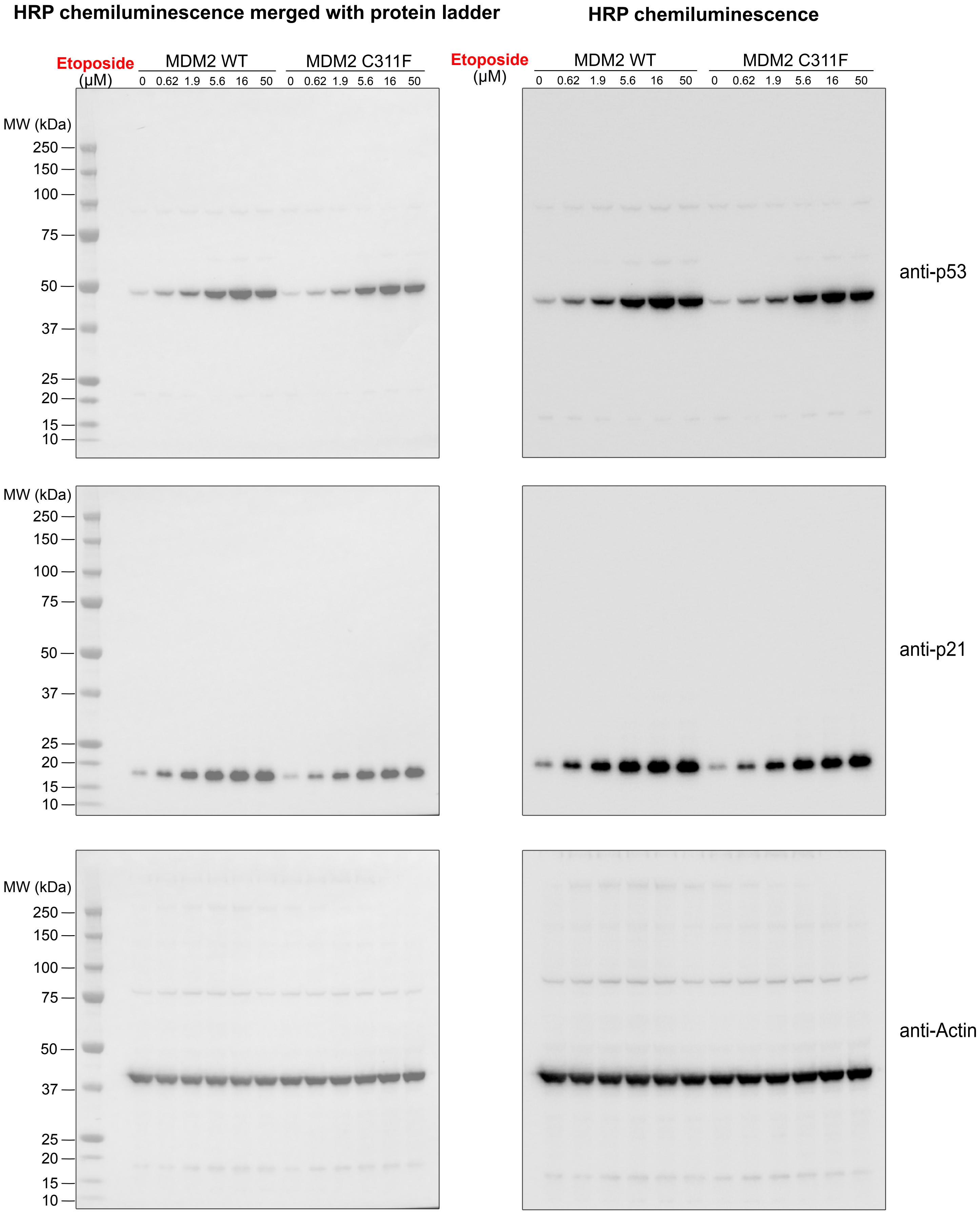

Full microscopy images shown in Fig. 5C.
Scale bar = 10 μm. White box indicates area used to create figure panels.

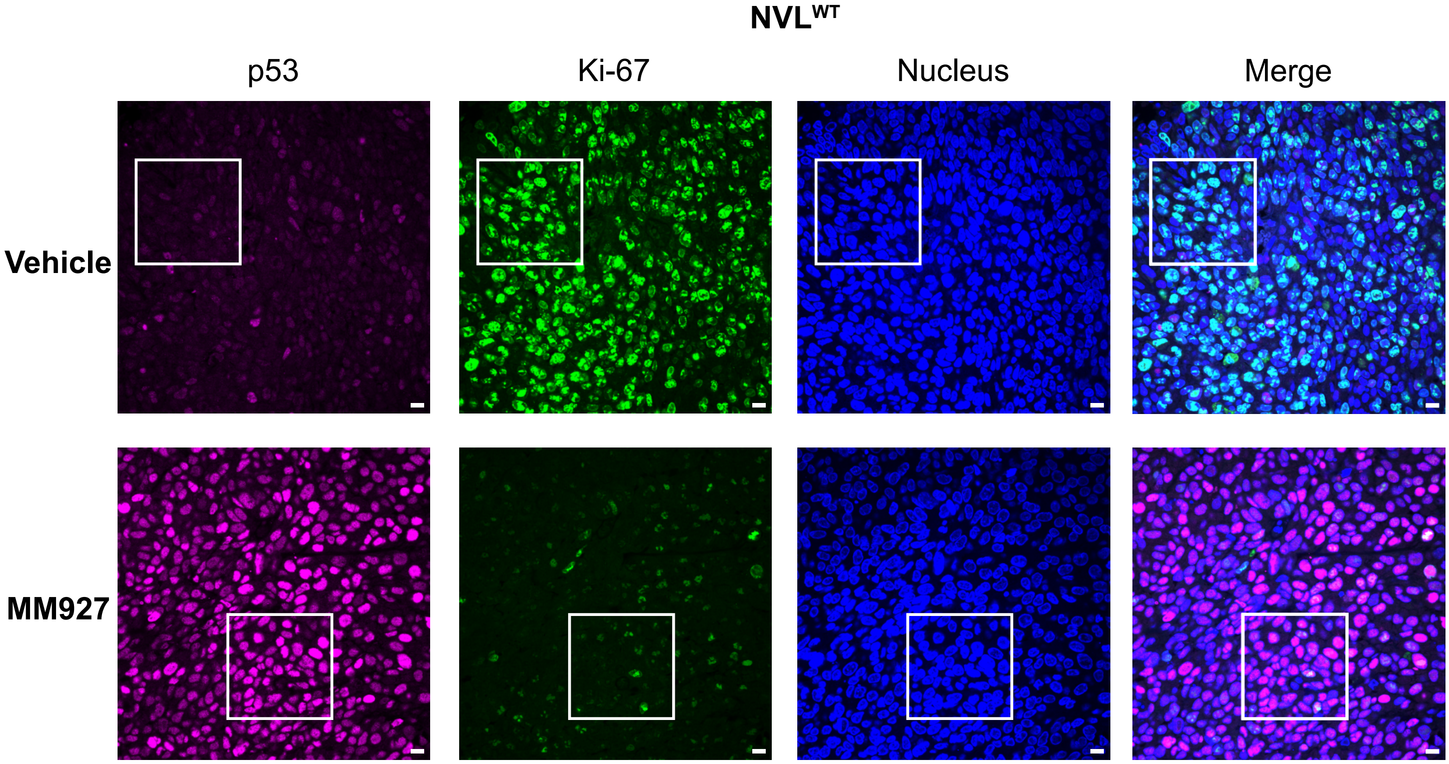


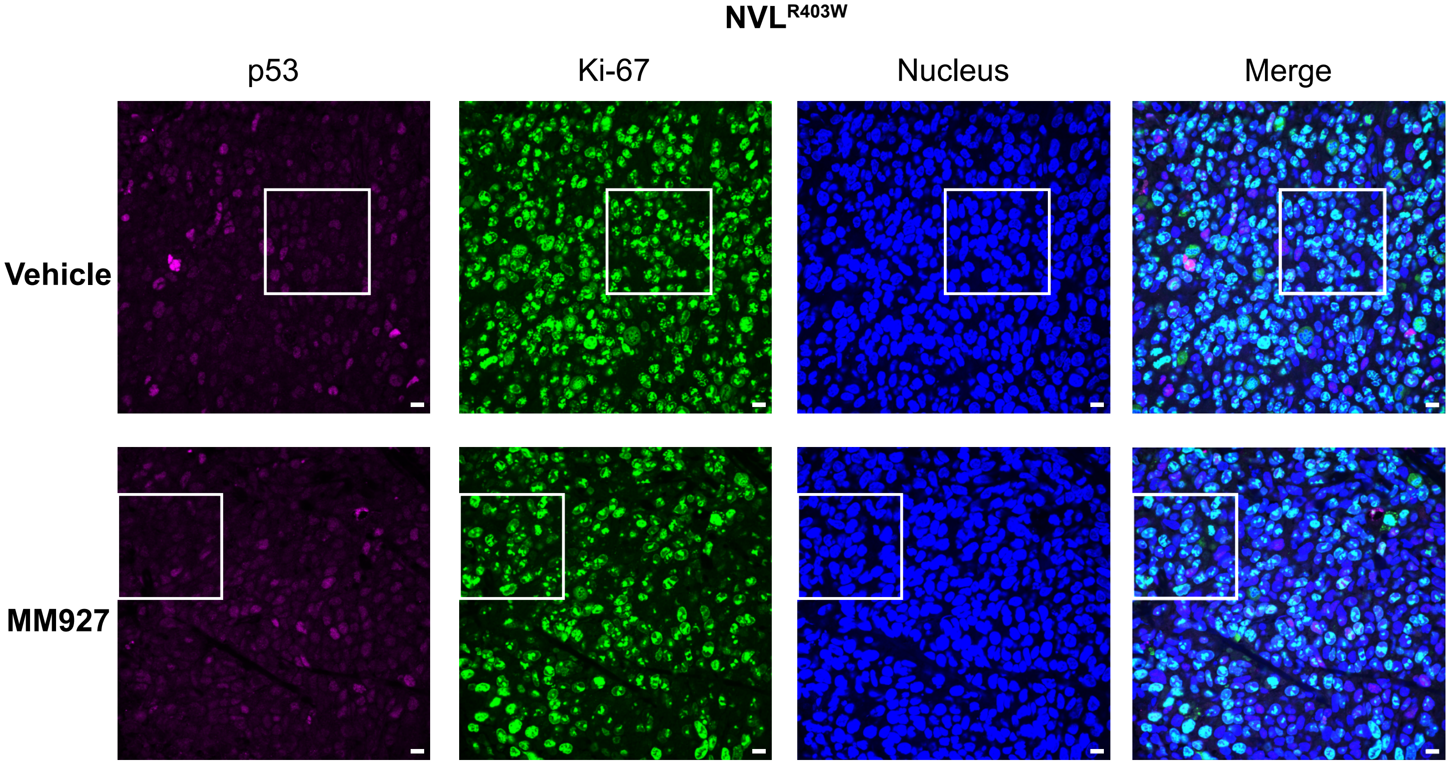

Full microscopy images shown in Fig. 6A.
Scale bar = 10 µm. White box indicates area used to create figure panels.

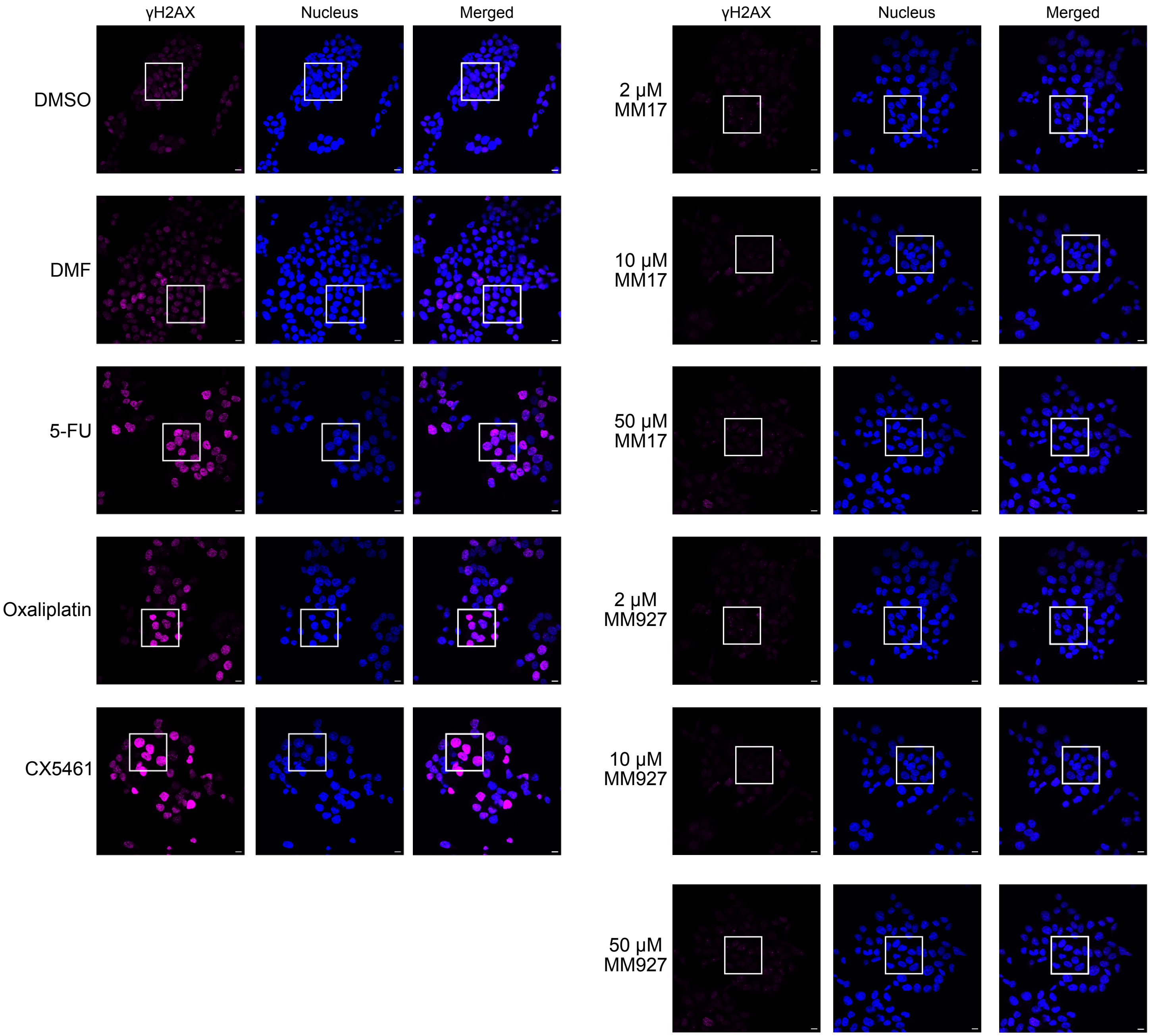

Full gel images shown in Fig. S2C (**A**), S2D (**B**), S2E (**C**).

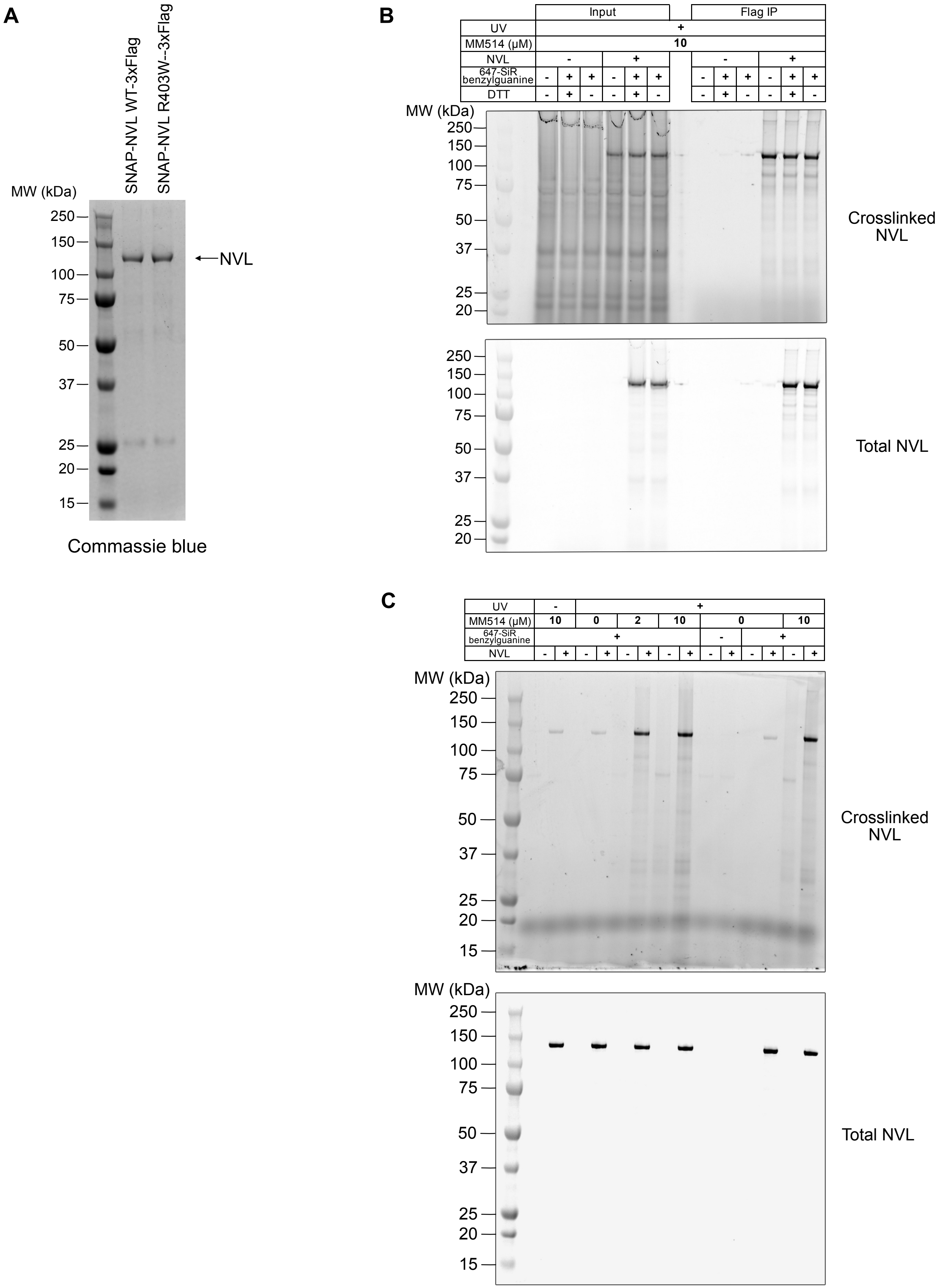

Full gel images shown in Fig. S2H (**A**) and Fig. S2I (**B**).

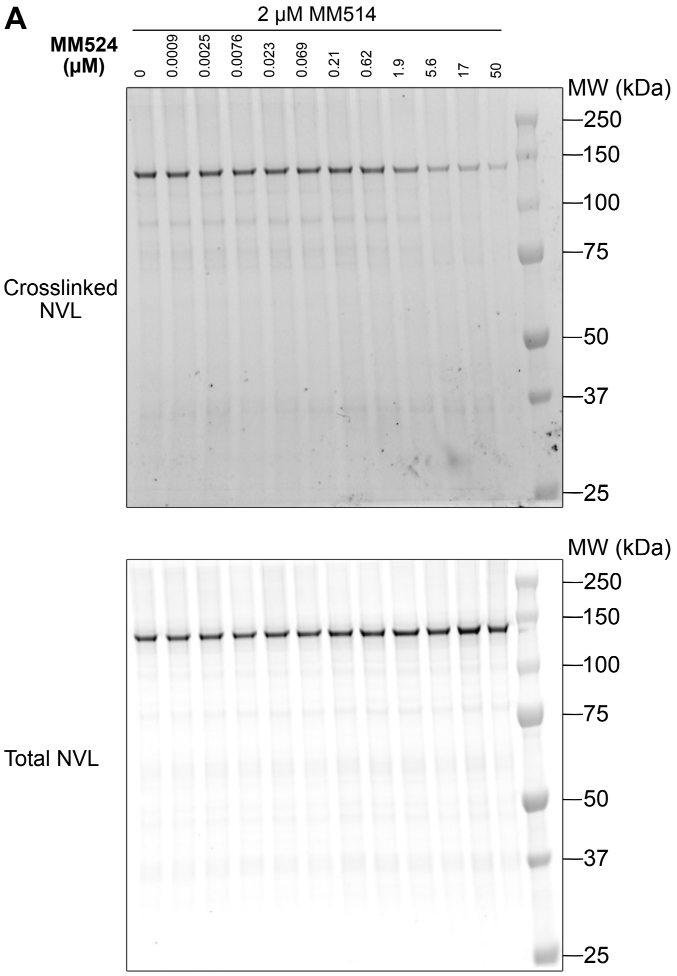


Note that the sample with 50 µM MM524 is not included in the figures or quantitation due to compound precipitation

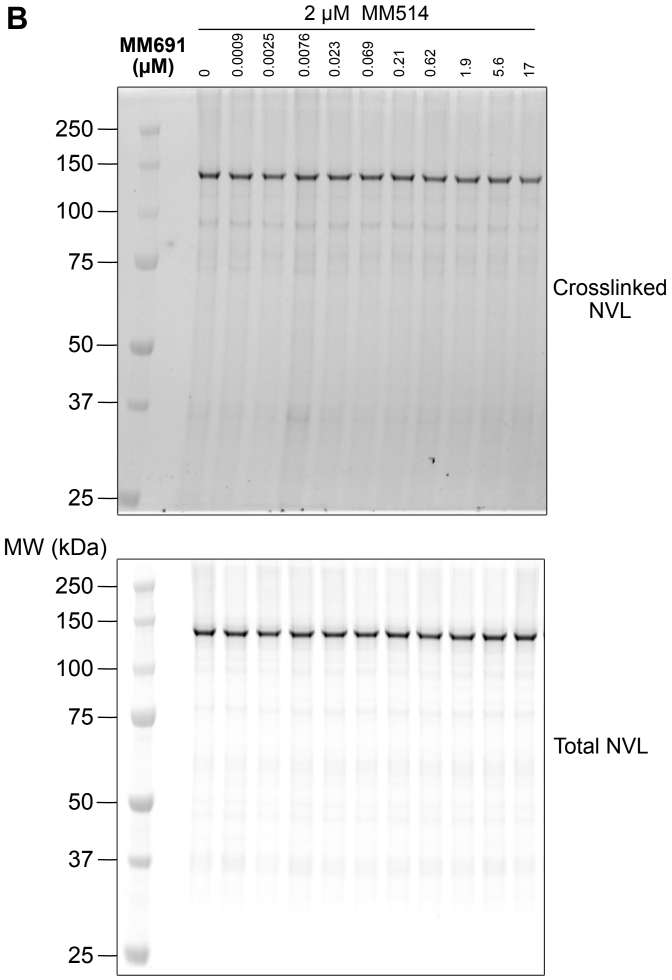

Full gel images shown in Fig. S2J (**A**) and Fig. S2K (**B**).

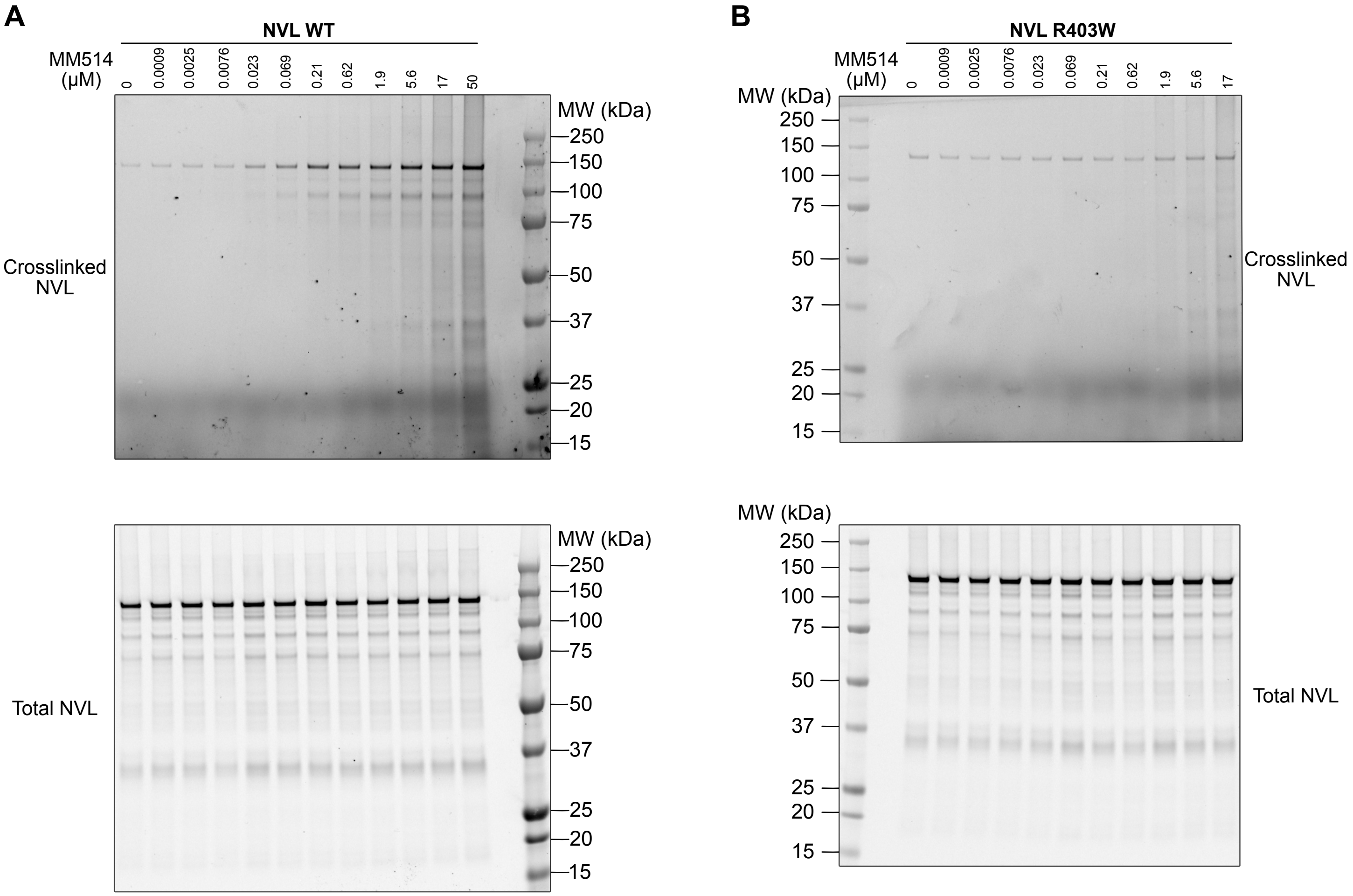


Note that the sample with 50 μM MM514 is not included in the figures or quantitation due to compound precipitation

Full Western blot images shown in Fig. S7C.
HRP chemiluminescence images are used in the figure panels. * non-specific band.

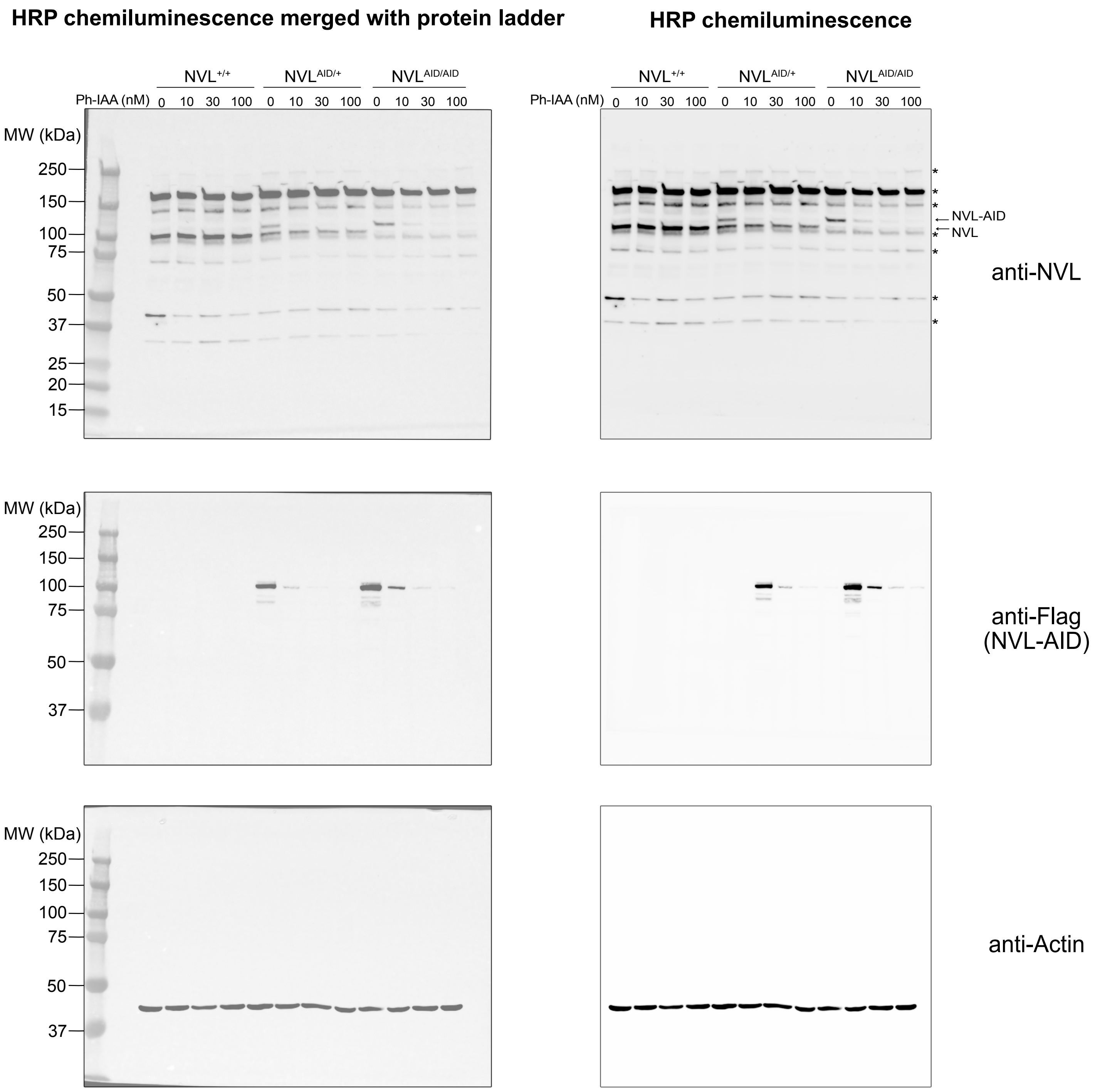

Full Western blot images shown in Fig. S7E.
HRP chemiluminescence images are used in the figure panels. * non-specific band.

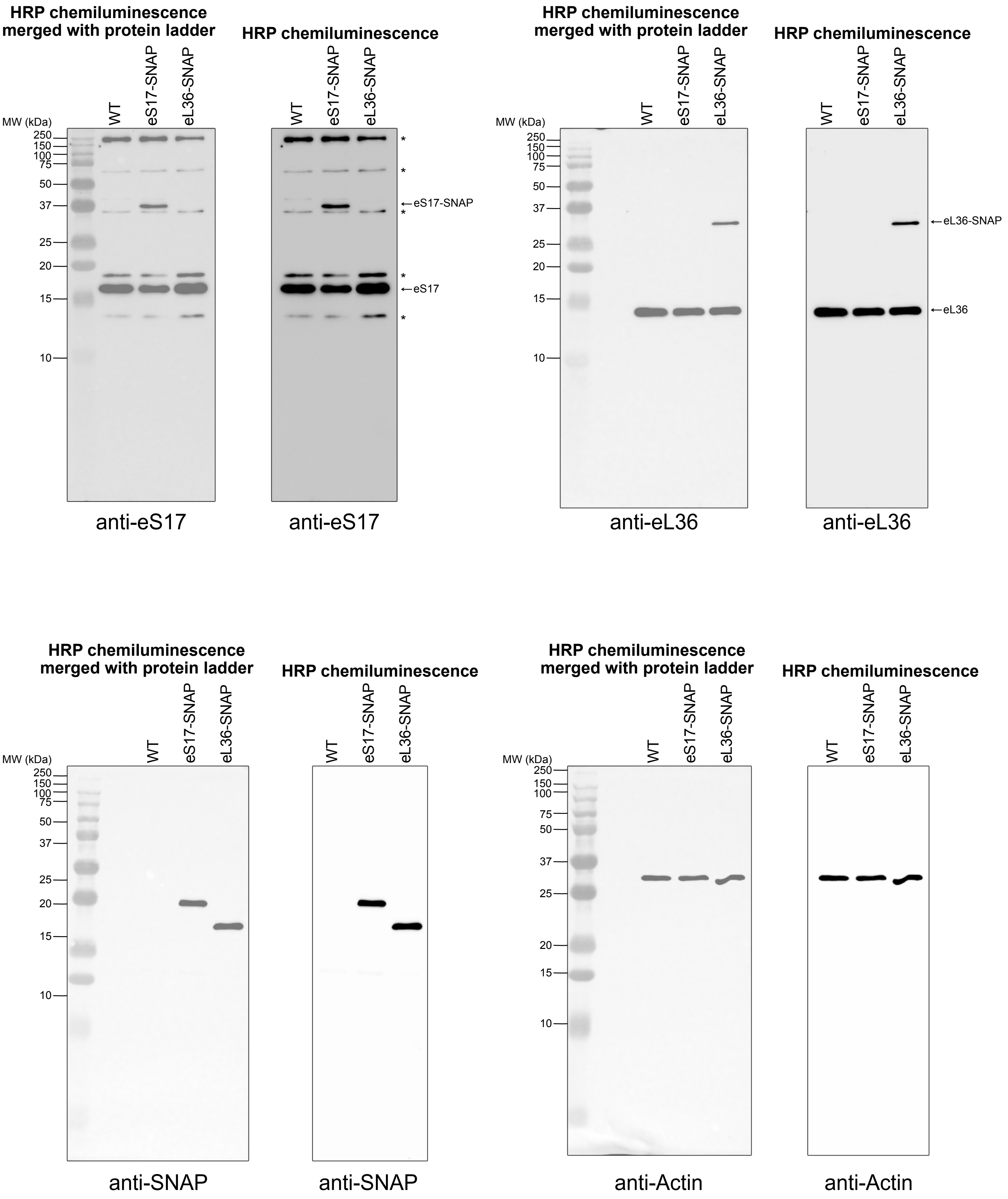

Full Western blot images shown in Fig. S7J.
HRP chemiluminescence images are used in the figure panels.

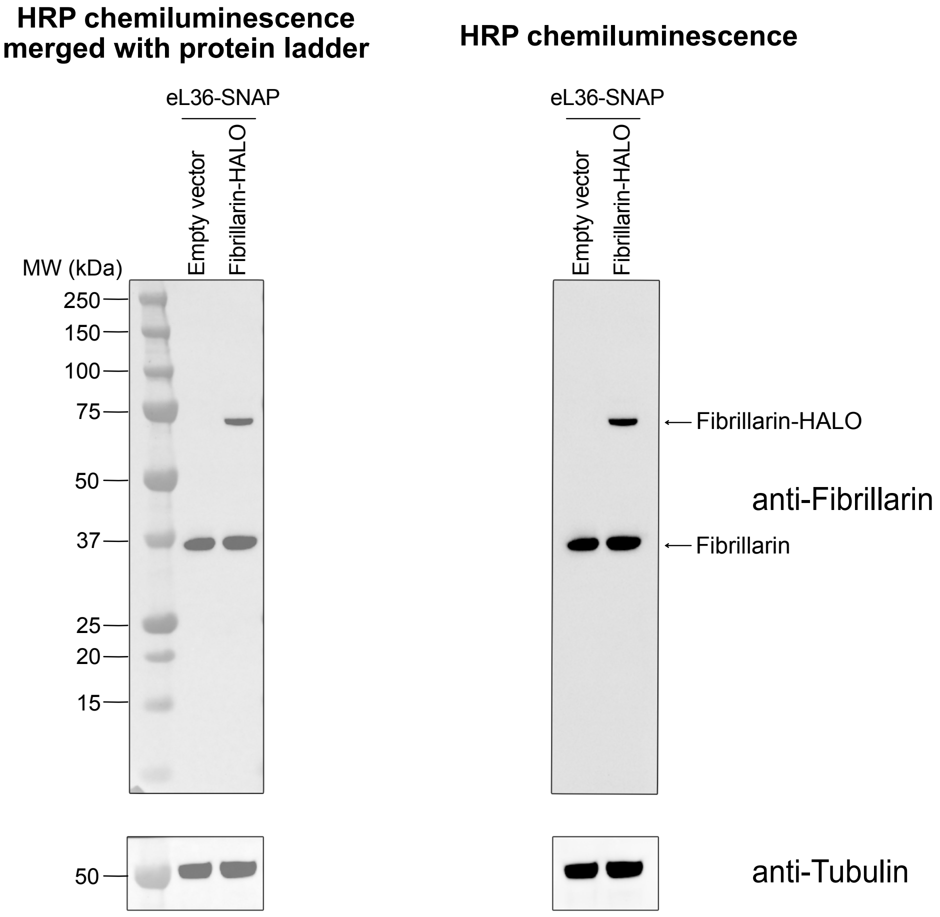


Membrane was first blotted and developed for anti-Fibrillarin,
then cut below the 75 kDa and above the 37 kDa ladder, and the middle membrane was
blotted and developed for anti-Tubulin

Full gel images shown in Fig. S7M.

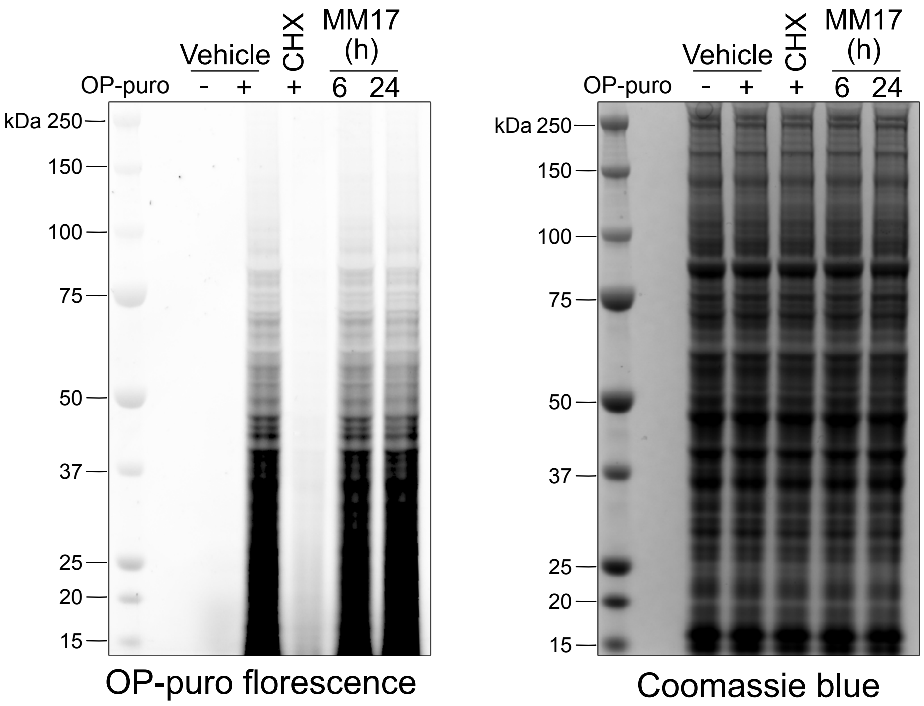

Full Western blot images shown in Fig. S8B.

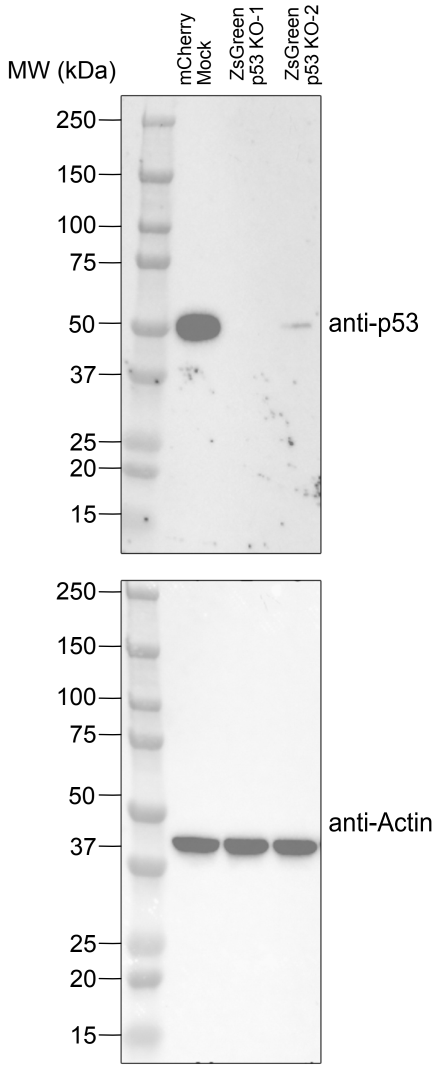

Full Western blot images shown in Fig. S8C.
HRP chemiluminescence images are used in the figure panels.

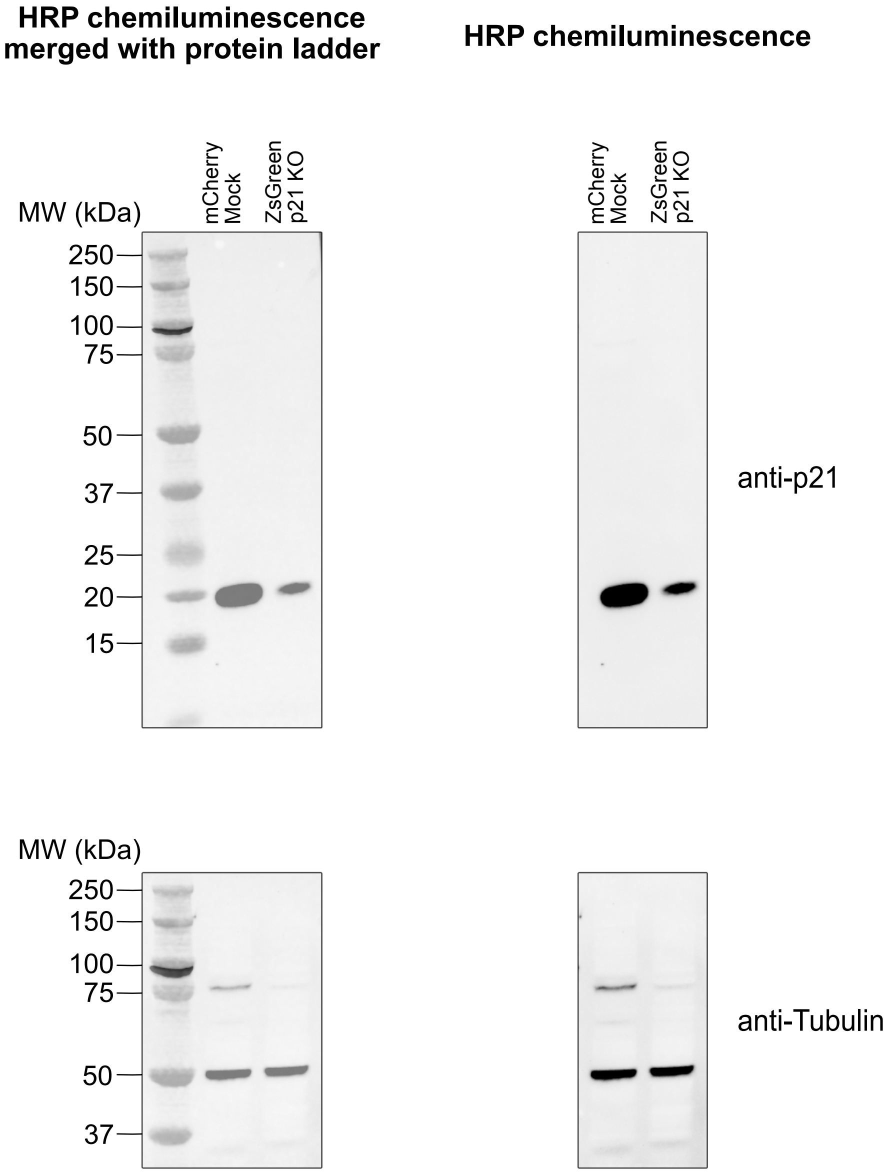


Membrane was first blotted and developed for anti-p21, then cut above the 25 kDa ladder, and top half was blotted and developed for anti-Tubulin.

Full Western blot images shown in Fig. S8G (**A**), Fig. S8I (**B**).
HRP chemiluminescence images are used in the figure panels.

**A**

**HRP chemiluminescence merged with protein ladder** **HRP chemiluminescence**

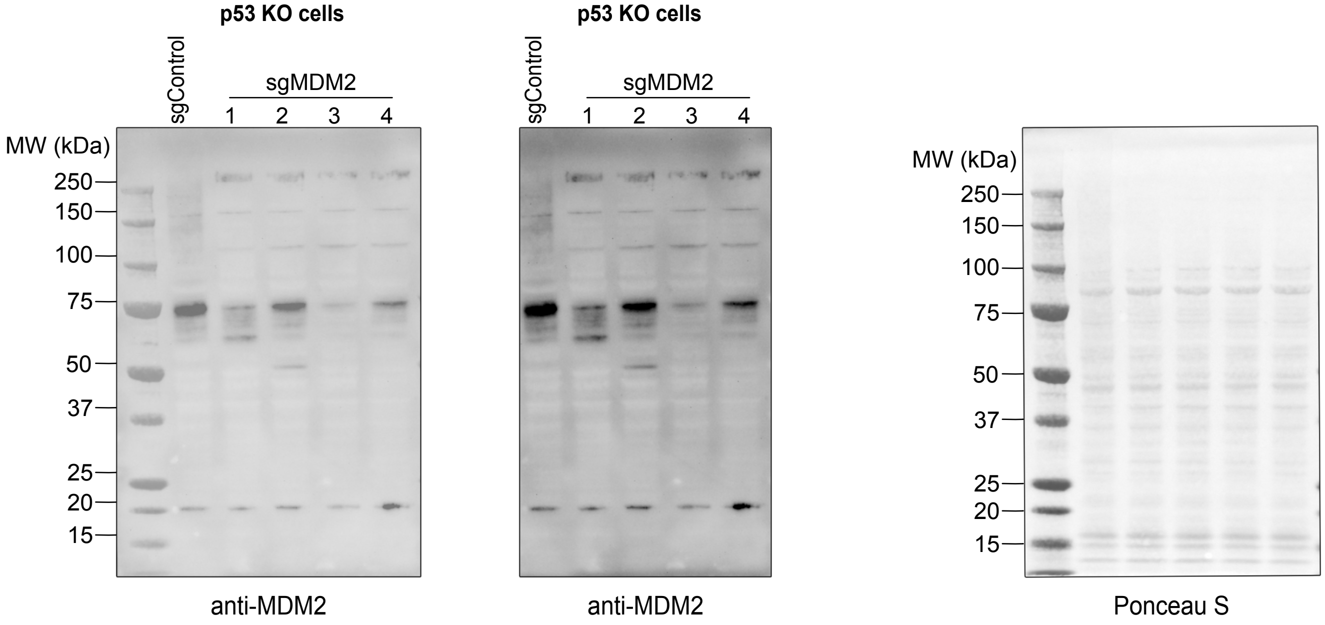


**B**

**HRP chemiluminescence merged with protein ladder** **HRP chemiluminescence**

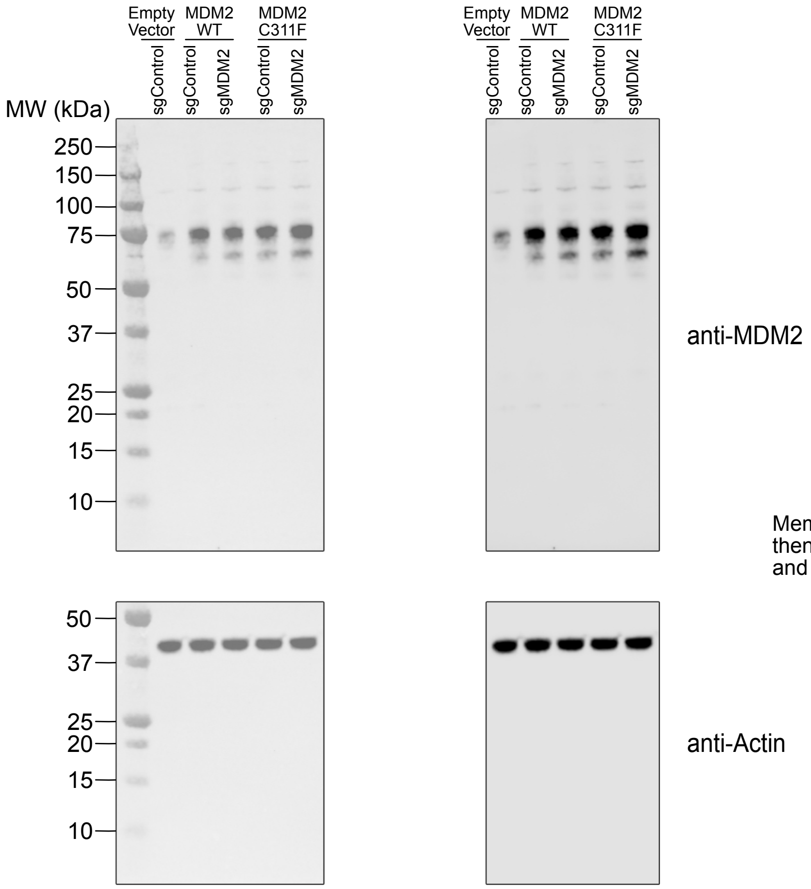


Membrane was first blotted and developed for anti-MDM2, then cut above the 50 kDa ladder, and bottom half was blotted and developed for anti-Actin.

Full Western blot images shown in Fig. S9A.
HRP chemiluminescence images are used in the figure panels.

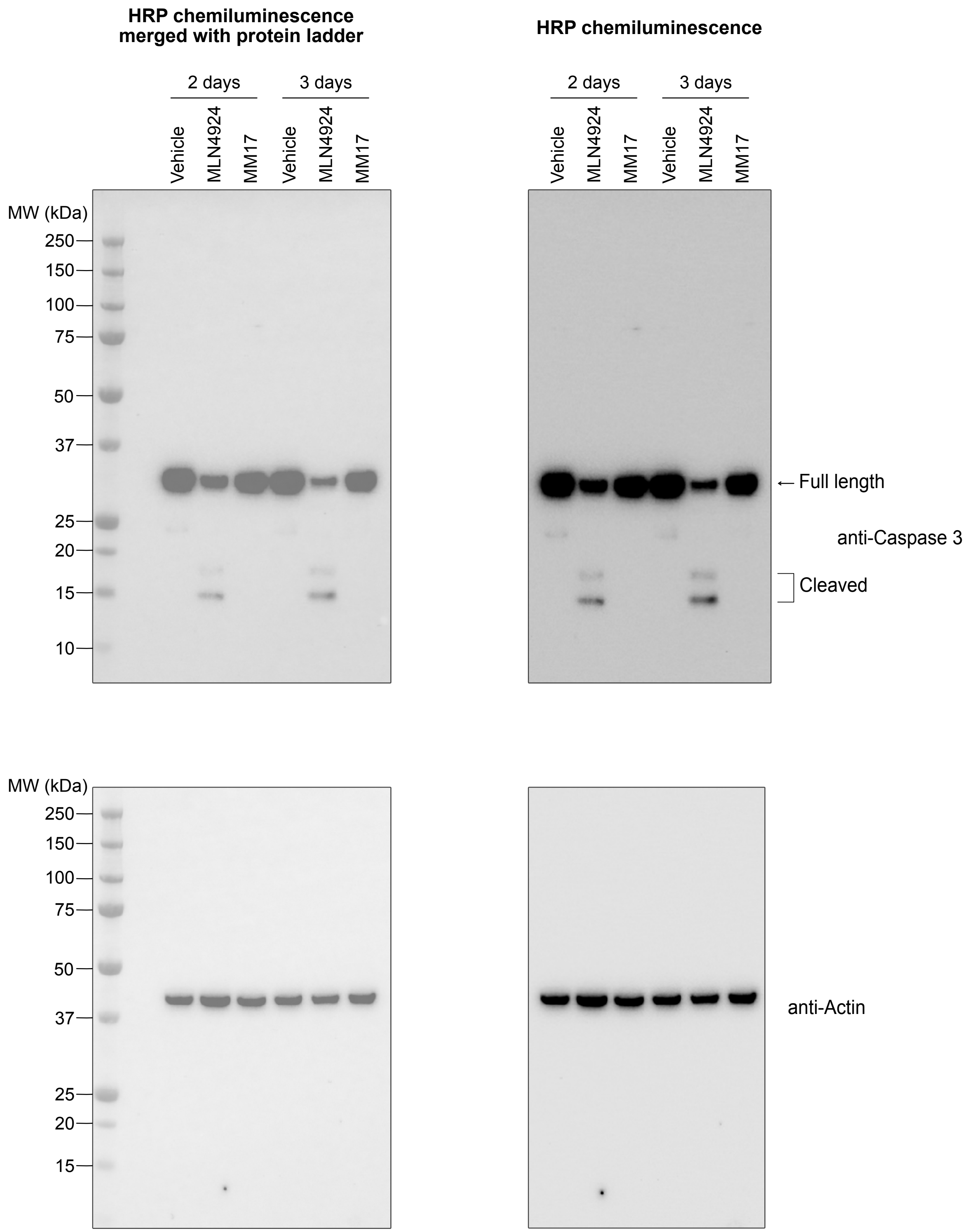

Full Western blot images shown in Fig. S9C.
HRP chemiluminescence images are used in the figure panels.

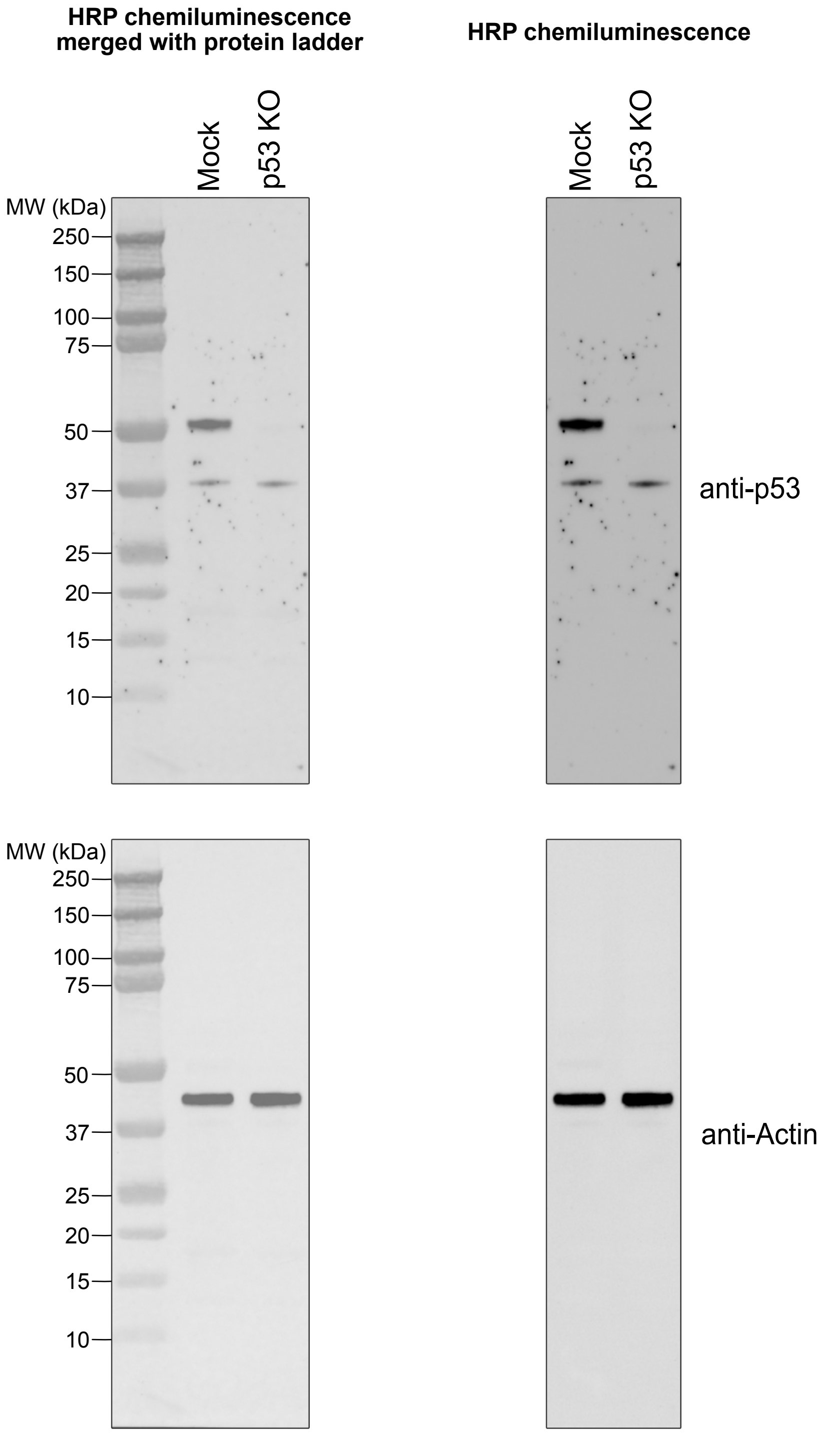

Full Western blot images shown in Fig. S14A.
HRP chemiluminescence images are used in the figure panels.

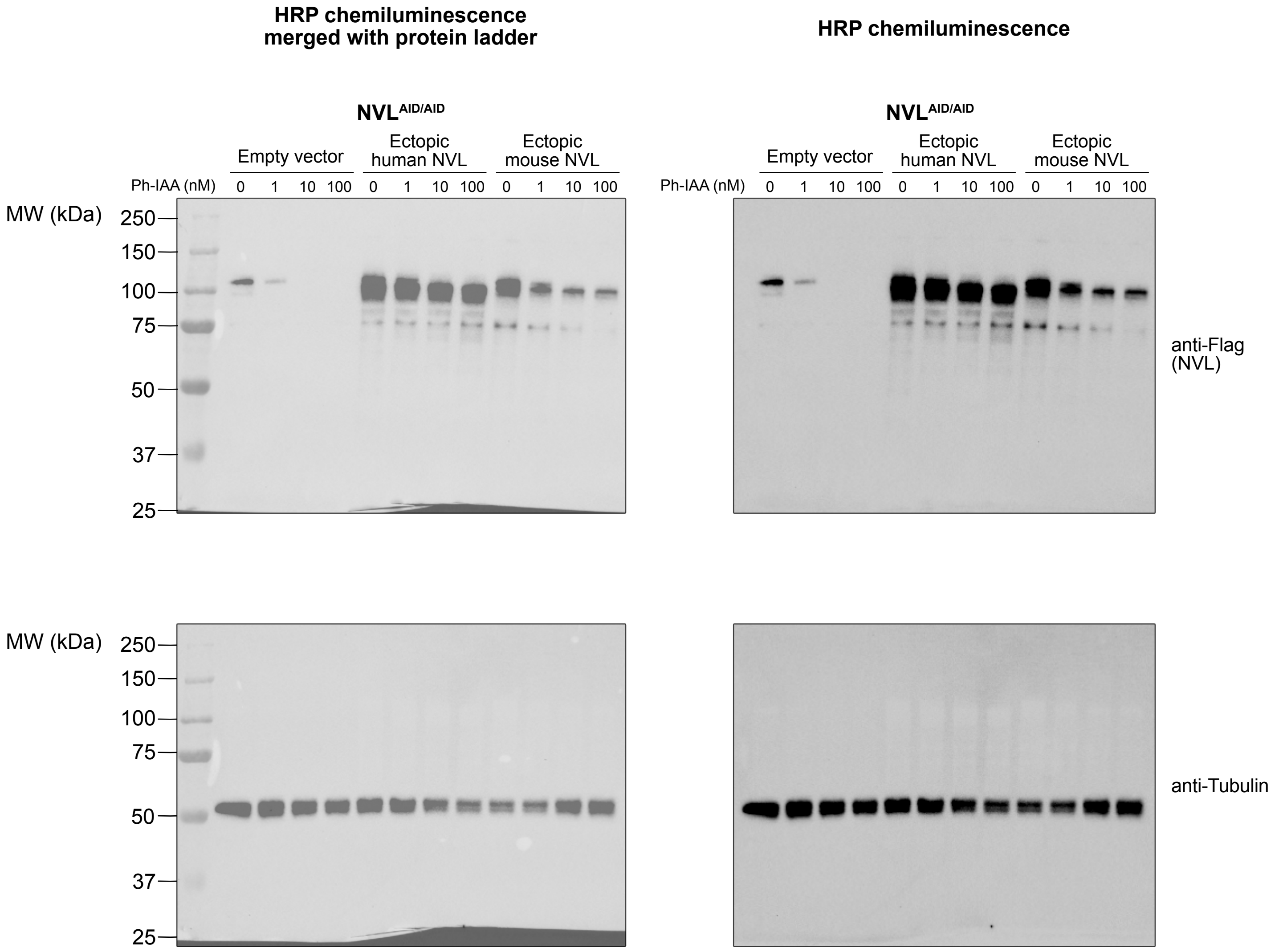

Full Western blot images shown in Fig. S14D (left).
HRP chemiluminescence images are used in the figure panels.

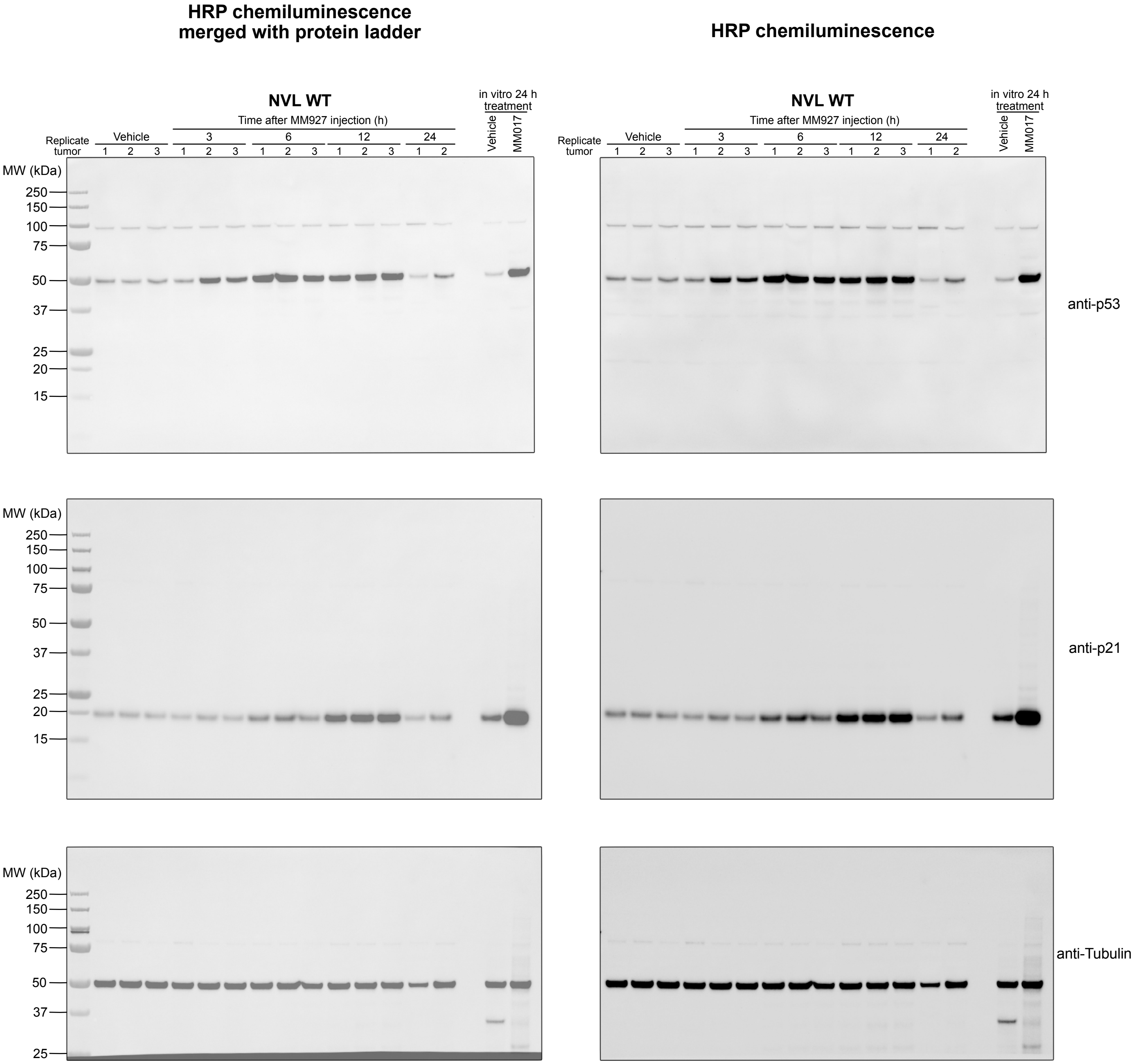


Membrane was first blotted and developed for anti-p21,
then cut below the 25 kDa ladder, and top half was blotted and
developed for anti-Tubulin.

Full Western blot images shown in Fig. S14D (right).
HRP chemiluminescence images are used in the figure panels.

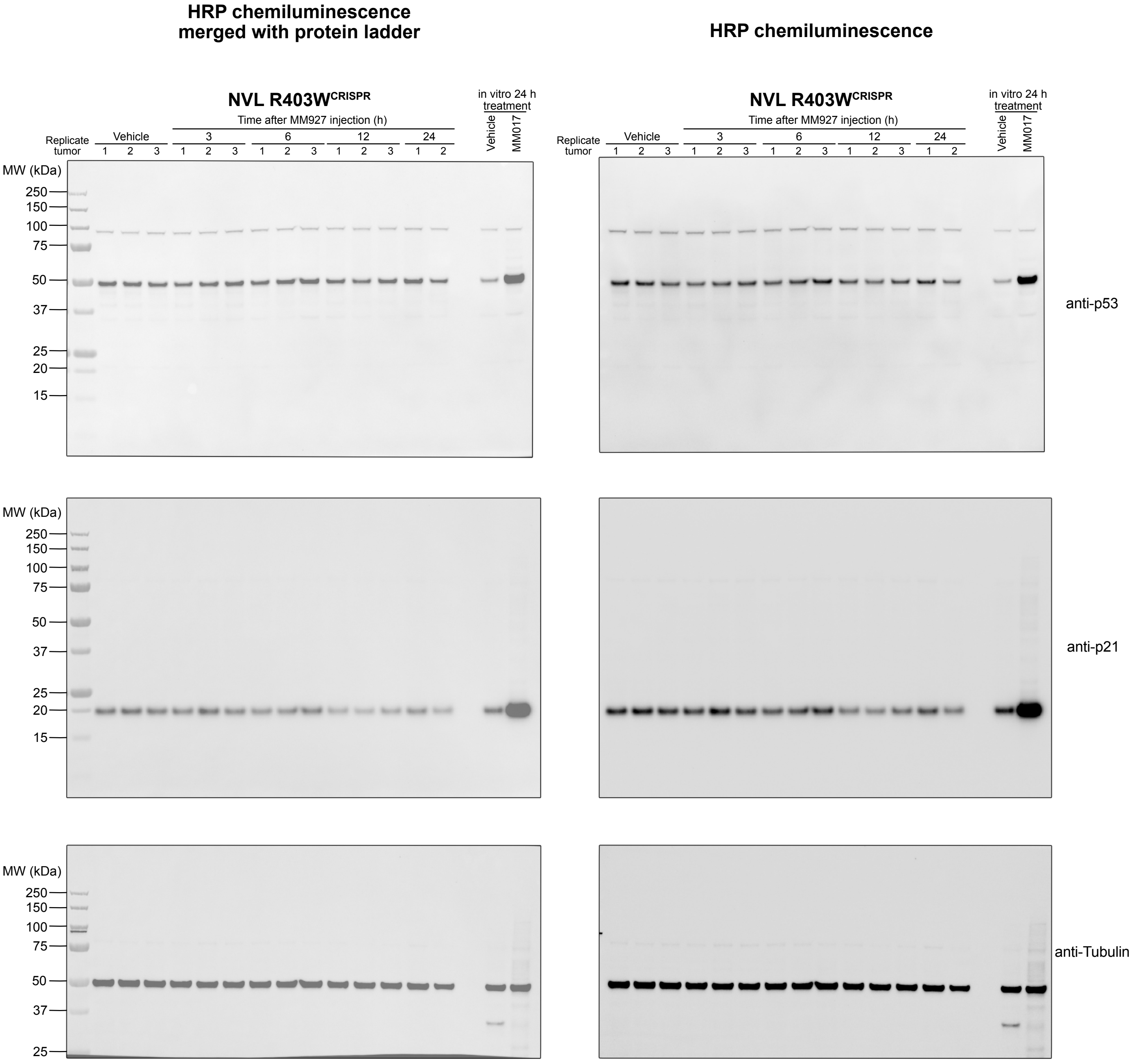


Membrane was first blotted and developed for anti-p21, then cut at the 25 kDa ladder, and top half was blotted and developed for anti-Tubulin.

Full Western blot images shown in Fig. S15A.
HRP chemiluminescence images are used in the figure panels.

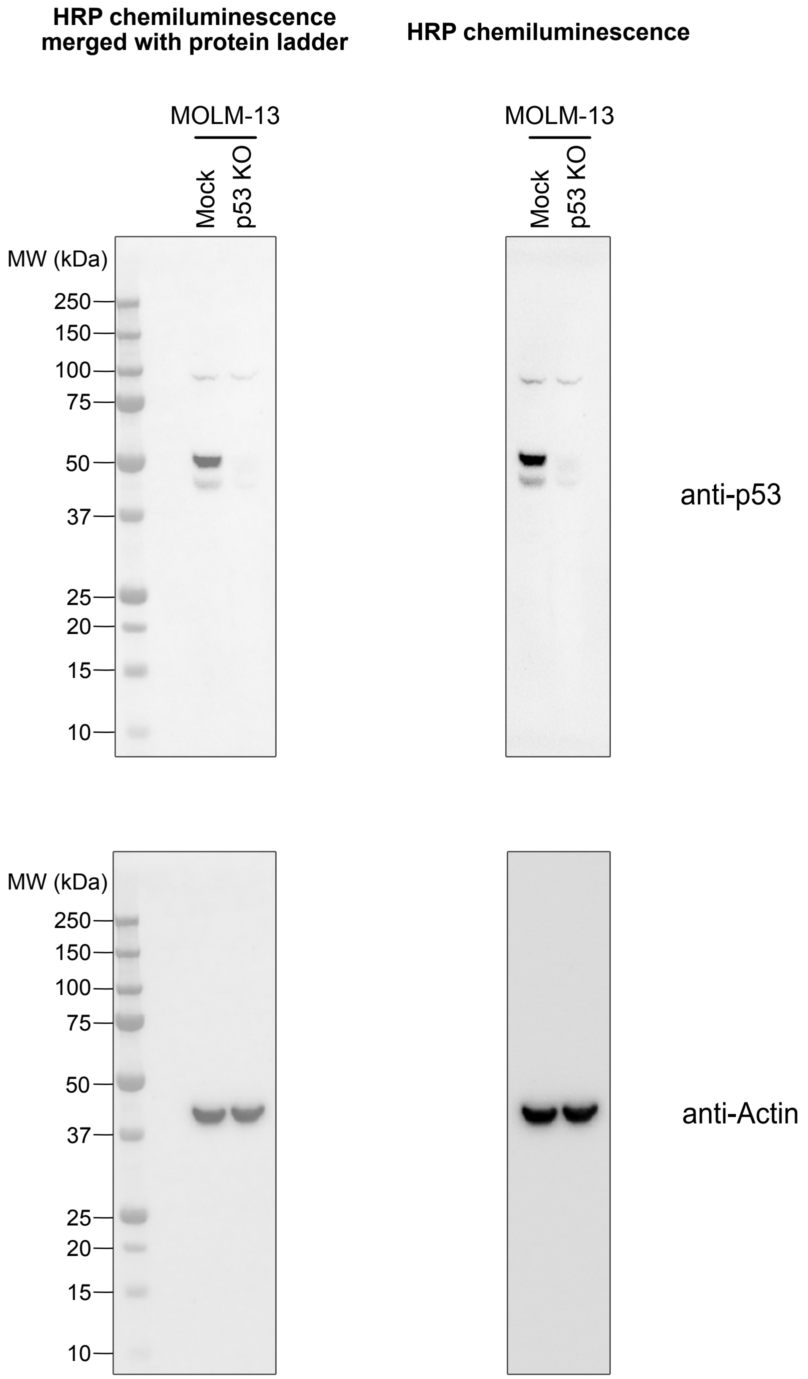

Full Western blot images shown in Fig. S15C.
HRP chemiluminescence images are used in the figure panels.

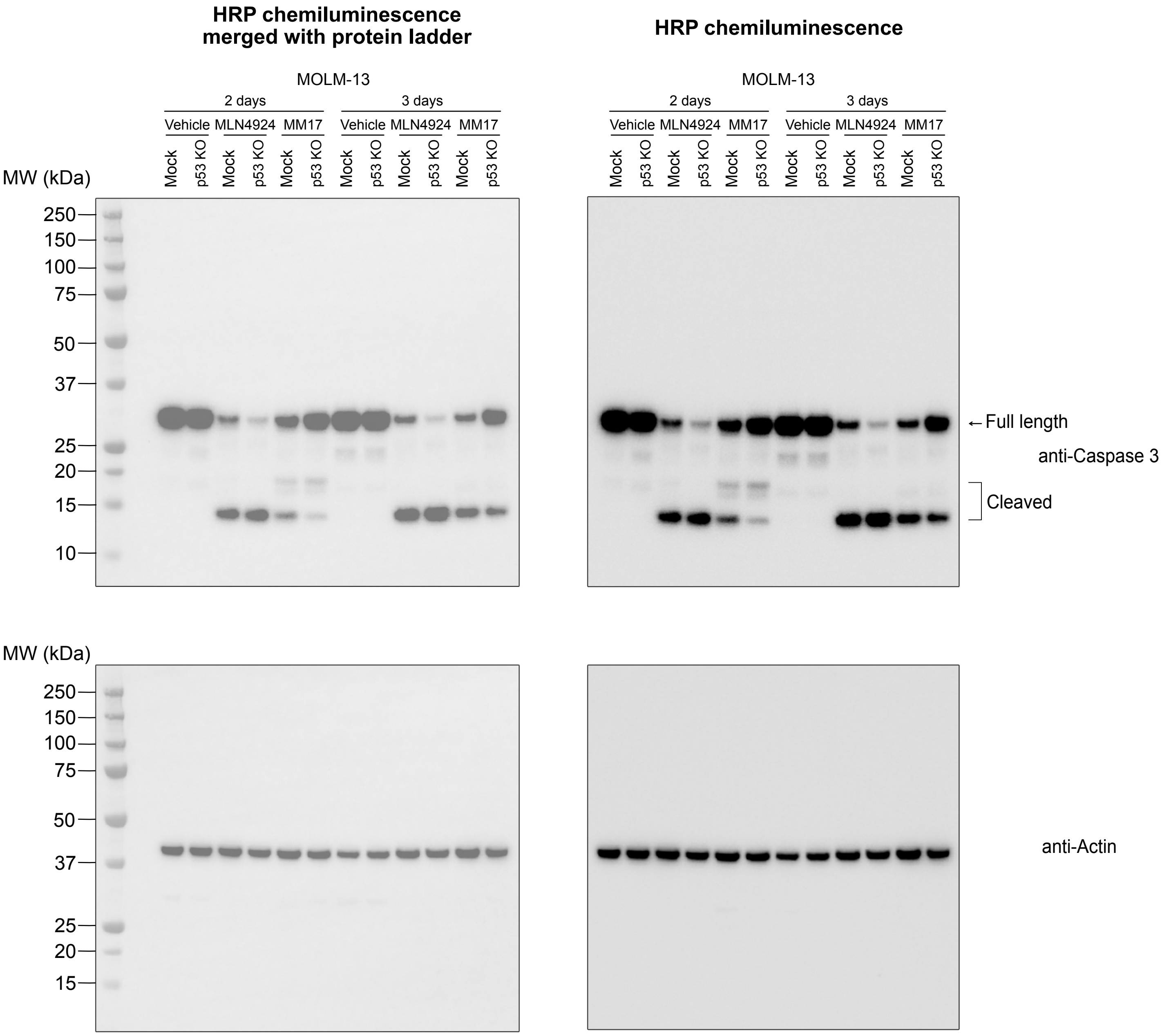

Full Western blot images shown in Fig. S15E.
HRP chemiluminescence images are used in the figure panels. * non-specific band.

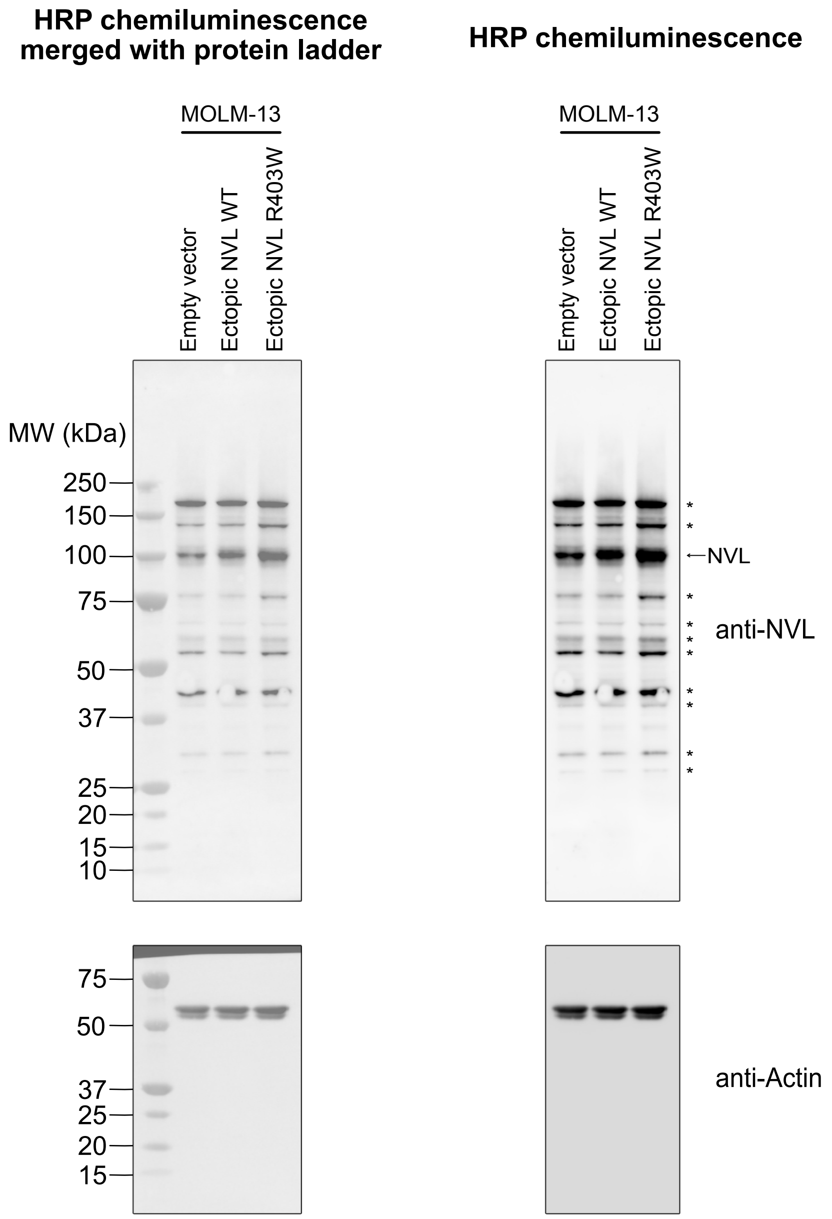
